# Logic-gating the HaloTag system with Conditional-Halo-ligator ‘CHalo’ reagents

**DOI:** 10.1101/2025.09.23.676741

**Authors:** Philipp Mauker, Carmen B.J. Zecha, Lucas Dessen-Weissenhorn, Luciano Román-Albasini, Joyce C.M. Meiring, Julia I. Brandmeier, Daniela Beckmann, Jan P. Prohaska, Martin Reynders, Yi Louise Li, Kristina Mrug, Hannah von Schwerin, Nynke A. Vepřek, Julia F. Gaisbauer, Leif Dehmelt, Perihan Nalbant, Jennifer Zenker, Anna Akhmanova, Martin Kerschensteiner, Angelika B. Harbauer, Julia Thorn-Seshold, Oliver Thorn-Seshold

**Affiliations:** Faculty of Chemistry and Food Chemistry, Dresden University of Technology, Dresden, DE; Department of Pharmacy, LMU Munich, Munich, DE; Max Planck Institute for Biological Intelligence; Martinsried, DE; Division of Cell Biology, Neurobiology and Biophysics, Department of Biology, Utrecht University, Utrecht, NL; Institute of Clinical Neuroimmunology, LMU University Hospital, LMU Munich, Munich, DE; Biomedical Center (BMC), Faculty of Medicine, LMU Munich, Martinsried, DE; Australian Regenerative Medicine Institute, Monash University, Clayton, Victoria, AU; Faculty of Biology, University of Duisburg-Essen, Essen, DE; Department of Chemistry and Chemical Biology, Technical University Dortmund, Dortmund, DE; Munich Cluster for Systems Neurology (SyNergy), Munich, DE; Technical University of Munich, Institute of Neuronal Cell Biology; Munich, DE; Institute for Clinical Chemistry, Medical Faculty, Dresden University of Technology, Dresden, DE

## Abstract

HaloTag proteins spontaneously ligate onto any chemical reagent featuring a chloroalkane motif (**CA**). We introduce the conditional **CHalo** motif, which ligates to HaloTag only after uncaging by light or enzymes. (1) Photo-triggered **CHalo** fluorogenic reagents allow spatiotemporally-specific labeling; (2) photo-triggered **CHalo** heterodimerisers can photocontrol protein recruitment; and (3) enzyme-triggered **CHalo** reagents can durably record diverse enzyme activities, and multiplexing them should allow quantitative ratiometric recording of multiple activities in parallel. **CHalo** thus permits manifold extensions to the HaloTag technology.

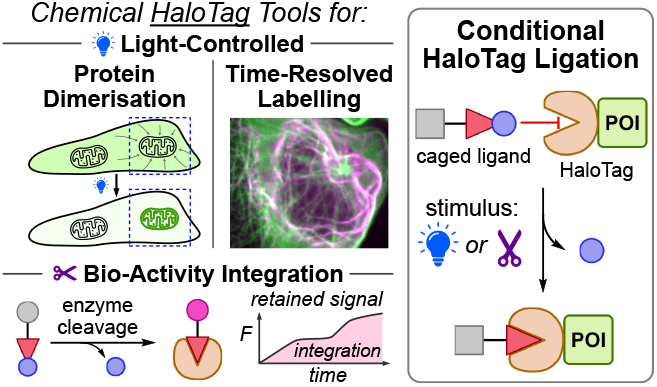

## Introduction (Fig. 1a-c)

HaloTag, SNAP-tag, and CLIP-Tag self-labelling proteins empower *chemical biology* by mediating the ligation of diverse *chemical* reagents to diverse *biological* targets (fusion proteins of interest, POIs). HaloTag is used throughout cell and *in vivo* biology,^1^ to visualise POIs with fluorogenic reagents (Halo-SiR), or to functionalise them as sensors (WHalo-CaMP), drug targets (T-REX, DART), or for degradation.^2^ Nonetheless, the utility of HaloTag was limited because it links the chemical and biological worlds *unconditionally*. Split-HaloTag sensors were recently engineered so their ligating reactivity is *conditional* on a biological event (calcium spike, GPCR-arrestin binding), making them “molecular activity recorder” proteins (CaProLa) that can reveal biology with unprecedented resolution.^3^

**Fig. 1.**
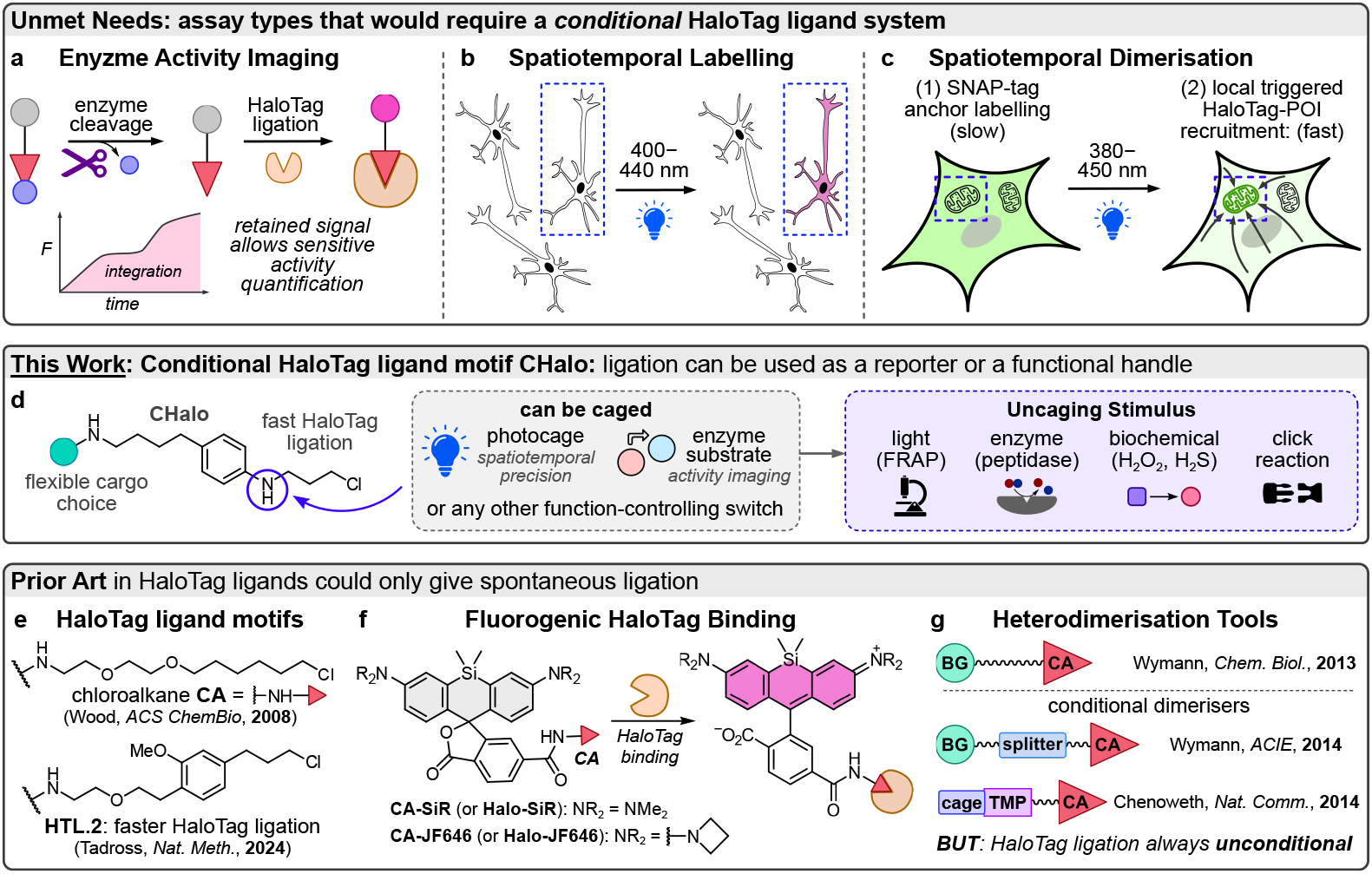
CHalo aims to unlock conceptually new chemigenetic assays for biology. **a-c**, Logic-gated HaloTag applications that would require a conditional HaloTag ligand. **d, CHalo** uses aniline uncaging to control its ligation to HaloTag. **e-g**, Prior HaloTag ligator motifs, popular reagents, and heterodimerisers.

A major cognate gap in the HaloTag toolbox remains: no *chemical motif* was known, that would ligate to HaloTag conditional on a (bio)chemical stimulus. Such a Conditional HaloTag Ligator motif could be immensely useful: e.g. with enzymatic activity as the stimulus, to durably record bioactivity with dyes (molecular imaging; **Fig. 1a**); or light as a spatiotemporally precise stimulus to track defined pools of POIs (photoactivated labelling; **Fig. 1b**) or trigger protein-protein interactions (photoactivated heterodimerisation; **Fig. 1c**). Compared to protein tools, it is much cheaper to engineer small molecules for new reactivities, faster to translate them between biological models, and it is feasible to apply several different chemical reagents in parallel to acquire multi-dimensional data (multiplexing). Conditional HaloTag Ligator reagents could also be applied directly into the thousands of established experimental HaloTag systems.^1^ We aimed to synthesise this useful, missing Conditional HaloTag Ligator modality (**Fig. 1d**).

### Development of the CHalo platform (Fig. 1d-g)

HaloTag ligates linear chain substrates like the “chloroalkane” motif (CA, **Fig. 1e-g**)^4^ that insert down a protein tunnel.^5^ We expected that adding a cage group on the chain would stop insertion: thus ligation would only be possible after a stimulus removes the cage. Since the ether chains of known HaloTag motifs like CA have no potential caging sites, we designed alternative motifs with anilines for flexible caging as stimulus-responsive carbamates (e.g. photocages, enzyme cages, bioorthogonal cages; **Supplementary Note 3, Fig. S9**). We first adapted Tadross’ methoxybenzene **HTL.2**^6^ (**Fig. 1e**) as isosteric **HL2**^**N**^ (cageable nitrogen 6 atoms from the chlorine; **Fig. S6**) which ligated very efficiently, but *N-*caging did not block ligation. After several redesign cycles we discovered **CHalo** (cageable nitrogen 3 atoms from chlorine, i.e. closest possible; **Fig. 1d**; **Supplementary Note 1**). Uncaged **CHalo** reagents can ligate very efficiently (∼10^6^ M^-1^s^-1^), only 10-fold slower than cognate CA reagents^5^ (**Fig. S10h−j**; **Supplementary Note 4**); while crucially, carbamate-**Caged-CHalo** did not ligate (<1% over 100 min; **Fig. S7cd**; **Supplementary Note 2**). **CHalo** thus fits the “off/ON” need for logic-gated HaloTag uses (**Fig. 1a–c**).

### Uncaging CHalo-SiR triggers strongly SiR-fluorogenic HaloTag ligation

Silicon-rhodamines (SiR: e.g. SiR-Halo/CA-SiR, Halo-JF646, etc; **Fig. 1f**) are valued for low-photodamage, far-red tracking of HaloTag proteins, with low background since they are fluorogenic upon HaloTag ligation.^2,7^ Ligated **CHalo-SiR** reached the same fluorescence brightness as ligated CA-SiR, though with even larger fluorogenicity (lower background; **Fig. S10bc, Supplementary Note 4**). Crucially, **caged-CHalo-SiR** reagents had as low background fluorescence as non-ligated CHalo-SiR (**Fig. S10fg**). Thus, **caged-CHalo-SiR** reagents should be highly fluorogenic upon uncaging then ligation, and should be bright enough for any settings that CA-SiR can be used in.

### CHalo reagents equip HaloTag to apply or record biological and biochemical stimuli

Using HaloTag as an “all-purpose anchor protein” to durably mark cells or proteins at a specific time or region, or upon specific stimuli, is attractive for tracing cell dynamics, trafficking, history, and fates (**Fig. 1a**). There are crucial differences between ligation prior to stimulus response (prior art), or ligation downstream of the stimulus (CHalo). Prior ligation uses HaloTag as a <u>single</u>-purpose anchor, focusing on its <u>fusion protein</u>: suiting e.g. high-resolution localisation (HaloTag-reactive caged-fluorophores like PaX^8^ or PA-JF549).

Downstream ligation instead focuses on the <u>uncaging step</u>: e.g. durably transducing or quantifying <u>stimuli</u>. We now showcase these new features with a set of CHalo reagents (**Fig. 2**). Conceptually, it should also allow “multipurpose” HaloTag assays: applying multiple CHalo reagents together, to record or transduce <u>multiple</u> stimuli in parallel, via one HaloTag type. These may be particularly attractive to quantitatively and ratiometrically record several enzyme activities at once (**Supplementary Note 5**).

**Fig. 2.**
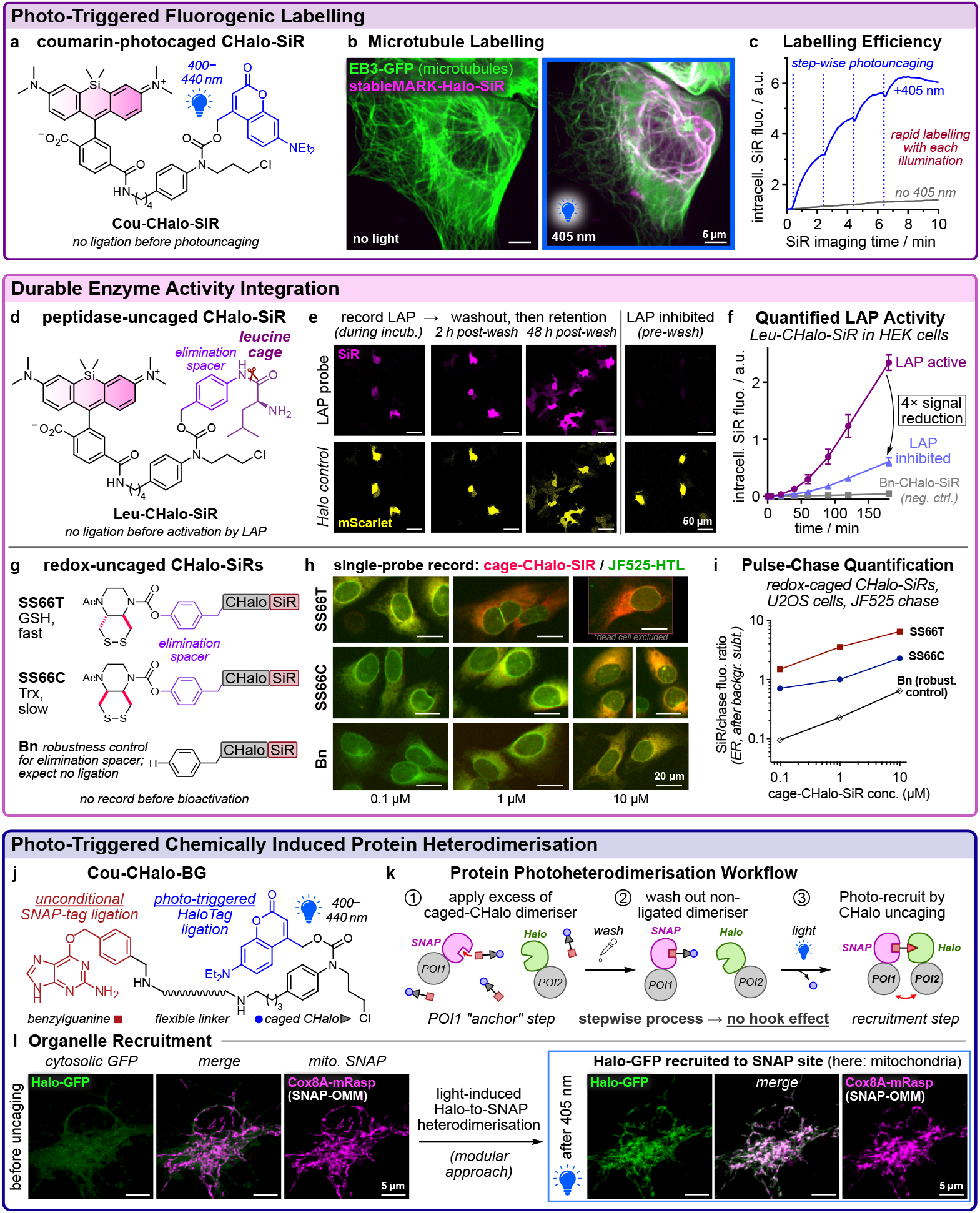
CHalo unlocks logic-gated HaloTag applications. **a-c**, Photoactivated label **Cou-CHalo-SiR**: temporally resolved photouncaging then HaloTag ligation activates red fluorescence (here: labelling StableMARK-Halo, a marker for stable microtubules, in U2OS cells; EB3-GFP overe xpressed to reveal the total microtubule network; 5 µM Cou-CHalo-SiR for 5 min; then 405 nm flashes; **Movie S1**). **d-f**, Model enzyme activity probe **Leu-CHalo-SiR**: durable imaging of leucine aminopeptidase activity (HEK cells with cytosolic HaloTag:mScarlet, 1 µM probe, 3 h; then 2× wash; optional pretreatment with inhibitor bestatin, 100 µM, 30 min). **g-i**, Bioreduction probes **SS66T-CHalo-SiR** (nonspecific, thiols) and **SS66C-CHalo-SiR** (thioredoxin) can be recorded and benchmarked by chase labelling to give coherent relative activities despite separate cell populations, over 100-fold differences of probe concentrations, and >20-fold different intrinsic activity levels (U2OS cells with HaloTag-Sec61 (ER-localised recorder) and GFP-NUP98 (nuclear marker, green), 0.1-10 µM SiR probes [red], 16 h; then chase with JF525-HTL [green], 1 µM). **j-l**, Photoactivated Halo-to-SNAP heterodimeriser **Cou-CHalo-BG**: modular, irreversible protein photoheterodimerisation (here: mouse primary neurons; mitochondrial SNAP-OMM anchor and mito-mRaspberry marker (magenta), cytosolic GFP-HaloTag target (green); SNAP labelled first (5 µM, 1 h) then 3× wash, 405 nm photoactivation, 5 min incubation, and re-imaged; **Movie S2**).

### Photocaged-CHalo-SiR is a fluorogenic label for HaloTag (Fig. 2a-c)

We created the coumarin-photocaged ligator-fluorogen reagent **Cou-CHalo-SiR** for spatiotemporally-photolocalised, durable cell marking (**Fig. 2a**). **Cou-CHalo-SiR** does not ligate HaloTag and is unaffected by GFP imaging (>470 nm); but violet light uncages it efficiently (ca. 50% conversion by 10 mJ/mm^2^ at 400-440 nm, **Fig. S10k**), enabling fluorogenic ligation (**Fig. S10l**). In cells, **Cou-CHalo-SiR** photolabels specific POIs, with temporal control (ligation half-times ca. 1 minute after uncaging, **Fig. 2c**) and minimal background (**Fig. 2b, Movie S1**: StableMARK-Halo, marker of stable microtubule subfraction; SiR-based background signal discussed at **Fig. S17**).

More generally, **photocaged-CHalo-Fluorogen** reagents will be high-sensitivity fluorogenic HaloTag labels that are compatible with multicolour as well as “multipurpose” multiplexed assays (**Supplementary Note 5**).

### Substrate-caged-CHalo-SiR probes are fluorogenic enzyme activity recorders (Fig. 2d-i)

There are no general platforms for sensitive fluorogenic chemical probes to allow *in vivo* enzyme activity integration and imaging, with durably quantitative cell-retained information, that do not impact native biology.^9^ Using enzyme activity to unleash CHalo ligation could leverage the diverse HaloTag systems and animal models already available,^1^ while harnessing the advantages of small molecule enzyme probes^10^ (e.g. unimpeded enzyme rates; **Supplementary Note 5**), but allowing clean persistent signal trapping.

Our test leucine-peptidase probe **Leu-CHalo-SiR** (**Fig. 2d**) gave excellent cellular signal retention over hours after washing (**Fig. 2e**), and plausible controls (signal low with peptidase inhibitor, no signal for no-peptide control **Bn-CHalo-SiR**; **Fig. 2ef**). This promises that CHalo can harness known substrate-mimic probe designs, including the use of chemical adapters to access diverse reactivities (**Fig. S9c**),^11^ but now converting them into durably trapped biologically innocent integrators: a meaningful step towards high-sensitivity enzyme imaging *in vivo*.

We next tested intracellular redox-activated fluorogenic probes, based on bioreductive unmasking of either the thioredoxin-selective cyclic disulfide **SS66C**^12,13^ or its essentially glutathione (GSH)-reporting diastereomer **SS66T**^12,13^ (**Fig. 2g**). As a step towards quantitative comparisons of recorded signals, we followed pulse labelling with these slow-turnover probes by chase labelling with the rapidly-labelling green fluorogenic ligand JF525-HTL; and we used image segmentation to focus on recorder regions (endoplasmic reticulum) while excluding non-ligated intracellular background (**Fig. 2h**). These data showed excellent linearity in the recorded signals over a ca. 100-fold concentration range (**Fig. 2i**), and were coherent with the specific activities expected (**SS66T**: high; **SS66C**: moderate; control **Bn** that can only be activated by unwanted hydrolysis: low). This suggests that CHalo-based recording may be able to reliably integrate and quantify even low enzymatic turnovers (low probe concentration and low activity), which has traditionally proved challenging for small molecule probes; and by recording a chemically different class of bioactivity, it underlines the modular applicability of the CHalo motif.

### Modular phototargeted SNAP/Halo heterodimerisation (Fig. 2j-l)

Chemically-induced protein (hetero)dimerisation (CID) is useful to probe interaction-dependent functions like receptor signaling, or single-protein functions dependent on localisation.^14^ SNAP-tag and HaloTag are proving valuable for covalent CID, complementing earlier noncovalent interaction domains (FKBP-FRB, eDHFR).^15^ Yet, because POI1-to-POI2 ligand designs suffer “hook effect” problems (incomplete dimerisation due to monovalent saturation of each POI), stepwise POI1-labelling / washout / POI2-labelling strategies are crucial.^16^ These require POI2-labelling to be unmasked upon external stimulus, typically photouncaging;^17^ but no photocaged^18^ SNAP-to-Halo designs are known.

CHalo solves the missing need for a modular, effective, photocaged SNAP-to-Halo double-covalent CID reagent. **Cou-CHalo-BG** performs the slower^5^ benzylguanine-SNAP-ligation first; then, after washout of unligated reagent, photouncaging allows HaloTag ligation (**Fig. 2j-k**). This allowed rapidly relocalising a cytosolic HaloTagged POI1 to a mitochondrial SNAP-tag in primary mouse neurons (**Fig. 2l, Fig. S26b**). **Cou-CHalo-BG** should be useful to photo-recruit *any* HaloTag protein to *any* SNAP-tag target; multiplexing analogues with spectrally distinct cages (**Fig. S9b**) could colour-code recruitment to various targets (e.g. Cage^2^-CHalo-TMP; Cage^3^-CHalo-CLIP); and recruiter-splitters^19^ could disconnect them on demand. The broad implementation of HaloTag tools may make such recruiters, which CHalo now unlocks, of particular practical impact.

## Conclusion

In conclusion, we developed the CHalo chemical motif which now allows diverse functional applications of the ubiquitous HaloTag protein. Recording enzyme activity, tracking protein pools, and recruiting proteins on demand, are three impactful examples of *chemigenetic logic-gating* that CHalo/HaloTag can deliver. In principle, almost any stimulus which cleaves a bond in a small molecule substrate can be used to activate the ligation; any HaloTagged protein can be targeted; and almost any chemical cargo can be delivered, including ligation-activated molecules (e.g. fluorogenic labels). We expect that CHalo can unite the broad availability of HaloTag models with the diversity of logic-gated chemistry to achieve substantial biological impact. We are particularly interested in multi-reagent/single-anchor assays, e.g. for “rainbow” ratiometric parallel recording with single cell resolution, since these should be able to test whether multiple enzymes’ activities are locked or are uncoupled (in physiological, or in challenged, stressed, or pathological conditions). Such quantitative multidimensional data should particularly impact systems biology, while more broadly empowering biochemistry and quantitative biology studies both *in vitro* and *in vivo*.

## Online content

All methods, additional references, reporting summaries, source data, extended data, supplementary information, acknowledgements, peer review information; details of author contributions and competing interests; and statements of data and code availability are available at *doi*.*org/xxxxxxxxx*. In brief, this includes the following:

## Supplementary Information

**Document S1**: Supplementary Notes 1–5, Figures S1–S26, Table S1, chemistry, materials and methods.

**Movie S1**: Timelapse imaging of photo-triggered fluorogenic labelling of stable microtubules with **Cou-CHalo-SiR**, corresponding to **Fig. 2b** (**MP4**)

**Movie S2**: Timelapse imaging of photo-triggered HaloTag-to-SNAP-tag recruitment with **Cou-CHalo-BG**, corresponding to **Fig. 2l** (**AVI**)

## Data and materials availability

All data needed to evaluate the conclusions in the paper are present in the paper and/or the Supplementary Materials, and are deposited and freely available on BioRxiv.

## Author Contributions Statement

P.M. performed synthesis, chemical analysis, photochemical evaluation, cell-free ligation experiments, analysed data and coordinated data assembly. J.T.-S. performed multiplexed cellular assays, and supervised L.D.-W. and C.Z. performing cell-free ligation experiments and quantification. J.P.P., M.R., J.I.B., H.S., and J.G. performed chemical synthesis. N.A.V. performed early cell-free ligation experiments. D.B. performed cell biology, confocal microscopy, image analysis and quantification for **Leu-CHalo-SiR**, supervised by M.K.. K.M. performed early live cell photo-heterodimerisation studies with **Cou-CHalo-BG**, supervised by P.N. and advised by L.D.. J.C.M.M. performed fluorogenic cell labelling with **Cou-CHalo-SiR**, supervised by A.A.. L.R.-A. performed live cell photo-heterodimerisation with **Cou-CHalo-BG**, supervised by A.B.H.. Y.L.L. performed mouse embryo live imaging with **Cou-CHalo-SiR**, supervised by J.Z.. O.T.-S. designed the study and the targets and supervised all other experiments. P.M. and O.T.-S. designed the experiments and co-wrote the manuscript with input from all authors.

## Acknowledgements

P.M. and J.T.-S. thank the Joachim Herz Foundation for fellowship support; P.M., and D.B. thank the Studienstiftung for fellowship support; J.Z. acknowledges a Sylvia & Charles Viertel Senior Medical Fellowship; and J.P.P. thanks the Fonds der chemischen Industrie for a Kekulé fellowship. We thank Alison Tebo (Janelia) for the advice about the role of the benzamide in the ligation step, which motivated us to keep developing reagents further despite having been disappointed in the performance of **HL2**^**N**^ and **HL3**; Luke Lavis (Janelia) for supportive discussions about the system concept; Julien Hiblot and Kai Johnsson (Max Planck Institute) for very kind gifts of **CA**-type reagents and HaloTag plasmids; Alf Honigmann (TU Dresden) for gifts of HaloTag plasmids; Gerti Beliu for kind gifts of HTL chase ligands and plasmids; Gerti Beliu and Made Budiarta (Uni Würzburg) and Ünal Coskun (TU Dresden) for producing HaloTag protein for cell-free assays; Dr. Inmaculada Segura (LMU) for her cloning expertise; the Imaging Facility at the Max Planck Institute for Biological Intelligence, Martinsried, Germany (RRID:SCR_026797) for facility access; and Thomas Misgeld (TUM) for supportive discussions. This work was supported by grants from the German Research Foundation (DFG: Emmy Noether grant 400324123 to O.T.-S.; SPP1926 grant 426018126 to O.T.-S., P.N., and L.D.; CRC1430 project A08 grant 424228829 to P.N. and K.M.; TRR353 ID 471011418 and SPP2453 ID 541742535 to A.B.H.; grant DE 823/10-1 to L.D.); the Max Planck Society (MPRGL to A.B.H.); the European Union (ERC StG Project 101077138 — MitoPIP to A.B.H.; Synergy grant PushingCell project number 101071793 to A.A.); the Munich Center for Systems Neurology (SyNergy EXC 2145; Project ID 390857198 to A.B.H. and M.K.); the Gravitation programme IMAGINE! (project number 24.005.009 to J.C.M.M and A.A.); and the NHMRC (APP2009409 to J.Z.). The Australian Regenerative Medicine Institute is supported by grants from the State Government of Victoria and the Australian Government.

## References

(1) Porzberg, N.; Gries, K.; Johnsson, K. Exploiting Covalent Chemical Labeling with Self-Labeling Proteins. Annu. Rev. Biochem. 2025, 94, 29–58. 10.1146/annurev-biochem-030222-121016.

(2) Deo, C.; Abdelfattah, A. S.; Bhargava, H. K.; Berro, A. J.; Falco, N.; Farrants, H.; Moeyaert, B.; Chupanova, M.; Lavis, L. D.; Schreiter, E. R. The HaloTag as a General Scaffold for Far-Red Tunable Chemigenetic Indicators. Nat. Chem. Biol. 2021, 17 (6), 718–723. 10.1038/s41589-021-00775-w.

(3) Huppertz, M.-C.; Wilhelm, J.; Grenier, V.; Schneider, M. W.; Falt, T.; Porzberg, N.; Hausmann, D.; Hoffmann, D. C.; Hai, L.; Tarnawski, M.; Pino, G.; Slanchev, K.; Kolb, I.; Acuna, C.; Fenk, L. M.; Baier, H.; Hiblot, J.; Johnsson, K. Recording Physiological History of Cells with Chemical Labeling. Science 2024, 383 (6685), 890–897. 10.1126/science.adg0812.

(4) Merrill, R. A.; Song, J.; Kephart, R. A.; Klomp, A. J.; Noack, C. E.; Strack, S. A Robust and Economical Pulse-Chase Protocol to Measure the Turnover of HaloTag Fusion Proteins. J. Biol. Chem. 2019, 294 (44), 16164–16171. 10.1074/jbc.RA119.010596.

(5) Wilhelm, J.; Kühn, S.; Tarnawski, M.; Gotthard, G.; Tünnermann, J.; Tänzer, T.; Karpenko, J.; Mertes, N.; Xue, L.; Uhrig, U.; Reinstein, J.; Hiblot, J.; Johnsson, K. Kinetic and Structural Characterization of the Self-Labeling Protein Tags HaloTag7, SNAP-Tag, and CLIP-Tag. Biochemistry 2021, 60 (33), 2560–2575. 10.1021/acs.biochem.1c00258.

(6) Shields, B. C.; Yan, H.; Lim, S. S. X.; Burwell, S. C. V.; Cammarata, C. M.; Fleming, E. A.; Yousefzadeh, S. A.; Goldenshtein, V. Z.; Kahuno, E. W.; Vagadia, P. P.; Loughran, M. H.; Zhiquan, L.; McDonnell, M. E.; Scalabrino, M. L.; Thapa, M.; Hawley, T. M.; Field, G. D.; Hull, C.; Schiltz, G. E.; Glickfeld, L. L.; Reitz, A. B.; Tadross, M. R. DART.2: Bidirectional Synaptic Pharmacology with Thousandfold Cellular Specificity. Nat. Methods 2024, 21 (7), 1288–1297. 10.1038/s41592-024-02292-9.

(7) Lukinavičius, G.; Umezawa, K.; Olivier, N.; Honigmann, A.; Yang, G.; Plass, T.; Mueller, V.; Reymond, L.; Corrêa Jr, I.R.; Luo, Z.-G.; Schultz, C.; Lemke, E. A.; Heppenstall, P.; Eggeling, C.; Manley, S.; Johnsson, K. A Near-Infrared Fluorophore for Live-Cell Super-Resolution Microscopy of Cellular Proteins. Nat. Chem. 2013, 5 (2), 132–139. 10.1038/nchem.1546.

(8) Lincoln, R.; Bossi, M. L.; Remmel, M.; D’Este, E.; Butkevich, A. N.; Hell, S. W. A General Design of Caging-Group-Free Photoactivatable Fluorophores for Live-Cell Nanoscopy. Nat. Chem. 2022, 14 (9), 1013–1020. 10.1038/s41557-022-00995-0.

(9) Mauker, P.; Dessen-Weissenhorn, L.; Zecha, C.; Vepřek, N.; Brandmeier, J. I.; Beckmann, D.; Kitowski, A.; Kernmayr, T.; Thorn-Seshold, J.; Kerschensteiner, M.; Thorn-Seshold, O. Cellularly-Retained Fluorogenic Probes for Sensitive Cell-Resolved Bioactivity Imaging. bioRxiv 2025. 10.1101/2025.04.17.649302.

(10) Chyan, W.; Raines, R. T. Enzyme-Activated Fluorogenic Probes for Live-Cell and in Vivo Imaging. ACS Chem. Biol. 2018, 13 (7), 1810–1823. 10.1021/acschembio.8b00371.

(11) Papot, S.; Tranoy, I.; Tillequin, F.; Florent, J.-C.; Gesson, J.-P. Design of Selectively Activated Anticancer Prodrugs: Elimination and Cyclization Strategies. Curr. Med. Chem. Anticancer Agents 2002, 2 (2), 155–185. 10.2174/1568011023354173.

(12) Zeisel, L.; Felber, J. G.; Scholzen, K. C.; Schmitt, C.; Wiegand, A. J.; Komissarov, L.; Arnér, E. S. J.; Thorn-Seshold, O. Piperazine-Fused Cyclic Disulfides Unlock High-Performance Bioreductive Probes of Thioredoxins and Bifunctional Reagents for Thiol Redox Biology. J. Am. Chem. Soc. 2024, 146 (8), 5204–5214. 10.1021/jacs.3c11153.

(13) Felber, J. G.; Kitowski, A.; Zeisel, L.; Maier, M. S.; Heise, C.; Thorn-Seshold, J.; Thorn-Seshold, O. Cyclic Dichalcogenides Extend the Reach of Bioreductive Prodrugs to Harness Thiol/Disulfide Oxidoreductases: Applications to Seco-Duocarmycins Targeting the Thioredoxin System. ACS Cent. Sci. 2023, 9 (4), 763– 776. 10.1021/acscentsci.2c01465.

(14) Broichhagen, J.; Trumpp, M.; Gatin-Fraudet, B.; Bruckmann, K.; Burdzinski, W.; Roßmann, K.; Levitz, J.; Knaus, P.; Jatzlau, J. Developing HaloTag and SNAP-Tag Chemical Inducers of Dimerization to Probe Receptor Oligomerization and Downstream Signaling. Angew. Chem. Int. Ed. 2025, 64 (35), e202506830. 10.1002/anie.202506830.

(15) Erhart, D.; Zimmermann, M.; Jacques, O.; Wittwer, M. B.; Ernst, B.; Constable, E.; Zvelebil, M.; Beaufils, F.; Wymann, M. P. Chemical Development of Intracellular Protein Heterodimerizers. Chem. Biol. 2013, 20 (4), 549–557. 10.1016/j.chembiol.2013.03.010.

(16) Ballister, E. R.; Aonbangkhen, C.; Mayo, A. M.; Lampson, M. A.; Chenoweth, D. M. Localized Light-Induced Protein Dimerization in Living Cells Using a Photocaged Dimerizer. Nat. Commun. 2014, 5 (1), 5475. 10.1038/ncomms6475.

(17) Chen, X.; Venkatachalapathy, M.; Kamps, D.; Weigel, S.; Kumar, R.; Orlich, M.; Garrecht, R.; Hirtz, M.; Niemeyer, C. M.; Wu, Y.-W.; Dehmelt, L. “Molecular Activity Painting”: Switch-like, Light-Controlled Perturbations inside Living Cells. Angew. Chem. Int. Ed. 2017, 56 (21), 5916–5920. 10.1002/anie.201611432.

(18) Caldwell, S. E.; Demyan, I. R.; Falcone, G. N.; Parikh, A.; Lohmueller, J.; Deiters, A. Conditional Control of Benzylguanine Reaction with the Self-Labeling SNAP-Tag Protein. Bioconjugate Chem. 2025, 36 (3), 540–548. 10.1021/acs.bioconjchem.5c00002.

(19) Zhang, H.; Aonbangkhen, C.; Tarasovetc, E. V.; Ballister, E. R.; Chenoweth, D. M.; Lampson, M. A. Optogenetic Control of Kinetochore Function. Nat. Chem. Biol. 2017, 13 (10), 1096–1101. 10.1038/nchembio.2456.

(20) Thorn-Seshold, O.; Mauker, P. Compound, Composition, Kit and Their Use for Protein Labelling. Patent Application 2025, LU602798.

