## Supplementary Information for "Logic-gating the HaloTag system with Conditional-Halo-ligator ‘CHalo’ reagents"

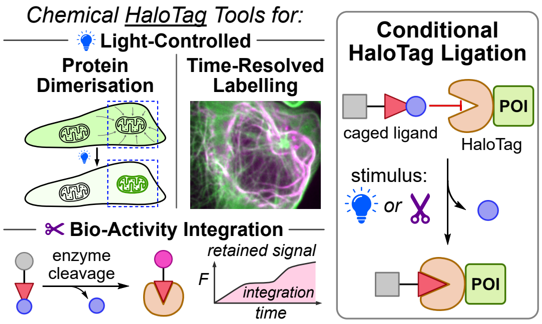

**For standard reagent use guides see Supplementary Note 1 (starting on page 3).** Particularly, with photocaged CHalo reagents (Cou-CHalo-SiR, Cou-CHalo-BG, etc) we strongly suggest experimenters to use red-only light sources for focusing on the microscope and for room lighting (not filtered white light bulbs).

8.5.3 Microscopy images for fluorescence quantification ... 27
8.6 Additional Data for Cou-CHalo-BG ... 29
**9 Methods: photochemical and biological characterisation ... 29**
9.1 Photochemical characterisation ... 29
9.2 Cell-free biological characterisation ... 30
9.3 Cellular characterisation ... 32
9.3.1 Cellular ligation rate ... 32
9.3.2 Spatiotemporally controlled cellular microtubule labelling: Cou-CHalo-SiR ... 33
9.3.3 Live cell imaging of enzyme activity: Leu-CHalo-SiR ... 34
9.3.4 Pulse-chase quantification of integrated enzyme activity: SS66T/C-CHalo-SiR ... 35
9.3.5 Spatiotemporally controlled protein heterodimerisation: Cou-CHalo-BG ... 35
**10 Synthetic Chemistry ... 36**
10.1 Chemistry methods and techniques ... 36
10.1.1 Analytical methods ... 36
10.1.2 Synthetic techniques ... 36
10.1.3 Chemicals ... 37
10.2 Synthetic procedures ... 37
10.2.1 General Procedures ... 37
10.2.2 CHalo Ligand building blocks ... 39
10.2.3 Fluorogenic CHalo-SiR conjugates ... 41
10.2.4 Light-controlled protein heterodimeriser Cou-CHalo-BG ... 48
10.2.5 Light-controlled & reporting protein heterodimeriser Cou-CHalo-SiR-BG ... 51
10.2.6 Proof-of-Concept and Design Optimisation ... 53
**11 References ... 69**
**12 NMR spectra ... 74**

### 1 User Guides for Photocaged CHalo Reagents (Fig S1-S3)

**We are happy to freely distribute Cou-CHalo reagents while our stocks last; we only ask receiving experimenters to send us their concentration and power integral settings / data, plus any other helpful assay or model parameters or identifiers, so that we can compile "typical benchmark settings" to recommend to the user community.**

#### 1.1 General handling

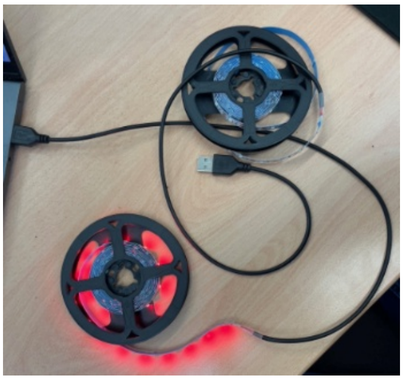

**Warning:** for photocaged CHalo reagents e.g. **Cou-CHalo-SiR**, **Cou-CHalo-BG**, etc, use **red-only** light sources for focusing and for transmission imaging on the microscope: since that light is focused through the objective, even minor wavelength components can reach high intensity and cause unexpected photouncaging. Red-only sources include red-emitting lasers (>630 nm) or red LEDs (typ. 630-660 nm) - these will always be good choices. Interference-filtered red-bandpass white sources *might* in principle be tolerated if the interference filtering is very effective (mileage may vary). In our experience, absorption-filtered white sources should always be avoided - such filtering is usually not >99.9% effective and the transmitted violet/blue light fraction then causes uncaging during setup that destroys reagent functionality.

**Lighting in the room and during handling:** Red-only LEDs for room lighting and experiment setup are very cheap; we use 3 euro red LED strips that plug into a USB socket with ca. 60 LEDs per strip (picture at right; each strip easily illuminates a workbench and/or hood; see e.g. this product link). Red-only room / handling lighting is not strictly required - as long as the light in the room / during handling is very dim (glow from computer monitor, etc), the amount of photouncaging it causes should be insignificant (note that an LED-based computer monitor can also be set to show pure red light). However, we are skeptical about "red light bulbs" that are actually absorption-filtered white light bulbs. Taken together we see no reason not to simply buy and use a USB-pluggable red LED strip for room lighting during handling.

**Storage**: To avoid the chemical degradation of reagent stocks during long-term storage, they should be aliquoted and then stored in the fridge/freezer (typically −20 °C; can depend on the solvent and reagent); wrapping the vials in aluminium foil will also prevent their photouncaging. For these irreversibly uncageable compounds, ideally, fresh aliquots should be used for each experiment, until the point where assays are confirmed to be working as expected and where satisfactory benchmark performance (for the compound in the assay settings) is available, so that any deviations from expected performance can be identified (although in our experience, assay problems rarely arise from stock degradation, but rather from on-stage mistakes).

**Treatment**: the reagents should always be added from 100X DMSO stocks (fresh dilution in two 10X steps on the experiment day), otherwise the reagents might precipitate from aqueous solutions at higher concentrations. If concentrations above 1 µM are used, experimenters should lock out for potential precipitation.

#### 1.2 Pre-tests: plausibility, working concentration, and uncaging settings

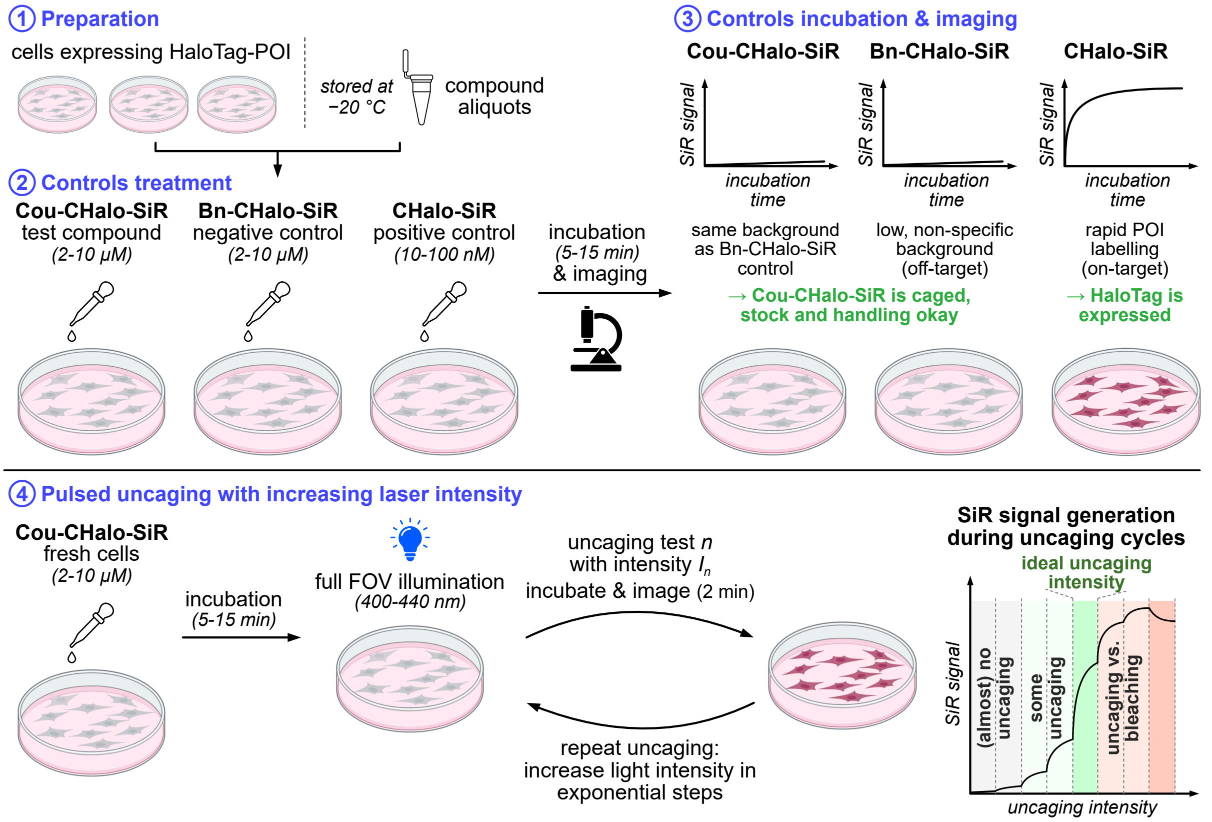

**Figure S1**: Experimental workflow to determine the ideal photouncaging settings using the **Cou-CHalo-SiR** fluorogenic dye with step-wise increase of the uncaging light "intensity" (in reality, the applied power integral, with units ~ $\mu J \cdot \mu m^{-2}$).

Determining the ideal photouncaging conditions with the microscope setup to be used in practice is crucial for straightforward and successful application of Cou-CHalo reagents. If too little light power is applied, local photouncaging will be insufficient; but too high light power will cause fluorophore bleaching, cellular phototoxicity, and can release too much CHalo species (i.e. allowing diffusion and out-of-target zone labelling) – finding the sweet spot is key. We recommend the following protocol to establish the uncaging conditions.

(A) Though non-ligated CHalo-SiR has zero fluorescence in cell-free buffer, subcellular compartmentalisation of any SiR reagents into mitochondria and ER gives non-specific background signals in these compartments (also see **Fig. S17**). The caged but non-activatable **Bn-CHalo-SiR** should be used as a control to report on this (expectedly minimal) non-specific "**compartment background**" signal (concentration-dependent) from CHalo-SiR and caged-CHalo-SiR reagents. Non-CHalo SiR reagents (e.g. SiR-Halo, CA-SiR, etc) can have much higher intensities of compartment background signal, and may also have slightly different localisation patterns, at the same concentration.

(B) If **Cou-CHalo-SiR** (at the same concentration) gives higher cellular signal in HaloTag-expressing cells *before uncaging* than **Bn-CHalo-SiR**, the **Cou-CHalo-SiR** must have partially degraded by losing the Cou cage, allowing it to ligate to HaloTag and generate strong ligated signal. In our experience, typically such unintentional photouncaging will have occurred on the stage during the microscopy experiment, e.g. from "white" light being focused through the objective (e.g. for a transmission image or for focussing), or from applying merely "filtered" red light where the filtering does not remove all UV/violet/blue light (instead of applying "red source" light from LEDs or lasers). In rare cases, photouncaging seems to have occurred by careless reagent stock and/or cell handling before reaching the stage (fix: handle stocks and cells under red source lighting and protect stocks from UV/violet/blue light exposure, e.g. wrapping vials in aluminium foil and storing them in the fridge/freezer).

(C) **CHalo-SiR** should be used as a positive control to find the maximum expected target-specific fluorescence, by titrating its concentration up (while comparing to the Bn-CHalo-SiR compartment background signal at the same concentration). It is also a good control for successful HaloTag-POI expression. **CHalo-SiR** is the photouncaging product from **Cou-CHalo-SiR** illumination. When **CHalo-SiR** is applied directly to cells, it ligates to HaloTag upon treatment; typically at low concentrations it will mainly give signal from the ligated product (signal/concentration gradient is high), while at intermediate concentrations all HaloTag protein will become saturated, such that at high concentrations its signal/concentration gradient matches that for the compartment background (low). In typical assays we aim to locally photogenerate an "acceptable" uncaged concentration of **CHalo-SiR** that may be in the 10-100 nM range (since photouncaging proceeds to the same percentage independent of the reagent concentration, we typically aim to minimise photodamge in these assays by applying a rather high **Cou-CHalo-SiR** concentration and only uncaging a small percentage of it). Thus we recommend applying the **CHalo-SiR** control at concentrations from 10−100 nM, typically by using 100−500X DMSO stocks (maximum DMSO concentration in cell medium: 1%).

***Experimental steps (Fig. S1)***:

**Pre-tests**: (1) Cells expressing the HaloTag-POI fusion protein and fresh stocks of **Cou-CHalo-SiR**, **Bn-CHalo-SiR** and **CHalo-SiR** are prepared. (2) The cells are treated with the respective compound (2−10 µM) and incubated for 5−15 min (*Note: all three dyes are well cell permeable allowing for short incubation times; higher concentrations of 2−10 µM should be used for* ***Cou-CHalo-SiR*** *to achieve rather high local concentrations since the photouncaging product is likewise permeable and will diffuse away from the activation site*). (3) The cells are imaged in the SiR channel (excitation: 650 nm, emission: 670 nm) during the incubation to determine the SiR signal increase. *Expected results*: **Bn-CHalo-SiR** and **Cou-CHalo-SiR** should give very low signal, that is approximately equal (minor non-specific "compartment background" signal in mitochondria and ER is expected, also see **Fig. S17**). If **Cou-CHalo-SiR** shows higher fluorescence than **Bn-CHalo-SiR**, typically the handling off- or on-stage has led to (partial) unwanted photouncaging of **Cou-CHalo-SiR**, so handling should be improved but reagent stocks are still fine (less likely: the stock used has been degraded and significant **CHalo-SiR** is now present inside it). **CHalo-SiR** should rapidly develop on-target fluorescence by ligating to HaloTag in the compartment where it is expressed (note though: signal does not *necessarily* reach plateau during the first 15 min at low concentrations).

**Parameter tuning**: If the pre-tests gave the expected results, experimenters can move forward to set experimental parameters for pulsed photouncaging assays. (4) A fresh well of HaloTag-POI expressing cells is treated with **Cou-CHalo-SiR** (typ. 10 µM as a start point*) for 5−15 min and then the whole field of view (FOV) is photouncaged step-wise several times (400−440 nm light) with ***significantly**** increasing light energies at each step, imaging the SiR channel over ca. 2 min after each uncaging step (e.g. taking 4-10 images). ***Assuming that the concentration has been well chosen****, a good uncaging energy is found by identifying which step gave the steepest specific SiR signal increase.

An expected **photouncaging** profile is shown in **Fig. S1**. Note that e.g. an uncaging light power integral of "0.1" will result in 10% photouncaging percentage that may give already e.g. 50% of maximum specific Halo-ligated signal (if Cou-CHalo-SiR concentration is 5 times higher than HaloTag concentration); but an uncaging light power integral of "1" that will result in ca. 66% uncaging (logarithmic response) but this only gives e.g. 100% of maximum specific signal (HaloTag is saturated), and any further increases in uncaging light power cannot increase the specific signal. In fact, **photobleaching** of the Halo-ligated SiR dye by the uncaging light always accompanies each uncaging step, so the total signal (at high Halo-ligation) can actually reduce with additional uncaging light.

*** Concentration:** A good control is to compare results with e.g. 3 µM and 30 µM of the Cou-CHalo-SiR reagent. The same photouncaging profile as for 10 µM should give nonlinearly scaled, but plausible, SiR signal profiles (e.g. with 30 µM, 3 fold lower uncaging light integral should be required to saturate the HaloTag and give maximum specific signal; with 3 µM, 3 fold higher light integral is required). If the concentration-comparison assays give plausible results then the 10 µM working concentration can be considered well-chosen. If not, adjust the concentration and run again.

**** Light Energy**: Ideally, the uncaging light power integrals should be scanned by logarithmic steps (1,2,4,8,16,...). Adjusting the laser intensity over a reproducible region (e.g. 2, 4, 8, 16, 32%) may be the most practical way to identify roughly the right power integral. However, ***once that power setting is found***, we strongly recommend to run a control that scans a smaller range of light power integrals by ***linearly*** adjusting the pixel dwell time over perhaps a 4-fold range, since that is an easy and reproducible way to linearly vary the applied power integral, without changing its spatial distribution, with good reproducibility. Since a "good" uncaging energy is by definition one that uncages a rather small amount of reagent (see point (C) above), that *linear* pixel dwell time scan series should be enough to deliver clear and ***linear*** **responses in HaloTag-ligated-SiR readout**, despite the complexities of e.g. photobleaching. If that linearity is not seen, saturation or thresholding effects must have been operating: thus the power integral would usually have been set vastly too high or too low, and severe photobleaching or absolute signal intensity problems may have been seen. **In such cases, the first step is to re-evaluate the working concentration**, and re-start the tuning procedure. *Nonlinearity can be a particular problem when overall HaloTag expression per cell is very low: e.g. it is easily possible that the step-wise illumination leads to HaloTag saturation after initial steps, preventing possible specific signal increase of higher light dose steps, while photobleaching starts to dominate. In that case, the photouncaging test should be repeated with a fresh well or fresh plate of cells (to avoid cases where unwanted light scattering may have photouncaged some reagent in neighbouring cells and wells), and starting from perhaps 5% of the uncaging energy that gave specific signal saturation.*

**Finally**: When the ideal light power integral (i.e. roughly the dwell time per pixel (ms) multiplied by energy density (W/µm$^2$ ***in the sample plane***)) has been identified, the experimenters should record the actual laser power for reference purposes (since the laser power can vary over time, this will allow reproducibly adjusting laser intensity for comparable future experiments). Please also send us the concentration / power integral settings that work for you, so we can add them to our **typical benchmark settings** compilation.

#### 1.3 Light-controlled fluorogenic labelling with Cou-CHalo-SiR (typical steps)

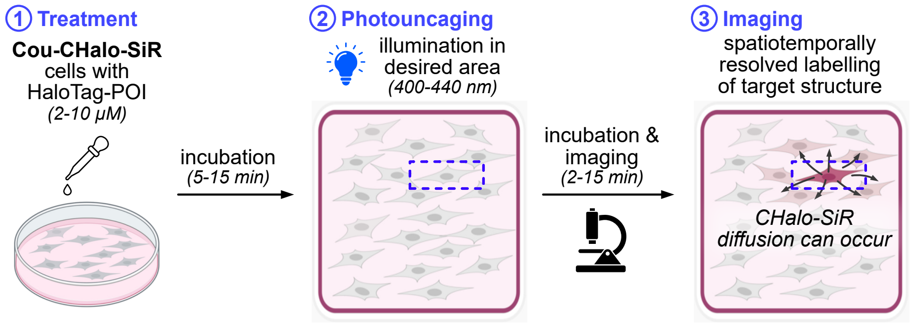

**Figure S2**: Experimental workflow for spatiotemporally controlled, fluorogenic HaloTag-POI labelling with **Cou-CHalo-SiR**.

**Cou-CHalo-SiR** can be used for spatiotemporally controlled fluorogenic labelling.

***Experimental steps (Fig. S2)***: (1) HaloTag-POI expressing cells are treated with **Cou-CHalo-SiR** for 5−15 min (the dye is well membrane permeable). We recommend starting with 2−10 µM concentrations to achieve high local concentrations since the released photouncaging product **CHalo-SiR** is rapidly diluted due to post-activation diffusion. (2) The desired region is illuminated with identical light doses as determined in the pre-tests (**Fig. S1**) and (3) SiR channel imaging is started immediately after uncaging to observe the fluorogenic HaloTag labelling of the target protein (excitation: 650 nm, emission: 670 nm). Typically, the signal plateaus within 5 min. Due to the cell-permeability of the **CHalo-SiR** photouncaging product before it ligates, the product can diffuse away from its activation site (**do not expect subcellular spatial resolution!**), and can exit the cell in which it was activated. After cell exit, **in 2D cell culture** it probably dilutes away in the cell culture medium, particularly if convective currents assist flow; whereas if cells are located **in a gel / matrix or multicellular organism**, "dilution away" is spatially limited and one can expect substantial labelling of neighbouring cells. Therefore, a fresh well of cells should be used for every experiment to achieve ideal temporal control for the labelling. (If neighbour labelling is seen in 2D cell culture without a gel/matrix, this probably indicates that the uncaging illumination was not truly cell- or subcellularly-localised.)

#### 1.4 Protein heterodimerisation with Cou-CHalo-BG

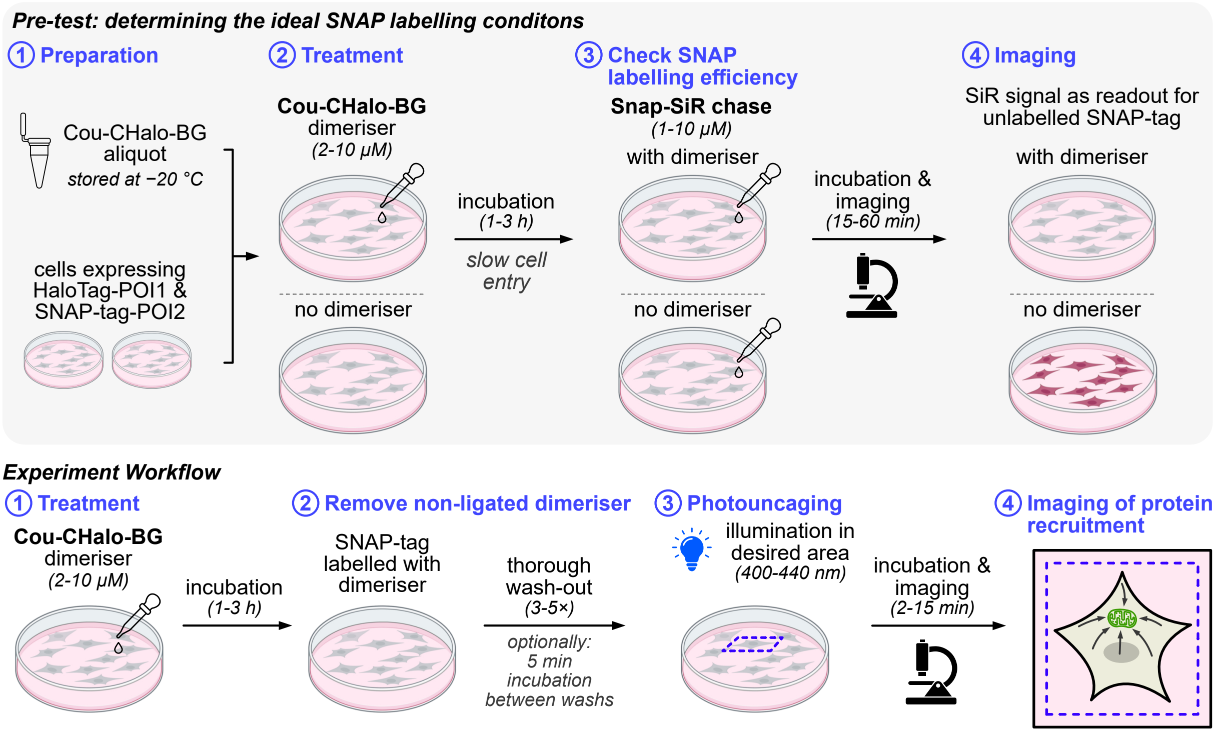

**Figure S3**: Experimental workflow for spatiotemporally controlled HaloTag-to-SNAP-tag protein heterodimerisation with **Cou-CHalo-BG**.

The photo-activatable SNAP-tag-to-HaloTag dimeriser **Cou-CHalo-BG** enables light-controlled protein heterodimerisation. For successful and quantitative dimerisation, efficient SNAP-tag labelling is crucial, therefore a pre-test to determine the SNAP labelling conditions should be performed.

***Experimental steps (Fig. S3)***: (1) Cells expressing the HaloTag-POI1 and SNAP-tag-POI2 proteins are prepared (*note: for successful dimerisation, both POIs must be expressed so they share access to the same compartment: e.g. cytsol to outer mitochondrial membrane or cytosol to inner plasma membrane dimerisation is possible, but not from ER to inner mitochondrial membrane*). (2) Cells are treated with fresh aliquots of **Cou-CHalo-BG** (2−10 µM) while in parallel control cells are cultured without treatment (DMSO control). Longer incubation times (1−3 h) than with **CHalo-SiR** are needed to reach high SNAP-tag anchor labelling, due to the lower membrane permeability of **Cou-CHalo-BG** compared to the **CHalo-SiR** dyes. (3) The medium is changed to remove the dimeriser (2× wash), then a SNAP chase dye (e.g. 1−10 µM Snap-SiR) is added to both the untreated cells and the cells pre-incubated with **Cou-CHalo-BG**, and is incubated for 15−60 min (needs longer incubation than for comparable HaloTag dyes), before the cells are imaged (checks for SNAP-tag labelling efficiency, by fluorogenic SNAP-tag ligation and SiR signal generation (excitation: 650 nm, emission: 670 nm)). **Ideally**, the cells pre-treated with the dimeriser show no SiR fluorescence and the untreated cells show high signal (alternatively a strong SiR signal reduction in the dimeriser cells allows to estimate the labelled portion of the SNAP-tag). If full labelling is achieved, the loading concentration (or, if really desired, the loading time) can be reduced so that it is easier to achieve full unligated wash-out in later applications. If very low labelling is observed, the loading concentration (or, in unusual cases, just the incubation time) should be increased.

***The actual experiment***: After establishing a SNAP-tag labelling protocol for the **Cou-CHalo-BG** dimeriser, it can be used for light-induced SNAP-tag-to-HaloTag protein heterodimerisation. (1) Cells expressing the HaloTag-POI1 and SNAP-tag-POI2 proteins are treated with fresh aliquots of **Cou-CHalo-BG** (2−10 µM) for 1−3 h. (*Note regarding protein expression: depending on the system and the readout, the experimenter should ensure that the ratio of HaloTag-POI1 and SNAP-tag-POI2 expression is tuned to support the desired effect. For example, for our experiments for cytosolic HaloTag-GFP recruitment to mitochondrial SNAP-tag, we aimed to achieve near-quantitative re-localisation of HaloTag-GFP and thus transfected it substoichiometrically (aim: higher expression of SNAP anchor than Halo target), then specifically chose to work with cells that had dim GFP fluorescence in the hope that the SNAP expression in those cells would be higher than the HaloTag expression (though, if there had been a SNAP reporter fluorescent protein, we would have used that to get a proper Halo:SNAP expression ratio estimate, which would have allowed better experiment design)*). (2) The excess of non-ligated dimeriser is removed by thorough washing (3−5× change medium, optionally with 5 min incubation after each medium change) to avoid undesired HaloTag ligation by residual non-SNAP-tag ligated dimeriser after uncaging. (3) The desired region is illuminated with identical light doses as determined before (see **Fig. S1**) and (4) the cells are imaged to observe the induced protein recruitment. For cytosolic HaloTag-GFP recruitment to mitochondrial SNAP-tag (on the outer mito-membrane) we saw relocalisation complete within 1 min, however, heterodimerisation kinetics will strongly dependent on the nature of the two POIs selected. If no relocalisation is observed, the wash-out protocol should be improved to avoid HaloTag ligation of remaining, non-SNAP ligated dimeriser (also see main **Fig. 1**).

**When the SNAP-tag anchor is prevented from freely diffusing in the cell (e.g. bound to an organelle surface or structure), subcellularly-spatially-specific uncaging should result in subcellularly-spatially-specific HaloTag recruitment.**

### 2 Reagent Overview (Fig S4-S5)

#### 2.1 CHalo Reagents

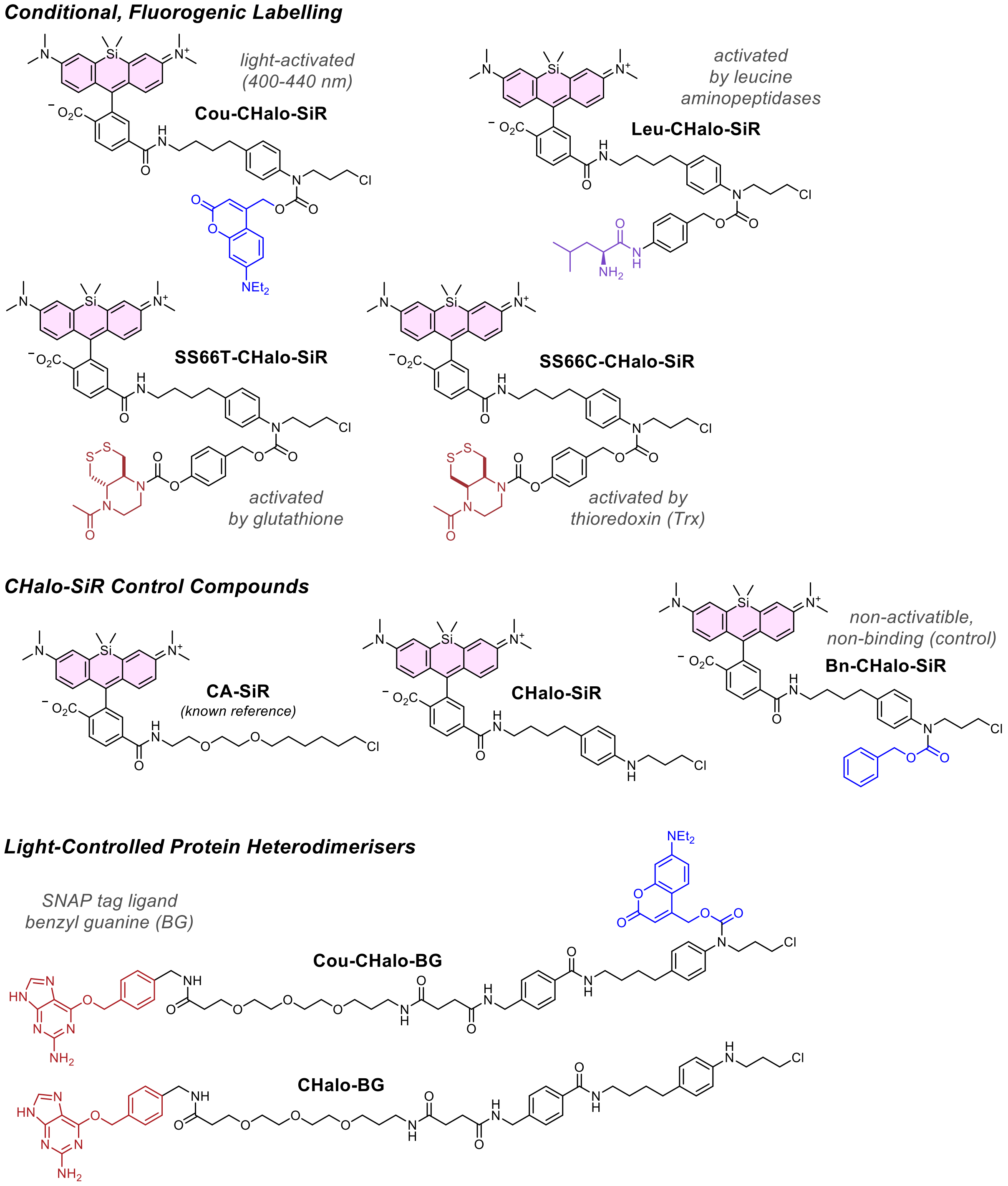

**Figure S4**: Chemical structures of **CHalo** reagents.

#### 2.2 Design Optimisation: proof-of-concept fluorescein conjugates

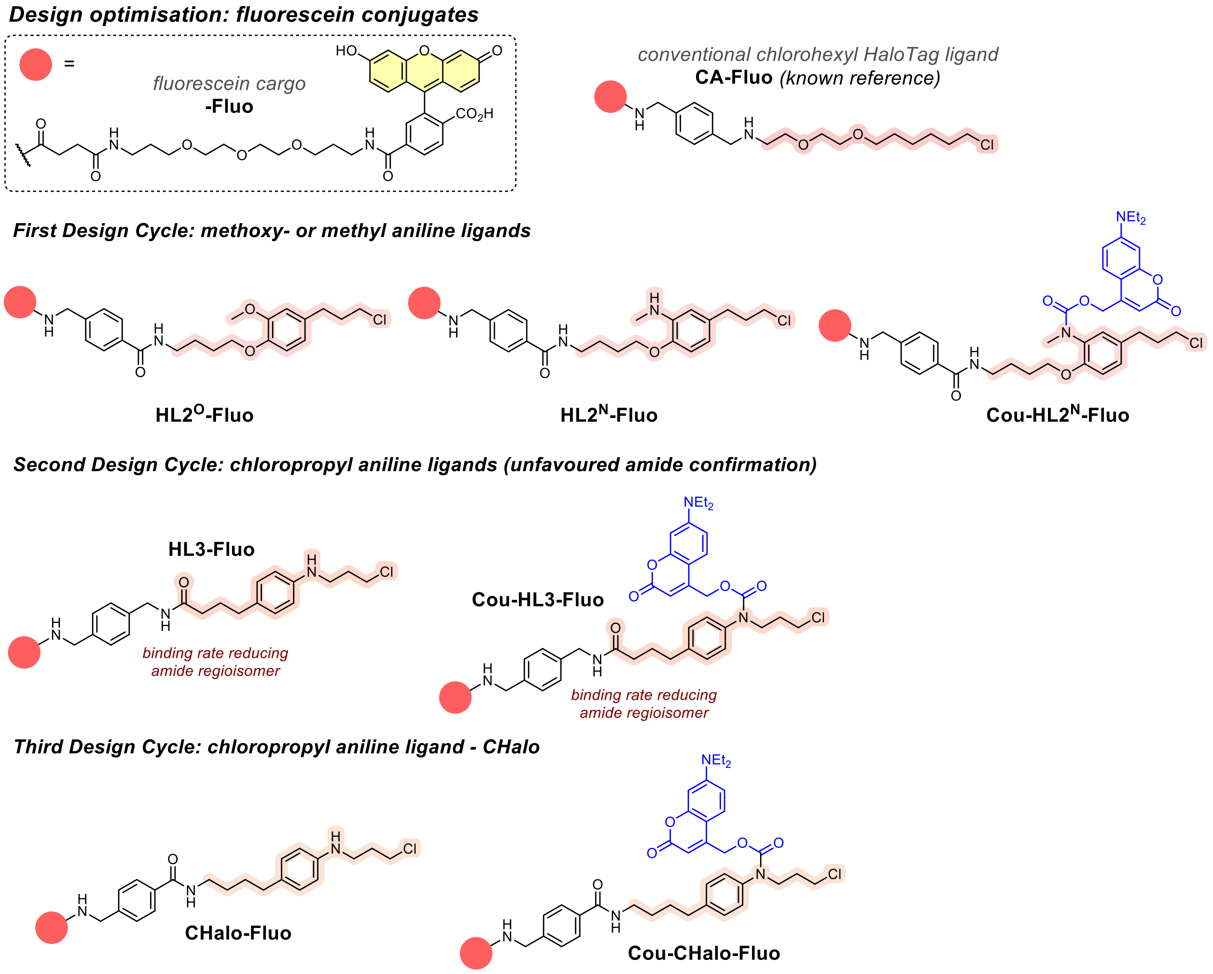

**Figure S5**: Chemical structures of initial HaloTag ligand designs as fluorescein conjugates which were iteratively optimised before arriving at the **CHalo** Design.

### 3 Supplementary Note 1: Ligand Design (Fig S6-S8)

#### 3.1 Initial Design ($HL2^N$ Ligand)

The HaloTag protein has been evolved to accept long, linear substrates like the classic chloroalkane motif (CA, **Fig. S6b**), with which it reacts at outstanding speed. The CA motif is used almost[1] exclusively for HaloTag reagents, so little is known about what structural changes are needed to make a CA derivative that *cannot* ligate to HaloTag. Since CA must insert down a long and seemingly narrow protein tunnel for ligation (**Fig. S6a**: crystal structure with CA-TMR[2]), it seemed likely that installing a bulky caging group on the chain, ≤10 atoms away from the reactive chlorine, would block ligation. However, the ethylene glycol ether backbone of the CA motif is chemically featureless, and does not offer a caging site, so we needed to install one. This site must be easy to derivatise with a range of stimulus-responsive cages in biochemically stable but easily uncaged form, which suggested installing an amine for caging as a carbamate since many such cage types are well known (for enzyme[3], light[4,5], or biochemical[6,7] uncaging). Yet, after uncaging this site, the motif must ligate HaloTag as efficiently as possible: which we imagined would be problematic for an aliphatic amine (e.g. as an isosteric replacement for one of the ether oxygens) due to its basicity and polarity.

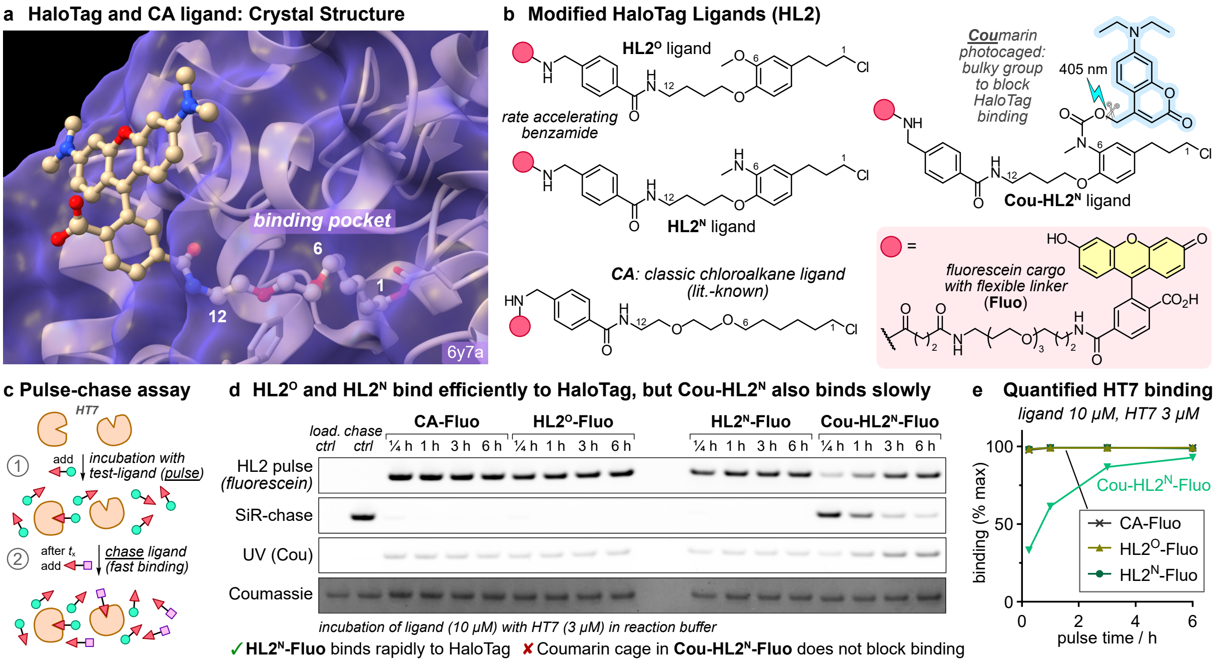

**Figure S6: Design of HaloTag ligand $HL2^N$ and its coumarin-photocaged version Cou-$HL2^N$ for light-induced HaloTag binding. (a)** Crystal structure of CA-rhodamine bound to HaloTag7 protein (HT7) with the chromophore positioned on the protein surface and the chloroalkane chain buried inside the protein *(pdb code: 6y7a)*; **(b)** Chemical structures of HL2-type fluorescein conjugates; **(c)** Schematic representation of the "pulse-chase" assay used to determine the binding efficiency of our ligands to HT7: (i) incubation with test-ligand (pulse), then (ii) addition of excess of **CA-SiR** which is known to rapidly bind HT7 (chase); **(d)** SDS-PAGE gel showing the binding efficiencies of **CA-Fluo** compared to **$HL2^O$-Fluo**, **$HL2^N$-Fluo** and **Cou-$HL2^N$-Fluo** (10 µM) each after incubation with purified HT7 (3 µM) for different times (15 min, 1 h, 3 h, 6 h); all rows show the HT7 band: (i) fluorescein fluorescence, (ii) SiR fluorescence, (iii) UV fluorescence (365 nm), (iv) Coumassie staining; **(e)** Quantified HT7-binding of HL2-type fluorescein conjugates (quantified from SiR-chase channel from panel d; quantification from fluorescein channel in panel d).

Thus, we seized on a recent report by Tadross *et al.*[8] who showed that a methoxybenzene is well tolerated as part of the HaloTag-ligating chain of **HTL.2** reagents (**Fig. 1e**). We exchanged the methoxy group for an only weakly basic *N*-methylaniline, yielding **$HL2^N$** where the carbamate-cageable nitrogen is six atoms away from the chlorine (**Fig. 1b**). We synthesised the fluorescein conjugate of the **$HL2^N$** ligand (**$HL2^N$-Fluo**), and its diethylaminocoumarin-photocaged derivative (**Cou-$HL2^N$-Fluo**), as well as the Tadross-inspired non-cageable analogue **$HL2^O$-Fluo** (which we expected should benchmark the rate penalty for installing an *N*-methylaniline instead of the **HTL.2**-type methoxybenzene). We performed pulse-chase assays (**Fig. S6c**) to assess their ability to ligate to the HaloTag protein: typically, a reagent ("pulse reagent") is incubated with purified HaloTag 7 protein (HT7) for a specific time ("pulse time") for ligation to take place, then an excess of a rapidly-ligating "chase ligand" (**CA-MaP555**[9] or **CA-SiR**[10]) is applied to saturate the remaining fraction of non-ligated HT7 so that the pulse labelling is essentially stopped after the given pulse time. The degree of labelling can be quantified by several methods, where different fluorophore wavelengths are used to quantify the pulse and the chase labels, e.g. by in-gel fluorescence analysis of an SDS-PAGE gel (which proves covalent attachment of the dye to the protein, otherwise the fluorescence signal would not overlap with the HT7 protein band). In our experience, pulse quantification is more reliably interpretable at low pulse labelling, with chase quantification becoming more reliably quantifiable with high pulse labelling.

Pleasingly, **HL2$^N$-Fluo** ligated very effectively to HaloTag (full binding within 15 min, like **HL2$^O$-Fluo**), which supported the aniline-type design logic. However surprisingly, caging it as a carbamate (**Cou-HL2$^N$**) did not stop the ligation reaction (binding half time $t_{½}$ ca. 0.5 h; **Fig. S6de**). We concluded that caging must be performed closer to the HaloTag ligation reaction site chlorine.

#### 3.2 CHalo Ligand Design

We now moved the cageable nitrogen as close as possible to the HaloTag ligation site. To avoid the potential for cytotoxic aziridine formation, we decided to place the cageable nitrogen not two but three methylenes away from the reactive chlorine, i.e. extending the chloroalkyl chain by one methylene unit from the Tadross design. This yielded **CHalo**, a simple substrate that is conveniently accessible from cheap starting materials yet surprisingly was never reported before in chemical literature for any purpose. The syntheses of caged cargo-bearing CHalo reagents were also simple and high-yielding (e.g. caging by reaction with a chloroformate, cargo attachment by amide coupling, **Fig. S7ab**). We anticipated that the potentially non-optimal positioning of the aryl ring might reduce ligation kinetics: but we now tested this for both free and caged analogues.

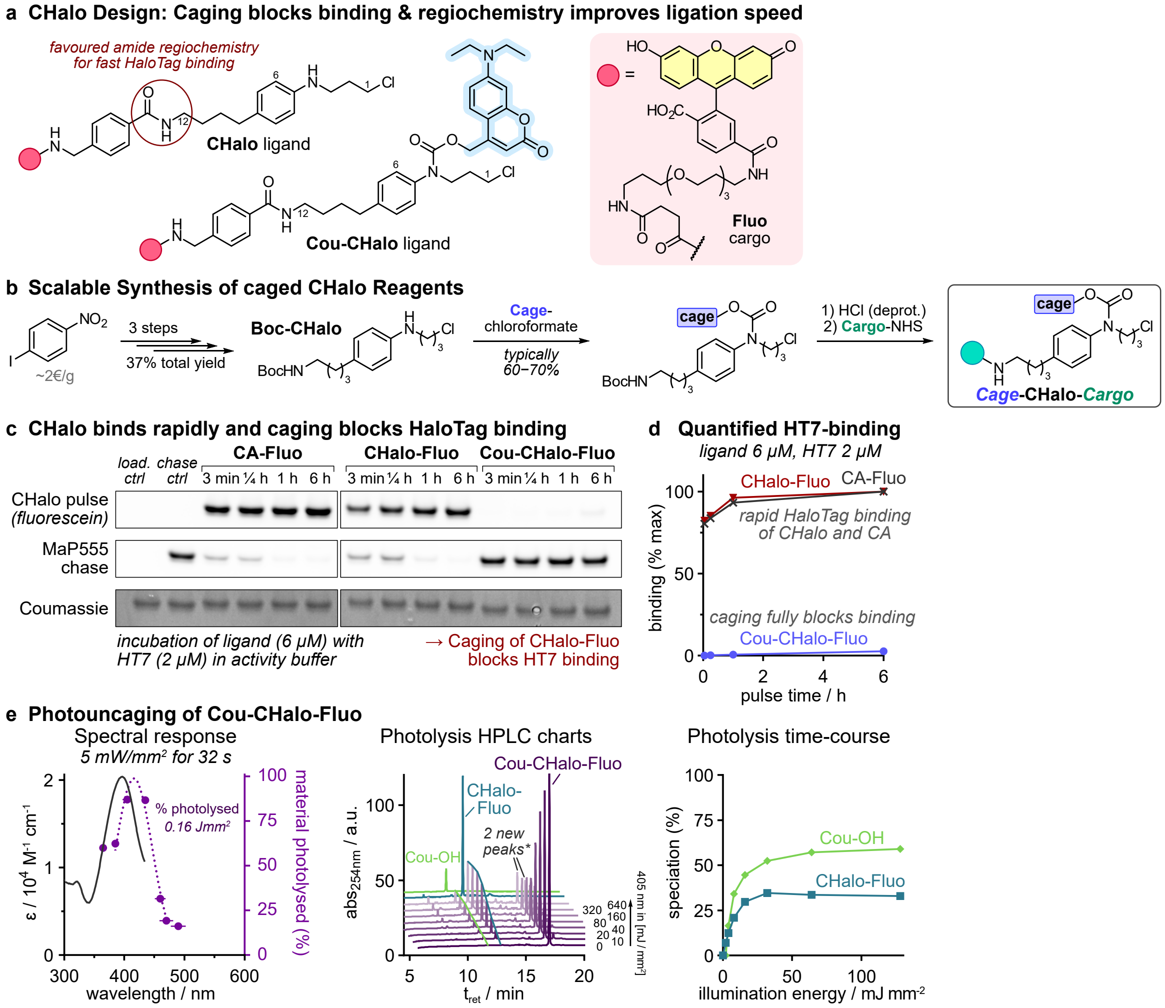

**Figure S7: CHalo Design: fast HaloTag Ligation and full binding suppression for caged Cou-CHalo ligand. (a)** Chemical structures of **CHalo-** and **Cou-CHalo-Fluo**; **(b)** Synthetic access to CHalo reagents; **(c)** SDS-PAGE gel showing the binding efficiencies of **CHalo-Fluo** and **Cou-CHalo-Fluo** (6 µM) each after incubation with purified HT7 (2 µM) for different times (3 min, 15 min, 1 h, 6 h); all rows show HT7 band: (i) fluorescein fluorescence, (ii) MaP555 chase fluorescence, (iii) Coumassie staining (full gel and quantification from SiR-chase channel in **Fig S14**); **(d)** Quantified HT7 binding of CHalo fluorescein conjugates (quantified from fluorescein channel in panel b); **(e) Cou-CHalo-Fluo** can be uncaged efficiently with blue light (ideal: 400−440 nm) and GFP orthogonally (no uncaging >470 nm; 50 µM sample in DMSO:water 7:3; illumination with the same integrated light intensity for each "wavelength" (horizontal error bars show FWHM of these quasi-Gaussian excitation light source LED "wavelengths"); HPLC speciation and photolysis time-course (50 µM in MeCN:water 1:1; illumination with 405 nm light, 5 mW/mm$^2$, applied for times from 1 to 128 seconds (factor 2 steps)); **the species in the "2 new peaks" close to the Cou-CHalo-SiR signal include the coumaryl-CHalo alkyl aniline which is a non-physiological byproduct of the high concentrations used in this assay (after an aniline has been photoliberated, it can trap another photogenerated coumarinyl unit: byproduct mass lacks $CO_2$).*

Free **CHalo** ligated rapidly to HaloTag protein with the fluorescein conjugates displaying similar ligation rates as the optimised linear CA substrate (**Fig. S7c**). Crucially though, the binding of carbamate-**caged-CHalo** was fully suppressed (under 1% after 100 minutes incubation, see **Supplementary Note 2** for analysis). Importantly, the CHalo ligand retains its HaloTag specificity and does not show off-target labelling of other proteins (**Fig. S16**). This now motivated us to create caged, logic-gated CHalo reagents for some of the applications that are impossible with any of the known, unconditional, HaloTag-ligating motifs; and began with the photoactivatable **Cou-CHalo-Fluo** which showed an expected uncaging action spectrum profile[11] with efficient uncaging from 380 to 450 nm (but no uncaging above 470 nm: which allows GFP imaging with lasers or filtered sources to be performed *in situ* without photouncaging), that converts **Cou-CHalo-Fluo** to free **CHalo-Fluo** with ca. >35% uncaging yield (expected for coumarin photocaged substrates[11], **Fig S7e**).

#### 3.3 Amide regiochemistry strongly effects ligation kinetics

Due to even easier synthetic accessibility, we had initially synthesised a cageable aniline ligand **HL3**, with flipped amide connectivity compared to the benzamide in the **CHalo** design (**Fig. S8a**). Caging **HL3** equally effectively blocked the HaloTag ligation, but the labelling speed of free **HL3** was significantly slower than free **CHalo** (**Fig. S8b**): showing the strong influence of amide regiochemistry on ligation kinetics.

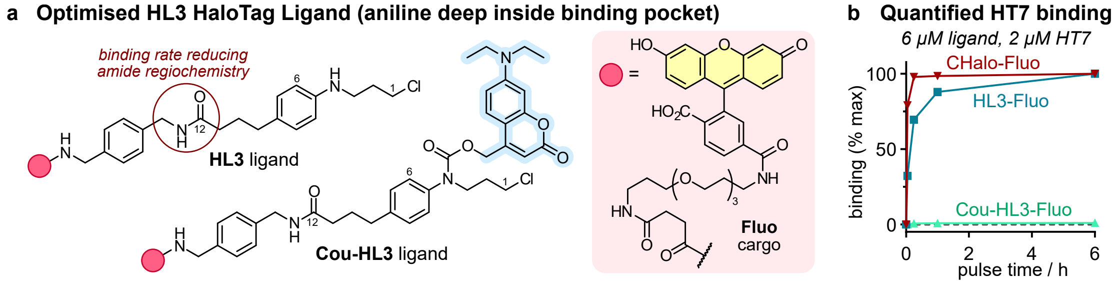

**Figure S8: HL3 Design shows that amide regiochemistry strongly effects HaloTag ligation kinetics. (a)** Chemical structures of **HL3-** and **Cou-HL3-Fluo**; **(b)** Quantified HT7 binding of **HL3** and **CHalo** fluorescein conjugates (quantified from fluorescein channel in SDS-PAGE gel from **Fig. S12**).

### 4 Supplementary Note 2: Rates & corrections for non-ligating compounds

In the text, for simplicity we state "less than 1% of **caged-CHalo** ligated within 100 minutes". In reality, this useful guide is also a broad generalisation: ligation is a bimolecular reaction, so the apparent rate depends largely on protein and ligand concentrations, as well as assay setup (temperature, medium, etc). But in practice, the more important question is: what was the real ligating species, when "caged ligation" was seen?

**We believe that the caged-CHalo reagents actually *do not* ligate at all**; and where ligation develops over time without a specific stimulus, what is occurring is an *in situ* hydrolysis or adventitious cleavage of the cage. The rates of those cleavages depend greatly upon assay settings and experimenters (hence e.g. our warning in the "user guide" **Supplementary Note 1**, to use red-emissive lights when handling Cou-caged reagents).

We do not want to under-sell the caging performance by falsely interpreting background signal as caged-ligation, but we also do not want to over-sell ligand performance by restricting the analysis to specific cage chemotypes and unusual assay settings (or by discussing only the quasi-non-existant caged-ligated fraction). Thus, in order to give a useful guideline about the real, in-practice utility of caged-CHalo reagents we gave this 1% figure, on the understanding that it essentially reflects their *O*-alkyl-*N*-aryl-*N*-alkyl carbamates' robustness, and should not be misinterpreted as "1% of the caged reagent will be found ligated onto HaloTag".

The "<1% in 100 minutes" figure is taken from our **Cou-CHalo-Fluo** assay with 6 µM ligand and 2 µM HaloTag, as used in standard benchmarking papers[12]: noting that Cou is a not-particularly-hydrolytically-resistant cage [compare: **Bn-CHalo-SiR** reaches 1% apparent signal only after 6 hours]; and though free **CHalo-Fluo** is also not the fastest-ligating substrate (compare to e.g. **CHalo-SiR**), the protein excess conditions ought to ensure that this "<1% in 6 hours" reflects rather a worst case scenario - **we believe that most experimenters will experience better stability and lower "unwanted ligation" in practice**.

**Data acquisition and correction for non-ligating compounds**

These assays were especially designed to test the *absence* of signal from caged compounds that are synthesised from a highly-signal-active precursor which can convert *in situ* to the signal-active form (by benzylic hydrolysis, and for the Cou reagents also by photouncaging from background lighting during assay setup and stop reagent addition). We chose to work in "typical" conditions that would be careful, but not overly controlled, hoping that these would reflect the experience of a typical experimenter in a real assay setup.

(1) As is typical for testing inactivity from a caged target, we expected the inevitable small amount of residual uncaged compound present at time zero in the assay to rapidly generate a signal plateau in our experiments, that might be minor (when roughly 1:1 ligand:protein are used) or significant (when vast excess of ligand is used). Therefore, when estimating the resistance to ligation of the caged compounds, we excluded a portion of this residual signal source, in a relatively bias-free way (that underestimates the likely residual signal by a factor ca. 2).

For the assays with 3:1 ligand:protein ratio, the direct ligand signal at the 5 min timepoint (fluorescein channel) was interpreted as the residual; for example, a loading-corrected gel band fluorescence value of 0.0053 (**Cou-CHalo-Fluo**) was interpreted as arising from 0.2% residual free **CHalo-Fluo**, and this value subtracted from all subsequent assay values (no adjustments for increasing reaction over time).

For the assays with 100:1 ligand:protein ratio that used the same **Cou-CHalo-Fluo** stock but were prepared on a different day, we expected only a slight difference in residual fit. Indeed, the fitted residual percentage was 0.1%.

For the assays that used **Bn-CHalo-SiR** and were prepared by the same final step purification as for **Cou-CHalo-Fluo**, we also expected a similar residual fit. Indeed, the fitted residual percentage was 0.1%.

(2) Both reagents are theoretically capable of uncaging over time inside the assay, by benzylic hydrolysis (expect: Cou much faster than Bn) and/or by photouncaging from stray light (Cou only). These uncaging modes should generate linear signal increases over time at low labelling percentages, since HaloTag protein will always be in vast excess compared to the *in situ* uncaged fraction.

For the Bn reagent assay (only run at 100:1 ratio, only readout by MaP555 chase), we saw only very small signal increase over time, and it was indeed apparently linear (fluorescence reaches 0.2% of theoretical maximum after 1 hour, 1% of theoretical maximum after 6 hours: i.e. *in situ* uncaging of only 0.01% of the applied **Bn-CHalo** reagent). These are very small numbers in an absolute sense and argue for the general stability and utility of the *O*-alkyl-*N*-aryl-*N*-alkyl carbamate caging strategies that CHalo was designed to harness.

For the Cou reagents, we believe that hydrolytic uncaging may be a more significant ongoing process, since the signal increases were concentration-dependent, and proportional to ligand concentration; but (a) the uncaging is still slow, such that the reagents can be useful binary indicators in photo-triggered applications in biology; and (b) we believe that its rate is specific to the Cou group, and need not be a feature of other photocages or enzyme cages. The 6 µM ligand : 2 µM HaloTag assays show a roughly linear signal increase (0.6% at 1 hour, 2.8% at 6 hours); the 20 µM : 0.2 µM assay has higher increase which is unsurprising from its higher concentration if free ligand is the limiting factor (1.4% at 1 hour, 18% at 6 hours, in the direct readout channel). We believe that our assay setup and quenching steps are the ones that apply light to the sample, whereas the incubation protected in the dark excludes it, therefore we believe that *in situ* uncaging does not reflect Cou photouncaging.

Although we used MaP555 chase as a counter-readout, we considered it ***not suitable for quantifying low ligation*** data, because in our benchmarking experiments the chase channel had assay-to-assay variability of typically ±15% of readout maximum when used to assess conversions at low CHalo ligation percentages. Therefore the MaP chase data are not shown on graphs for **Cou-CHalo-Fluo** systems because their intrinsic Fluo channel readout is more reliable (though we quantified e.g. 25% readout at 6 hours for **Cou-CHalo-Fluo** which roughly matches the 18% from intrinsic readout), and MaP chase data are only graphed for the 100:1 **Bn-CHalo-SiR** run which has no intrinsic channel but should have the maximum possible parasitic labelling.

(3) Once these overly simplified corrections for 0.1-0.2% residual free ligand were introduced, the caged-reagent assay data are much more plausibly interpretable than before.

The 3:1 assay now has a linear fit appropriate to e.g. an ongoing pseudo-first-order reaction where the availability of *in situ* uncaged **CHalo-Fluo** is the limiting factor, rather than its previous form (a saturation curve form with half-conversion by ~2 hours but plateauing at just 3.5%).

The same is true for the 100:1 assay with **Bn-CHalo-SiR** (now ongoing linear; previously a saturation curve with half-conversion after about 5 minutes [similar to the kinetic for free **CHalo-SiR**] but plateauing at just 11%); and the 100:1 assay with **Cou-CHalo-Fluo** (previously pausing at 10% conversion from 15 min to 1 hour, then reaching 30% at 6 hours; now nearly linear increase at approx. 3% per hour).

**Plausibility**: In all caged-reagent assays, the residual free ligand corrections applied only alter the signal curves by subtracting a value that is less than the uncorrected signal developed within 15 minutes. If signals specific to the actual reagent species had been developing, they would simply appear to be delayed by about 15 minutes as a result of these corrections, without changing their form. Instead, the form of these data now match plausible expectations for their mechanism (caged forms never ligate, tiny portions that get uncaged are what ligates effectively): confirming the suitability of this correction.

##### Caged-Ligation with $HL2^N$ reagents?

We had been surprised that **$HL2^N$** reagents apparently ligated even when caged, so checked the plausibility of caged-ligation with different methods than just the standard pulse/chase imaging channels. In the end, we found reliable confirmation by imaging the in-gel Cou motif fluorescence from ***HaloTag-ligated Cou-$HL2^N$-Fluo*** reagents using a broad "UV excitation" setting (filtering for roughly 365 nm excitation, 450 emission) which acquires signal more intensively from the Cou group than from the fluorescein group (approx ratio 75:25, determined after using an SiR chase to confirm the percentage occupancy, then assuming that fluorescein motif UV intensity within ligated **Cou-$HL2^N$-Fluo** is identical to the motif UV intensity for **$HL2^N$-Fluo** and subtracting the occupancy-adjusted partial contribution of the fluorescein motif).

In that reagent case, the efficiency of caged ligation was so high (ca. 1/3 the rate of uncaged **$HL2^N$-Fluo**) that the levels of (residual plus *in situ*) uncaged **$HL2^N$-Fluo** ligating to HaloTag proved much smaller than the level of caged-reagent ligation, which both allows robust confirmation that the caged-reagent was really the ligating species, and allows robust analysis of its binding kinetics by the "UV channel" signal intensity (the fluorescein motif component does not even need to be normalised away since the stoichiometry is enforced as 1:1). Additional corrections that might be desired for fine quantification, e.g. for fluorescence intensity modulation due to the dual chromophore ligand, were considered superfluous because the main effect, that the caged reagent ligates, was already visible as confirmed by the 3-fold higher fluorescence intensity (from the Cou motif) which dominates the UV band and which is not a feature of the free **$HL2^N$-Fluo** comparator (**Fig. S6**). (Future experimenters interested in this phenomenon might consider using an isosteric coumarinyl-propionamide rather than coumarinyl-hydroxymethylcarbamate, to prevent photolability of the product and allow for e.g. MS-based quantification).

This observation seems in line with recently emerging results that argue that HaloTag is rather more tolerant for alkylator ligands than had previously been assumed (viz. the assumption that the linearity and length of the classic chloroalkane linker is strongly required so that a reagent can dock and ligate); and suggests additional modes of functionalising HaloTag-reactive ligands with two orthogonal cargos, while preserving decent to very good ligation kinetics (probably the aryl ring and substituent disposition that the **$HL2^N$** design inherited from Tadross and coworkers, is crucial in this respect, since it provides a very fast basis for ligation).

### 5 Supplementary Note 3: Harnessing known cage reactivities with CHalo

The CHalo system can in principle harness a variety of useful cages to allow ligation after different stimuli. Such caging groups have been broadly developed in prior literature, e.g. for triggering uncaging with light (photocages), enzymatically, or with a biochemical stimulus; and these caging groups can easily be introduced onto the CHalo ligand as carbamates (**Fig. S9a**) allowing straightforward implementation of diverse gating applications - a crucial flexibility advantage of small molecule reagents versus genetic tools.

A variety of **photocages** are available with tuned uncaging wavelengths (**Fig. S9b**). The most commonly used *ortho*-nitrobenzyl (*o*-NB) photocages are cleaved with UV light, with some modifications up to 380 nm (disadvantageous: relatively high phototoxicity). Dialkylaminocoumarins (e.g. DEACM) can be activated with violet light and we used the DEACM cage since it allows parallel GFP imaging (490 nm excitation) without photo-cleavage. Depending on the application, blue/green/yellow-responsive coumarins or other cage types such as BODIPYs or xanthenes[4] could be designed instead; and even far-red to near-infrared uncaging by cyanine cages[13,14] or SiR-assisted cleavage of *o*-NB cages[15] are feasible: i.e. the whole visible light range is conceptually accessible for CHalo. For more literature examples we refer the readers to this review[5].

**Enzyme cages** are widely used for fluorogenic enzyme activity probes (**Fig. S9c**), with a broad range of target enzymes such as nitroreductases, peptidases, esterases, glucosidases or oxidoreductases.[3,16,17] Such probes enable the sensitive detection of enzyme turnover (instead of just enzyme expression level) and can reveal changes in enzyme activity (e.g. in disease) rendering them invaluable research tools. **Biochemical** cages for sensing stimuli such as hydrogen sulfide or hydrogen peroxide are also known (**Fig. S9d**).[6,7]

All these cages are well known for releasing either phenols or anilines (optionally with the use of chemical adapters[18]), so they can conceptually be used straightforwardly to cage the CHalo ligand and thus create diverse photo-/enzyme-/biochemically-caged probes. For more examples we refer the readers to this review[3].

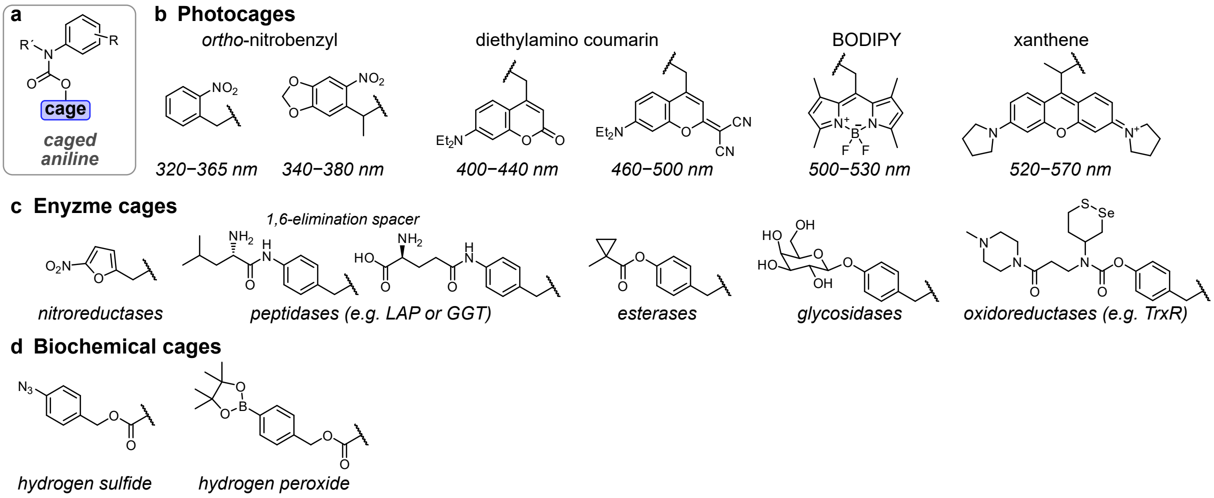

**Figure S9: A selection of known caging groups** that should be chemically compatible with the CHalo motif, and could in the future be applied to harness photo-, enzyme- or biochemical uncaging (optionally using chemical adaptors).

### 6 Supplementary Note 4: CHalo Ligation and CHalo-SiR Fluorogenicity

**<u>Conclusions:</u>** (1) CHalo's intrinsic ligation rate (ca. 1/10 that of CA) as well as its cell permeability are "good enough" to perform strongly in cells in settings where normal CA reagents succeed. The target protein choice and the CHalo reagent cargo's barrier crossing ability will be the key determinants of any given reagent's *actual* labelling speed in biology. (2) **CHalo-SiR** is a powerful fluorogenic label, much like SiR-CA. (3) **Caged-CHalo-SiR** reagents are cage-conditional fluorogenic labels that inherit the uncaging features known for their chosen cage, then (when uncaged) give the ligation and fluorogenicity known for CHalo-SiR.

**<u>Background:</u>** Fluorogenic chloroalkane-siliconrhodamine ligands are perhaps the most widely used of all HaloTag reagents (sold as SiR-Halo, SiR-CA, Halo-JF646, etc). These ligands excel for low-background, low-photodamage, far-red imaging of HaloTag fusion proteins, since their fluorescence in the cell environment is low until they ligate to HaloTag (typically quoted: 5 to 20-fold fluorescence turn-on upon ligation)[10]. While the bimolecular reaction rate for TMR/CPY-chloroalkane ligating to HaloTag protein can be incredibly fast in purified cell-free conditions (up to ca. $10^8$ $M^{-1}s^{-1}$)[12], this is not the rate-limiting aspect for performance in biology: because the ligation reaction requires first that the reagent encounters its target protein, i.e. typically it must cross at least the plasma membrane, while remaining soluble and bioavailable, and the rate of crossing barriers is almost always much slower than the rate of ligation once they are crossed.[9] The assay setting and the cellular location of the HaloTag fusion protein determine what barriers must be crossed and therefore greatly control how fast the overall crossing-then-ligation can proceed: e.g. very low hindrance for extracellular labelling in 2D cell culture following reagent addition to the medium (very fast fluorogenicity), high hindrance for intracellular labelling in the brain *in vivo* following i.v. administration (very slow fluorogenicity). The structure of the cargo attached to the chloroalkane (or CHalo) motif also greatly controls barrier crossing rates: e.g. high membrane permeability for "MaP" dyes that favour a spirocyclised unligated state, vs low membrane permeability for permanently open (zwitterionic) dyes.

**<u>Goals:</u>** Given that target encounter rate (restricted by barrier crossing rate e.g. by cell permeability) is usually rate-limiting for HaloTag ligation in live biology, we consider that the best outcomes for the CHalo motif *as a conditional ligation motif* would be that (a) neither CHalo nor typical caged-CHalo motifs make their reagents dramatically less permeable than the known chloroalkane motif; and (b) CHalo reagents are not dramatically slower to ligate than chloroalkane reagents. We expected that if our (intentionally) simple CHalo reagents had slow apparent labelling rates in live biology, the first point of improvement would be to switch cargos to the known faster-cell-penetrating dyes (like the MaP series) as cargos; and we would only consider the apparent intracellular labelling rates or the apparent cell-free labelling rates as problematic features of the CHalo motif *if* those rates were >100x slower than the rates for cognate chloroalkane compounds.

To determine what performance both **CHalo-Cargo** and **Caged-CHalo-Cargo** reagents could have, as compared to known CA-SiR, we also used silicon-rhodamine as the cargo, and made **CHalo-SiR** as well as caged **Cou-CHalo-SiR** and **Bn-CHalo-SiR** as testbed reagents.

**<u>Solubility and aggregation problems call for caution about ligation rate and fluorogenicity assays:</u>** At least in our hands, SiR-Halo, **CHalo-SiR**, **caged-CHalo-SiR**, and presumably other similarly-sized reagents with high $logD_{7.4}$ values (e.g. expected for those with CPY as cargo), seem prone to aggregation in biological media even at low concentrations (e.g. 500 nM), and presumably also to adsorption effects (e.g. onto the plastic of well plate or cuvette walls or pipette tips depending on the solvent, as well as onto e.g. albumin protein). This may be reflected in the choice of high BSA concentrations (10 µM) as a standard component of the "activity buffer" used for much development work around dye ligands for HaloTag. At least in our hands, the strong tendency to aggregation is coupled with large variation in how much aggregation is actually ongoing (increases over time, also depends on ligand concentration, buffer / cosolvent, assay temperature, handling and dilution steps, other assay components, etc) as well as how reversible the aggregation is, as well as how problematic that aggregation is (depending on readout).

For example, we knew already from work with well-water-soluble and permanently-fluorescent **Caged-CHalo-Fluo** reagents that caged CHalo species should not ligate to HaloTag over even 6 hours (**Fig S7cd**, **Supplementary Note 2**). Indeed, with caged-CHalo-SiR (which we expected would have low fluorescence in cellular environments, due to its preferred biolocalisation into apolar environments where the non-red-absorbing and hence non-fluorescent spirocyclised form would dominate), <u>cellular</u> assays did not show strong fluorescence: so we were confident that no spontaneous ligation or uncaging-then-ligation was occurring (since this would lock the SiR into place on the HaloTag protein such that the fluorescent zwitterionic form now dominates). <u>Cell-free</u> assays are coherent with this lack of spontaneous ligation or uncaging-then-ligation: the caged-CHalo-SiR reagent has very low UV-Vis absorption and very low fluorescence (**Fig S10fg**). In our assays, CA-SiR had apparently 9-fold lower absorbance and 5-fold lower fluorescence emission intensity in cuvette (aqueous activity buffer, including 10 µM BSA) in the absence of HaloTag protein, than when HaloTag protein was supplied in excess and ligation was allowed to plateau (**Fig S10b**). Under the same conditions, CHalo-SiR had apparently 40-fold lower absorbance and 400-fold lower fluorescence emission intensity in the absence of HaloTag protein as compared to the presence of HaloTag (**Fig S10c**). Notably, the UV-Vis absorbances of CHalo-SiR and CA-SiR incubated with excess HaloTag protein were essentially identical (scaling by a factor 0.85, which is within assay variation limits, **Fig S10bc**); and even the raw data values of their fluorescence emission spectra were essentially identical (**Fig S10bc**: although each emission spectrum is technically on an arbitrary unit scale, the experiments were acquired under otherwise identical conditions, and the raw value scaling was by only 0.91).

**We trust the conclusions that** ligated CHalo-dye reagents will have similar or identical fluorescence intensity and spectrum as their ligated chloroalkane-dye counterparts; and that non-ligated CHalo-dye *or* caged-CHalo-dye reagents will have similar or identical fluorescence properties as their non-ligated chloroalkane-dye counterparts. **However, we caution against** simplistic claims on the basis of this assay type that e.g. "CHalo-SiR is 400-fold fluorogenic upon target binding whereas CA-SiR is only 5-fold fluorogenic", and we caution against interpreting the without-vs-with HaloTag data in terms of open/closed ratios and/or fluorescence quantum yields. The absorbance is entirely dependent on the open/closed ratio, which depends on the microenvironment (e.g. should have one value if the dye is ligated to protein; but if not it will depend on the solvent nature and its dielectric if the dye is not ligated but is in molecular solution, or else it will depend on how the dye is part of an aggregate/adsorbate/precipitate if not in molecular solution); and the fluorescence depends both on the absorption as well as on additional microenvironment features that affect e.g. quantum yield. Therefore, when there is low or no ligation, the measured fluorescence values in cell-free settings may actually have more to do with whether the unligated reagent is in molecular solution or part of an aggregate/adsorbate (extrinsic features of the cell-free assay setup that do not translate to cellular settings), rather than reflecting intrinsic parameters that do translate to any possible cell assay setting.

**$D_{50}$ values**: Those solubility-related problems were raised previously[9], with the recommendation that determining open/closed ratios across a range of solvent dielectrics is a better way to characterise the ability of a dye to be fluorogenic in cells thanks to ligation-induced de-lactonisation.[9] For example, $D_{50}$ values can be defined as the dielectric constant at which a dye has 50:50 open:closed ratio when all of the dye is in true molecular solution. SiR reagents are reported to have $D_{50}$ values ca. 30−35,[10] showing that this dye has a high tendency to spirocyclise even in somewhat polar environments: and due to most SiR reagents' overall hydrophobicity, this should ensure that in cells, most unligated SiR-bearing compounds will be nonfluorescent lactones since they will preferentially partition into hydrophobic environments with low dielectrics (**Fig. S10e**). We measured the absorbance of CHalo-SiR in water-dioxane mixtures; at high dielectrics (<20% dioxane) we saw a drop in solubility which compromised the high-end of titration and made it uncertain whether a plateau was reached (all-zwitterionic state), however we could confidently determine that its $D_{50}$ value by this method was ≥53: so it likewise favours dielectric-and-partitioning-based fluorogenicity in the cellular setting.

**Ligation Rates?** Given that the data so far support caging-control of ligation as well as ligation-control over the open/closed ratio and fluoresecence, and given the arguments about fusion protein and barrier crossing determining biological performance more than cell-free ligation rates, we felt it would be necessary but also sufficient for typical performance in cellular settings that uncaged CHalo-SiR should show somewhat similar ligation rate as SiR-CA in reasonably robust cell-free settings; and we did **not** attempt to measure cell-free ligation rates under any particular conditions with high precision.

We screened several cell-free parameter choices until we were satisfied with the robustness of our analysis. For example, under standard conditions from the literature ("activity buffer" with 10 µM BSA, 4 µM HaloTag, 2 µM ligand, 1% DMSO), we found that apparent ligation rates were drastically affected by changing the DMSO percentage and handling procedure in ways that indicate that aggregation/insolubility of CHalo-SiR (and also of SiR-CA) were compromising the actual ligation rates (e.g. easily making ligation seem 10-100 fold slower than when solubility was ensured by using higher cosolvent percentage, lower ligand concentrations, etc). We approximated the **CHalo-SiR** ligation rate particularly by using minimal dye concentrations and by varying DMSO concentrations until the apparent rate was stable (**Fig. S10i**; 1-10% DMSO).

Using the conditions we feel were reliable, we found that **CHalo-SiR ligated to HaloTag with a bimolecular reaction rate constant of ca. $10^6$ $M^{-1}s^{-1}$ at 23°C in cell-free assays** (**Fig S10i**: 5 nM dye, 200 nM HaloTag, 3-10% DMSO), indicating that the rate constant would be higher still at physiological 37°C. This is about 20-fold slower than typical values quoted for SiR-CA ligation at 37 °C;[12] and although in our hands SiR-CA ligated too fast to be well-resolved by the measurement techniques we employed for CHalo compounds, our estimate for the SiR-CA ligation rate was **10-fold faster** than CHalo-SiR ligation rate (**Fig S10h**), which we assume reflects an intrinsic 10-fold slower ligation of the CHalo than the CA motif. We feel that this intrinsic ligation rate difference is molecularly plausible (given the change of ligand structure compared to the optimised Tadross scaffold that the need for conditional ligation had enforced). Also, since CA reagents are considered useful over a vast range of ligation kinetics (from $10^4$ to $10^8$ $M^{-1}$ $s^{-1}$ depending on their cargo), we considered that (a) the only 10-fold difference would not make CHalo reagents useless, especially when the cargo is one that favours higher reaction rates; (b) the cell-free ligation rate difference would still be irrelevant in almost all biological practical applications since it leaves the CHalo ligation rate as being much faster than the other rate steps we expected to be important or rate-determining for practical performance.

Indeed, **cellular assays** showed much slower CHalo and CA labelling rates (half-lives 0.6-3 hours, **Fig S10j**) than in cell-free assays (seconds to minutes). They were also only 5-fold different from each other.

Finally, to validate the principle of **CHalo uncaging**, we studied the aminocoumarin-photocaged fluorogenic reagent **Cou-CHalo-SiR**. The reagent inherited the expected properties known for its cage: it was efficiently uncaged with blue light (50% conversion with 10 mJ/mm$^2$ at 380-440 nm, **Fig. S10k**), yet unaffected by wavelengths used for typical GFP imaging (no uncaging above 470 nm). Incubation with purified HaloTag protein showed no ligation (and therefore no fluorescence generation) before uncaging light, but rapidly generated fluorescence after being illuminated with 405 nm light (**Fig. S10l**).

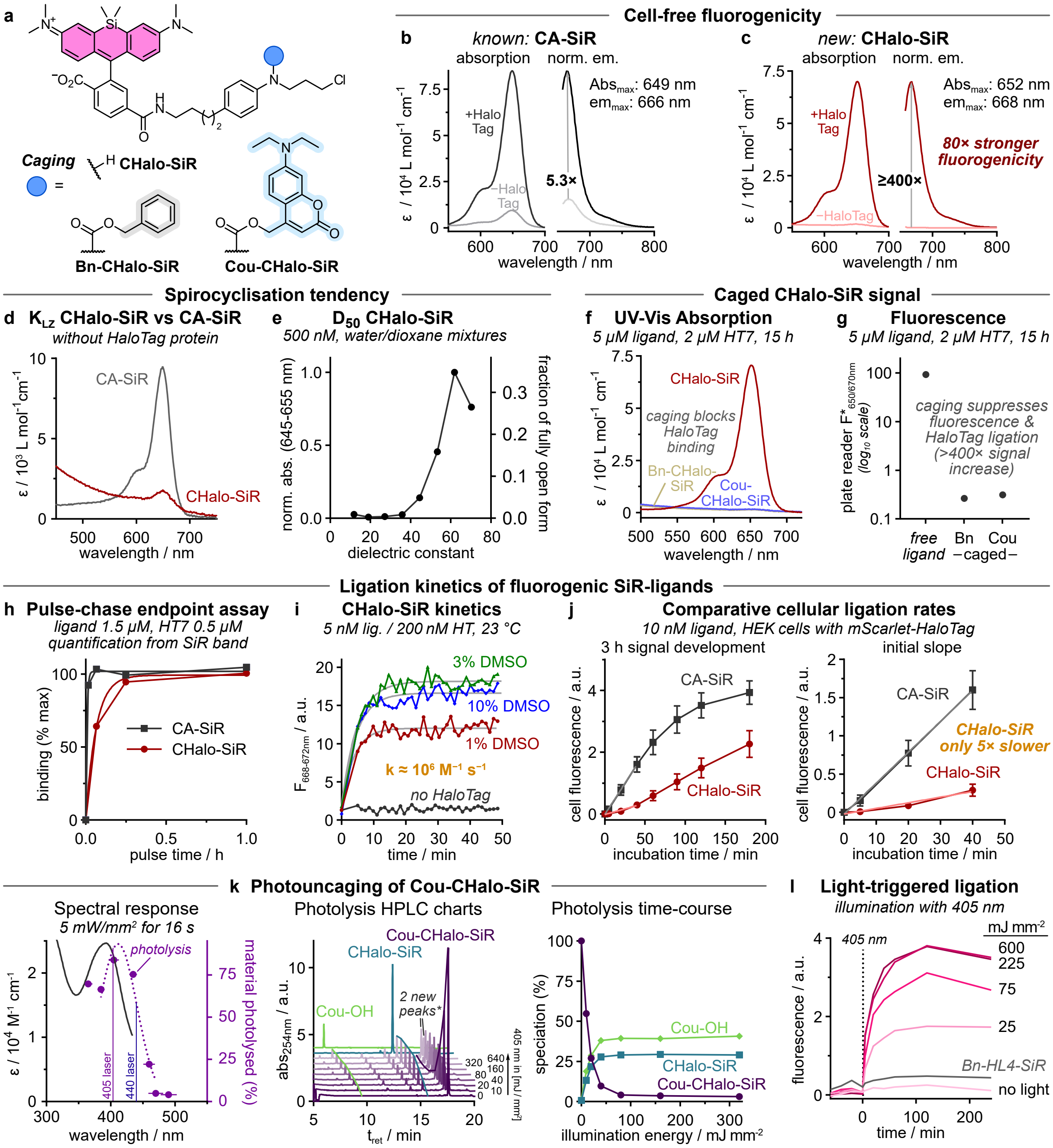

**Figure S10: Absorbance, fluorogenicity and photouncaging properties of Cou-CHalo-SiR. (a)** Chemical structures of **CHalo-SiR** and its caged derivatives **Bn-** and **Cou-CHalo-SiR**; **(b-c) CHalo-SiR** has 400-fold far-red fluorescence turn-on upon HaloTag binding (2 µM ligand ± 4 µM HaloTag7). **(d)** Investigation of the lactone-zwitterion equilibria of non-ligated **CHalo-SiR** and **CA-SiR** (UV-vis spectra without HaloTag protein). **(e)** CHalo-SiR absorbance as a function of the dielectric constant (absorbance normalised to maximum absorbance, measured in water/1,4-dioxane mixtures; 0.5 µM CHalo-SiR); aggregation effects cause low absorbance value at high dielectric constant. **(fg)** Visible region absorption spectra and fluorescence emission of free **CHalo-SiR**, and caged **Bn-** and **Cou-CHalo-SiR** after incubation with HaloTag protein (5 µM ligand, 2 µM HT7, 15 h); **(h)** Ligation kinetics of CHalo-SiR compared to CA-SiR quantified from SiR in-gel fluorescence (SDS-PAGE gel and quantification in **Fig. S13**); **(i)** Binding kinetics of **CHalo-SiR** determined by fluorescence increase upon HaloTag binding (5 nM ligand, 200 nM HT7, in PBS with 1−10% DMSO, 23 °C); **(j)** Cellular ligation rates of **CHalo-SiR** and **CA-SiR** (10 nM) in HEK cells transfected with cytosolic Halo-mScarlet (quantification from microscopy images; *n*=3). **(k) Cou-CHalo-Fluo** can be uncaged efficiently with blue light (ideal: 400−440 nm) and GFP orthogonally (no uncaging >470 nm; 50 µM sample in DMSO:water 7:3; illumination with the same light intensity for each wavelength, *horizontal error bars: FWHM of excitation light*; HPLC speciation and photolysis time-course (50 µM in MeCN:water 1:1; illumination with 405 nm LED light, 5 mW/mm$^2$, applied for times from 2 to 128 seconds (factor 2 steps)); **the species in the "2 new peaks" close to the Cou-CHalo-SiR signal include the coumaryl-CHalo alkyl aniline which is a nonphysiological byproduct of the high concentrations used in this assay (after an aniline has been photoliberated, it can trap another photogenerated coumarinyl unit: byproduct mass lacks $CO_2$)*; **(l) Cou-CHalo-SiR** generates fluorescence upon photouncaging followed by HaloTag binding (5 µM ligand with 2 µM HT7; 1 h pre-incubation, then 405 nm, 5 mW/mm$^2$).

### 7 Supplementary Note 5: HaloTag as an anchor; and biological background

##### *§1: Biological Background for prior usage of SLPs (expanded from the main text's Introduction)*

The HaloTag[19,20], SNAP-tag[21,22], and CLIP-tag[21,23] self-labelling proteins (SLPs) have enormously expanded the possibilities of *chemical biology* because they offer a high-specificity, orthogonal connection, latching the functionality of very diverse *chemical* reagents onto very diverse *biological* targets. The HaloTag has been broadly used,[24] especially for fluorescently labelling proteins of interest (POIs) with typically fluorogenic, small molecule fluorophores[9,10,25,26] (**Fig 1a**) that have been applied for studying protein localisation[27], movement and trafficking[28], intracellular protein concentrations,[29] or protein turnover,[1] in living cells; or for ultrafast staining of thick tissues after fixation[30]. Other major fields of application are chemically induced protein dimerisation (CID)[31–33], targeted localisation of biosensors (e.g. for calcium)[34], specific protein degradation,[35] or for increasing the local concentration of a reagent to enforce binding or reaction, through tethering (DART,[36] T-REX[37]). Conceptually, HaloTag has been the most attractive SLP since (1) its ligation rate to its conventional ligands can be up to 1000 times faster than that of SNAP and CLIP to theirs[38], (2) it is reported to be entirely monomeric, rather than introducing partial aggregation tendency due to its uniform negative surface charge[39], (3) it is reported to have no organelle localisation preference, (4) its classic HaloTag-reactive motif, i.e. the 6-chlorohexyl bisether **CA** (**Fig. 1a**) is more apolar than SNAP/CLIP motifs which generally favours the passive cell entry of Halo ligands over those of SNAP and CLIP[40].

##### *§2: General features for and against "using HaloTag as an All-Purpose Anchor"*

Using HaloTag as an "all-purpose anchor protein" to durably mark cells or proteins with molecular reporters at a specific time or region, or upon specific stimuli, potentially against a non-emissive background, is an attractive prospect for tracing cell dynamics, trafficking, history, and fate. (1) The HaloTag system is very broadly **established** - it is available in many model systems, with many tissue / cell-type specificities and subcellular compartmentalisations. (2) The covalent ligation of the ligand with the HaloTag protein prevents post-ligation diffusion of the small molecule from the protein (either localised at cellular structures or compartementalised with the HaloTag-POI), thus allowing **durable** labelling and in the case of enzyme-cages enables reliable signal integration as a real-time and cumulative readout for the enzyme activity. (3) Using the **conditional ligation** approach (instead of ligation followed by conditional activation) has a number of unique advantages, which are covered in detail in the paragraphs below: from multiplexing and multipurpose assays (§3), through to successfully harnessing enzyme substrate probe designs (§5-6). (4) The recent disclosure of "xHTL" exchangeable HaloTag ligands (moderate to high affinity but without ligation reactivity[41]) hints at noncovalent **exchangeable CHalo** (XCHalo) systems, simply by replacing its chlorine with e.g. a trifluoromethanesulfonamide, which may be particularly effective in multipurpose assays (e.g. sequential recruitment to different targets).

##### *§3: Multiplexing: Multicolour and Multipurpose uses of CHalo/HaloTag*

By **multiplexing CHalo reagents**, their conditional ligation after activation can, at least conceptually, be used for e.g. **multi-target multi-colour imaging** (e.g. Cage$^n$-CHalo-Dye$^n$ panels with five channels separable via intensiometric imaging (blue, green, yellow, red, far-red), or, more by FLIM). This should durably capture the activities of different enzymes in parallel with single-cell resolution, such that the **relative enzyme activities are quantitatively reflected** in the relative HaloTag-recorded reporter signals (expecting proportionality factors *between* enzymes, which crucially however are *invariant* between different cells at least in 2D cell culture assays). Such relative activity assays have been previously impossible with known small-molecule enzyme activity probes (that suffer from post-activation signal loss by diffusion or export out of cells) or genetic intensiometric sensors (which give signals dependent on their expression levels, and therefore make absolute and relative quantification difficult). These multicolour CHalo/HaloTag assays may also be useful e.g. for fingerprinting to distinguish correlated vs non-correlated enzyme activities within or outside networks, as a marker of normal or dysregulated functions. The multicolour reporters can be direct live cell reporters (fluorogenic reagents), or endpoint readouts (if permanently fluorescent reagents are used, since these need washout before quantification). Our ongoing research aims at multicolour panels to fingerprint and record the relative activities of related oxidoreductases with quantitative accuracy at single cell resolution, that we believe will be able to harness the *in vivo* applicability of HaloTag protein systems to allow panel translation from cell culture to live animal settings.

Multiplexing can also include applications beyond just imaging, e.g. including both reporter reagents and effector reagents (well beyond the one phototriggered heterodimeriser we have shown here) for **multipurpose assays**: e.g. a 550 nm photouncaged XCHalo-based recruiter to CLIP-Tag and a 450 nm photouncaged CHalo-based recruiter to SNAP-tag, for two-colour-two-target sequential recruitments, applied in parallel with 460 nm and 580 nm emissive reporters whose ligations are dependent on enzyme activities and with a 650 nm emissive reporter dependent on a reactive small molecule. Such multipurpose assays can leverage the many suitable "channels" of photocage as well as the diversity of click chemistry, biochemical, and enzymatic cage groups for the CHalo uncaging step, married to the broad range of functional cargos.

##### *§4: Side note on SLPs in the context of photo-triggered fluorogenic labels (expanded from main text):*

Photo-triggered (or photoactivated, or photoconvertible) fluorescent proteins are widely used throughout biology, e.g. paGFP. Chemigenetic counterparts of these protein-only tools have only recently been developed, driven by the need for vastly more photostable and higher-performance fluorophores at custom wavelength regions [which are feasible for small molecule chemistry to access]: but so far these have been restricted to *unconditionally-ligating but photoactivated-fluorescence reagents* (HaloTag-reactive caged-fluorophores, e.g. PaX[42]). Yet, outside their target uses in high-resolution structural imaging of the SLPs' fusion protein targets, these approaches have several limitations for broader chemical biology: e.g., they prevent more powerful assays because all ligation sites are saturated with the pre-fluorophore, so HaloTag becomes a single-purpose anchor; and, during photoactivation, background from non-ligated but photoactivated forms can lower the signal-to-noise, and can diffuse unchecked (all cells' binding sites saturated). A minor point too is that conditional ligation keeps the target protein and fluorophore spatially separate during uncaging, which may reduce on-target photodamage during photouncaging (or, equivalently, while imaging ongoing enzymatic uncaging). The comparative advantages and unique applications of photoactivated-ligation rather than post-ligation-photoactivated-fluorescence systems, with their focus on revealing or harnessing a stimulus or stimulus-defined spatiotemporal location, parallel those discussed above for other prior ligation vs downstream ligation reagent designs.

##### *§5: Generalities on molecular imaging*

**Fluorogenic probes** are crucial tools for visualising and quantifying bioactivity by linearly generating fluorescence upon biological stimuli, from zero background. They are often used for non-invasive bioactivity imaging for enzymes (e.g. peptidases, esterases, phosphatases, glycosidases, and oxidoreductases[3,43–48]) or reactive analytes[3] such as hydrogen peroxide and hydrogen sulfide.[6,7] However, **post-activation signal loss** problematically impairs cell-resolved activity imaging, since it lowers the sensitivity and reliability of signal quantification. There are only a few generalised methods that can increase the **cellular signal retention**: mainly, charge trapping[7,49–51], fluorophore precipitation,[52] or non-specific electrophilic labelling of impermeable biomolecules (SPiDER probes[53]). Each has its specific **disadvantages and limitations**[51], e.g. with charge-trapping, retention is often only moderately improved (ion exporters[54]) and probes require additional delivery strategies; whereas precipitation- or electrophilic-trapping are pro-inflammatory and cell toxic in the longer term (crystal formation / protein alkylation[55]).

The **ideal enzyme activity integrating probe** would be as non-invasive / non-toxic as typical non-retained (soluble) probes but have long-term retention across a range of different cell lines (variable exporter expression profiles etc). We propose that clean, specific, efficient (quantitative), bioorthogonal reaction of a probe product with an introduced innocent intracellular protein target like HaloTag should achieve full product retention without disturbing the *activating target enzyme* or without nonspecific / off-target cell-toxic effects (signal now protected from loss due to small molecule transporters, bioorthogonal reaction allows high dosage without toxic side effects, and the system is immediately *in vivo* compatible since many examples have been shown for expressing HaloTag *in vivo*).

We use CHalo as a modular basis for reagents to image and integrate the bioactivity of an enzyme or reactive biochemical, since *the step which is cell-retaining is the same as the step which is signal generating* (HaloTag ligation step with SiR cargo), and that step is efficient, and entirely conditional on prior CHalo uncaging. This should then allow high-sensitivity imaging and integration with e.g. caged CHalo-SiR fluorogenic reagents. Advantageously, since CHalo's aniline carbamate caging chemistry is compatible with auto-immolative spacers that act as chemical adapters allowing a diversity of chemical reaction types and biochemical / enzyme activators to act as the upstream trigger for uncaging[56] and thus bioactivity integration. The proof of concept for this design principle will be a demonstration via a leucine aminopeptidase (LAP) probe.

##### *§6: SLPs in the context of enzyme-triggered fluorogenic labels and molecular imaging / recording*

There are no generally useful platforms for sensitive fluorogenic chemical probes that allow quantitative *in vivo* enzyme activity integration and localisation, and are cell-retained without impacting native biology. Failings often include e.g. limited signal retention, and biological non-innocence (see ref[51]). A similar concept, briefly mentioned in the main text, has been the use of single-purpose protein-based integrators, like CaProLa[57], that transduce macromolecular rearrangements of the sensor (e.g. a cpHaloTag) which have been caused by cellular activity (e.g. analyte binding) into durable ligation events that can be read out later. These sensors are however laborious to construct and to translate between model systems; they seem adapted to only one activity readout per sensor; and the space of "activity" that they can sense is, probably, mostly separate from the activity space that classic small molecule enzyme substrates can reveal.

CHalo's principle of **<u>triggered ligation</u>** uses enzyme activity to allow a probe product to ligate cleanly and durably to HaloTag: which seems an ideally quantitative, non-invasive, high-sensitivity approach that could harness the diverse HaloTag-expressing animal models already available. The CHalo design could ensure that the enzyme reaction proceeds *unimpeded* on the free Caged-CHalo substrate, resulting in a *rapidly ligating uncaged CHalo product* that is then irreversibly trapped by HaloTag (for accumulating readout and/or sensitive endpoint analysis). This seems a reliable way to convert the decades of knowledge on freely-diffusing small molecule enzyme probes[3] into biologically innocent quantitative integrators. The successful performance of **Leu-CHalo-SiR**, which contains a common PABA 1,6-elimination spacer as a chemical adapter, also promises that other types of chemical adapters (benzylic or other elimination, or cyclisation, etc) which access a huge diversity of enzymatic or chemical reactivities,[56] can be used to patch a diversity of enzyme-uncaged motifs onto the CHalo system without case-by-case needs to re-assess compounds for potential instability or kinetic problems with their uncaging or ligation steps (**Fig. 2ef**).

It is not otherwise obvious how enzyme activity should be used to gate a different step of the process without introducing major problems. The only easily conceivable alternative approach is one that we call **prior ligation**: anchoring a small-molecule activity probe onto HaloTag then generating fluorescence upon reaction of the target enzyme with the HaloTag-probe conjugate. Prior ligation reagents could in principle be created very easily with molecularly simple substrate designs using modules that have been known for decades (e.g. CA-fluorescein-*O*-esters). However, we are unaware of any probes utilising prior ligation of a pro-fluorophore to HaloTag that managed to be effective fluorogenic sensors for enzyme activity. A prior ligation concept would require the activating enzyme to have equally good recognition of the protein-bound substrate motif, as for the freely-diffusing small molecule substrate: which seems highly unlikely. More probably, the HaloTag protein would block or restrict enzyme access to the protein-bound motif, whereas unligated pro-fluorophore would react without hindrance: so suppressing specific bound signal while giving a freely diffusing unwanted background, and in turn, reducing probe sensitivity and utility. Even more problematically, pre-activation anchoring requires the HaloTag and the activating enzyme to reside in the same cellular compartment (since proteins cannot cross intracellular membranes) and even if they are accessible for one another, diffusion of a HaloTag-probe construct to e.g. a plasma membrane bound enzyme of interest would be much slower than a freely diffusible small molecule probe limiting the turnover and thus the sensitivity.

##### *§7: SLPs in the context of chemically induced protein dimerisation (CID)*

Simple reagent examples have tested spontaneous (unconditional) HaloTag-to-SNAP-tag dimerisers that could be cleaved upon illumination,[31] or unconditional HaloTag-to-SNAP-tag dimerisers that target exofacial proteins.[58] Although we did not find assays in those reports that characterise the degree of non-productive monovalent labelling depending on ligand : SNAP : Halo stoichoimetry (also known as the "hook effect" for bifunctional molecules), it seems impossible that those designs would escape this otherwise entirely general problem for saturable systems, which is particularly problematic when stoichiometries are far from equal and the overall concentration is low (i.e. in all but a few special cases).

To overcome the hook effect, photo-activatable protein ligation motifs have instead proven to be invaluable tools in applications around and towards chemically induced dimerisation (CID) reagents. Some early examples tested benzylguanine caging for conditional SNAP-tag ligation[59] (which although not done for SNAP-to-Halo designs, is still conceptually on the same level: see below). However, benzylguanine (BG) caging designs were then essentially abandoned, with few "tool creation" reports and so far without followup "tool use" papers: likely due to the problems of benzylguanine caging chemistry and slow post-uncaging kinetics which have recently been discussed[60] (although the recent report of "second generation" SNAP-tag substrates may revitalise design efforts leveraging SNAP-caged reagents[61]). Matching that analysis, most studies using phototriggered heterodimerisers in the last decade have focused instead on unconditional HaloTag labelling followed by trimethoprim (TMP) photouncaging for noncovalent recruitment of *Escherichia coli* dihydrofolate reductase (eDHFR), allowing conditional protein dimerisation: since the TMP ligand can be more easily caged than benzylguanine[60] and results in very rapid, good-affinity binding to eDHFR (even if it is noncovalent and technically reversible). Such photo-activatable dimerisers and photosplitters (which incorporate an additional photocleavable group in the linker between the two ligand motifs) were used for applications ranging from light-induced recruitment of cytosolic proteins to kinetochores or mitochondria, to controlling peroxisome transport and mitotic checkpoint signalling, to molecular activity painting and studying signalling processes.[33,62–65]

However, to the best of our knowledge, no photocaged double-covalent heterodimeriser has yet been reported. Conceptually, while uncaging Halo ligation had been impossible, photouncaged-Halo-to-SNAP heterodimerisation could have been done before since the SNAP-tag ligand offers not only an aniline-type nitrogen that could be suitable for caging, but also imidazole-type nitrogens that have already been explored for caging, even if both are considered difficult[60] caging sites (and we had indeed worked towards Halo-to-SNAP reagents in early stages of this project before abandoning them). However, any such SNAP-to-Halo designs would have had to photouncage the SNAP ligation step (10-100 fold slower ligation than HaloTag ligation[38]) which seems unpromising for tool utility; and in any case, no photouncaged-SNAP-to-Halo designs were reported to be successful. **Cou-CHalo-BG** instead can perform the slower[38] SNAP-ligation first; then, after washout of unligated reagent, photouncaging unleashes the expectedly faster HaloTag-ligation to complete the heterodimerisation (**Fig. 2j-l**): a labelling order that best retains the spatiotemporal resolution of photouncaging, and which may in the future be adaptable to work with *fluorogenic* HaloTag ligation (i.e. fluorescent turn-on reports on the successful completion of heterodimerisation).

### 8 Additional Data

#### 8.1 SDS-PAGE Gels and Quantification

For full discussion of (caged) HL2$^{N}$ ligands see **Fig. S6**. In **Fig. S11** we additionally show the quantification of in-gel fluorescence from HL2$^{O/N}$ ligands from the SiR-chase and the fluorescein channels as well as the increase of the UV fluorescence (ex.: 365 nm) proving the undesired binding of the caged **Cou-HL2$^{N}$-Fluo**.

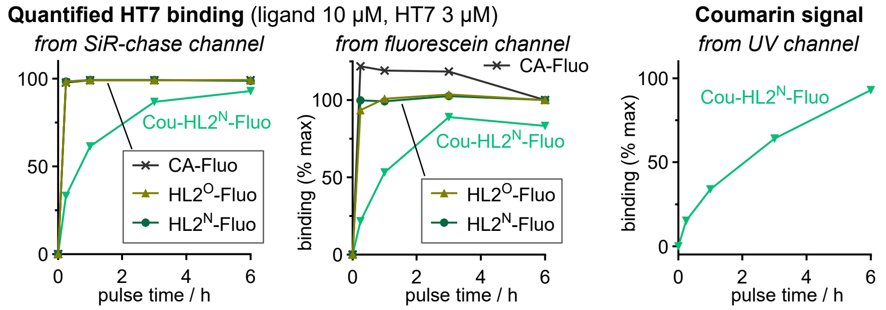

**Figure S11: HL2$^{N}$-Fluo efficiently ligates to HaloTag but ligation of Cou-HL2$^{N}$-Fluo is not blocked by caging:** additional in-gel fluorescence quantification from SiR-chase and fluorescein channel for SDS-PAGE gel shown in **Fig. S6**; the coumarin signal increases upon ligation of **Cou-HL2$^{N}$-Fluo** while no signal increase occurs from free **HL2$^{N}$-Fluo** which is not coumarin-caged (quantified from UV channel, normalised to 93% at 6 h from SiR-chase; incubation with 10 µM ligand and purified HT7 (3 µM) for different times (15 min, 1 h, 3 h, 6 h)).

For full discussion of the HL3 ligand see **Fig. S8**. **Fig. S12** shows the SDS-PAGE gel used for quantifying the HL3 binding and the signal quantification from both the fluorescein and the SiR-chase channels.

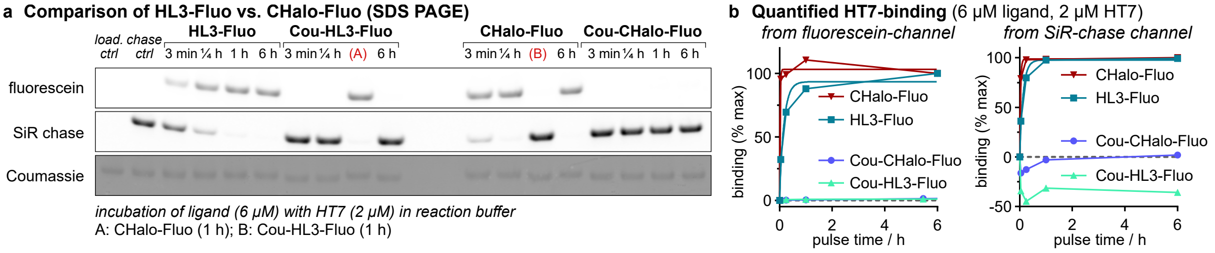

**Figure S12: HL3-Ligand shows slower HaloTag ligation kinetics than CHalo. (a)** SDS-PAGE gel showing the binding efficiencies of **HL3-Fluo** and **Cou-HL3-Fluo** compared to **CHalo-Fluo**, and **Cou-CHalo-Fluo** (6 µM) each after incubation with purified HT7 (2 µM) for different times (3 min, 15 min, 1 h, 6 h); A: **CHalo-Fluo**, 1 h; B: **Cou-HL3-Fluo**, 1 h; all rows show HT7 band: (i) fluorescein fluorescence, (ii) SiR chase fluorescence, (iii) Coumassie staining; **(b)** Quantified HT7-binding of HL3/4-type fluorescein conjugates (quantified from fluorescein or SiR channel in panel b).

The development of the CHalo ligand is discussed in detail at **Fig. S7** together with SDS-PAGE data showing rapid binding of CHalo reagents and fully blocked binding of caged CHalo reagents. In **Fig. S13** we additionally show comparative SDS-PAGE data for CA- and CHalo- conjugates with fluorescein and silicon-rhodamine SiR to assess the relative binding speed by in-gel fluorescence (quantification from fluorescein or SiR fluorescence as well as from MaP555 chase fluorescence). Despite incubation of ligand and HaloTag protein at low concentrations (1.5 : 0.5 µM) the ligation is still too fast to reliably resolve it by in-gel fluorescence (>60% ligation after 3 min) supporting that the CHalo ligand gives very rapid HaloTag ligation (detailed kinetics discussion at **Fig. S10ef**).

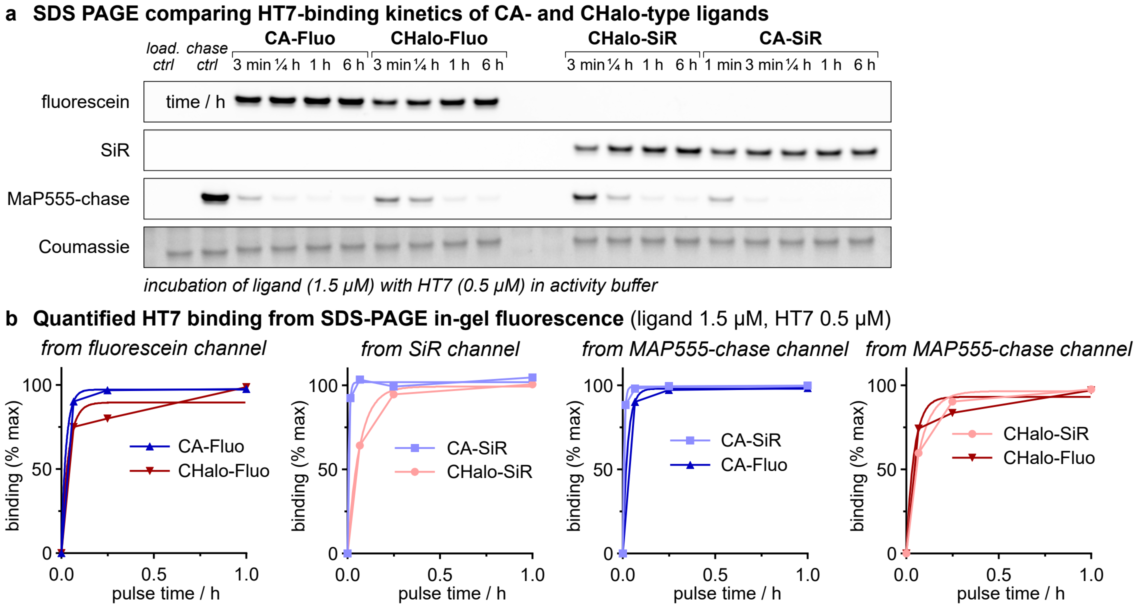

**Figure S13: CHalo-Fluo and CHalo-SiR efficiently ligate to HaloTag (comparison to CA reagents). (a)** SDS-PAGE gel for determining the binding kinetics of the fluorescein conjugates **CA-Fluo** and **CHalo-Fluo** and the SiR conjugates **CA-SiR** and **CHalo-SiR** (1.5 µM) each after incubation with purified HT7 (0.5 µM) for different times (3 min, 15 min, 1 h, 6 h, and after 1 min for **CA-SiR**); all rows show the HT7 band: (i) fluorescein fluorescence, (ii) SiR fluorescence, (iii) MaP555 fluorescence, (iv) Coumassie staining; **(b)** HT7-binding quantified from fluorescein, SiR, and MaP555-chase channel in panel a).

For full discussion of the CHalo binding and how caging blocks binding see **Supplementary Note 2** and discussion at **Fig. S7**. **Fig. S14** shows the SDS-PAGE analysis and in-gel fluorescence quantification of (caged) CHalo fluorescein conjugates (quantified from fluorescein channel and MaP555 chase channel).

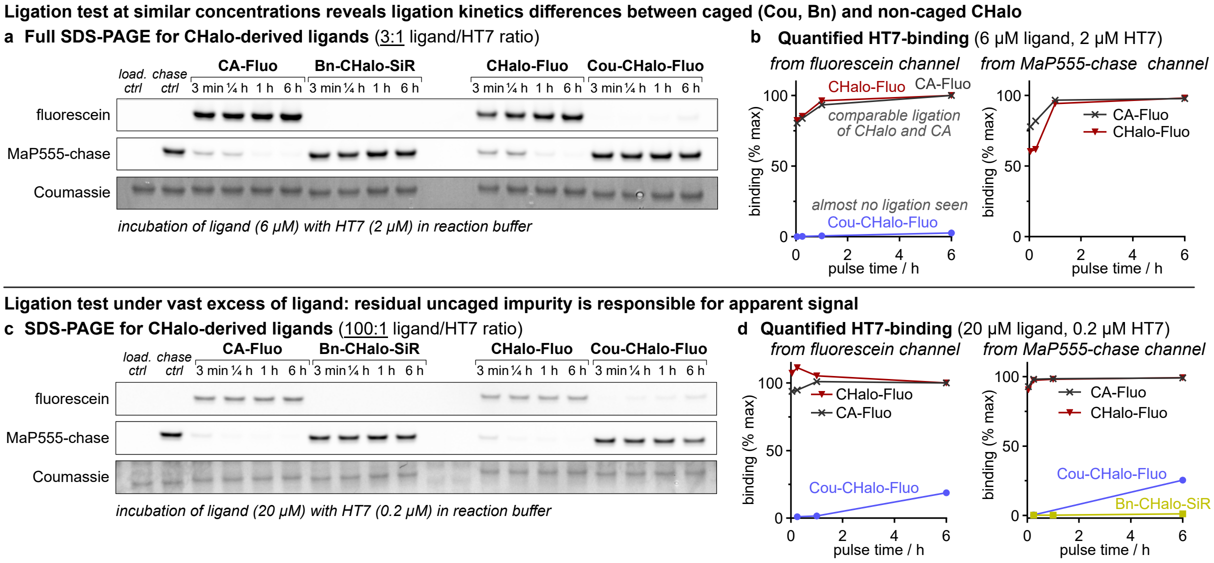

**Figure S14: Caging of CHalo ligand effectively blocks HaloTag ligation. (a)** SDS-PAGE gel showing the binding efficiencies of **CA-Fluo** compared to **Bn-CHalo-SiR**, **CHalo-Fluo**, and **Cou-CHalo-Fluo** (6 µM) each after incubation with purified HT7 (2 µM, *ratio = 3:1*) for different times (3 min, 15 min, 1 h, 6 h); all rows show HT7 band: (i) fluorescein fluorescence, (ii) MaP555 fluorescence, (iii) Coumassie staining (parts of gel shown in **Fig S7c**); **(b)** Quantified HT7-binding of CHalo-type fluorescein conjugates (quantified from MaP555 channel from panel b); **(c)** SDS-PAGE gel showing the binding efficiencies of **CA-Fluo** compared to **Bn-CHalo-SiR**, **CHalo-Fluo**, and **Cou-CHalo-Fluo** (20 µM) each after incubation with purified HT7 (0.2 µM, *ratio = 100:1*) for different times (3 min, 15 min, 1 h, 6 h); all rows show HT7 band: (i) fluorescein fluorescence, (ii) MaP555 fluorescence, (iii) Coumassie staining; **(d)** Quantified HT7-binding quantified from fluorescein and MaP555 chase channel from panel d).

#### 8.2 Fluorescence Properties of Fluorescein Conjugates

For the proof-of-concept studies, we conjugated fluorescein to the HaloTag ligands as permanently fluorescent markers to trace ligation by in-gel fluorescence. All conjugates show the expected absorption and emission maxima (ca. 500/520 nm) with extinction coefficients 40–80·10³ M⁻¹ cm⁻¹ (**Fig. S15**, **Table S1**). The quantum yields, however, strongly differed: highest fluorescence is observed for the ethylene glycol ligand **CA** (Φ = 0.83) which is strongly reduced for the phenyl group containing ligand **HL2$^{O}$** (Φ = 0.28) and even lower for the aniline ligands **HL2$^{N}$** and **CHalo** (Φ ca. 0.15) which might be explained by photoinduced electron transfer partially quenching the fluorescence.[66] The quenching seems even stronger when another quencher is installed in the **Cou-HL2$^{N}$** and **Cou-CHalo** ligands (Φ ca. 0.05) giving a 15-fold lower quantum yield compared to **CA-Fluo**. Interestingly, our fluorescein aryl-chain conjugates revealed small fluorogenicity upon HaloTag ligation that is coherent with suppressed PET quenching efficiency once ligation fixes the chain arene out of contact with the fluorophore (reminiscent of the larger unquenching seen with rhodamines), whereas alkyl-chain **CA-Fluo** shows reduced fluorescence upon binding (potentially due to unfavourable interactions of the normally anionic fluorescein with the protein surface that is best evolved for cationic rhodamines).

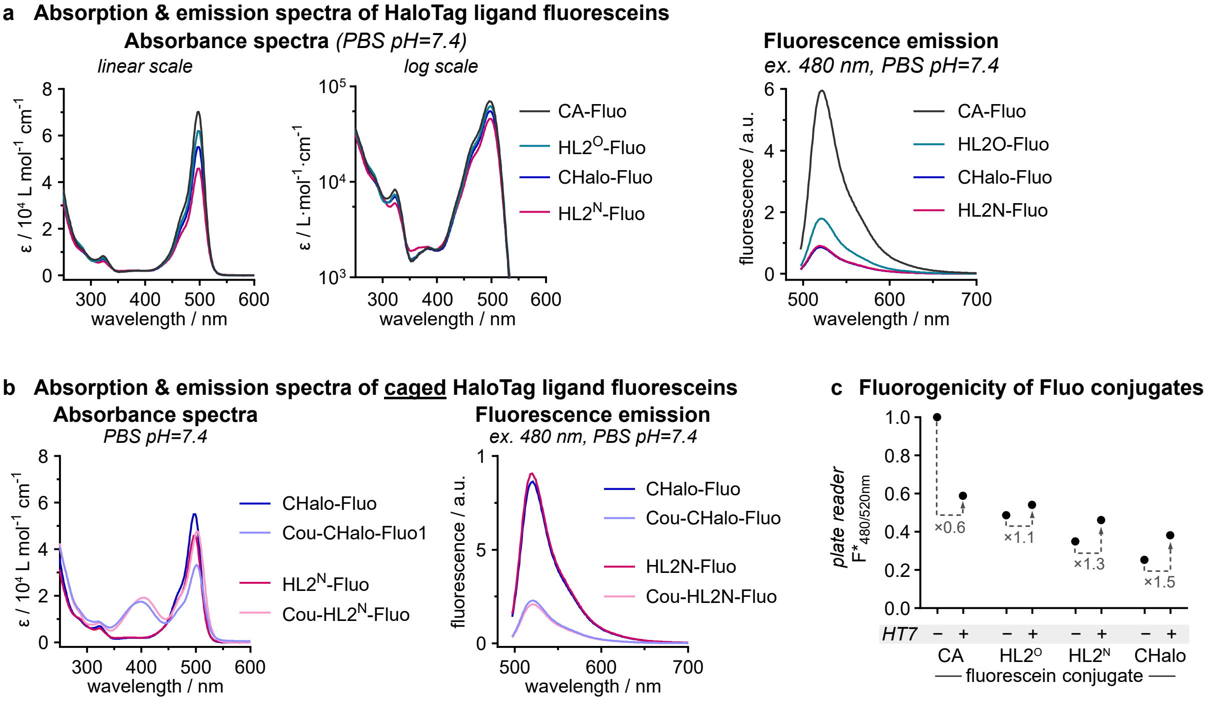

**Figure S15: Absorption and fluorescence properties of HaloTag ligand fluorescein conjugates. (a)** Absorption and emission spectra of **CA-**, **HL2$^{O}$-**, **HL2$^{N}$-** and **CHalo-Fluo** (10 µM, PBS, pH = 7.4); **(b)** Absorption and emission spectra of coumarin-caged conjugates **Cou-HL2$^{N}$-Fluo** and **Cou-CHalo-Fluo** (10 µM, PBS, pH = 7.4); **(c)** Fluorescence emission differences of free and HaloTag-bound Fluo-conjugates (2 µM ligand, 4 µM HT7, activity buffer, 15 h).

| Fluorophore | $\lambda_{Abs,max}$ / nm | $\lambda_{emission,max}$ / nm | Stokes Shift / nm | $\varepsilon_{\lambda,max}$ / L mol⁻¹ cm⁻¹ | $\Phi_{FL}$ | brightness / L mol⁻¹ cm⁻¹ |
|---|---|---|---|---|---|---|
| CA-Fluo | 497 | 522 | 25 | 70·10³ | 0.83 | 59·10³ |
| HL2$^{O}$-Fluo | 498 | 521 | 23 | 62·10³ | 0.28 | 17·10³ |
| HL2$^{N}$-Fluo | 498 | 520 | 22 | 46·10³ | 0.17 | 7.7·10³ |
| Cou-HL2$^{N}$-Fluo | 502 | 521 | 19 | 48·10³ | 0.04 | 1.9·10³ |
| CHalo-Fluo | 498 | 521 | 23 | 55·10³ | 0.14 | 7.8·10³ |
| Cou-CHalo-Fluo | 501 | 520 | 19 | 33·10³ | 0.06 | 1.8·10³ |

**Table S1**: Fluorescence properties of fluorescein conjugates **as free (unbound) ligands, in aqueous buffer** (PBS, pH = 7.4).

Quantum yields of the novel fluorophores were determined the following equation (Resch-Genger and coworkers[67]), albeit with the refractive indices cancelling each other (same medium):

$$\Phi_{f,x} = \Phi_{f,st} \cdot \frac{F_x}{F_{st}} \cdot \frac{1 - 10^{-A_{st}(\lambda_{ex})}}{1 - 10^{-A_x(\lambda_{ex})}} \cdot \frac{n_x(\lambda_{em})^2}{n_{st}(\lambda_{em})^2}$$

Fluorescein was used as a reference fluorophore with a quantum yield of $\Phi_{f,st}$ = 0.85 (in PBS, pH=7.4).[68]

#### 8.3 HaloTag specificity

We supported the HaloTag-specificity of **CHalo-Fluo** ligation by competitive incubations of the ligand (5 µM) with HaloTag (0.1 µg, 0.5 µM) and cell lysates (up to 10 µg cellular protein). In-gel fluorescence indicated that the ligand only binds to HaloTag even though at least 4.5 µM of unreacted ligand remains exposed to cell lysate proteins (**Fig S16a**).

Relatedly, incubating the fluorogenic **CHalo-SiR** with cell lysate only (no HaloTag protein) shows no unwanted fluorescence generation, supporting that no undesired covalent or non-covalent interactions with the SiR conjugate occur that would generate off-target fluorescence (**Fig S16b**).

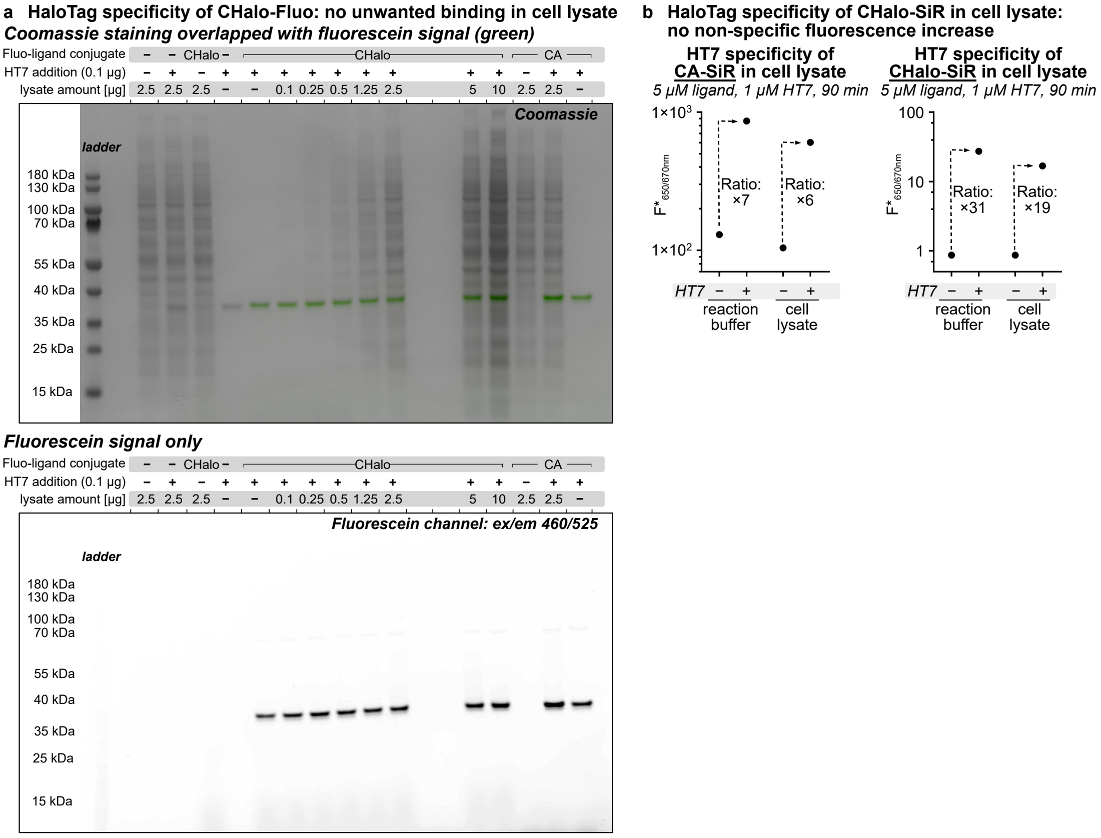

**Figure S16: HaloTag specific binding of CHalo-Fluo in cell lysate. (a)** SDS-PAGE gel of **CHalo-Fluo** with different amounts of HeLa cell lysate with and without HT7 (treatment of 5 µM **CHalo-Fluo** for 1 h with or without HT7 (0.5 µM), comparison: **CA-Fluo**); **(b)** Fluorescence generation of **CA-SiR** and **CHalo-SiR** with reaction buffer or with HeLa cell lysate with and without HT7 (treatment of 5 µM ligands for 1.5 h with 1 µM HT7).

#### 8.4 Localisation of non-specific CHalo-SiR background in cells

The fluorescence of **CHalo-SiR** reagents is mostly, but not entirely, suppressed before ligation to HaloTag. Live cell imaging experiments showed minor non-specific background fluorescence of **CHalo-SiR** reagents in cells without HaloTag ligation (compare **Fig S18**: Leu-CHalo-SiR without HaloTag). Treatment of U2OS cells without HaloTag showed SiR signal in mitochondria, nucleus, and endoplasmic reticulum (ER) (**Fig. S17**). This localisation stems from the SiR dye cargo (not an intrinsic feature of the CHalo motif logic gate), which is logical since lipophilic, delocalised cations like rhodamines and triphenylphosphoniums are known as mitochondrial targeting groups with some ER localisation.[69]

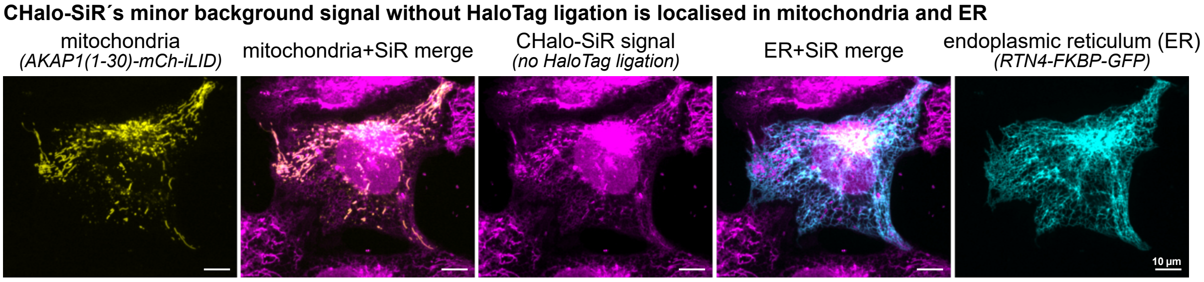

**Figure S17: CHalo-SiR gives minor non-specific fluorescence (no HaloTag ligation) in cells that is localised in mitochondria and ER.** Incubation of U2OS cells (no HaloTag) with **CHalo-SiR** (5 µM) for 20 min.

#### 8.5 Additional Data for Leu-CHalo-SiR

##### 8.5.1 HaloTag dependent signal generation and post-wash retention

We used the CHalo system for durable enzyme-activity imaging by uncaging **Leu-CHalo-SiR** with leucine aminopeptidase (LAP) followed by fluorogenic HaloTag ligation of the released **CHalo-SiR**. **Leu-CHalo-SiR** enables very sensitive detection due to its full fluorescence suppression before binding and the >400-fold fluorogenicity upon ligation (**Fig. 1h-j**). Cruicially, **caged-CHalo-SiR** cannot ligate to HaloTag (**Fig. 2f:** no signal for **Bn-CHalo-SiR**) and the signal generation from the released probe is fully dependent on the ligation to HaloTag (**Fig S18a**: no signal in cells not expressing HaloTag). After LAP activation (leucine peptidolysis, then spontaneous 1,6-elimination of the carbamic acid anion from the *para*-aminobenzyl spacer, then its spontaneous $CO_2$ evolution to give the uncaged CHalo-SiR reagent) followed by ligation to HaloTag protein, the HaloTag----CHalo-SiR conjugate is now membrane-impermeable and intracellularly retained. The actual SiR signal is limited by the rate of degradation of the HaloTag fusion protein employed, which depends on the cell line and protein target.[1] Nonetheless, signal integration by HaloTag ligation over hours to days is a feasible approach, even *in vivo* as shown by the calcium signal recorder CaProLa in flies.[70]

**Fig. S18b** shows the post-wash retention of the SiR signal in the specific settings we used (cytosolic HaloTag-mScarlet target, HEK cells): both unconditional ligators (CA-SiR and **CHalo-SiR**) are well retained for 2 h; but their signal decreases by 85% over the course of 24 h. We conjecture that the HaloTag-mScarlet fusion protein has a turnover half-life of ca. 8 h, explaining that signal loss (plausible rate). Such signal reduction was not observed for **Leu-CHalo-SiR** which showed bright cellular signal even after 48 h. We do not attribute that observation to retaining the ligated product from time zero, but rather to retention of the unligated probe that ligated later, allowing continuous signal generation after washing - although this still indicates remarkable detection sensitivity at presumably very low concentrations (although the retention is likely here fostered by the leucine trigger, and therefore cannot be expected with other trigger-CHalo probes). Nonetheless, since protein half-life can be tuned or chosen to fit desired applications, we see clear potential for more-stable (or faster-turnover) HaloTag protein anchors. (*Images used in quantification are shown in* ***Figs. S20−25****)*.

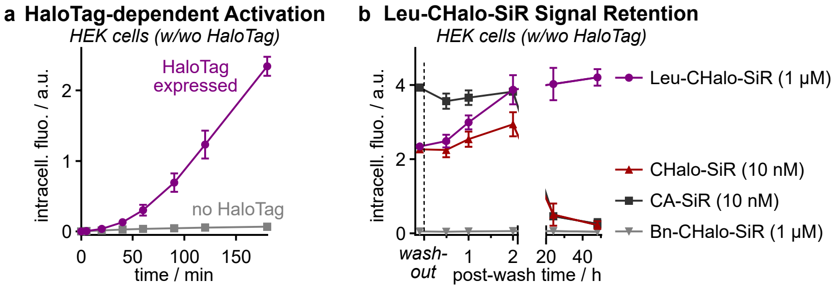

**Figure S18**: **(a)** HaloTag dependent **Leu-CHalo-SiR** signal generation in cells (1 µM, HEK cells with or without cytosolic HaloTag-mScarlet protein; data quantified from microscopy images); **(b)** Intracellular signal retention after removing the SiR reagent (2× medium change, pre-incubation of HEK cells with cytosolic HaloTag-mScarlet with each ligand for 3 h; data quantified from microscopy images; *n*=3).

##### 8.5.2 Full data for Figure 2e

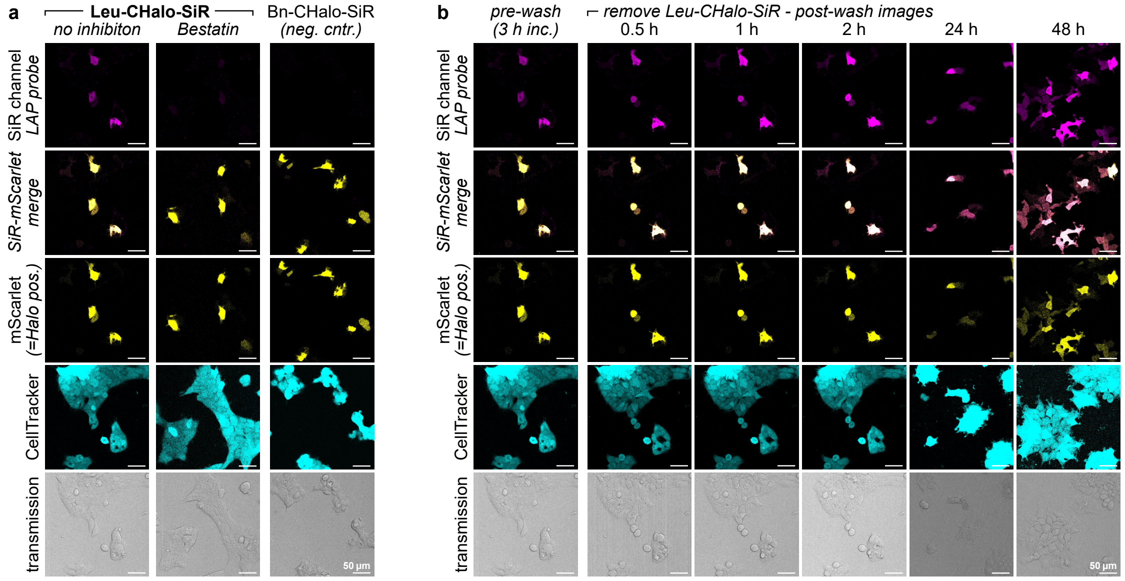

**Figure S19**: Full microscopy data for main **Fig. 2e** (HEK cells, 1 µM probe, 3 h, then 2× wash; optional pre-treatment with bestatin (LAP inhibitor, 100 µM, 30 min)).

##### 8.5.3 Microscopy images for fluorescence quantification

**Leu-CHalo-SiR** (1 µM)

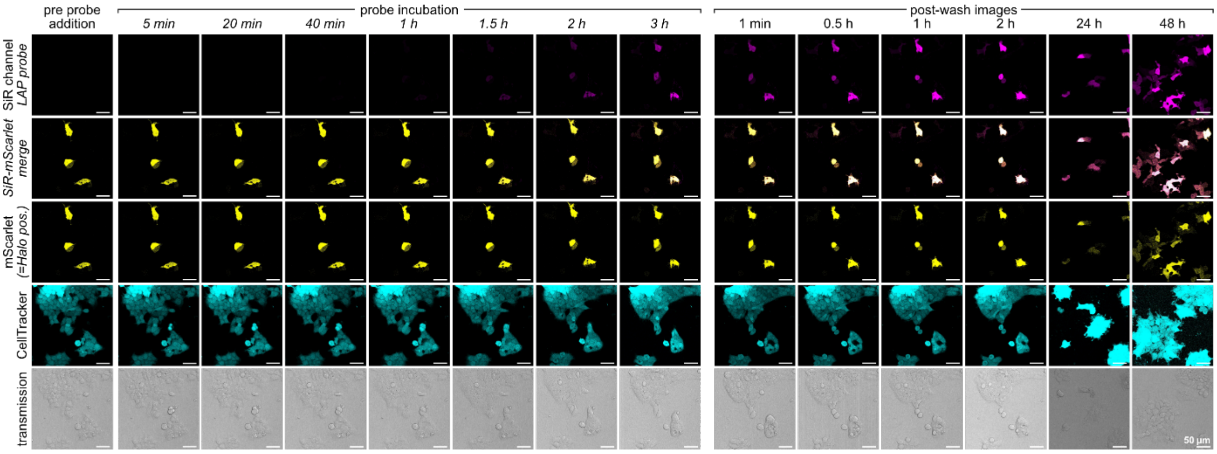

**Figure S20**: Microscopy images for fluorescence quantification for **Leu-CHalo-SiR** (1 µM probe in HEK cells transfected with cytosolic HaloTag-mScarlet; 3 h probe incubation, then 2× wash and post-wash imaging for 48 h; images from one of three replicates which were used for quantification).

**Leu-CHalo-SiR + LAP inhibitor Bestatin** (1 µM probe, pre-treatment with 100 µM Bestatin for 30 min)

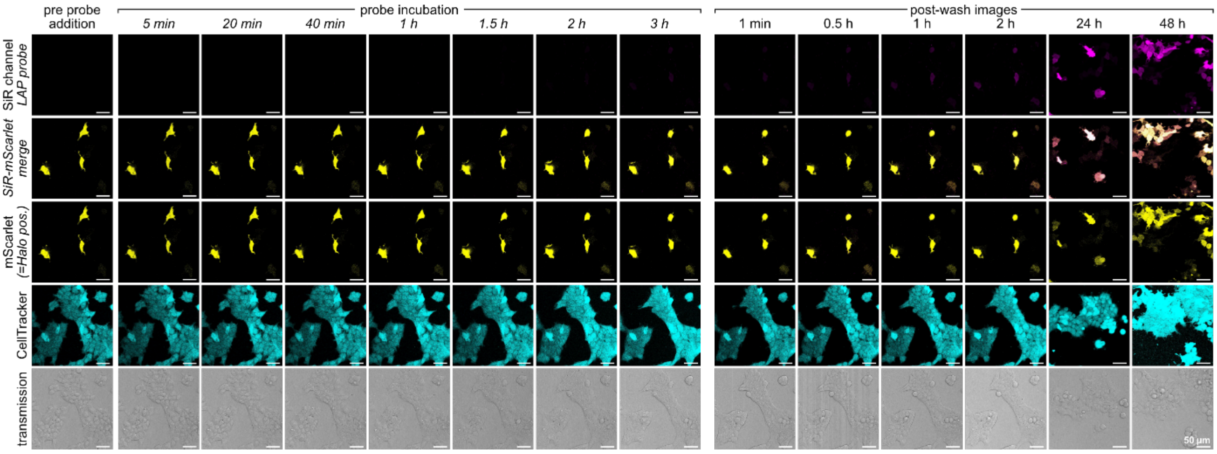

**Figure S21**: Microscopy images for fluorescence quantification for **Leu-CHalo-SiR with LAP inhibitor bestatin** (pre-treatment with bestatin (100 µM, 30 min); then 1 µM probe in HEK cells transfected with cytosolic HaloTag-mScarlet; 3 h probe incubation, then 2× wash and post-wash imaging for 48 h; images from one of three replicates which were used for quantification). These can be interpreted as: washout removes bestatin, but only partially removes **Leu-CHalo-SiR**.

**Bn-CHalo-SiR** (1 µM)

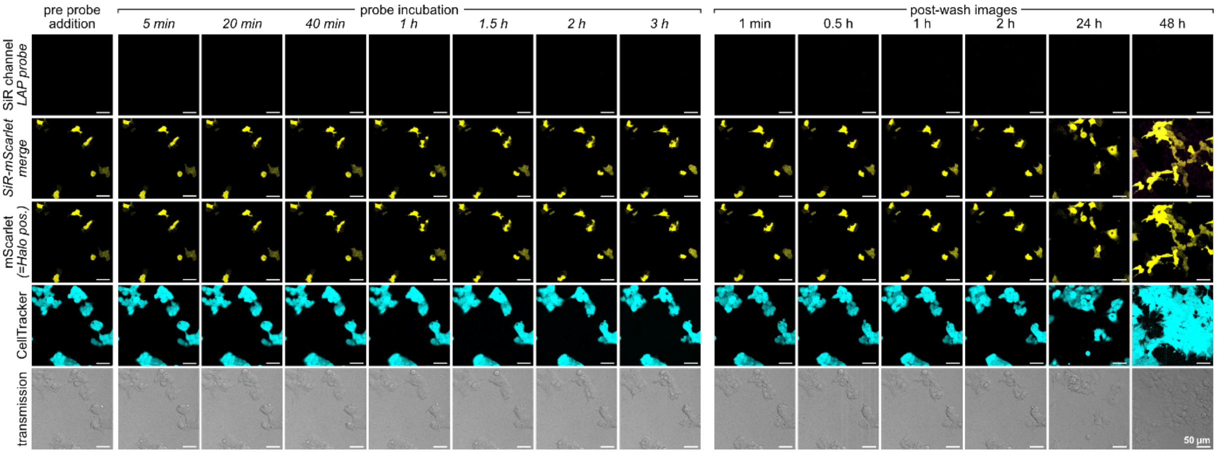

**Figure S22**: Microscopy images for fluorescence quantification for **non-activatable Bn-CHalo-SiR** (1 µM probe in HEK cells transfected with cytosolic HaloTag-mScarlet; 3 h probe incubation, then 2× wash and post-wash imaging for 48 h; images from one of three replicates which were used for quantification).

**HEK cells without HaloTag-mScarlet, Leu-CHalo-SiR** (1 µM)

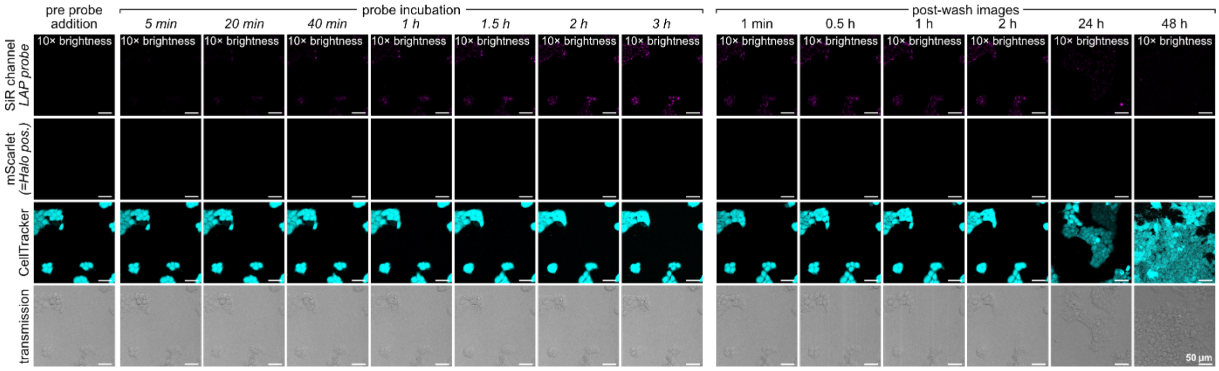

**Figure S23**: Microscopy images for fluorescence quantification for **Leu-CHalo-SiR in HEK cells without HaloTag-mScarlet** (1 µM probe; 3 h probe incubation, then 2× wash and post-wash imaging for 48 h; SiR brightness 10× enhanced compared to the images for the other probes to make the minor background fluorescence visible; images from one of three replicates which were used for quantification).

**CHalo-SiR** (10 nM)

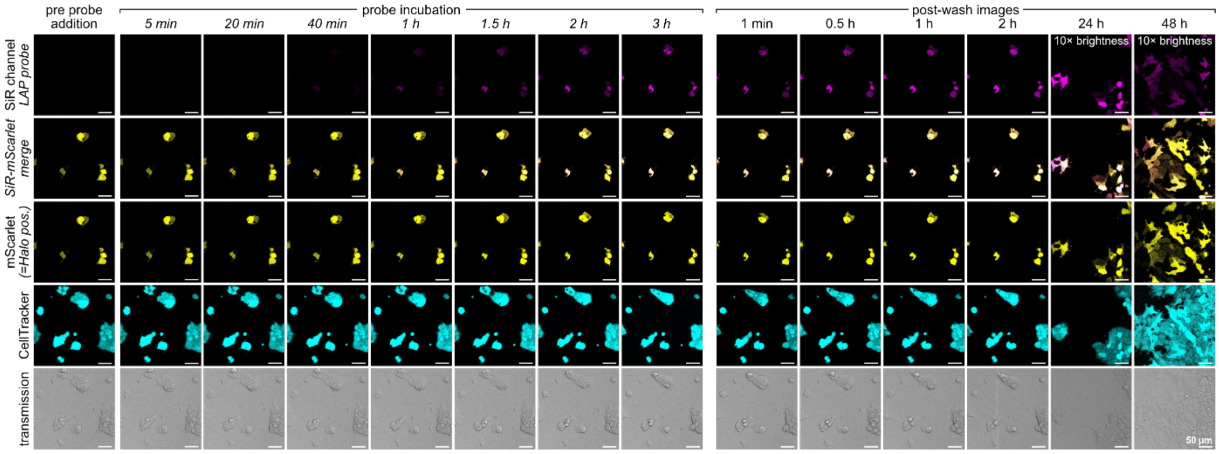

**Figure S24**: Microscopy images for fluorescence quantification for **free CHalo-SiR** (10 nM probe in HEK cells transfected with cytosolic HaloTag-mScarlet; 3 h probe incubation, then 2× wash and post-wash imaging for 48 h; images from one of three replicates which were used for quantification).

**CA-SiR** (10 nM)

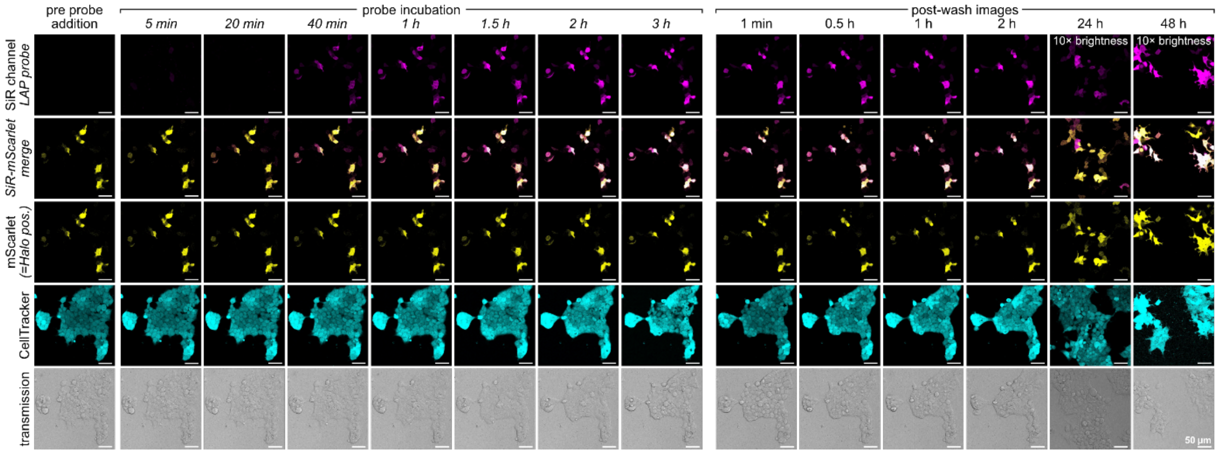

**Figure S25**: Microscopy images for fluorescence quantification for **known CA-SiR** (10 nM probe in HEK cells transfected with cytosolic HaloTag-mScarlet; 3 h probe incubation, then 2× wash and post-wash imaging for 48 h; images from one of three replicates which were used for quantification).

#### 8.6 Additional Data for Cou-CHalo-BG

**Cou-CHalo-BG** can be efficiently uncaged with violet/blue light (ca. 50% conversion with 10 mJ/mm$^2$ at 405 nm, **Fig. S26a**, panel 3) while being unaffected by GFP imaging (no uncaging above 470 nm) as expected for aminocoumarin cages.

We quantified the co-localisation upon SNAP-Halo dimerisation (see main **Fig. 2l**) using Mander's coefficient (**Fig. S26b**) showing rapid and strong increase of the signal co-localisation upon light-activation of SNAP-ligated **Cou-CHalo-BG** (pre-incubation for 1 h, then light-activation, imaging after 5 min).

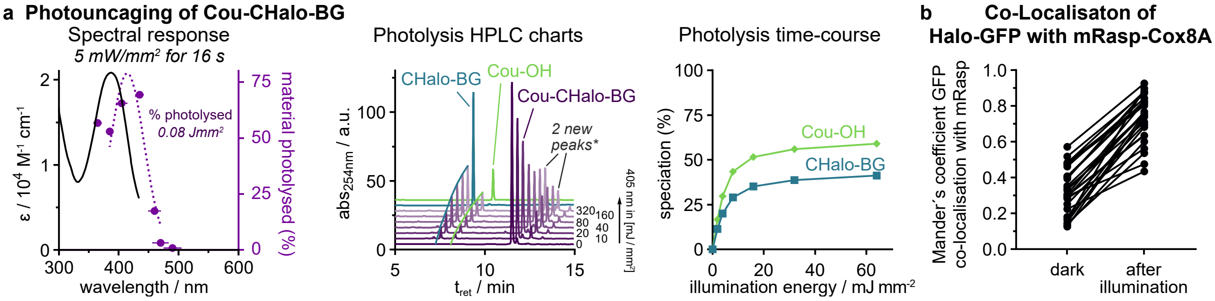

**Figure S26: (a) Cou-CHalo-BG** can be uncaged efficiently with blue light (ideal: 400−440 nm) and GFP orthogonally (no uncaging >470 nm; 50 µM sample in DMSO:water 7:3; illumination with the same light intensity for each wavelength, *horizontal error bars: FWHM of excitation light*; HPLC speciation and photolysis time-course (50 µM in MeCN:water 1:1; illumination with 405 nm light, 5 mW/mm$^2$, applied for times from 1 to 64 seconds (factor 2 steps)); **the species in the “2 new peaks” close to the Cou-CHalo-SiR signal include the coumaryl-CHalo alkyl aniline which is a nonphysiological by-product of the high concentrations used in this assay (after an aniline has been photoliberated, it can trap another photo-generated coumarinyl unit: byproduct mass lacks $CO_2$)*; **(b)** Halo-SNAP dimerisation efficiency quantified by Mander´s coefficient of GFP co-localisation with mRaspberry (quantification for main **Fig. 2l**; each data point represents one cell; *n* = 3).

### 9 Methods: photochemical and biological characterisation

#### 9.1 Photochemical characterisation

##### Absorption spectroscopy

*(several figures, indicated at specific methods section)*

UV-Vis spectra were recorded on a Cary 60 UV-Vis spectrophotometer from *Agilent Technologies Inc*., Santa Clara (USA) using 1 cm quartz or PMMA cuvettes. The scan rate was set to 600 nm/min and 2.5 nm slit width was used. Unless stated otherwise, the probes and fluorophores were dissolved in PBS (pH = 7.4, ≤1 % DMSO). All spectra of CHalo ligands with purified HaloTag7 protein were performed in reaction buffer (25 mM PBS (PanReac AppliChem, A0965.9010), 150 mM NaCl (Bernd Kraft, 04610.2600), 0.5 g/l BSA (Carl Roth, 3737.2) (pH = 7.5)).

##### Fluorescence spectroscopy

*(several figures, indicated at specific methods section)*

Fluorescence spectroscopy was performed on a Cary Eclipse Fluorescence Spectrometer from *Agilent Technologies Inc*., Santa Clara (USA) using quartz cuvettes (scan rate: 120 nm/min, 5 nm slit width) or on a Tecan Infinite M1000 plate reader. All spectra of CHalo ligands with purified HaloTag7 protein were performed in reaction buffer (25 mM PBS (PanReac AppliChem, A0965.9010), 150 mM NaCl (Bernd Kraft, 04610.2600), 0.5 g/l BSA (Carl Roth, 3737.2) (pH = 7.5)).

##### Photouncaging efficiency (HPLC analysis)

*(Figures: S7e, S10k, S26)*

*Instrument*: Analytical HPLC analysis was conducted using an Agilent 1100 SL system equipped with a binary pump to deliver water:acetonitrile eluent mixtures containing 0.1% formic acid at a 1 mL/min flow rate, an Agilent 1100 series diode array detector, and a Hypersil Gold HPLC column. The eluent was a mixture of water (analytical grade, 0.1 % formic acid) and MeCN (analytical grade, 0.1 % formic acid).

The samples were illuminated in 384-well plate wells (50 µM, 50 µL, solvent: mixtures of acetonitrile/water 1:1 or DMSO/water 7:3) using a pE-4000 illumination system (CoolLED) set to 5 mW/cm$^2$. The pE-4000 illumination system was attached to a liquid light guide for light output that enables convenient handling and output uniformity in the illuminated wells. The liquid light guide was placed directly over the well to ensure uniform and comparable illumination conditions.

#### 9.2 Cell-free biological characterisation

##### SDS-PAGE

*(several figures, indicated at methods for pulse-chase assay & HaloTag specificity)*

Samples were denatured in NuPAGE LDS-buffer 4× (Invitrogen, NP0007) and NuPAGE DTT buffer 10× (Invitrogen, NP0004) at appropriate volumes and heated to 70 °C in a Biometra TSC ThermoShaker (Analytik Jena) for 10 min. Gels (NuPage 4−12%, Bis-Tris Midi-Protein-Gel (Invitrogen, WG1402BOX)) were run using a SureLock Tandem Midi Gel Tank (Thermo Fisher, STM1001) and NuPAGE MES running buffer (Invitrogen, NP0002) at 165V for 40 min using a PowerEase Touch 350W Power Supply (Thermo Fisher, PS0350). 5 µL of PageRuler protein ladder (thermos scientific, 26616) and 10 µL of the samples were added to the wells. The in-gel fluorescence was imaged on an Amersham ImageQuant 800 Fluor (Cytiva) using the Cy2-channel (ex: 460 nm, em: 525bp20) for fluorescein ligands, Cy3-channel (ex: 535 nm, em: 605bp40) for MaP555-ligand and Cy5-channel (ex: 635 nm, em: 705bp40) for SiR-ligands; UV (ex. 365 nm, Cy3(UV)-filter) was used for coumarin fluorescence. The total protein stain was imaged after staining the gel with InstantBlue (abcam, ab119211) for 1 h.

Gels were evaluated using Bio-Rad Image Lab Software (quantification of absorption/fluorescence intensity), Microsoft Excel and GraphPad Prism. Quantified fluorescence intensities were corrected for the total protein amount quantified from InstantBlue staining to account for differences in the loaded amount protein. The quantified values from the fluorescein channel were normalised to the value after 6 h incubation (to the corresponding free ligand for Cou-caged fluoresceins), values from the chase channel gave the percentage of chase binding and were transformed to percentage of fluorescein ligand binding (100%−[percentage chase]). The coumarin binding of **Cou-HL2$^{N}$-Fluo** was quantified from the UV signal (subtracted by background from fluorescein determined from **HL2$^{N}$-Fluo**, normalisation of 6 h time-point to percentage calculated from chase-channel). For low ligation of Cou-CHalo reagents, further corrections were performed (see **Supplementary Note 2**).

##### HaloTag7 expression and purification

*(preparation for all experiments using purified HaloTag7 protein)*

<u>Method A</u>: The HT7 plasmid pET51b-His-TEV-HaloTag7 was a gift from Kai Johnsson (Addgene plasmid # 167266; http://n2t.net/addgene:167266 ; RRID:Addgene_167266), transformed into C41 (DE3) E. coli (Sigma-Aldrich, CMC0017) and plated onto 2xYT agar (Carl Roth, X967.1) with 100 µg/mL ampicillin (Carl Roth, K029.1). After incubation at 37 °C overnight a single colony was picked, inoculated in 13 mL tubes (Sarstedt, 62.515.006) with 8 mL 2xYT medium (Carl Roth, X966.1) supplemented with 100 µg/mL ampicillin (Carl Roth, K029.1) and incubated at 37 °C 200 RPM overnight. 100 µL were then added into 10 mL 2xYT medium supplemented with 100 µg/mL ampicillin in 100 mL Erlenmeyer flasks. Expression was induced by adding 0.5 mM IPTG (Carl Roth, 2316.4) and 1 mM $MgCl_2$ (Merck, M8266) at an $OD_{600}$ of 0.8. After incubation at 37 °C overnight cultures were pelleted for 10 min (4800g). Bacteria were lysed for 20 min in protease inhibitor cocktail buffer (Sigma-Aldrich, I3911), DNAse (PanReac, A3778), lysozyme (Carl Roth, 8259.2) and Bugbuster 10× protein extraction reagent (Merck Millipore, 70921) and the lysate clarified by centrifuging for 15 min (20.000g). The proteins were purified using a HisTrap HP 1 mL (Cytiva, GE29-0510-221) on an Äkta Pure 25 M (Cytiva) with 50 mM Tris (Carl Roth, 4855.2), 500 mM NaCl (Bernd Kraft, 04610.2600), 250 mM imidazole (tcichemicals, I0001) elution buffer (pH = 7.5) and a HiLoad Superdex 200 PG 16/600 column (Cytiva, GE28-9893-35) with 25 mM Tris (Carl Roth, 4855.2), 1 mM EDTA (VWR, 7125.1000), 1 M NaCl (Bernd Kraft, 04610.2600) elution buffer. Proteins were assessed by SDS-PAGE and then transferred into 25mM PBS (PanReac AppliChem, A0965.9010), 150 mM NaCl (Bernd Kraft, 04610.2600) buffer (pH = 7.5) using zeba 7k MWCO spin columns (thermo scientific, 89882), concentrated using an Amicon Ultra centrifugational filter (merck Millipore, UFC8010) and stored at 4°C until use.

<u>Method B</u>: *E. coli BL21* were transformed with pET51b-His-TEV-HaloTag7 (Addgene #167266; gift from Kai Johnsson) and plated on LB agar supplemented with 100 µg/mL ampicillin. After overnight incubation at 37 °C, a single colony was used to inoculate 30 mL of LB medium containing 100 µg/mL ampicillin and grown at 37 °C, 200 rpm. 10 mL of the culture was put into 700 mL LB medium containing 100 µg/mL ampicillin and grown at 37 °C, 200 rpm until OD600 of 0.6. Protein expression was induced with 0.5 mM IPTG and cultures were incubated overnight at 28 °C. Cells were harvested by centrifugation (6.000 × g, 10 min) and pellets were resuspended in Lysis buffer (50 mM HEPES, 300 mM NaCl, 10 mM Imidazol, pH 7.4) and the slurry was subjected to cell lysis (LM10 microfluidizer; 15000 PSI, 5 passes, on ice). The lysate was clarified by centrifugation (10,000 × g, 20 min). The supernatant was applied to Ni-NTA Sepharose Fast Flow resin (Qiagen) and bound proteins were eluted with buffer containing 280 mM imidazole, 50 mM Tris, 300 mM NaCl, pH 7.4. Fractions containing the target protein were pooled and subjected to size exclusion chromatography (SEC) on a HiLoad Superdex 200 16/60 column (Cytiva) equilibrated in 25 mM HEPES, 150 mM NaCl, pH 7.5. The major elution peak, corresponding to HaloTag7, was identified by SDS-PAGE. Peak fractions were concentrated using 10 kDa MWCO centrifugal filters and further purified on a Superdex 75 Increase 10/300 GL column (Cytiva). Final peak fractions were pooled, concentrated, aliquoted, and stored at 4 °C. Protein purity was confirmed by SDS-PAGE. Thermal stability was assessed by nanoDSF (Prometheus NT.48, NanoTemper), giving a melting temperature ($T_m$) of ~63°C.

##### HaloTag dependent fluorogenicity

*(Figures: SiR: S10b−d; fluoresceins: S15a−c)*

The fluorogenicity of the ligand-**SiR** conjugates was determined using a Cary Eclipse Fluorescence Spectrometer (excitation: 652 nm, scan rate: 120 nm/min, 5 nm slit width). Purified HaloTag 7 protein (4 µM) and the respective ligand (**CA-SiR** and **CHalo-SiR**, 2 µM respectively) were mixed in reaction buffer (25 mM PBS (PanReac AppliChem, A0965.9010), 150 mM NaCl (Bernd Kraft, 04610.2600), 0.5 g/l BSA (Carl Roth, 3737.2) (pH = 7.5)) and incubated until binding was completed (maximum fluorescence intensity is reached: 15 min for **CA-SiR**, 8 h for **CHalo-SiR**). Then the ligand-SiR conjugates were incubated in reaction buffer (2 µM) without HaloTag 7 protein for the same time and the fluorescence was measured. The turn-on ratio was determined by the fluorescence difference of dye with or without HaloTag 7 at emission 665−672 nm. Absorption spectra were measured on a Cary 60 UV-Vis spectrophotometer as described above.

The fluorogenicity of the ligand-**fluorescein** conjugates was determined using a Tecan Infinite M1000 plate reader (ex: 480 nm; em: 488−700 nm). The fluorescein ligand-conjugates (**HL1-Fluo**, **HL2$^{O}$-Fluo**, **HL2$^{N}$-Fluo** and **CHalo-Fluo**, 2 µM concentration) were incubated for 15 h with or without purified HaloTag 7 protein (4 µM) in reaction buffer (25 mM PBS (PanReac AppliChem, A0965.9010), 150 mM NaCl (Bernd Kraft, 04610.2600), 0.5 g/l BSA (Carl Roth, 3737.2) (pH = 7.5)). The fluorescence emission was normalised to the signal of HL1-Fluo at 520 nm and the turn-on ratio was determined from the signal ratio of dye with or without HaloTag 7 at 520 nm. Absorption and emission spectra (ex: 480 nm) in PBS (pH = 7.4) were recorded as described above.

##### Dielectric constant dependent UV-Vis absorbance

*(Figure S10e)*

**CHalo-SiR** (0.5 µM) was dissolved in water/1,4-dioxane mixtures containing 10−80vol% 1,4-dioxane (1% DMSO) and the absorbance at 645−655 nm was measured at ambient temperature (23 °C). The absorbance was normalised to the highest value and plotted against the dielectric constant of the solvent mixture.[71] The fraction of the fully open (zwitterionic) form was calculated as the fraction of the HaloTag ligated CHalo-SiR absorbance ($7.3 \cdot 10^{4}$ $M^{-1}$ $cm^{-1}$).

##### Pulse-chase assays

*(Figures: S6de, S7cd, S8b, S10h, S11, S12ab, S13ab, S14a−d)*

HaloTag7 (HT7) protein was incubated with the tested ligand for different times, after which a commercially available, fast-binding HT7 ligand was added in excess to stop the reaction and bind to the remaining unbound HT7 protein. All assays working with photocaged ligands were performed in a darkened room and whenever possible covered with aluminium foil. To compare the binding kinetics of **Cou-CHalo-Fluo** (photocaged), **CHalo-Fluo**, **CA-Fluo** and **BnCHalo-SiR** to HT7 protein, assays were performed with 2 µM HT7 and 6 µM ligand or with 0.2 µM HT7 and 20 µM ligand. Protein and ligand were mixed in reaction buffer (25 mM PBS (PanReac AppliChem, A0965.9010), 150 mM NaCl (Bernd Kraft, 04610.2600), 0.5 g/l BSA (Carl Roth, 3737.2) (pH = 7.5)) and incubated at room temperature for 3, 15, 60 and 360 min respectively. To compare the binding kinetics of CA and CHalo as fluorescein and SiR conjugates, a ratio of 1:3 protein:ligand (0.5 µM:1.5 µM) was mixed in reaction buffer and incubated for 1 (only **CA-SiR**), 4, 15, 60 and 240 min respectively. After incubation, the chase ligand (**MaP555**[9] or **CA-SiR**[10]), was added at 10 µM concentration for 1 h to bind remaining HT7. The samples were analysed by SDS-PAGE as described above. Controls: HT7 protein only; HT7 protein with chase ligand. The unused SDS-PAGE pockets were filled with reaction buffer.

##### Binding & fluorogenicity of caged SiR ligands

*(Figure S10fg)*

The ligand SiR conjugates (free **CHalo-SiR** and caged **Bn-CHalo-SiR** and **Cou-CHalo-SiR**, 5 µM respectively) were incubated with HT7 (2 µM) in reaction buffer (25 mM PBS (PanReac AppliChem, A0965.9010), 150 mM NaCl (Bernd Kraft, 04610.2600), 0.5 g/L BSA (Carl Roth, 3737.2) (pH = 7.5)) for 15 h to complete binding. The fluorescence intensity was determined using a Tecan Infinite M1000 plate reader (ex: 650 nm; em: 665 nm). Absorption spectra were recorded as described above.

##### Binding kinetics

*(Figure S10i)*

The ligand-SiR binding kinetics was determined from the fluorescence turn-on kinetics upon HaloTag binding at ambient temperature (23 °C). HaloTag protein (200 nM final concentration) and the ligand (**CA-SiR** or **CHalo-SiR**, 5 nM final concentration) were prepared as 2X stocks in PBS (pH = 7.5) with varying DMSO concentration in both stock solutions (1, 3 and 10% DMSO) to assess solubility / aggregation effects with different DMSO content. Both components were mixed 1:1 (25 µL ligand + 25 µL HaloTag) and the fluorescence intensity (ex: 645 nm; em: 668−672 nm) was measured every 1.2 s for 60 s (delay from mixing HaloTag and the ligand to first measurement: 5 s ±1 s) using a Cary Eclipse Fluorescence Spectrometer (scan rate: 300 nm/min, 5 nm slit width).

The emission values were averaged and plotted against the incubation time and fitted with a “one-phase association fit” which provided the “plateau value” with GraphPad Prism (version 10.4.2). The approximated rate constant k was determined by plotting ln(F*(t)) against the first 10 s of incubation time and applying a linear fit ($F^*(t) = ([\text{plateau}]-[F_{668\text{-}672nm}(t)])/([\text{plateau}]-[S_0])$; $S_0$ = fluorescence emission of ligand without HaloTag). The rate constants were calculated as k = −slope(ln(F*(t))) / c(HaloTag).

**Light dependent binding of Cou-CHalo-SiR with purified HT7**

*(Figure S10l)*

Under light exclusion, **Cou-CHalo-SiR** (5 µM) was incubated with purified HT7 (2 µM) in reaction buffer (25 mM PBS (PanReac AppliChem, A0965.9010), 150 mM NaCl (Bernd Kraft, 04610.2600), 0.5 g/L BSA (Carl Roth, 3737.2) (pH = 7.5)) for 1 h (fluorescence measurement after 20, 40 and 60 min). Then, the samples were illuminated with 405 nm light (using a pE-4000 illumination system (CoolLED) set to 5 mW/cm$^2$) for different times (5, 15, 45, 120 s) and the fluorescence intensities were measured after 5 min, 20 min, 1 h, 2 h and 4 h using a Tecan Infinite M1000 plate reader (ex: 650 nm; em: 665 nm).

**HT7 specific protein binding**

*(Figure S16ab)*

*Cell culturing and lysis*: HeLa cells were obtained from German Collection of Microorganisms and Cell Cultures (DSMZ Cat No. ACC57) and grown in Dulbecco’s modified Eagle’s medium (DMEM, Sigma-Aldrich, D1145) supplemented with 10% heat-inactivated fetal bovine serum (Biochrom S0615), 1% L-glutamine (Sigma-Aldrich, G7513), 1 mM Sodium Pyruvate (Sigma-Aldrich, S8636) and 100 nM sodium selenite (Sigma-Aldrich, 214485-5G) at 37°C and 5% $CO_2$. Washing was performed with Dulbecco’s PBS (Sigma-Aldrich, D8537), cell detachment was performed using Trypsin-EDTA solution (Sigma-Aldrich, T4174) diluted to 1x Dulbecco’s PBS (Sigma-Aldrich, D8537). Cell growth was monitored using an inverted microscope (Nikon Eclipse Ti), passage was kept between 2 and 20. Cells are tested regularly for mycoplasma contamination and only mycoplasma negative cells were used in assays. HeLa cells were lysed using M-PER reagent (thermos scientific, 78501) according to the manufacturer’s protocol and protein concentration was measured using the Pierce BCA Protein Assay Kit (Thermo Scientific, 23227), afterwards the lysate was stored on ice until used (2.8 mg protein / mL).

*In-gel fluorescence of fluorescein conjugates*: 0.5 µM HT7 protein and 5 µM of **CA-Fluo** or **CHalo-Fluo** ligand with differing amounts of HeLa cell lysate (0.1–10 µg, prepared as described above) were mixed in reaction buffer (25 mM PBS (PanReac AppliChem, A0965.9010), 150 mM NaCl (Bernd Kraft, 04610.2600), 0.5 g/L BSA (Carl Roth, 3737.2) (pH = 7.5)) and incubated for typically 1 h (or times as specified) at room temperature. The samples were analysed by SDS-PAGE as described above. Controls: lysate only; HaloTag7 only; ligands only; HT7 protein with ligand. The unused SDS-PAGE pockets were filled with reaction buffer.

*Fluorescence measurement of SiR conjugates*: HT7 (1 µM) and the ligand SiR conjugate (5 µM; **CA-SiR** or **CHalo-SiR**) were incubated with HeLa cell lysate or reaction buffer (25 mM PBS (PanReac AppliChem, A0965.9010), 150 mM NaCl (Bernd Kraft, 04610.2600), 0.5 g/L BSA (Carl Roth, 3737.2) (pH = 7.5)) and incubated at room temperature for typically 90 min (full binding was not reached for CHalo reagents). The fluorescence intensity was determined using a Tecan Infinite M1000 plate reader (ex: 650 nm; em: 665 nm).

#### 9.3 Cellular characterisation

##### 9.3.1 Cellular ligation rate

*(Figure S10j)*

HEK 293T cells were grown in DMEM (ThermoFisher 21885108) supplemented with 10% FBS (Biochrom S0615) and 1% Penicillin/Streptomycin (ThermoFisher 15140122) at 37 °C and 5% $CO_2$. Cell growth was monitored using an inverted microscope (Leica DMi1). Assays were performed in 8-well glass bottom chambered coverslips (ibidi 80827). These were coated two days before experiment by applying a 0.1 mg/mL Poly-D-Lysine (Sigma-Aldrich P7280) solution for 2 h at 37 °C and HEK cells seeded into the coated wells at a density of 20.000 cells per cm².

Transfection with Halo-mScarlet plasmid (kind gift from Kai Johnsson (Addgene plasmid # 167266 ; http://n2t.net/addgene:167266 ; RRID:Addgene_167266)) was performed 6 h after seeding. Per well (1 cm²) to be transfected, 26 µL of DMEM were combined with 0.26 µg of plasmid DNA and 0.78 µL of TransIT-LT1 transfection reagent (Mirus Bio MIR 2304), incubated at room temperature for 15–30 min, then added to cells dropwise. 27 µL of DMEM only were added to non-transfected control wells.

Confocal live cell imaging was performed at the Core Facility Bioimaging of the Biomedical Center with an inverted Leica SP8X microscope, equipped with Argon laser, WLL2 laser (470−670 nm) and acusto-optical beam splitter. Live cells were treated and recorded at 37 °C. For the duration of the assay, cells were kept in Hank’s Balanced Salt Solution (HBSS, ThermoFisher 14025092) to allow incubation without $CO_2$ atmosphere.

Experiments were performed with the following timeline: Cells were stained with 0.5 µM CMFDA (green cell tracker dye (abcam ab145459)) at 37 °C for 15 min. The medium was then changed to 250 µL HBSS and 50 µL of 6x concentrated solutions of CA-SiR and CHalo-SiR were added to achieve 10 nM final concentration. Images were acquired after 5, 20, 40, 60, 90, 120 and 180 min of incubation.

The microscope was programmed to take three images of different fields of view per condition and time point tested, which were focused using reflection-based adaptive focus control. Bias deriving from time delays between probes imaged in different wells was minimized by permutation of the positioning of probes between replicates. Images were acquired with a 20× 0.75 objective and additional 2× optical zoom. Image pixel size was 569 nm. The following fluorescence settings were used: CMFDA excitation 488 nm (Argon), emission 500–540 nm; mScarlet excitation 569 nm (WLL), emission 580–600 nm; SiR excitation 652 nm (WLL), emission 665–705 nm. Recording was performed sequentially to avoid bleed-through. CMFDA, mScarlet and SiR were recorded with hybrid photo detectors (HyDs), a transmitted light image was generated with a conventional photomultiplier tube. (Note: the represented brightness of the CMFDA CellTracker images was increased for the time-points 24 and 48 h due to signal loss after long incubations.)

Images were analysed using Fiji ImageJ. Images of transfected cells were segmented by thresholding on the mScarlet channel. Images of non-transfected cells were segmented by thresholding on the CMFDA channel. The resulting masks were then applied to the SiR channel to obtain a mean fluorescence intensity value. Analysed images were in focus and field of view outliers were excluded from analysis (for each biological replicate at least two field of view images were evaluated, otherwise the replicate was repeated). Data was plotted using GraphPad Prism.

##### 9.3.2 Spatiotemporally controlled cellular microtubule labelling: Cou-CHalo-SiR

*(Figures: labelling of stable microtubules: 2bc, Movie S1; localisation to endomembranes: S17)*

**Cell culture**

U2OS cells (ATCC, CVCL_0224) were cultured in Dulbecco's Modified Eagle Medium (DMEM, Sigma) supplemented with 10% Fetal Bovine Serum (FBS, Corning), 100 U/mL penicillin and 100 µg/mL streptomycin (1% Penstrep, Sigma) at 37 °C with 5% $CO_2$.

**DNA constructs**

StableMARK-Halo-2xFKBP was cloned into a pSIN-TRE vector described in Noordstra *et al.*[72] with a puromycin resistance cassette by Gibson assembly. Prior to this a StableMARK-Halo-2xFKBP construct was cloned into a pB80 vector from Nijenhuis *et al.*[73] (gift from L. Kapitein, UU, Utrecht, Netherlands; Addgene, #174625) with flexible GS linkers between all the domains by Gibson assembly, using the StableMARK from StableMARK-2xmNeon (Jansen *et al.*[74], gift from L. Kapitein, UU, Utrecht, Netherlands; Addgene, #174649) and FKBP from Vim-mCh-FKBP (Pasolli *et al.*[75], Addgene; #240423) and Halo from Halo-Rab6A (Meiring *et al.*[76], Addgene; #190171). Lentiviral packaging constructs pMD2.G (Addgene, #12259) and psPAX2 (Addgene, #12260) were gifts from D. Trono, EPFL, Lausanne, Switzerland. EB3-GFP (Addgene, #190164) was described in Stepanova *et al.*[77]. RTN4-FKBP-GFP (Farias *et al.*[78]) was a gift from G.G. Farias, UU, Utrecht, Netherlands) and AKAP1(1-30)-mCh-iLID was a gift from L. Kapitein, UU, Utrecht, Netherlands.

**Lentivirus production**

HEK293T cells at 90% confluency were transfected with 15 µg pSIN-TRE StableMARK-Halo-2xFKBP, 5 µg pMD2.G and 10 µg psPAX2 using 90 µL MaxPEI (1 mg/mL Polyethylenimine, Polysciences) in optiMEM (1.2 mL final volume). Transfection mix was vortexed at low speed and pre-incubated at room temperature for 10 min before adding to HEK293T cells. Medium was refreshed the following 3 days, only collected for virus harvest on the last 2 days. Medium was filtered through a 0.045 µm filter before applying to a Amicon Ultra-15 filter column (Merck, UFC903029) and centrifuging for 25 min at 1400×g to purify virus. Supernatant was aliquoted and stored at −80 °C until used for cell transduction.

**Cell line transduction and selection**

U2OS cells were transduced with virus when they were at 40% confluency, medium was refreshed the following day and 3 µg/mL puromycin was added 48 h after transduction for 20 h to select for pSIN-TRE StableMARK-Halo-2xFKBP containing cells. From here single cells were plated and expanded to make a clonal cell line, subsequently selected for optimal expression levels upon 16 h treatment with 500 ng/mL doxycycline (Doxycycline-hyclate, Abcam, ab141091).

**Cell transfection**

U2OS cells were transfected with EB3-GFP or RTN4-FKBP-GFP and AKAP1(1-30)-mCh-iLID using Fugene 6 (Promega, #E2691) according to manufacturer's instructions at a ratio of 1 µg DNA to 3 µL Fugene 6.

**Cou-CHalo-SiR uncaging to label stable microtubules in U2OS**

pSIN StableMARK-Halo-2xFKBP U2OS were seeded into an 8 well labtek chamber (Thermo Scientific, #155409) at 40% together with EB3-GFP Fugene 6 transfection mix and 500 ng/mL doxycycline the day before the experiment. At the day of the experiment, the medium on the cells was replaced with 200 µL fresh medium. In a dark room without blue/UV light a 2x **Cou-Chalo-SiR** solution (10 µM **Cou-Chalo-SiR**, 2% DMSO in cell media) was prepared and prewarmed to 37 °C. Samples were imaged on a custom spinning disc confocal microscope with a Nikon Eclipse Ti2-E body with Perfect Focus, a Yokogawa CSU-X1-A1 spinning disc unit, an ASI MS-2000-XYZ stage with Piezo Top plate and STXG-PLAMX-SETZ21L TokaiHit incubation chamber keeping sample at 37 °C with 5% $CO_2$ during the experiment. The sample was imaged using a Plan Apo VC 100x / 1.40 oil objective and Prime BSI sCMOS camera (Teledyne Photometrics). Voltran Stradus lasers 405 (wavelength for uncaging), 488 (GFP: microtubule marker) and 642 nm (SiR) were used to illuminate sample in combination with ET460/50m, ET525/50m and ET700/75m Chroma filters respectively. MetaMorph 7.10 software was used to control the microscope. 200 µL of the 2x **Cou-Chalo-SiR** solution was added to a well such that the final concentration was 5 µM **Cou-Chalo-SiR** and 1% DMSO, 5 min prior to commencing imaging. EB3-GFP (488 nm, 50 µW, 0.1 W/cm$^2$) and **Cou-Chalo-SiR** (642 nm, 0.52 mW, 1 W/cm$^2$) were imaged every 10 s using 500 ms exposure times. A single 2 s long pulse of 405 nm (0.74 mW, 2 W/cm$^2$) was applied to the sample to uncage **Cou-Chalo-SiR** at the indicated time points, spaced 2 min apart. *(Figures: 2bc; Movie S1)*

**Localization of CHalo-SiR to ER and mitochondria**

WT U2OS were seeded at 15% onto 25 mm glass #1.5 coverslips in a 6 well cell culture plate. The following day cells were transfected with RTN4-FKBP-GFP and AKAP1(1-30)-mCh-iLID. One day after transfection, the coverslips with cells were mounted in Attofluor cell chambers (Invitrogen, #A7816) with fresh medium. Samples were imaged on a custom spinning disc confocal microscope with a Nikon Eclipse Ti2-E body with perfect focus, a Yokogawa CSU-W1-T1 spinning disc unit, ASI MS-2000-XYZ stage with Piezo Top plate and STXG-PLAMX-SETZ21L TokaiHit incubation chamber keeping sample at 37 °C with 5% $CO_2$ during the experiment. Sample was imaged using a Plan Apo λD 100x / 1.45 objective and Prime BSI sCMOS camera (Teledyne Photometrics). Voltran Stradus lasers 488 (GFP: ER marker) and 642 nm (SiR) and Coherent OBIS 561 nm (mCherry: mitochondrial marker) lasers were used to illuminate sample in combination with ET525/50m, ET700/75m and ET630/75m Chroma filters respectively. MetaMorph 7.10 software was used to control the microscope. RTN4-FKBP-GFP, AKAP1(1-30)-mCh-iLID and **CHalo-SiR** were imaged once every 1 min for 20 min after addition. **CHalo-SiR** (5 µM, 1% DMSO in cell medium) was applied to cells immediately after the acquisition started.

##### 9.3.3 Live cell imaging of enzyme activity: Leu-CHalo-SiR

*(Figures: 2ef, S18ab, S19−25)*

Bestatin is both a broad-acting ***aminopeptidase inhibitor*** (reason why we employ it to transiently chemically inhibit the function of peptidases: this is a way to test the cellular enzyme-specificity and activity-responsiveness of our leucine-anilide-caged model probe; the unnatural substrate design of leucine anilides makes them primarily processed in human cells only by the more tolerant enzyme leucine aminopeptidase), and an inhibitor of cellular inhibitor of apoptosis protein 1 (cIAP1, an E3 ligase i.e. having roles in the proteosomal degradation of other proteins). In the conditions we use and with the readout that we are monitoring, we do not expect that its cIAP1 inhibiting effects are relevant to assay outcomes or conclusions.

HEK 293T cells were grown in DMEM (ThermoFisher 21885108) supplemented with 10% FBS (Biochrom S0615) and 1% Penicillin/Streptomycin (ThermoFisher 15140122) at 37 °C and 5% $CO_2$. Cell growth was monitored using an inverted microscope (Leica DMi1). Assays were performed in 8-well glass bottom chambered coverslips (ibidi 80827). These were coated two days before experiment by applying a 0.1 mg/mL Poly-D-Lysine (Sigma-Aldrich P7280) solution for 2 h at 37 °C and HEK cells seeded into the coated wells at a density of 20.000 cells per cm².

Transfection with Halo-mScarlet plasmid (kind gift from Kai Johnsson (Addgene plasmid # 167266 ; http://n2t.net/addgene:167266 ; RRID:Addgene_167266)) was performed 6 h after seeding. Per well (1 cm²) to be transfected, 26 µL of DMEM were combined with 0.26 µg of plasmid DNA and 0.78 µL of TransIT-LT1 transfection reagent (Mirus Bio MIR 2304), incubated at room temperature for 15–30 min, then added to cells dropwise. 27 µL of DMEM only were added to non-transfected control wells.

Confocal live cell imaging was performed at the Core Facility Bioimaging of the Biomedical Center with an inverted Leica SP8X microscope, equipped with Argon laser, WLL2 laser (470−670 nm) and acusto-optical beam splitter. Live cells were treated and recorded at 37 °C. For the duration of the assay, cells were kept in Hank's Balanced Salt Solution (HBSS, ThermoFisher 14025092) to allow incubation without $CO_2$ atmosphere.

Experiments were performed with the following timeline: Cells were stained with 0.5 µM CMFDA (green cell tracker dye (abcam ab145459)) at 37 °C for 15 min. The medium was then changed to 250 µL HBSS with or without 100 µM bestatin (Carl Roth 2937.1). After 25 min of bestatin incubation, pre-treatment images were recorded. After 30 min of bestatin incubation, 50 µL of 6x concentrated solutions of the different silicon-rhodamine (SiR) HaloTag ligand dyes were added on top of the wells to achieve the final concentrations of 10 nM (for **CA-SiR**, **CHalo-SiR**) or 1 µM (for **Leu-CHalo-SiR**, **Bn-CHalo-SiR** and all ligands on non-transfected cells) respectively. Images were acquired after 5, 20, 40, 60, 90, 120 and 180 min of incubation. All wells were then washed twice with HBSS and post-wash images acquired at 0, 30, 60 and 120 min thereafter. Cells were then returned to the incubator (37°C, 5% $CO_2$) in DMEM (+FCS, +Penicillin/Streptomycin) supplemented with 100 µg/mL primocin (invivogen ant-pm) and again transferred to the microscope for image acquisition at 24 h (± 1.5 h) and 48 h (± 1.5 h) post washing.

The microscope was programmed to take three images of different fields of view per condition and time point tested, which were focused using reflection-based adaptive focus control. Bias deriving from time delays between probes imaged in different wells was minimized by permutation of the positioning of probes between replicates. Images were acquired with a 20× 0.75 objective and additional 2× optical zoom. Image pixel size was 569 nm. The following fluorescence settings were used: CMFDA excitation 488 nm (Argon), emission 500–540 nm; mScarlet excitation 569 nm (WLL), emission 580–600 nm; SiR excitation 652 nm (WLL), emission 665–705 nm. Recording was performed sequentially to avoid bleed-through. CMFDA, mScarlet and SiR were recorded with hybrid photo detectors (HyDs), a transmitted light image was generated with a conventional photomultiplier tube. (Note: the represented brightness of the CMFDA CellTracker images was increased for the time-points 24 and 48 h due to signal loss after long incubations.)

Images were analysed using Fiji ImageJ. Images of transfected cells were segmented by thresholding on the mScarlet channel. Images of non-transfected cells were segmented by thresholding on the CMFDA channel. The resulting masks were then applied to the SiR channel to obtain a mean fluorescence intensity value. Analysed images were in focus and field of view outliers were excluded from analysis (for each biological replicate at least two field of view images were evaluated, otherwise the replicate was repeated). Data was plotted using GraphPad Prism.

##### 9.3.4 Pulse-chase quantification of integrated enzyme activity: SS66T/C-CHalo-SiR

U2OS cells with stable GFP-NUP98 marker (nuclear pore complex) and transient HaloTag7-Sec61 fusion protein (ER-localised protein transport complex) as recorder were cultured and passaged in DMEM with penicillin (100 U/mL) and streptomycin (100 µg/mL), 10% FCS, 100 nM $Na_2SeO_3$ (for optimal functional translation of selenoproteins, relevant for TrxR-dependent redox biology), and 300 µg/mL G418 (selection for HaloTag-Sec61). 10 k cells were seeded per well in Greiner µ-clear 96 well plate, in 100 µL medium, one day before treatment; cells were then treated with caged-CHalo-SiR pulse ligands (10 µL of 10X probe stocks that had been freshly prepared from addition of 9 µL of DMEM to 1 µL of pulse ligand solution in 100% DMSO) establishing 1% DMSO final concentration. After 16 h incubation, JF525-HTL was added (chase ligand; 1 µM final concentration, by directly adding 1 µL of 100X stock in pure DMSO). GFP channel imaging showed image intensities stabilised after 15 min (saturation of remaining HaloTag binding sites by JF525). Imaging was then conducted with a BioTek Cytation 5 Cell Imaging Multimode Reader (epifluorescence microscope), using CY5 filter cube for SiR imaging ("623" nm LED with 628/40 bandpass excitation filter and 685/40 bandpass emission filter, red channel in **Fig 2h**) and YFP filter cube for GFP and JF525 imaging ("505" nm LED with 500/24 bandpass excitation filter and 542/27 bandpass emission filter, green channel in **Fig 2h**). Images were analysed using Fiji. ROIs containing the ER, excluding the nucleus and the cell periphery, were used to quantify the SiR/JF525 pulse/chase ratios; (minor) contributions from extracellular background to the per-pixel fluorescence were first subtracted from the average ROI fluorescence values in each channel, then the SiR/JF525 intensity ratios were calculated.

##### 9.3.5 Spatiotemporally controlled protein heterodimerisation: Cou-CHalo-BG

*(Figures: 2l, S26b; Movie S2)*

**Primary Neuronal Cultures**

C57BL/6 mice were obtained from the animal facility at the Max Planck Institute for Biological Intelligence in Martinsried, Germany. All procedures involving mice were performed in compliance with the Government of Upper Bavaria, approved by the license number ROB-55.2-2532.Vet_02-23-213. Primary neurons were collected as previously described.[79–81] E16.5 timed-pregnant dams were euthanized by $CO_2$ and the mouse embryos were rapidly collected and decapitated in ice-cold dissection media (Hank´s Balanced Salt Solution supplemented with 100 mM of $MgCl_2$, 100 mM of HEPES and 10 mM kynurenic acid (Sigma-Aldrich, K3375)). The brains were dissected and stripped of meninges. Hippocampi were collected in ice-cold dissection media and dissociated into a single-cell suspension using of papain (0.2 mg/mL; Roche, 10108014001) for 5 min at 37 °C. Enzyme digestion was quenched using ovomucoid trypsin inhibitor (1 mg/mL; Abnova, P5243), followed by washes and trituration in complete media: Neurobasal (Gibco, 21103049) with B27 (Gibco, 17504044), penicillin/streptomycin and *L*-glutamine (Gibco, 10378016). Cells were then counted and $10^5$ cells were plated onto glass-bottom 24-well plates previously coated with poly-*L*-lysine and laminin. Neurons were maintained in a humidified incubator at 37 °C and 5% $CO_2$ with a half-media change on DIV 5 and 7 (DIV = days *in vitro*).

**Treatments, Microscopy and analysis**

On DIV 5-7, neurons were co-transfected with 50 ng of GFP-Halo (subcloned from Addgene #67764 by removing the PEX3 sequence using MfeI and AscI digestion followed by T4 re-ligation (all enzymes from New

England Biolabs) and transformation), 400 ng of SNAP-OMM (Addgene, #69599) and 50 ng of mito-mRaspberry (Addgene, #55931) per well, using Lipofectamine 2000 (Invitrogen, 11668019). The transfection ratio of GFP-Halo and SNAP-OMM was found to be optimal between 1:4 and 1:8. Cells were treated and imaged at least 48 h post-transfection (DIV 7-9).

Neurons were imaged on an inverted laser-scanning confocal microscope Stellaris DMI8 (Leica), controlled by the LAS-X software (Leica) using a HC PL APO CS2 63×/1.40 OIL objective, kept at 37 °C by a Okolab black cage incubator. Imaging settings: excitation with a white light laser (GFP: 489 nm and mRaspberry: 598 nm) and emission collection in a single sequence using 2 hybrid detectors for the GFP (500–530 nm) and mRaspberry (609–653 nm) channels.

The neurons were treated with **Cou-CHalo-BG** (final concentration: 5 µM, 0.5% DMSO) or DMSO only (negative control) for 1 h in complete media (Neurobasal with B27, penicillin/streptomycin and *L*-glutamine). The excess of the **Cou-CHalo-BG** was removed by medium change and incubation for 5 min (the wash cycle was repeated 3×), then the medium was changed to Hibernate E without phenol red for imaging (Transnetyx Tissue, HEPR500). The acquisitions for the timelapse experiments were performed under the Lightning mode at 598 Hz on 50 × 50 µm (1176 × 1176 px) frames at 30 s intervals (1 frame before illumination, then "uncaging frame", then imaging for 5 min). Uncaging was performed using a 405 nm laser diode (16.1−213 µW, 45.6−603 kW/cm$^2$) during the acquisition of the "uncaging frame" (pixel dwell time of 750 ns).

The re-localisation of the GFP signal to the mitochondrial mRaspberry signal was quantified by comparing the images before uncaging and 5 min after uncaging. The images were analyzed with JaCoP in FIJI/ImageJ (version 1.54) and the Mander´s coefficient for the ratio of GFP overlapping mitochondria was calculated.

### 10 Synthetic Chemistry

#### 10.1 Chemistry methods and techniques

##### 10.1.1 Analytical methods

High resolution mass spectrometry (**HRMS**) was conducted on the following instruments: (1) a *Thermo Finnigan LTQ FT Ultra FourierTransform* ion cyclotron resonance spectrometer from *ThermoFisher Scientific GmbH* applying electron spray ionisation (ESI) with a spray capillary voltage of 4 kV at temperature 250 °C with a method dependent range from 50 to 2000 u;(2) a *Finnigan MAT 95 from Thermo Fisher Scientific* applying electron ionisation (EI) at a source temperature of 250 °C and an electron energy of 70 eV with a method dependent range from 40 to 1040 u; and (3) a Waters Xevo G2-XS Q-TOF applying electron spray ionisation (ESI) with a spray capillary voltage of 2 kV at a source temperature of 140 °C with a method dependent range from 50 to 1200 u.

Nuclear magnetic resonance (**NMR**) spectroscopy was performed using the following instruments: (1) a *Bruker Avance* (600/150 MHz, with TCI cryoprobe) or (2) a *Bruker Avance III HD Biospin* (400/100 MHz, with BBFO cryoprobe™) from Bruker Corp. or (3) a Bruker Avance III HD (800 MHz, with cryoprobe) or (4) a Bruker Avance Neo (600/150 MHz, with cryoprobe). NMR-spectra were measured at 298 K, unless stated otherwise, and were analysed with the program *MestreNova 12* developed by *MestreLab Ltd.* $^1$H-NMR spectra chemical shifts (δ) in parts per million (ppm) relative to tetramethylsilane (δ = 0 ppm) are reported using the residual protic solvent ($CHCl_3$ in $CDCl_3$ : δ = 7.26 ppm, DMSO-$d_5$ in DMSO-$d_6$: δ = 2.50 ppm, $CHD_2OD$ in $CD_3OD$: δ = 3.31 ppm) as an internal reference. For $^{13}$C-NMR spectra, chemical shifts in ppm relative to tetramethylsilane (δ = 0 ppm) are reported using the central resonance of the solvent signal ($CDCl_3$: δ = 77.16 ppm, DMSO-$d_6$: δ = 39.52 ppm, $CD_3OD$: δ = 49.00 ppm) as an internal reference. For $^1$H-NMR spectra in addition to the chemical shift the following data is reported in parenthesis: multiplicity, coupling constant(s) and number of hydrogen atoms. The abbreviations for multiplicities and related descriptors are s = singlet, d = doublet, t = triplet, q = quartet, or combinations thereof, m = multiplet and br = broad. When rotamers were observed in the NMR spectra, the corresponding signals are separated by a slash ("/"). Where known products matched literature analysis data, only selected data acquired are reported.

Analytical high performance liquid chromatography (**HPLC**) analysis was conducted either using an *Agilent 1100* system from *Agilent Technologies Corp.,* Santa Clara (USA) equipped with a DAD detector and a *Hypersil Gold* HPLC column from *ThermoFisher Scientific GmbH*, Dreieich (Germany) or a *Agilent 1200 SL* system *Agilent Technologies Corp.,* Santa Clara (USA) equipped with a DAD detector, a *Hypersil Gold* HPLC column from *ThermoFisher Scientific GmbH*, Dreieich (Germany) and consecutive low-resolution mass detection using a LC/MSD IQ mass spectrometer applying ESI from *Agilent Technologies Corp.,* Santa Clara (USA). For both systems mixtures of water and MeCN (both analytical grade, 0.1 % formic acid) were used as eluent systems.

##### 10.1.2 Synthetic techniques

Unless stated otherwise, all reactions were performed without precautions regarding potential air- and moisture-sensitivity and were stirred with Teflon-coated magnetic stir bars. For work under inert gas (nitrogen) atmosphere, a Schlenk apparatus and a high vacuum pump from Vacuubrand GmbH, Wertheim (Germany) were used. For solvent evaporation a *Laborota 400* from Heidolph GmbH, Schwabach (Germany) equipped with a vacuum pump was used. Flash column chromatography was conducted with a Biotage® Isolera One Chromatograph with Biotage® Sfär Silica D columns (10 g or 25 g silica) for normal-phase (np) chromatography or with Biotage® Sfär C18 D columns (12 g or 30 g silica) for reversed-phase (rp) chromatography. Reactions were monitored by thin layer chromatography (TLC) on TLC plates (*Si 60 F254 on aluminium sheets*) provided by Merck GmbH and visualised by UV irradiation and by analytical HPLC-MS. The procedures and yields are not optimised.

##### 10.1.3 Chemicals

All chemicals, which were obtained from BLDpharm, Sigma-Aldrich, TCI, Alfa Aesar, Acros, abcr or carbolution were used as received and without purification. Tetrahydrofuran (THF), dichloromethane (DCM) and dimethylformamide (DMF) were provided by Acros and were stored under argon atmosphere and dried over molecular sieves. TLC control, extractions and column chromatography were conducted using distilled, technical grade solvents. Whenever the term *hexanes (Hex)* is used, the applied solvent actually comprised isomeric mixtures of hexane (2-methylpentane, 3-methylpentane, 2,2-dimethylbutane, 2,3-dimethylbutane).

#### 10.2 Synthetic procedures

##### 10.2.1 General Procedures

**General Procedure A: deprotection of Boc-amines**

The Boc-amines were dissolved in anhydrous DCM (0.02 M–0.25 M) and hydrogen chloride in dioxane (4 M) was added to give 2 M HCl concentration in the mixture. The reaction mixture was stirred at room temperature for 1 h, the volatiles were removed *in vacuo* and the product was used without further purification for the next synthetic step.

**General Procedure B: amide coupling with HATU**

The carboxylic acid (1.0 eq) was dissolved in anhydrous DMF (0.02 M–0.25 M), DIPEA (5.0 eq) and *O*-(7-Azabenzotriazol-1-yl)-*N*,*N*,*N*′,*N*′-tetramethyluronium-hexafluorphosphat (HATU, 1.0 eq–1.2 eq) were added and the mixture was stirred at room temperature for 20 min. Then the amine (0.5 eq–2.0 eq, equivalents varied depending on the synthetic accessibility of respective carboxylate and amine) was added and the reaction mixture was stirred at room temperature for 1–12 h until completion (monitored by HPLC-MS). The solvent was removed *in vacuo* and the crude product was diluted with ethyl acetate and a half-saturated solution of sodium bicarbonate. The layers were separated, the aqueous layer was extracted with ethyl acetate (3×) and the combined organic layers were washed with brine and dried over sodium sulfate. The crude product was purified by flash column chromatography.

**General Procedure C: DEACM-carbamate formation with triphosgene**

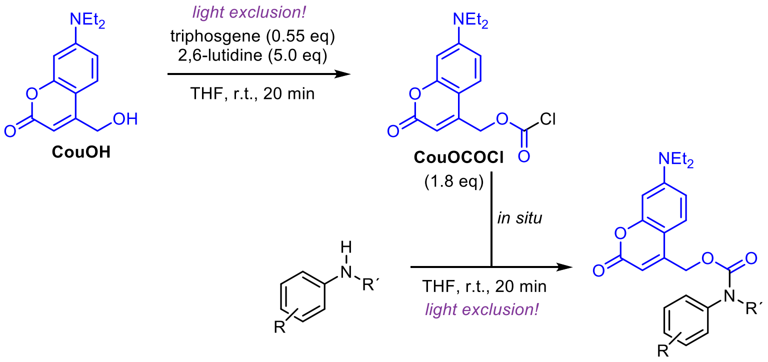

*Note: the reaction must be performed under light exclusion.*

Adaptation from previously described procedure.[82] Under nitrogen atmosphere, **CouOH** (1.0 eq) was dissolved in anhydrous THF (0.1 M). Triphosgene (0.55 eq) and 2,6-lutidine (5.0 eq) were added and the reaction mixture was stirred at room temperature for 20 min (HPLC-MS analysis of the reaction progress by quenching with methyl piperazine in acetonitrile).

In a separate flask, the aniline was dissolved in anhydrous THF (0.02 M–0.2 M) under nitrogen atmosphere. Then the reaction mixture with the **CouOCOCl** (1.8 eq) was added dropwise and the mixture was stirred at room temperature for 20 min. Upon full conversion of the aniline, the excess of the **CouOCOCl** was reacted with a secondary amine (piperidine or methyl piperazine depending on the desired polarity for the following purification). The mixture was diluted with ethyl acetate and water, extracted (3×), the organic layers were washed with brine (1x) and the crude product was purified by flash column chromatography.

*Note 1: the chloroformate formation shows unwanted chloride formation (giving CouCl) with tertiary amine bases like DIPEA. A previously described procedure[82] solved the problem by reacting CouOH and phosgene without a base in THF, which requires at least 6 h reaction time for decent conversion. We found that using the sterically hindered pyridine base 2,6-lutidine enables selective formation to CouOCOCl within minutes.*

*Note 2: the carbamate formation with CouOCOCl is much cleaner than the reaction with CouO-PFPC (see General Procedure D) which mostly gives the hydrolysed byproduct CouOH and minor amounts of CouCl. We still used an excess of the chloroformate to ensure full conversion of the aniline since we often observed partial hydrolysis of the chloroformate during the reaction.*

**General Procedure D: DEACM-carbamate formation with bis(pentafluorophenyl)carbonate**

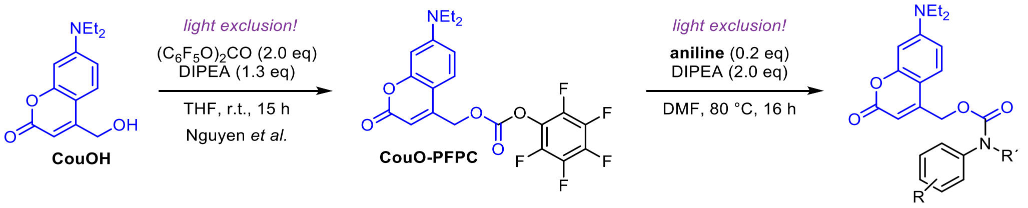

*Note: the reaction must be performed under light exclusion.*

**CouO-PFPC** was prepared according to a previously described procedure (Nguyen *et al.*[82], compound 14) using bis(pentafluorophenyl)-carbonate to afford the coumarin pentafluorophenylcarbonate (PFPC) which can be purified and stored at –20 °C.[82] For photocaging (secondary) anilines, the aniline (0.2 eq) was dissolved in anhydrous DMF (0.05 M–0.2 M), DIPEA (2.0 eq) and **CouO-PFPC** (1.0 eq) were added and the reaction mixture was heated to 80 °C for 16 h. The solvent was removed in vacuo and the crude product was purified by flash column chromatography.

*Note: while the CouO-PFPC is a convenient reagent due to its stability, the carbamate formation reaction gives unwanted side products (ether CouO-$C_6F_5$, carbonate $(CouO)_2CO$ and the hydrolysis product CouOH, therefore CouO-PFPC is used in excess). Separation of these byproducts from the desired product is often inconvenient without prep-HPLC purification, thus, for larger scale reactions the use of triphosgene is more advantageous.*

**General Procedure E: Siliconrhodamine (SiR) conjugation**

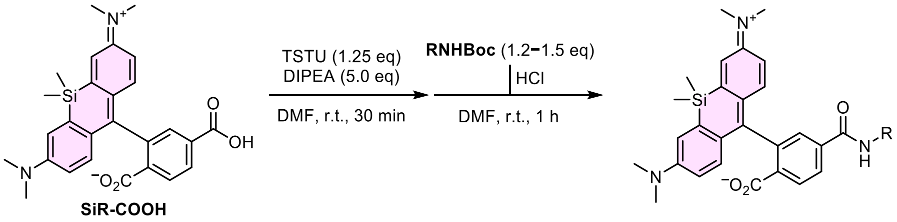

6-Carboxy tetramethyl siliconrhodamine (**SiR-COOH**) was dissolved in anhydrous DMF (4.2 mM), DIPEA (5.0 eq) and TSTU (50 mM in DMF, 1.6 eq) were added and the reaction mixture was stirred at room temperature for 30 min. In parallel the Boc-amine was deprotected following General Procedure A, subsequently added to the SiR-NHS ester and stirred at room temperature for 1 h. The mixture was purified by preparative HPLC.

#### 10.2.2 CHalo Ligand building blocks

##### Synthetic route for CHalo-Boc

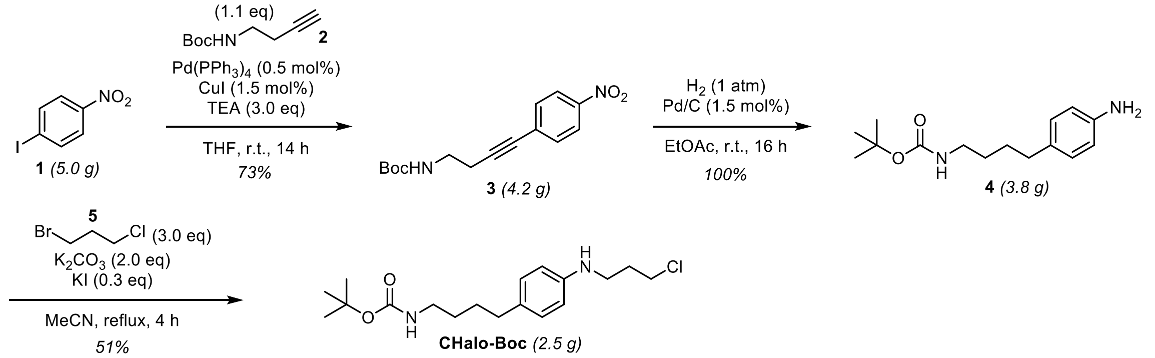

##### Compound 3

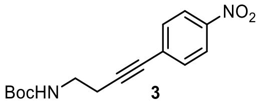

1-Iodo-4-nitrobenzene (**1**, 5.00 g, 19.9 mmol, 1.0 eq) was dissolved in anhydrous THF (50 mL, 0.4 M), triethylamine (8.3 mL, 60 mmol, 3.0 eq) and *tert*-butyl but-3-yn-1-ylcarbamate (**2**, 3.83 g, 21.5 mmol, 1.1 eq) were added and the solution was degassed by bubbling nitrogen through the mixture for 10 min. Copper iodide (57 mg, 0.30 mmol, 1.5 mol%) and tetrakis(triphenylphosphine)palladium(0) (0.12 g, 99 µmol, 0.50 mol%) were added and the reaction mixture was stirred at room temperature for 14 h. The volatiles were removed *in vacuo*, the residue was diluted with water (50 mL), brine (10 mL) and diethyl ether (50 mL), the layers were separated and the aqueous layer was extracted with diethyl ether (3 × 30 mL). The combined organic layers were dried over sodium sulfate and the solvent was removed *in vacuo*. The crude product was purified by flash column chromatography (*iso*-hexanes/ethyl acetate; 5→25% EtOAc) affording **3** (4.23 g, 14.6 mmol, 73%) as an orange solid.

**TLC** Rf = 0.10 (*np*, Hex:EtOAc 9:1); Rf of **1** = 0.62
**$^1$H-NMR** (600 MHz, $CDCl_3$): *δ* (ppm) = 8.16 (d, J = 8.8 Hz, 2H), 7.53 (d, J = 8.9 Hz, 2H), 4.85 (s, 1H), 3.38 (s, 2H), 2.65 (t, J = 6.6 Hz, 2H), 1.45 (s, 9H).
**$^{13}$C-NMR** (151 MHz, $CDCl_3$): *δ* (ppm) = 155.8, 147.0, 132.5, 130.6, 123.7, 93.4, 80.6, 79.8, 39.4, 28.5, 21.4.
**HRMS** (ESI+): m/z calc. for $C_{10}H_{11}N_2O_2^+$ $[M{-}Boc{+}H]^+$: 191.0815, found: 191.0822.

##### Compound 4

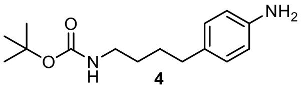

**3** (4.18 g, 14.4 mmol, 1.0 eq) was dissolved in ethyl acetate (30 mL, 0.5 M) under nitrogen atmosphere, palladium on charcoal (Pd/C, 0.26 g, 0.22 mmol, 1.5 mol%) was added and the reaction was stirred under hydrogen atmosphere (1 atm) for 17 h. The Pd/C was removed by filtration over Celite (washed with ethyl acetate (7×) and the filtrate was concentrated under reduce pressure affording **4** (3.81 g, 14.4 mmol, 100%) as a colourless solid.

**TLC** Rf = 0.15 (*np*, Hex:EtOAc 3:1); Rf of **3** = 0.47
**$^1$H-NMR** (600 MHz, $CDCl_3$): *δ* (ppm) = 6.96 (d, J = 8.3 Hz, 2H), 6.63 (d, J = 8.3 Hz, 2H), 4.50 (s, 1H), 3.45 (s, br, 2H), 3.12 (d, J = 6.3 Hz, 2H), 2.51 (t, J = 7.6 Hz, 2H), 1.62 – 1.54 (m, 2H), 1.48 (dt, J = 13.5, 6.6 Hz, 2H), 1.44 (s, 9H).
**$^{13}$C-NMR** (151 MHz, $CDCl_3$): *δ* (ppm) = 156.1, 144.0, 132.7, 129.3, 115.5, 79.1, 40.6, 34.7, 29.7, 29.0, 28.6.
**HRMS** (ESI+): m/z calc. for $C_{15}H_{24}N_2NaO_2^+$ $[M{+}Na]^+$: 287.1730, found: 287.1733.

##### CHalo-Boc

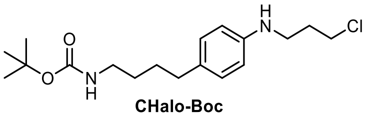

**CHalo-Boc**

**4** (3.74 g, 14.1 mmol, 1.0 eq) was dissolved in anhydrous acetonitrile (40 mL, 0.35 M) under nitrogen atmosphere. Potassium carbonate (3.91 g, 28.3 mmol, 2.0 eq), potassium iodide (0.71 g, 4.2 mmol, 0.30 eq) and 1-bromo-3-chloropropane (**5**, 4.2 mL, 42 mmol, 3.0 eq) were added and the reaction mixture was heated to reflux for 4 h. The volatiles were removed under reduced pressure and the residue was diluted with water (50 mL), brine (30 mL) and ethyl acetate (30 mL). The layers were separated and the aqueous layer was extracted with ethyl acetate (2 × 30 mL). The combined organic layers were washed with brine (40 mL), dried over sodium sulfate and the solvent was removed *in vacuo*. The crude product was purified by flash column chromatography (*iso*-hexanes/ethyl acetate; 0→25% EtOAc) affording **CHalo-Boc** (2.45 g, 7.19 mmol, 51%) as an off-white solid. The starting material **4** was partially recovered (0.92 g, 3.5 mmol, 25%).

*Note: Although the reaction did not proceed to full conversion, longer reaction times and more equivalents of 1-bromo-3-chloropropane were avoided due to increasing double alkylation.*

**TLC** *Rf* = 0.43 (*np*, Hex:EtOAc 3:1); Rf of **4** = 0.15
**$^1$H-NMR** (600 MHz, $CDCl_3$): *δ* (ppm) = 6.99 (d, J = 8.4 Hz, 2H), 6.56 (d, J = 8.4 Hz, 2H), 4.52 (s, 1H), 3.65 (t, J = 6.3 Hz, 2H), 3.31 (t, J = 6.6 Hz, 2H), 3.12 (q, J = 6.7 Hz, 2H), 2.51 (t, J = 7.4 Hz, 2H), 2.06 (p, J = 6.5 Hz, 2H), 1.63 – 1.53 (m, 2H), 1.53 – 1.46 (m, 2H), 1.44 (s, 9H).
**$^{13}$C-NMR** (151 MHz, $CDCl_3$): *δ* (ppm) = 156.1, 146.0, 131.4, 129.3, 113.0, 79.1, 42.8, 41.3, 40.6, 34.6, 32.1, 29.7, 29.0, 28.5.
**HRMS** (ESI+): m/z calc. for $C_{18}H_{29}ClN_2NaO_2^+$ $[M+Na]^+$: 363.1810, found: 363.1812.

##### Cou-CHalo-Boc

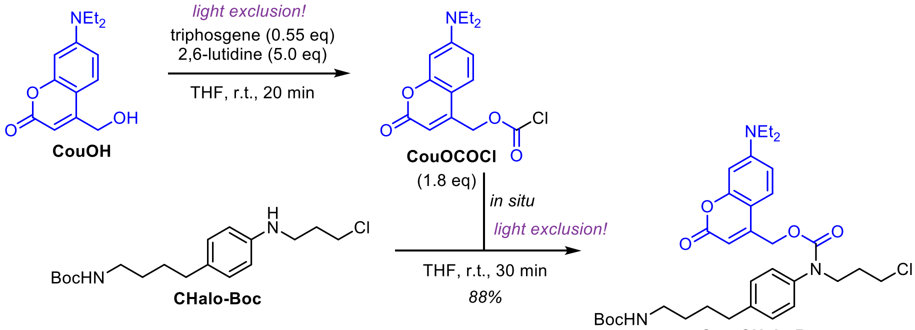

*<u>Note</u>: the reaction must be performed under light exclusion.*

According to General Procedure C: **CouOH** (71 mg, 0.29 mmol, 1.0 eq) was dissolved in anhydrous THF. Triphosgene (47 mg, 0.16 mmol, 0.55 eq) was added, then 2,6-lutidine (0.17 mL, 1.4 mmol, 5.0 eq) was added dropwise and the reaction mixture was stirred at room temperature for 20 min to give **CouOCOCl**.

In a separate flask, **CHalo-Boc** (54 mg, 0.16 mmol, 1.0 eq) was dissolved in anhydrous THF and the reaction mixture of the **CouOCOCl** (1.8 eq) was added dropwise. The mixture was stirred at room temperature for 30 min. *N*-Methylpiperazine (53 µL, 0.48 mmol, 3.0 eq) was added to react with the excess **CouOCOCl** and the mixture was stirred for 10 min. The volatiles were removed *in vacuo* and the residue was diluted with water (15 mL), brine (10 mL) and ethyl acetate (15 mL). The layers were separated and the aqueous layer was extracted with ethyl acetate (2 × 15 mL). The combined organic layers were washed with brine (20 mL), dried over sodium sulfate and the solvent was removed *in vacuo*. The crude product was purified by flash column chromatography (*iso*-hexanes/ethyl acetate; 5→25% EtOAc) affording **Cou-CHalo-Boc** (86 mg, 0.14 mmol, 88%) as a yellow solid.

**TLC** *Rf* = 0.27 (*np*, Hex:EtOAc 2:1); Rf of **CHalo-Boc** = 0.66
**$^1$H-NMR** (600 MHz, $CDCl_3$): *δ* (ppm) = 7.21 (d, J = 8.1 Hz, 3H), 7.13 (d, J = 6.9 Hz, 2H), 6.59 (s, 1H), 6.52 (s, 1H), 5.69 (s, 1H), 5.22 (s, 2H), 4.73 (s, 1H), 3.85 (s, 2H), 3.56 (t, J = 6.3 Hz, 2H), 3.40 (q, J = 7.0 Hz, 4H), 3.15 (d, J = 6.3 Hz, 2H), 2.66 (t, J = 7.4 Hz, 2H), 2.07 (p, J = 6.6 Hz, 2H), 1.67 (p, J = 7.7 Hz, 2H), 1.55 – 1.47 (m, 2H), 1.43 (s, 9H), 1.20 (t, J = 7.1 Hz, 6H).
**$^{13}$C-NMR** (151 MHz, $CDCl_3$): *δ* (ppm) = 161.9, 156.2, 154.9, 150.2, 142.0, 138.6, 129.6, 127.4, 124.4, 109.2, 98.4, 79.1, 62.6, 48.6, 45.2, 42.2, 40.6, 35.2, 31.3, 29.8, 29.2, 28.6, 12.5.
**HRMS** (ESI+): m/z calc. for $C_{33}H_{44}ClN_3NaO_6^+$ $[M+Na]^+$: 636.2811, found: 636.2793

#### 10.2.3 Fluorogenic CHalo-SiR conjugates

##### SiR-COOH

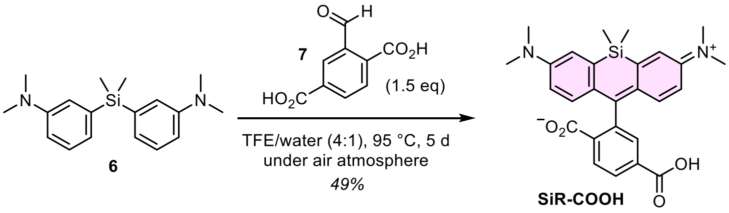

**6** (previously described[83]) was reacted with **7** (previously described[84]) adapting known procedures for xanthene synthesis.[84,85]

**6** (0.17 g, 0.58 mmol, 1.0 eq) was dissolved in trifluoroethanol/water (4:1, 4 mL), **7** (0.17 g, 0.88 mmol, 1.5 eq) was added and the reaction was heated to 95 °C in a pressure tube for 5 d. The volatiles were removed under reduced pressure and the crude product was purified by *rp*-flash column chromatography (acetonitrile/water; 25→60% MeCN) to give **SiR-COOH** (0.13 g, 0.28 mmol, 49%) as a blue solid. Analytical data matched literature values.[10]

**CA-SiR** was synthesised according to a previously described procedure (Lukinavičius *et al.*[10], compound SiR-Halo).

##### CHalo-SiR

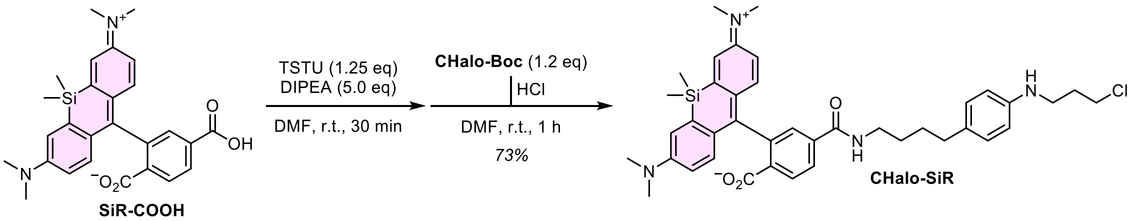

**CHalo-Boc** (13 mg, 38 µmol, 1.2 eq) was coupled with **SiR-COOH** following General Procedure E. Purification by *rp*-flash column chromatography (acetonitrile/water; 20→70% MeCN) affording **CHalo-SiR** (16 mg, 23 µmol, 73%) as a light-blue solid.

**$^1$H-NMR** (600 MHz, $CDCl_3$): *δ* (ppm) = 7.97 (dd, J = 8.0, 0.8 Hz, 1H), 7.88 (dd, J = 8.0, 1.4 Hz, 1H), 7.61 (dd, J = 1.4, 0.8 Hz, 1H), 6.98 – 6.94 (m, 4H), 6.76 (d, J = 8.9 Hz, 2H), 6.56 (dd, J = 9.0, 2.9 Hz, 2H), 6.52 (d, J = 8.5 Hz, 2H), 6.10 (t, J = 5.8 Hz, 1H), 3.64 (t, J = 6.3 Hz, 2H), 3.43 – 3.36 (m, 2H), 3.29 (t, J = 6.6 Hz, 2H), 2.97 (s, 12H), 2.52 (t, J = 7.1 Hz, 2H), 2.04 (p, J = 6.5 Hz, 2H), 1.66 – 1.54 (m, 4H), 0.67 (s, 3H), 0.59 (s, 3H).
**$^{13}$C-NMR** (151 MHz, $CDCl_3$): *δ* (ppm) = 170.1, 166.3, 155.4, 149.5, 146.2, 140.1, 136.9, 131.2, 131.0, 129.3, 129.2, 128.3, 127.7, 126.0, 122.9, 116.6, 113.6, 113.0, 92.1, 42.8, 41.2, 40.4, 40.3, 34.6, 32.1, 29.1, 29.0, 0.5, -1.1.
**HRMS** (ESI+): m/z calc. for $C_{40}H_{48}ClN_4O_3Si^+$ $[M]^+$: 695.3179, found: 695.3162

##### Synthetic route for Bn-CHalo-SiR

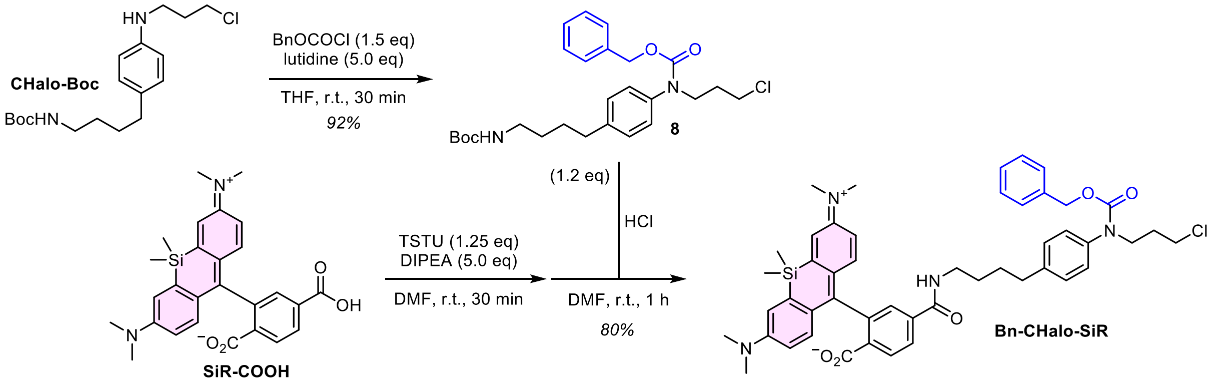

**Compound 8**

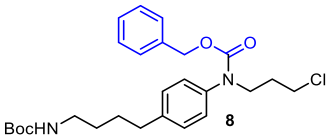

Adaptation from previously described procedure.[82] Under nitrogen atmosphere, **CHalo-Boc** (50 mg, 0.15 mmol, 1.0 eq) was dissolved in anhydrous THF (3 mL). 2,6-Lutidine (85 µL, 0.73 mmol, 5.0 eq) and benzyl chloroformate (32 µL, 0.22 mmol, 1.5 eq) were added and the reaction mixture was stirred at room temperature for 30 min. The solvent was removed under reduced pressure, water (20 mL) was added and the mixture was extracted with ethyl acetate (3 × 15 mL). The organic layer was dried over sodium sulfate, the desiccant was filtered off and the solvent was removed *in vacuo*. The crude product was purified by *np*-flash column chromatography (*iso*-hexanes/ethyl acetate; 0→40% EtOAc) to give **8** (64 mg, 0.14 mmol, 92%) as a colourless solid.

**TLC** *Rf* = 0.33 (n*p*, Hex:EtOAc 3:1); Rf of **CHalo-Boc** = 0.43
**$^1$H-NMR** (400 MHz, $CDCl_3$): *δ* (ppm) = 7.35 – 7.21 (m, 5H), 7.16 (d, *J* = 8.3 Hz, 2H), 7.09 (d, *J* = 7.7 Hz, 2H), 5.14 (s, 2H), 4.50 (s, 1H), 3.85 – 3.78 (m, 2H), 3.53 (t, *J* = 6.5 Hz, 2H), 3.18 – 3.10 (m, 2H), 2.62 (t, *J* = 7.6 Hz, 2H), 2.04 (p, *J* = 6.6 Hz, 2H), 1.69 – 1.63 (m, 2H), 1.56 – 1.48 (m, 2H), 1.44 (s, 9H).
**$^{13}$C-NMR** (101 MHz, $CDCl_3$): *δ* (ppm) = 156.1, 155.7, 141.1, 139.4, 136.7, 129.2, 128.5, 128.0, 127.4, 127.1, 79.3, 67.3, 48.3, 42.4, 40.5, 35.2, 31.5, 29.8, 28.6.
**HRMS** (ESI+): m/z calc. for $C_{26}H_{35}ClN_2NaO_4^+$ [M+Na]$^+$: 497.2178, found: 497.2161

**Bn-CHalo-SiR**

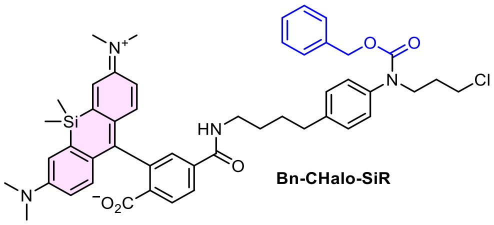

**8** (18 mg, 38 µmol, 1.2 eq) was coupled with **SiR-COOH** following General Procedure E. Purification by *rp*-flash column chromatography (acetonitrile/water; 25→85% MeCN) affording **Bn-CHalo-SiR** (16 mg, 23 µmol, 73%) as a light-blue solid.

**$^1$H-NMR** (600 MHz, $CDCl_3$): *δ* (ppm) = 7.96 (dd, J = 8.0, 0.7 Hz, 1H), 7.89 (dd, J = 8.0, 1.4 Hz, 1H), 7.62 (s, 1H), 7.36 – 7.19 (m, 5H), 7.12 (d, J = 8.0 Hz, 2H), 7.06 (d, J = 8.0 Hz, 2H), 6.96 (d, J = 2.9 Hz, 2H), 6.76 (d, J = 9.0 Hz, 2H), 6.55 (dd, J = 9.0, 2.9 Hz, 2H), 6.28 (s, 1H), 5.12 (s, 2H), 3.82 – 3.77 (m, 2H), 3.51 (t, J = 6.5 Hz, 2H), 3.39 (q, J = 6.8 Hz, 2H), 2.96 (s, 12H), 2.61 (t, J = 7.4 Hz, 2H), 2.02 (p, J = 6.7 Hz, 2H), 1.65 (p, J = 7.9 Hz, 2H), 1.58 (q, J = 7.2 Hz, 2H), 0.66 (s, 3H), 0.59 (s, 3H).
**$^{13}$C-NMR** (151 MHz, $CDCl_3$): *δ* (ppm) = 170.1, 166.3, 155.6, 155.4, 149.5, 140.8, 140.0, 139.5, 136.8, 136.7, 131.2, 129.2, 129.1, 128.5, 128.3, 128.0, 127.7, 127.1, 126.0, 123.0, 116.6, 113.6, 92.1, 67.3, 48.3, 42.3, 40.3, 40.2, 35.1, 31.5, 29.2, 28.6, 0.4, -1.1.
**HRMS** (ESI+): m/z calc. for $C_{48}H_{54}ClN_4O_5Si^+$ [M+H]$^+$: 829.3547, found: 829.3532

**Cou-CHalo-SiR**

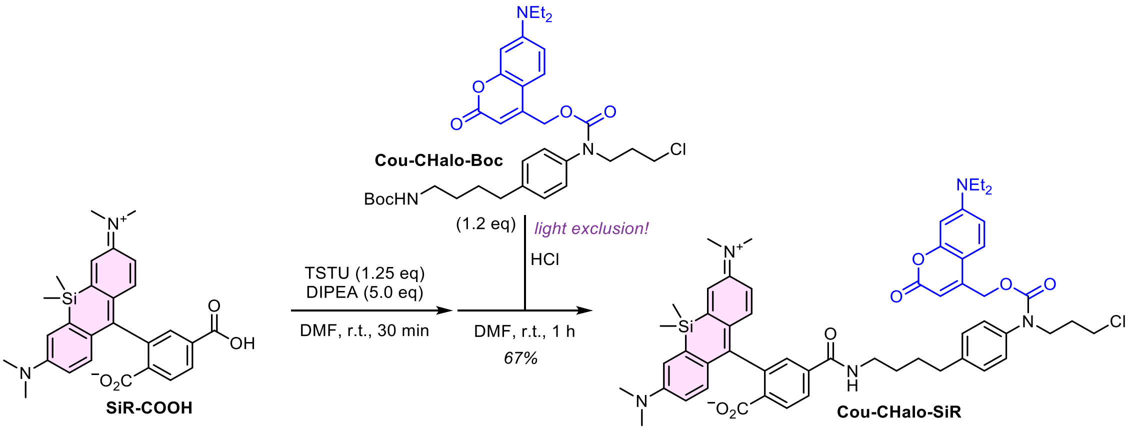

*Note: the reaction must be performed under light exclusion.*

**Cou-CHalo-Boc** (32 mg, 52 µmol, 1.2 eq) was coupled with **SiR-COOH** following General Procedure E. Purification by *rp*-flash column chromatography (acetonitrile/water; 30→90% MeCN) followed by *np*-flash column chromatography (DCM/MeOH; 0→3% MeOH) affording **Cou-CHalo-SiR** (28 mg, 29 µmol, 67%) as a yellow solid.

**TLC** *Rf* = 0.44 (*np*, DCM:MeOH 19:1)
**$^1$H-NMR** (400 MHz, DMSO-d$_6$): $\delta$ (ppm) = 8.73 (t, J = 5.5 Hz, 1H), 8.06 (d, J = 8.8 Hz, 1H), 8.01 (d, J = 8.0 Hz, 1H), 7.64 (s, 1H), 7.42 – 7.34 (m, 1H), 7.26 – 7.17 (m, 5H), 7.01 (d, J = 1.7 Hz, 2H), 6.66 – 6.60 (m, 5H), 6.50 (s, 1H), 5.24 (s, 2H), 3.78 – 3.73 (m, 2H), 3.63 (t, J = 6.4 Hz, 2H), 3.41 (q, J = 6.9 Hz, 4H), 3.27 – 3.21 (m, 2H), 2.90 (s, 12H), 2.62 – 2.55 (m, 2H), 1.90 (p, J = 6.6 Hz, 2H), 1.63 – 1.45 (m, 4H), 1.10 (t, J = 7.0 Hz, 6H), 0.63 (s, 3H), 0.51 (s, 3H).
**$^{13}$C-NMR** (151 MHz, CDCl$_3$): $\delta$ (ppm) = 170.1, 166.4, 162.3, 156.0, 154.8, 154.3, 150.8, 150.4, 149.4, 141.9, 140.2, 138.6, 137.2, 131.7, 129.7, 129.1, 128.2, 127.8, 127.6, 125.7, 124.3, 123.9, 116.8, 113.3, 108.9, 105.6, 104.5, 97.7, 92.4, 62.4, 48.2, 44.8, 42.2, 40.3, 35.1, 31.2, 28.7, 12.5, 0.6, -1.5.
**HRMS** (ESI+): m/z calc. for $C_{55}H_{63}ClN_5O_7Si^+$ [M+H]$^+$: 968.4180, found: 968.4176

##### Synthetic route for Leu-CHalo-SiR

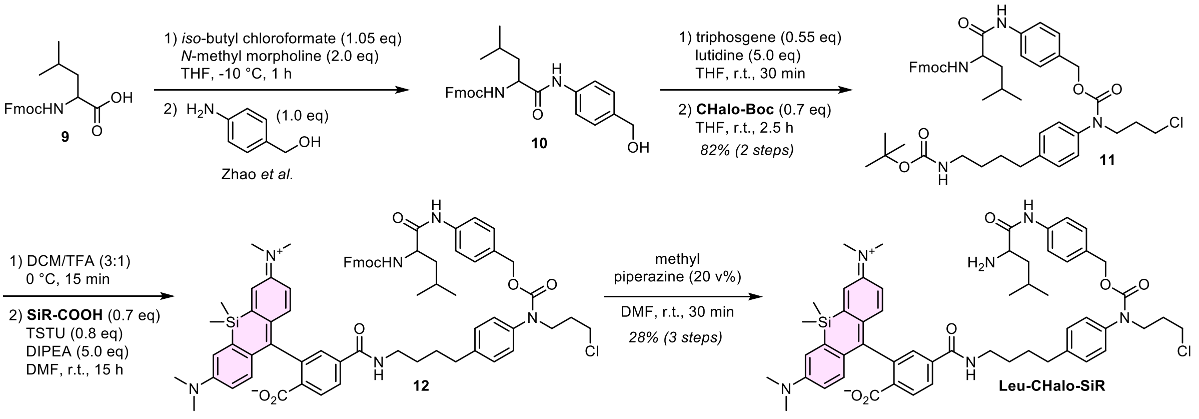

Compound **10** was prepared according to a previously described procedure (Zhao *et al.*[86], compound 3).

##### Compound 11

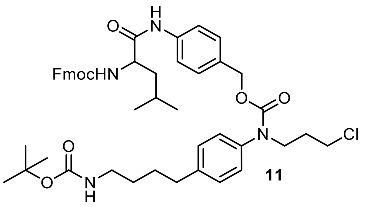

Adaptation from previously described procedure.[82] Under nitrogen atmosphere, **10** (10 mg, 22 µmol, 1.0 eq) was dissolved in anhydrous THF (1 mL). Triphosgene (0.6 eq) and 2,6-lutidine (5.0 eq) were added and the reaction mixture was stirred at room temperature for 15 min. In a separate flask, **CHalo-Boc** (2 mg, 6 µmol, 0.7 eq) was dissolved in anhydrous THF (0.5 mL) under nitrogen atmosphere, the **10**-chloroformate was added. dropwise and the reaction mixture was stirred at room temperature for 20 min. Methyl piperazine (2 µL, 18 µmol, 3.0 eq) was added to react with the excess of the chloroformate. The mixture was diluted with water (10 mL) and extracted with ethyl acetate (3 × 10 mL). The organic layer was dried over sodium sulfate, the desiccant was filtered off and the solvent was removed *in vacuo*. The crude product was purified by *rp*-flash column chromatography (acetonitrile/water; 20→100% MeCN) to give **11** (4 mg, 5 µmol, 82% over 2 steps) as a colourless solid.

**TLC** *Rf* = 0.69 (*np*, Hex:EtOAc 1:1)
**$^1$H-NMR** (600 MHz, CDCl$_3$): $\delta$ (ppm) = 8.14 (s, 1H), 7.76 (d, J = 7.4 Hz, 2H), 7.57 (t, J = 7.0 Hz, 2H), 7.43 (s, 2H), 7.38 (t, J = 7.3 Hz, 2H), 7.29 – 7.27 (m, 2H), 7.14 (d, J = 8.2 Hz, 2H), 7.08 (d, J = 8.3 Hz, 4H), 5.30 (s, 1H), 5.05 (s, 2H), 4.57 (s, 1H), 4.48 – 4.38 (m, 2H), 4.29 (s, 1H), 4.20 (t, J = 6.7 Hz, 1H), 3.80 (t, J = 6.9 Hz, 2H), 3.53 (s, 2H), 3.13 (s, 2H), 2.62 (t, J = 7.5 Hz, 2H), 2.06 – 1.99 (m, 2H), 1.70 (s, 1H), 1.66 – 1.58 (m, 4H), 1.55 – 1.46 (m, 2H), 1.43 (s, 9H), 1.01 – 0.91 (m, 6H).
**$^{13}$C-NMR** (151 MHz, CDCl$_3$): $\delta$ (ppm) = 155.6, 143.8, 141.4, 139.3, 132.7, 129.2, 127.9, 127.2, 127.2, 125.1, 120.2, 119.9, 67.2, 54.4, 48.1, 47.3, 42.4, 40.8, 40.5, 35.2, 31.5, 29.9, 29.8, 28.7, 28.6, 24.9, 23.1, 22.1.
**HRMS** (ESI+): m/z calc. for $C_{47}H_{57}ClN_4NaO_7^+$ [M+Na]$^+$: 847.3808, found: 847.3806.

#### Leu-CHalo-SiR

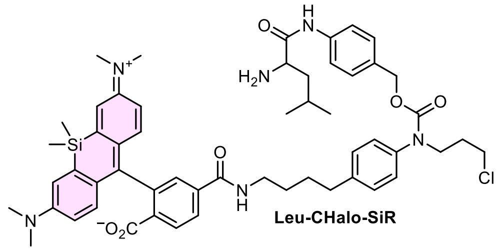

Leu-CHalo-SiR

**11** (2 mg, 2 µmol, 1.3 eq) was Boc-deprotected in DCM / trifluoroacetic acid (3:1, 1.33 mL) at 0 °C for 15 min. The trifluoroacetic acid was quenched by slowly adding aqueous sodium carbonate solution until no gas development was observed upon addition. The intermediate was coupled with the NHS ester of **SiR-COOH** (1.0 eq) following General Procedure E to give **12** which was Fmoc-deprotected by adding methyl piperazine (20 vol%) to the coupling reaction mixture in DMF which was stirred at room temperature for 30 min. The crude product was purified by preparative HPLC (acetonitrile/water, 0.1% formic acid; 15→70% MeCN, 20 min) affording **Leu-CHalo-SiR** (0.37 mg, 0.39 µmol, 28% over 3 steps) as a light-blue solid.

*Note regarding the Boc-deprotection: 2M HCl in DCM/dioxane (1:1) decomposed the carbamate to the undesired benzyl chloride; DCM/TFA (1:1) at r.t. gave the benzylic TFA-adduct.*

*Note regarding the Fmoc-deprotection: methyl piperazine was used instead of the standard protocol with piperidine for easier separation of the fluorenyl deprotection side product.*

**$^1$H-NMR** (600 MHz, DMSO-d$_6$): $\delta$ (ppm) = 8.74 (d, J = 5.7 Hz, 1H), 8.43 (s, 4H), 8.06 (dd, J = 8.1, 1.3 Hz, 1H), 8.01 (d, J = 8.1 Hz, 1H), 7.64 (s, 1H), 7.58 (d, J = 7.4 Hz, 1H), 7.17 (d, J = 8.5 Hz, 2H), 7.13 (d, J = 8.5 Hz, 2H), 7.01 (d, J = 2.6 Hz, 2H), 6.66 – 6.60 (m, 4H), 4.99 (s, 2H), 3.74 – 3.68 (m, 2H), 3.60 (t, J = 6.4 Hz, 2H), 2.91 (s, 12H), 2.68 (t, J = 4.8 Hz, 3H), 2.58 – 2.53 (m, 4H), 2.53 – 2.51 (m, 4H), 1.91 – 1.83 (m, 2H), 1.59 – 1.52 (m, 2H), 1.50 (d, J = 6.8 Hz, 2H), 0.89 (d, J = 6.6 Hz, 3H), 0.87 (d, J = 6.6 Hz, 3H), 0.63 (s, 3H), 0.52 (s, 3H).
**HRMS** (ESI+): m/z calc. for $C_{55}H_{63}ClN_5O_7Si^+$ [M+H]$^+$: 968.4180, found: 968.4176

#### Synthetic route for SS66T-CHalo-SiR (GSH activation)

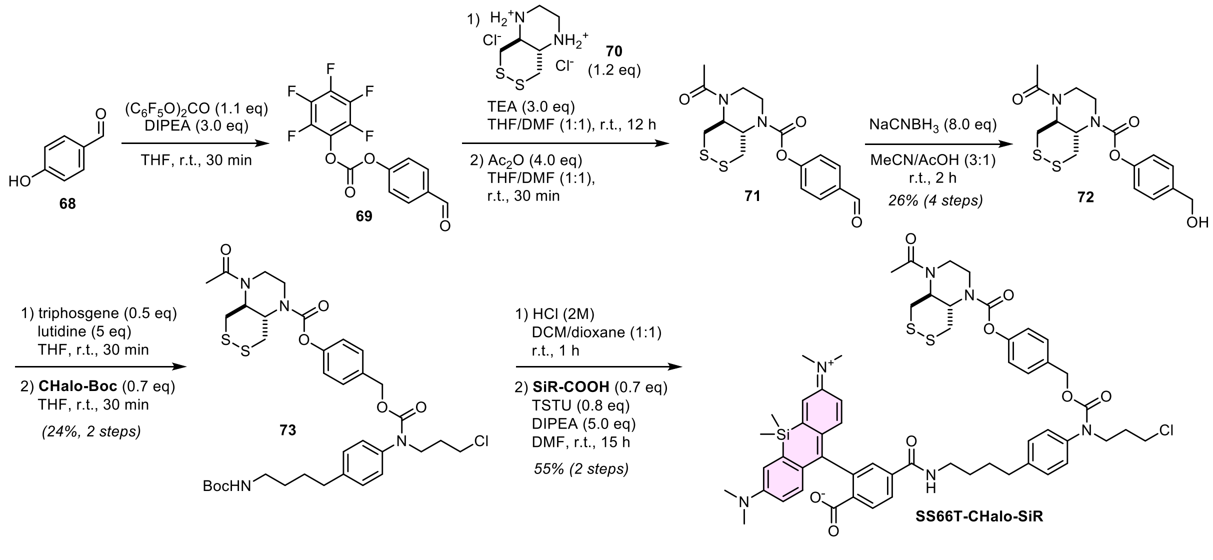

#### Compound 72

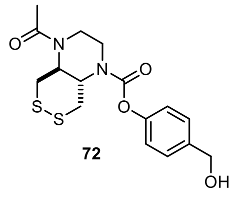

4-Hydroxybenzaldehyde (**68**, 37 mg, 0.30 mmol, 1.0 eq) was dissolved in anhydrous THF (1.1 mL). DIPEA (0.15 mL, 0.90 mmol, 3.0 eq) and bis(pentafluorophenyl)carbonate (201 mg, 0.51 mmol, 1.7 eq) were added and the reaction mixture was stirred at room temperature for 30 min. Diamine **70** (90 mg, 0.36 mmol, 1.2 eq), DIPEA (0.15 mL, 0.90 mmol, 3.0 eq) and anhydrous DMF (1.1 mL) were added and the mixture stirred at room temperature for 12 h. Then, acetic anhydride (0.11 mL, 1.2 mmol, 4.0 eq) was added, the mixture was stirred at room temperature for 30 min and the excess of the acetic anhydride was removed upon addition of *N*-methyl piperazine (0.20 mL, 1.8 mmol, 6.0 eq). The volatiles were removed *in vacuo* and the crude product was semi-purified by *rp*- flash column chromatography (acetonitrile/water; 5→50% MeCN). **71** was dissolved in acetonitrile (1.5 mL) and acetic acid (0.5 mL), sodium cyanoborohydride (75 mg, 1.2 mmol, 4.0 eq) was added and the reaction mixture was stirred at room temperature for 2 h. DCM (10 mL) and a half-saturated solution of sodium bicarbonate (15 mL) were added, the layers were separated and the aqueous layer was extracted with DCM (2 ×10 mL). The organic layer was dried over sodium sulfate, the desiccant was filtered off and the solvent was removed *in vacuo*. The crude product was purified by *rp*-flash column chromatography (acetonitrile/water; 5→35% MeCN) to give **72** (29 mg, 79 µmol, 26% over 4 steps) as a colourless oil.

**TLC** *Rf* = 0.37 (*np*, DCM:MeOH 19:1)
**$^1$H-NMR** (400 MHz, $CDCl_3$): *δ* (ppm) = 7.35 (d, *J* = 8.5 Hz, 2H), 7.05 (d, *J* = 8.0 Hz, 2H), 4.66 (d, *J* = 2.7 Hz, 2H), 4.29 (s, 1H), 4.16 – 3.81 (m, 2H), 3.80 – 3.41 (m, 4H), 3.30 (dd, *J* = 13.6, 2.6 Hz, 1H), 3.24 – 2.84 (m, 1H), 2.11 (s, 3H), 2.02 (s, 1H).
**HRMS** (ESI+): m/z calc. for $C_{16}H_{20}N_2NaO_4S_2^+$ [M+Na]$^+$: 391.0757, found: 391.0749.

**Compound 73**

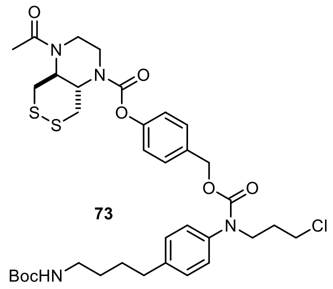

73

Under nitrogen atmosphere, **72** (5.0 mg, 14 µmol, 1.2 eq) was dissolved in anhydrous THF (0.25 mL). Triphosgene (2.8 mg, 9.5 µmol, 0.8 eq) was added from a fresh stock solution, then 2,6-lutidine (7.9 µL, 68 µmol, 6.0 eq) was added and the reaction mixture was stirred at room temperature for 20 min to form the chloroformate (HPLC-MS analysis of the reaction progress by quenching with piperidine in acetonitrile).

In a separate flask, **CHalo-Boc** (3.9 mg, 11 µmol, 1.0 eq) was dissolved in anhydrous THF (0.3 mL) under nitrogen atmosphere, added to the chloroformate and the reaction mixture was stirred at room temperature for 30 min. The mixture was diluted with DCM (10 mL) and water (15 mL), the layers were separated and the aqueous layer was extracted with DCM (2 ×10 mL). The organic layer was dried over sodium sulfate, the desiccant was filtered off and the solvent was removed *in vacuo*. The crude product was purified by *rp*-flash column chromatography (acetonitrile/water; 20→100% MeCN) to give **73** (2 mg, 3 µmol, 24% over 2 steps) as a colourless solid.

**TLC** *Rf* = 0.74 (*np*, DCM/MeOH 47:3)
**$^1$H-NMR** (400 MHz, $CDCl_3$): *δ* (ppm) = 7.23 (d, *J* = 8.3 Hz, 2H), 7.16 (d, *J* = 8.4 Hz, 2H), 7.07 (d, *J* = 7.8 Hz, 2H), 7.04 – 6.98 (m, 2H), 5.10 (s, 2H), 4.71 – 4.17 (m, 2H), 3.98 (d, *J* = 48.0 Hz, 1H), 3.84 – 3.76 (m, 2H), 3.67 (s, 3H), 3.53 (t, *J* = 6.5 Hz, 2H), 3.34 – 2.88 (m, 5H), 2.63 (t, *J* = 7.6 Hz, 2H), 2.12 (s, 3H), 2.08 – 1.98 (m, 2H), 1.70 – 1.57 (m, 4H), 1.57 – 1.47 (m, 2H), 1.44 (s, 9H).
**HRMS** (ESI+): m/z calc. for $C_{35}H_{47}ClN_4NaO_7S_2^+$ [M+Na]$^+$: 757.2467, found: 757.2438.

**SS66T-CHalo-SiR**

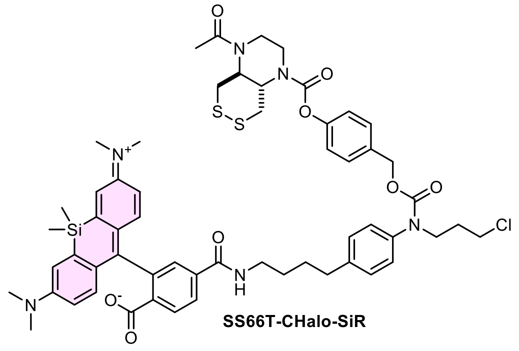

SS66T-CHalo-SiR

**73** (2 mg, 3 µmol, 1.7 eq) was coupled with **SiR-COOH** following General Procedure E. Purification by reparative HPLC (acetonitrile/water, 0.1% formic acid; 20→80% MeCN, 20 min) affording **SS66T-CHalo-SiR** (0.99 mg, 0.91 µmol, 57%) as a light-blue solid.

**HRMS** (ESI+): m/z calc. for $C_{57}H_{66}ClN_6O_8S_2Si^+$ [M+H]$^+$: 1089.384, found: 1089.382.

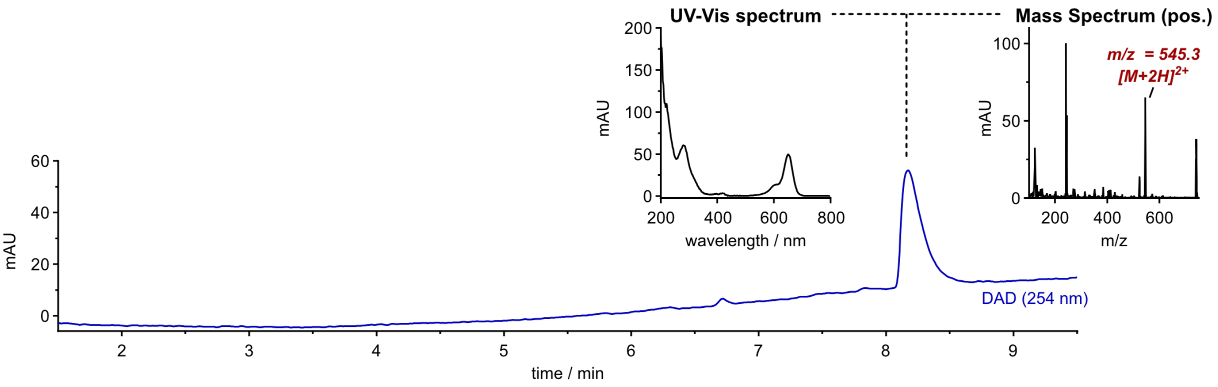

#### Synthetic route for SS66C-CHalo-SiR (thioredoxin (Trx) activation)

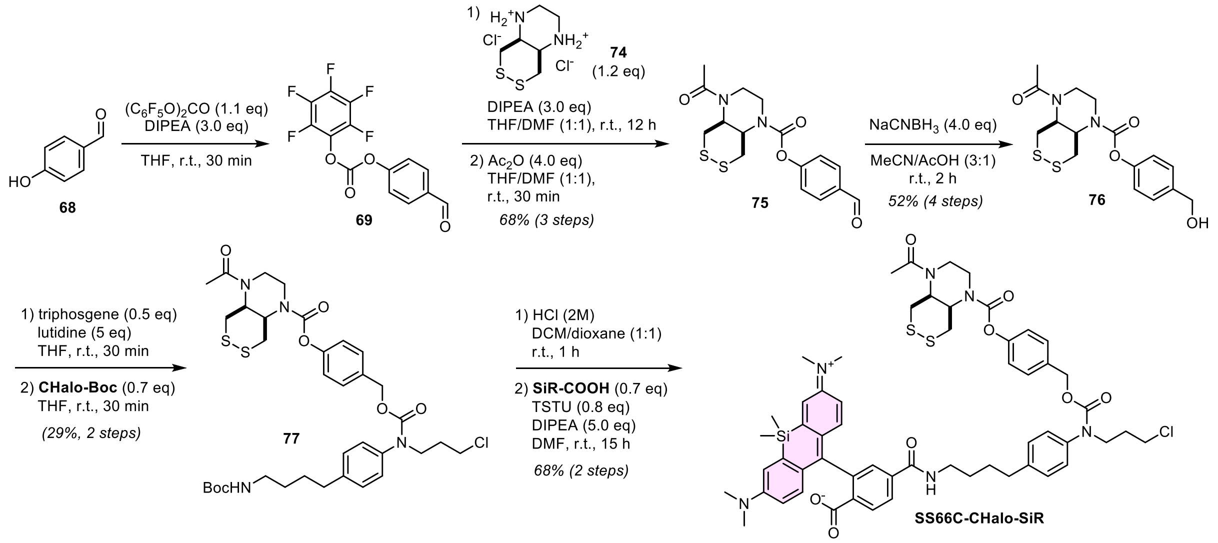

#### Compound 76

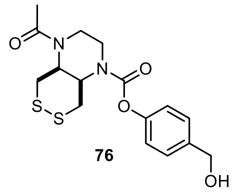

76

4-Hydroxybenzaldehyde (**68**, 12 mg, 0.10 mmol, 1.0 eq) was dissolved in anhydrous THF (0.4 mL). DIPEA (0.05 mL, 0.30 mmol, 3.0 eq) and bis(pentafluorophenyl)carbonate (67 mg, 0.17 mmol, 1.7 eq) were added and the reaction mixture was stirred at room temperature for 30 min. Diamine **74** (30 mg, 0.12 mmol, 1.2 eq), DIPEA (0.05 mL, 0.30 mmol, 3.0 eq) and anhydrous DMF (0.4 mL) were added and the mixture stirred at room temperature for 12 h. Then, acetic anhydride (38 µL, 0.40 mmol, 4.0 eq) was added, the mixture was stirred at room temperature for 30 min and the excess of the acetic anhydride was removed upon addition of *N*-methyl piperazine (67 µL, 0.60 mmol, 6.0 eq). The volatiles were removed *in vacuo* and the crude product was semi-purified by *rp*- flash column chromatography (acetonitrile/water; 5→50% MeCN). **75** was dissolved in acetonitrile (1.5 mL) and acetic acid (0.5 mL), sodium cyanoborohydride (25 mg, 0.40 mmol, 4.0 eq) was added and the reaction mixture was stirred at room temperature for 2 h. DCM (10 mL) and a half-saturated solution of sodium bicarbonate (15 mL) were added, the layers were separated and the aqueous layer was extracted with DCM (2 ×10 mL). The organic layer was dried over sodium sulfate, the desiccant was filtered off and the solvent was removed *in vacuo*. The crude product was purified by *rp*-flash column chromatography (acetonitrile/water; 5→35% MeCN) to give **72** (19 mg, 52 µmol, 52% over 4 steps) as a colourless oil.

**TLC** *Rf* = 0.37 (*np*, DCM:MeOH 19:1)
**$^1$H-NMR** (400 MHz, CDCl$_3$): *δ* (ppm) = 7.38 (d, *J* = 8.5 Hz, 2H), 7.10 (d, *J* = 8.6 Hz, 2H), 4.69 (s, 2H), 4.40 (td, *J* = 6.1, 3.4 Hz, 1H), 4.23 (s, 1H), 3.90 – 3.80 (m, 2H), 3.79 – 3.65 (m, 2H), 3.50 (dd, *J* = 14.0, 9.0 Hz, 1H), 3.16 (dd, *J* = 14.4, 3.4 Hz, 1H), 2.97 (dd, *J* = 13.5, 4.1 Hz, 1H), 2.15 (s, 3H), 1.83 (d, *J* = 11.5 Hz, 1H).
**$^{13}$C-NMR** (151 MHz, CDCl$_3$): *δ* (ppm) = 170.3, 154.0, 150.2, 138.5, 128.1, 121.7, 64.7, 51.3, 43.3, 42.0, 38.6, 36.2, 21.9; *one expected carbon peak not observed.*
**HRMS** (ESI+): m/z calc. for $C_{16}H_{20}N_2NaO_4S_2^+$ $[M+Na]^+$: 391.0757, found: 391.0746.

**Compound 77**

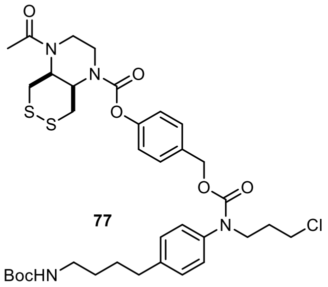

Under nitrogen atmosphere, **76** (6.3 mg, 17 µmol, 1.2 eq) was dissolved in anhydrous THF (0.25 mL). Triphosgene (3.6 mg, 12 µmol, 0.8 eq) was added from a fresh stock solution, then 2,6-lutidine (10 µL, 86 µmol, 6.0 eq) was added and the reaction mixture was stirred at room temperature for 20 min to form the chloroformate (HPLC-MS analysis of the reaction progress by quenching with piperidine in acetonitrile).

In a separate flask, **CHalo-Boc** (4.9 mg, 14 µmol, 1.0 eq) was dissolved in anhydrous THF (0.3 mL) under nitrogen atmosphere, added to the chloroformate and the reaction mixture was stirred at room temperature for 30 min. The mixture was diluted with DCM (10 mL) and water (15 mL), the layers were separated and the aqueous layer was extracted with DCM (2 ×10 mL). The organic layer was dried over sodium sulfate, the desiccant was filtered off and the solvent was removed *in vacuo*. The crude product was purified by *rp*-flash column chromatography (acetonitrile/water; 20→100% MeCN) to give **77** (3 mg, 4 µmol, 29% over 2 steps) as a colourless solid.

**TLC** *Rf* = 0.74 (*np*, DCM/MeOH 47:3)
**$^1$H-NMR** (400 MHz, $CDCl_3$): *δ* (ppm) = 7.23 (d, *J* = 8.4 Hz, 2H), 7.15 (d, *J* = 8.3 Hz, 2H), 7.07 (t, *J* = 7.3 Hz, 4H), 5.11 (s, 2H), 4.40 (td, *J* = 6.1, 3.3 Hz, 1H), 4.28 – 4.17 (m, 1H), 3.90 – 3.63 (m, 6H), 3.60 – 3.45 (m, 3H), 3.19 – 3.07 (m, 3H), 3.03 – 2.93 (m, 1H), 2.66 – 2.59 (m, 2H), 2.15 (s, 3H), 2.07 – 1.98 (m, 2H), 1.72 – 1.57 (m, 4H), 1.57 – 1.47 (m, 2H), 1.44 (s, 9H).
**HRMS** (ESI+): m/z calc. for $C_{35}H_{47}ClN_4NaO_7S_2^+$ [M+Na]$^+$: 757.2467, found: 757.2452.

**SS66C-CHalo-SiR**

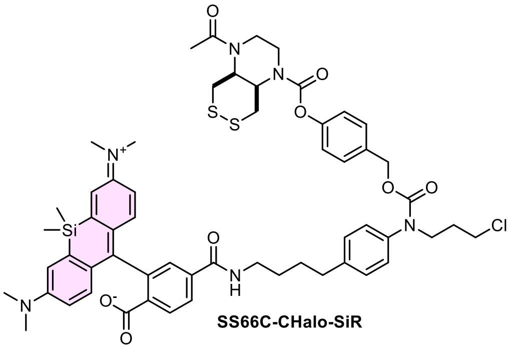

**77** (3 mg, 4 µmol, 2.6 eq) was coupled with **SiR-COOH** following General Procedure E. Purification by reparative HPLC (acetonitrile/water, 0.1% formic acid; 20→80% MeCN, 20 min) affording **SS66T-CHalo-SiR** (1.2 mg, 1.1 µmol, 68%) as a light-blue solid.

**HRMS** (ESI+): m/z calc. for $C_{57}H_{66}ClN_6O_8S_2Si^+$ [M+H]$^+$: 1089.384, found: 1089.382.

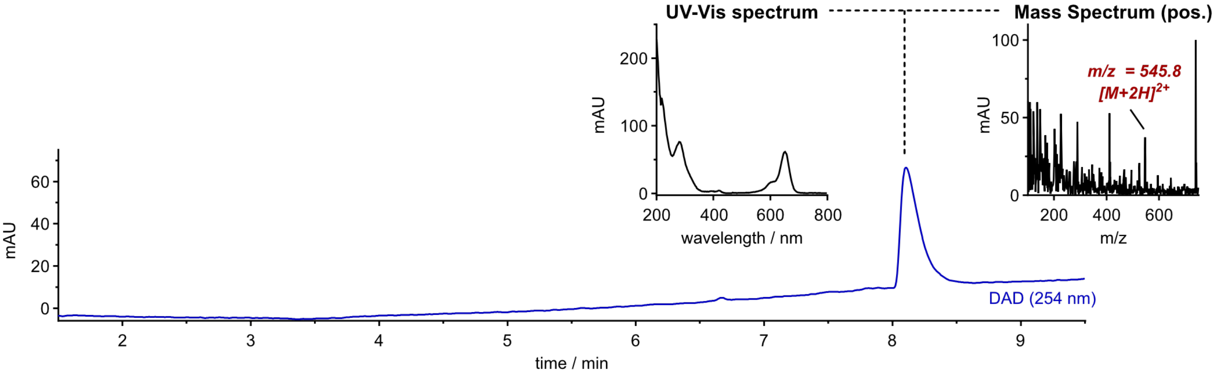

##### 10.2.4 Light-controlled protein heterodimeriser Cou-CHalo-BG

###### Synthetic route for Cou-CHalo-BG

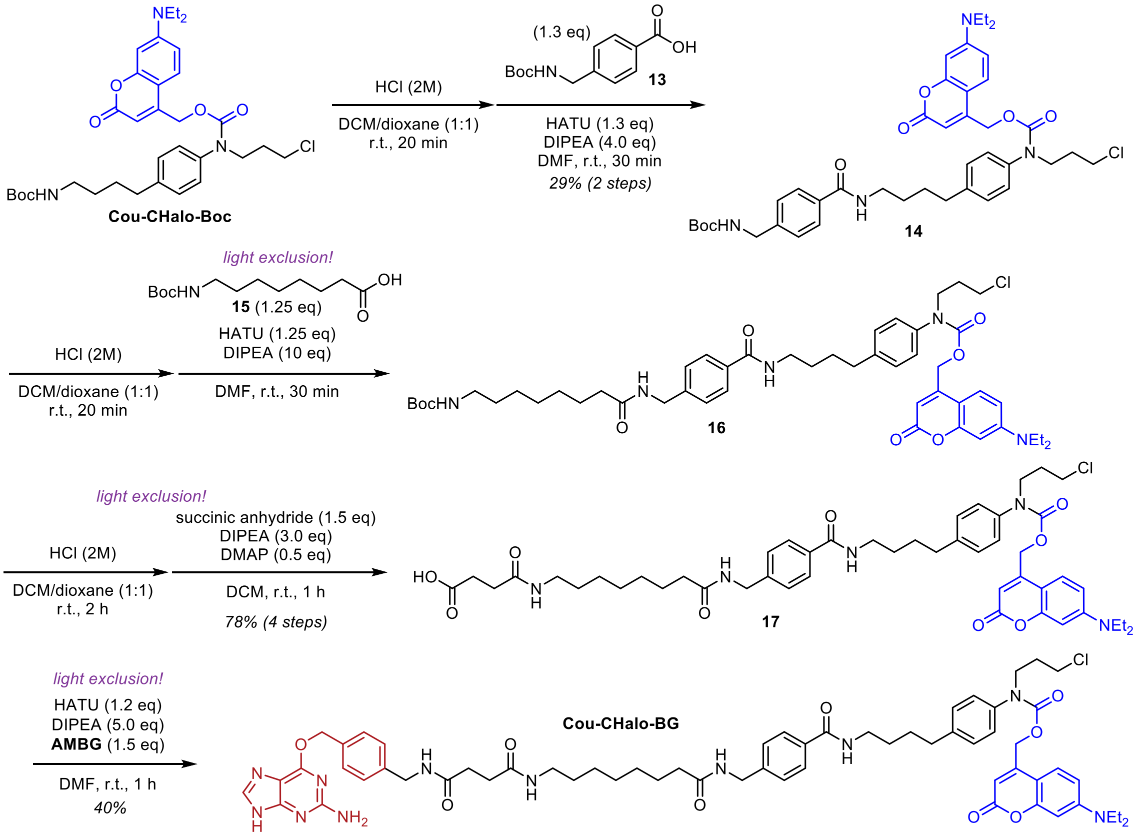

###### Compound 14

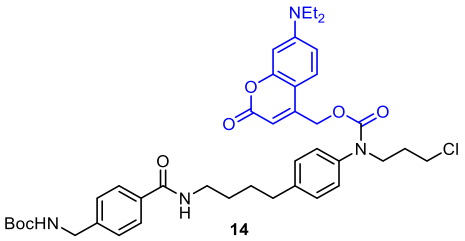

14

*Note: the reaction must be performed under light exclusion.*

**Cou-CHalo-Boc** (29 mg, 47 µmol, 1.0 eq) was deprotected according to General Procedure A (0.1 M) and coupled with carboxylate **13** (16 mg, 61 µmol, 1.3 eq, 0.02 M) following General Procedure B (1.0 eq HATU, 30 min reaction time). Piperidine (6 µL, 0.06 mmol, 1.3 eq) was added to react with excess carboxylate. The volatiles were removed *in vacuo*, the residue was diluted with water (15 mL) and brine (10 mL) and extracted with ethyl acetate (3 × 10 mL). The organic layer was dried over sodium sulfate, the desiccant was filtered off and the solvent was removed *in vacuo*. The crude product was purified by flash *np*-column chromatography (*iso*-hexanes/ethyl acetate; 10→70% EtOAc) to give **14** (31 mg, 42 µmol, 88%) as a yellow solid.

**TLC** *Rf* = 0.17 (*np*, Hex:EtOAc 1:1); 0.47 (*np*, DCM:MeOH 19:1)
**$^1$H-NMR** (600 MHz, $CDCl_3$/MeOD (4:1)): $\delta$ (ppm) = 7.70 (s, 2H), 7.24 (d, J = 7.9 Hz, 2H), 7.17 (d, J = 7.6 Hz, 3H), 7.07 (d, J = 7.6 Hz, 2H), 6.54 (s, 1H), 6.43 (s, 1H), 5.48 (s, 1H), 5.14 (s, 2H), 4.23 (s, 2H), 3.81 – 3.75 (m, 2H), 3.51 (t, J = 5.9 Hz, 2H), 3.40 – 3.31 (m, 6H), 2.66 – 2.61 (m, 2H), 2.00 (p, J = 6.6 Hz, 2H), 1.72 – 1.63 (m, 2H), 1.58 – 1.51 (m, 2H), 1.37 (s, 9H), 1.13 (t, J = 7.0 Hz, 6H).
**$^{13}$C-NMR** (151 MHz, $CDCl_3$/MeOD (4:1)): $\delta$ (ppm) = 167.9, 162.5, 156.3, 155.9, 154.9, 150.6, 142.4, 141.9, 138.4, 133.5, 129.6, 127.4, 127.3, 124.2, 109.1, 105.8, 104.7, 97.7, 79.8, 62.5, 48.3, 44.9, 44.1, 42.1, 39.9, 35.0, 31.0, 28.8, 28.6, 28.3, 12.3.
**HRMS** (ESI+): m/z calc. for $C_{41}H_{52}ClN_4O_7^+$ $[M+H]^+$: 747.3519, found: 747.3524.

**Compound 17**

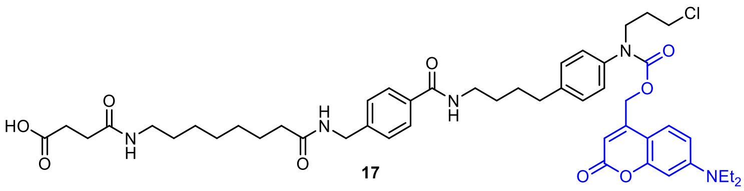

17

*Note: the reaction must be performed under light exclusion.*

**14** (31 mg, 42 µmol, 1.0 eq) was deprotected according to General Procedure A (0.04 M) and coupled with carboxylate **15** (14 mg, 52 µmol, 1.25 eq, 0.04 M) following General Procedure B (1.0 eq HATU, 30 min reaction time). The volatiles were removed *in vacuo* to give crude **16** which was Boc-deprotected according to General Procedure A (0.05 M) and dissolved in anhydrous DCM (2 mL). DIPEA (35 µL, 0.21 mmol, 5.0 eq), DMAP (3 mg, 0.02 mmol, 0.5 eq) and succinic anhydride (6 mg, 0.06 mmol, 1.5 eq) were added and the reaction mixture was stirred at room temperature for 20 min. The volatiles were removed *in vacuo* and the crude product was purified by *rp*-flash column chromatography (acetonitrile/water; 10→70% MeCN) to give **17** (28 mg, 32 µmol, 76% over 4 steps) as a yellow solid.

**$^1$H-NMR** (600 MHz, $CDCl_3$/MeOD (1:1)): $\delta$ (ppm) = 7.72 (d, J = 7.8 Hz, 2H), 7.27 (d, J = 8.3 Hz, 2H), 7.21 (d, J = 7.8 Hz, 3H), 7.11 (d, J = 7.9 Hz, 2H), 6.59 (s, 1H), 6.47 (s, 1H), 5.58 (s, 1H), 5.34 – 5.10 (m, 2H), 4.36 (s, 2H), 3.81 (s, 2H), 3.53 (t, J = 6.1 Hz, 2H), 3.40 (q, J = 6.1 Hz, 6H), 3.12 (t, J = 7.1 Hz, 2H), 2.70 – 2.63 (m, 2H), 2.57 (t, J = 6.9 Hz, 2H), 2.41 (t, J = 7.0 Hz, 2H), 2.18 (t, J = 7.5 Hz, 2H), 2.03 (p, J = 6.5 Hz, 2H), 1.70 (s, 2H), 1.60 (dq, J = 14.5, 7.4 Hz, 4H), 1.42 (q, J = 7.0 Hz, 2H), 1.27 (s, 6H), 1.17 (t, J = 7.1 Hz, 6H).
**$^{13}$C-NMR** (151 MHz, $CDCl_3$/MeOD (1:1)): $\delta$ (ppm) = 209.5, 175.4, 174.9, 173.0, 168.6, 163.1, 156.2, 155.2, 151.3, 150.8, 142.4, 142.0, 138.6, 133.6, 129.7, 127.8, 127.6, 127.5, 124.7, 109.5, 62.8, 45.2, 43.1, 42.2, 40.1, 39.6, 36.3, 35.3, 31.3, 30.9, 29.9, 29.3, 29.1, 29.1, 28.9, 28.8, 26.7, 25.8.
**HRMS** (ESI+): m/z calc. for $C_{48}H_{62}ClN_5NaO_9^+$ $[M+Na]^+$: 910.4128, found: 910.4131.

**Cou-CHalo-BG**

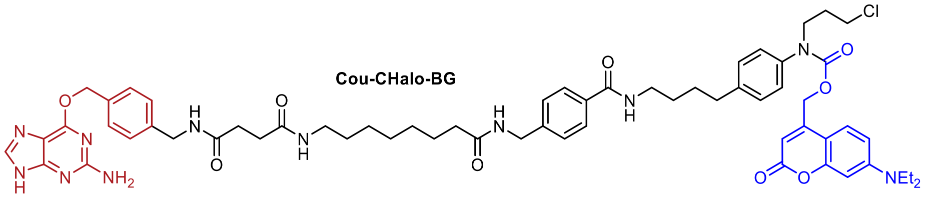

*Note: the reaction must be performed under light exclusion.*

6-((4-(aminomethyl)benzyl)oxy)-9H-purin-2-amine (**AMBG**, 13 mg, 47 µmol, 1.5 eq) was coupled with carboxylate **17** (28 mg, 32 µmol, 1.0 eq, 0.01 M) following General Procedure B (1.2 eq HATU, 1 h reaction time). The reaction mixture was filtered over Celite to remove the insoluble **AMBG**, the volatiles were removed under reduced pressure and the crude product was purified by *rp*-flash column chromatography (acetonitrile/water; 20→70% MeCN) followed by *np*-flash column chromatography (DCM/MeOH; 0.5→15% MeOH) affording **Cou-CHalo-BG** (14.4 mg, 13 µmol, 40%) as a yellow solid.

**TLC** *Rf* = 0.27 (*np*, DCM:MeOH 9:1)
**$^1$H-NMR** (600 MHz, DMSO-$d_6$): $\delta$ (ppm) = 12.49 (s, 1H), 8.41 (t, J = 5.7 Hz, 1H), 8.35 (td, J = 6.0, 3.5 Hz, 2H), 7.83 – 7.76 (m, 4H), 7.43 (d, J = 8.1 Hz, 2H), 7.37 (s, 1H), 7.28 (d, J = 8.1 Hz, 2H), 7.27 – 7.23 (m, 6H), 6.67 – 6.62 (m, 1H), 6.51 (s, 1H), 6.26 (s, 2H), 5.45 (s, 2H), 5.25 (s, 2H), 4.28 (d, J = 6.0 Hz, 2H), 4.25 (d, J = 5.9 Hz, 2H), 3.77 (s, 2H), 3.64 (t, J = 6.4 Hz, 2H), 3.41 (q, J = 7.0 Hz, 4H), 3.28 (q, J = 6.6 Hz, 2H), 3.00 (q, J = 6.6 Hz, 2H), 2.61 (t, J = 7.5 Hz, 2H), 2.36 (ddd, J = 7.8, 6.2, 1.7 Hz, 2H), 2.31 (ddd, J = 8.6, 6.3, 1.7 Hz, 2H), 2.13 (t, J = 7.4 Hz, 2H), 1.92 (p, J = 6.6 Hz, 2H), 1.65 – 1.58 (m, 2H), 1.57 – 1.53 (m, 2H), 1.52 – 1.47 (m, 2H), 1.35 (p, J = 7.0 Hz, 2H), 1.23 (d, J = 4.3 Hz, 6H), 1.10 (t, J = 7.0 Hz, 6H).
**$^{13}$C-NMR** (151 MHz, DMSO-$d_6$): $\delta$ (ppm) = 172.2, 171.4, 171.1, 165.9, 160.6, 159.6, 155.7, 154.0, 151.2, 150.4, 142.9, 141.1, 139.5, 138.6, 135.2, 133.1, 129.0, 128.5, 127.2, 127.2, 126.8, 125.4, 108.7, 105.1, 96.8, 66.5, 62.5, 47.6, 44.0, 42.6, 42.6, 41.8, 41.7, 38.9, 38.5, 35.3, 34.4, 30.9, 30.9, 30.7, 29.1, 28.9, 28.6, 28.5, 28.4, 26.3, 25.3, 12.3.
**HRMS** (ESI+): m/z calc. for $C_{61}H_{74}ClN_{11}NaO_9^+$ $[M+Na]^+$: 1162.525, found: 1162.526.

#### Synthetic route for CHalo-BG

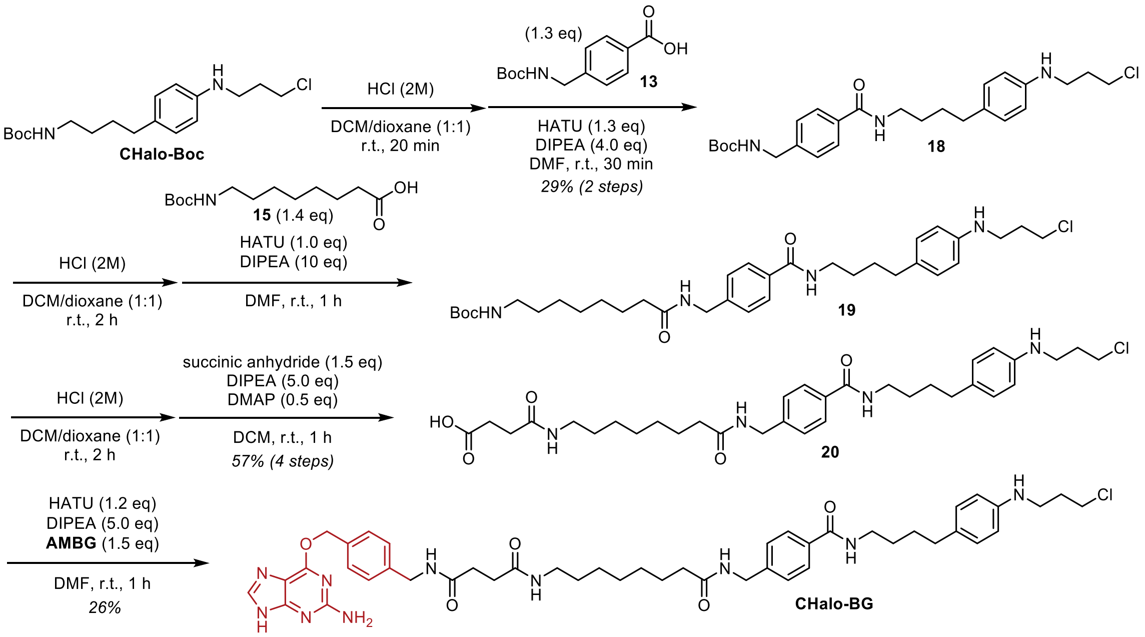

#### Compound 18

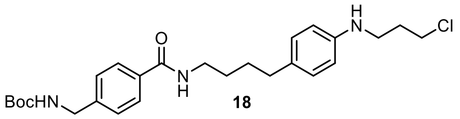

18

**CHalo-Boc** (0.16 g, 0.48 mmol, 1.0 eq) was deprotected according to General Procedure A (0.02 M) and coupled with carboxylate **13** (0.15 g, 0.59 mmol, 1.3 eq, 0.25 M) following General Procedure B (1.0 eq HATU, 30 min reaction time). *N*-methyl piperazine (66 µL, 0.59 mmol, 1.3 eq) was added to react with excess carboxylate. The volatiles were removed *in vacuo*, the residue was diluted with water (15 mL) and extracted with ethyl acetate (3 × 10 mL). The organic layer was dried over sodium sulfate, the desiccant was filtered off and the solvent was removed *in vacuo*. The crude product was purified by flash *rp*-column chromatography (acetonitrile/water; 10→80% MeCN) to give **18** (72 mg, 0.15 mmol, 29% over 2 steps) as a light-yellow solid.

**$^1$H-NMR** (400 MHz, $CDCl_3$): *δ* (ppm) = 7.69 (d, J = 8.1 Hz, 2H), 7.31 (d, J = 8.0 Hz, 2H), 6.99 (d, J = 8.3 Hz, 2H), 6.56 (d, J = 8.4 Hz, 2H), 6.14 (s, 1H), 4.97 (s, 1H), 4.33 (d, J = 5.6 Hz, 2H), 3.65 (t, J = 6.3 Hz, 2H), 3.44 (q, J = 6.4 Hz, 2H), 3.31 (t, J = 6.6 Hz, 2H), 2.54 (t, J = 6.8 Hz, 2H), 2.06 (p, J = 6.5 Hz, 2H), 1.68 – 1.59 (m, 4H), 1.46 (s, 9H).
**$^{13}$C-NMR** (101 MHz, $CDCl_3$): *δ* (ppm) = 167.3, 156.0, 146.1, 142.6, 133.8, 131.3, 129.4, 127.5, 127.3, 113.1, 79.9, 44.4, 42.8, 41.3, 40.1, 34.6, 32.1, 29.3, 29.1, 28.5.
**HRMS** (ESI+): m/z calc. for $C_{26}H_{37}ClN_3O_3^+$ $[M+H]^+$: 474.2518, found: 474.2501

#### Compound 20

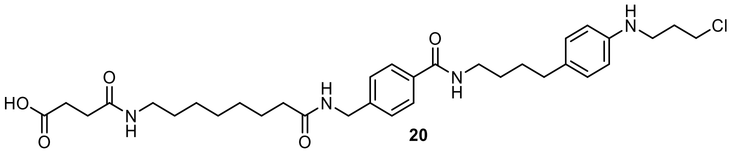

20

**18** (11 mg, 23 µmol, 1.0 eq) was deprotected according to General Procedure A (0.02 M) and coupled with carboxylate **15** (8 mg, 32 µmol, 1.4 eq, 0.03 M) following General Procedure B (1.0 eq HATU, 1 h reaction time). The volatiles were removed *in vacuo* to give crude **19** which was Boc-deprotected according to General Procedure A (0.02 M) and dissolved in anhydrous DCM (2 mL). DIPEA (20 µL, 0.12 mmol, 5.0 eq), DMAP (1.4 mg, 12 µmol, 0.5 eq) and succinic anhydride (2.8 mg, 28 µmol, 1.2 eq) were added and the reaction mixture was stirred at room temperature for 2 h. The volatiles were removed *in vacuo* and the crude product was purified by *rp*-flash column chromatography (acetonitrile/water; 5→50% MeCN) to give **20** (8 mg, 13 µmol, 57% over 4 steps) as a colourless solid.

**$^1$H-NMR** (600 MHz, MeOD/$CDCl_3$ (4:1)): *δ* (ppm) = 7.72 (d, J = 8.3 Hz, 2H), 7.30 (d, J = 8.4 Hz, 2H), 6.96 (d, J = 8.5 Hz, 2H), 6.57 (d, J = 8.5 Hz, 2H), 4.38 (s, 2H), 3.63 (t, J = 6.4 Hz, 2H), 3.36 (t, J = 6.5 Hz, 2H), 3.24 (t, J = 6.7 Hz, 2H), 3.13 (t, J = 7.1 Hz, 2H), 2.57 (t, J = 7.1 Hz, 2H), 2.52 (t, J = 6.9 Hz, 2H), 2.43 (t, J = 7.1 Hz, 2H), 2.21 (t, J = 7.5 Hz, 2H), 2.02 (p, J = 6.6 Hz, 2H), 1.65 – 1.57 (m, 6H), 1.45 (p, J = 7.2 Hz, 2H), 1.30 (s, 6H).

$^{13}$**C-NMR** (151 MHz, MeOD/CDCl$_3$ (4:1)): *δ* (ppm) = 175.8, 175.5, 173.6, 169.2, 146.9, 142.9, 134.0, 131.9, 129.7, 128.0, 127.9, 113.9, 43.3, 43.1, 41.9, 40.5, 39.9, 36.6, 35.1, 32.6, 31.2, 30.1, 29.7, 29.7, 29.5, 29.5, 29.3, 27.1, 26.2.
**HRMS** (ESI+): m/z calc. for $C_{33}H_{48}ClN_4O_5^+$ $[M+H]^+$: 615.3308, found: 615.3308.

**CHalo-BG**

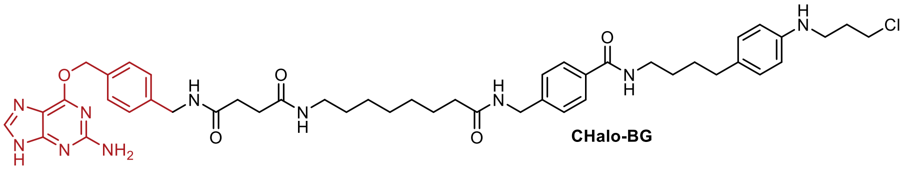

CHalo-BG

6-((4-(aminomethyl)benzyl)oxy)-9*H*-purin-2-amine (**AMBG**, 5 mg, 20 µmol, 1.5 eq) was coupled with carboxylate **20** (8 mg, 13 µmol, 1.0 eq, 0.05 M) following General Procedure B (1.2 eq HATU, 1 h reaction time). The crude product was purified by preparative HPLC (acetonitrile/water, 0.1% formic acid; 5→50% MeCN, 20 min) affording **CHalo-BG** (3.0 mg, 3.4 µmol, 26%) as a colourless solid.

$^{1}$**H-NMR** (600 MHz, DMSO-d$_6$): *δ* (ppm) = 12.42 (s, 1H), 8.39 (t, J = 5.6 Hz, 1H), 8.37 – 8.32 (m, 2H), 7.79 (d, J = 3.3 Hz, 2H), 7.76 (d, J = 8.3 Hz, 2H), 7.43 (d, J = 7.8 Hz, 2H), 7.28 (d, J = 8.3 Hz, 2H), 7.25 (d, J = 8.1 Hz, 2H), 6.89 (d, J = 8.4 Hz, 2H), 6.48 (d, J = 8.5 Hz, 2H), 6.29 (s, 2H), 5.44 (s, 2H), 5.42 (t, J = 5.6 Hz, 1H), 4.28 (d, J = 5.9 Hz, 2H), 4.24 (d, J = 5.9 Hz, 2H), 3.72 (t, J = 6.5 Hz, 2H), 3.24 (q, J = 6.4 Hz, 2H), 3.09 (q, J = 6.3 Hz, 2H), 3.00 (q, J = 6.8 Hz, 2H), 2.43 (t, J = 6.9 Hz, 2H), 2.36 (dd, J = 11.0, 4.6 Hz, 2H), 2.31 (dd, J = 11.1, 4.6 Hz, 2H), 2.13 (t, J = 7.4 Hz, 2H), 1.95 (p, J = 6.6 Hz, 2H), 1.56 – 1.45 (m, 6H), 1.39 – 1.31 (m, 2H), 1.23 (s, 6H).
$^{13}$**C-NMR** (151 MHz, DMSO-d$_6$): *δ* (ppm) = 172.3, 171.4, 171.1, 165.8, 159.9, 159.7, 155.2, 146.8, 142.9, 137.8, 135.2, 133.1, 129.2, 128.8, 128.5, 127.2, 127.2, 126.8, 113.5, 112.1, 66.5, 43.4, 41.8, 41.7, 40.1, 39.1, 38.5, 35.3, 34.0, 31.7, 30.9, 30.9, 29.1, 28.9, 28.8, 28.6, 28.5, 26.3, 25.3.
**HRMS** (ESI+): m/z calc. for $C_{46}H_{59}ClN_{10}NaO_5^+$ $[M+Na]^+$: 889.4251, found: 889.4246.

##### 10.2.5 Light-controlled & reporting protein heterodimeriser Cou-CHalo-SiR-BG

**Synthetic route for Cou-CHalo-SiR-BG**

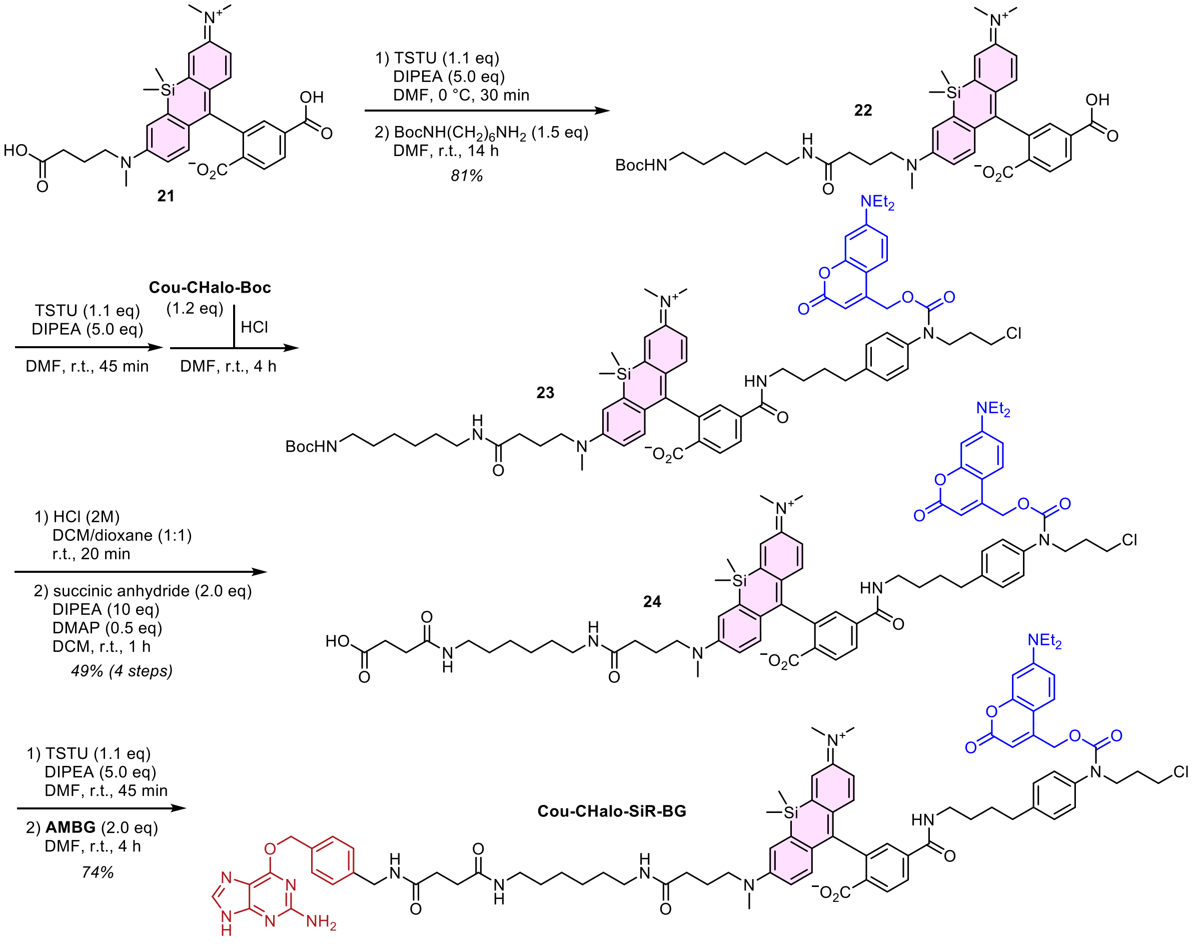

**Compound 22**

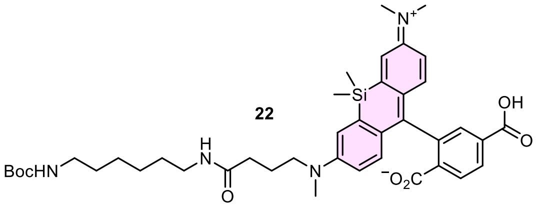

22

**21** was a kind gift from the laboratory of Kai Johnsson.

**21** (10 mg, 19 µmol, 1.0 eq) was dissolved in anhydrous DMF (0.3 mL), DIPEA (16 µL, 92 µmol, 5.0 eq) was added and the mixture was cooled to 0 °C. TSTU (6.1 mg, 20 µmol, 1.1 eq) was dissolved in anhydrous DMF (0.4 mL), added to the solution of **21** and the mixture was stirred at 0 °C for 45 min. Then, Boc-1,6-hexanedimanine (6.0 mg, 28 µmol, 1.5 eq) was added and the mixture was stirred at room temperature for 45 min. The solvent was removed *in vacuo* and the crude product was purified by *rp*-flash column chromatography (acetonitrile/water; 10→80% MeCN) to give **22** (11 mg, 15 µmol, 81%) as a blue solid.

**$^{1}$H-NMR** (600 MHz, $CDCl_3$/MeOD 2:1): *δ* (ppm) = 7.91 (d, J = 8.0 Hz, 1H), 7.67 (d, J = 8.0 Hz, 1H), 7.63 (s, 1H), 6.67 (d, J = 2.7 Hz, 1H), 6.65 (d, J = 2.7 Hz, 1H), 6.40 (d, J = 8.9 Hz, 1H), 6.38 (d, J = 9.0 Hz, 1H), 6.27 – 6.23 (m, 2H), 3.06 – 3.02 (m, 2H), 2.79 (t, J = 7.1 Hz, 2H), 2.68 (t, J = 7.0 Hz, 2H), 2.64 (s, 6H), 2.63 (s, 3H), 1.85 (t, J = 7.3 Hz, 2H), 1.54 (p, J = 7.3 Hz, 2H), 1.11 (dd, J = 14.2, 7.2 Hz, 4H), 1.08 (s, 9H), 0.98 – 0.94 (m, 4H), 0.33 (s, 3H), 0.25 (s, 3H).
**$^{13}$C-NMR** (151 MHz, $CDCl_3$/MeOD 2:1): *δ* (ppm) = 173.4, 170.4, 166.9, 156.7, 154.0, 149.3, 148.1, 136.9, 136.8, 136.0, 130.5, 130.0, 129.8, 129.5, 127.9, 127.8, 125.7, 125.1, 116.4, 116.0, 113.1, 112.7, 78.5, 51.1, 39.7, 39.4, 38.8, 37.3, 32.7, 29.2, 28.6, 27.6, 25.9, 25.8, 22.5, -0.6, -2.4.
**HRMS** (ESI+): m/z calc. for $C_{41}H_{55}N_4O_7Si^+$ $[M+H]^+$: 743.3835, found: 743.3843.

**Compound 24**

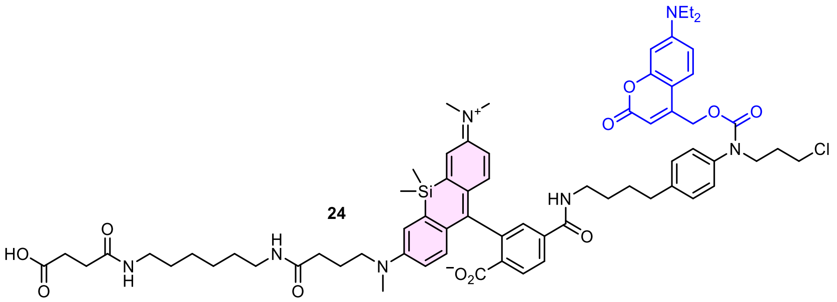

24

**22** (11 mg, 15 µmol, 1.0 eq) was dissolved in anhydrous DMF (0.5 mL) and DIPEA (13 µL, 74 µmol, 5.0 eq) was added. TSTU (5.4 mg, 18 µmol, 1.2 eq) was dissolved in anhydrous DMF (0.4 mL), added to the solution of **22** and the mixture was stirred at room temperature for 30 min. **Cou-CHalo-Boc** (10 mg, 17 µmol, 1.2 eq) was deprotected according to General Procedure A, dissolved in anhydrous DMF (0.4 mL) and added to the reaction mixture, the resulting mixture was stirred at room temperature for 4 h. The solvent was removed *in vacuo* and the crude product was semi-purified by *rp*-flash column chromatography (acetonitrile/water; 20→100% MeCN) to give **23** (12 mg) as green solid. **23** (12 mg, 10 µmol, 1.0 eq) was deprotected according to General Procedure A (0.01 M) and dissolved in anhydrous DCM (1 mL). DIPEA (8.2 µL, 48 µmol, 5.0 eq), DMAP (0.6 mg, 5 µmol, 0.5 eq) and succinic anhydride (2 mg, 19 µmol, 2.0 eq) were added and the reaction mixture was stirred at room temperature for 1 h. The volatiles were removed *in vacuo* and the crude product was purified by *rp*-flash column chromatography (acetonitrile/water; 20→80% MeCN) to give **24** (9 mg, 7 µmol, 49% over 4 steps) as a green solid.

**$^{1}$H-NMR** (600 MHz, $CDCl_3$/MeOD 4:1): *δ* (ppm) = 7.97 (s, 2H), 7.93 (d, J = 7.9 Hz, 1H), 7.74 (s, 1H), 7.19 (d, J = 7.9 Hz, 3H), 7.07 (d, J = 7.0 Hz, 2H), 6.91 (d, J = 2.4 Hz, 1H), 6.89 (d, J = 2.8 Hz, 1H), 6.68 (d, J = 8.9 Hz, 1H), 6.66 (d, J = 9.0 Hz, 1H), 6.55 (d, J = 7.8 Hz, 1H), 6.50 – 6.42 (m, 2H), 6.35 (s, 1H), 5.42 (s, 1H), 5.14 (s, 2H), 3.80 (s, 2H), 3.53 (t, J = 5.8 Hz, 2H), 3.40 – 3.33 (m, 6H), 3.28 (d, J = 7.1 Hz, 2H), 3.13 – 3.07 (m, 4H), 2.88 (s, 9H), 2.68 – 2.63 (m, 2H), 2.54 (t, J = 6.9 Hz, 2H), 2.39 (t, J = 6.9 Hz, 2H), 2.13 (t, J = 7.3 Hz, 2H), 2.02 (p, J = 6.5 Hz, 2H), 1.82 (p, J = 7.3 Hz, 2H), 1.69 (s, 2H), 1.56 (p, J = 7.5 Hz, 2H), 1.44 – 1.37 (m, 4H), 1.23 (dq, J = 11.6, 5.8 Hz, 6H), 1.15 (t, J = 7.1 Hz, 6H), 0.60 (s, 3H), 0.53 (s, 3H).
**$^{13}$C-NMR** (151 MHz, $CDCl_3$/MeOD 4:1): *δ* (ppm) = 175.8, 173.7, 173.2, 171.0, 167.3, 167.2, 163.0, 151.1, 149.7, 148.5, 142.0, 140.5, 138.6, 137.5, 131.2, 130.8, 129.7, 128.8, 128.4, 128.3, 127.5, 125.8, 124.4, 124.2, 116.9, 116.4, 113.5, 113.2, 109.3, 97.6, 93.5, 62.7, 51.7, 48.4, 44.9, 42.2, 40.4, 40.3, 39.3, 39.3, 38.1, 35.2, 33.5, 31.2, 31.1, 30.2, 29.8, 29.2, 29.2, 28.9, 28.8, 26.4, 26.3, 23.1, 12.5, 0.4, -1.6.
**HRMS** (ESI+): m/z calc. for $C_{68}H_{85}ClN_7O_{11}Si^+$ $[M+H]^+$: 1238.5759, found: 1238.5757.

##### Cou-CHalo-SiR-BG

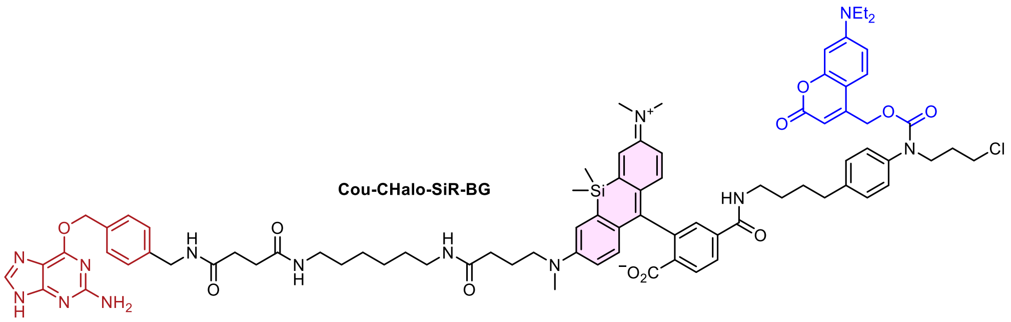

**24** (9 mg, 7 µmol, 1.0 eq) was dissolved in anhydrous DMF (0.3 mL), DIPEA (6.2 µL, 36 µmol, 5.0 eq) and TSTU (50 mM in DMF, 0.17 mL, 8.7 µmol, 1.2 eq) were added and the reaction mixture was stirred at room temperature for 30 min. 6-((4-(aminomethyl)benzyl)oxy)-9*H*-purin-2-amine (**AMBG**, 4 mg, 15 µmol, 2.0 eq) was added and the solution was sonicated to disperse the hardly soluble AMBG. The reaction mixture was stirred for 45 min, the mixture was filtered over celite and the crude product was purified by preparative HPLC (acetonitrile/water, 0.1% formic acid; 25→75% MeCN, 20 min) to give **Cou-CHalo-SiR-BG** (8.0 mg, 5.4 µmol, 74%) as a green solid.

$^{1}$**H-NMR** (600 MHz, $CDCl_3$/MeOD 2:1): $\delta$ (ppm) = 8.28 (t, J = 5.6 Hz, 1H), 8.13 (s, 1H), 8.06 (t, J = 5.7 Hz, 1H), 8.00 (d, J = 7.4 Hz, 1H), 7.95 (d, J = 8.0 Hz, 1H), 7.75 (s, 1H), 7.71 (s, 1H), 7.57 (d, J = 4.9 Hz, 1H), 7.43 – 7.38 (m, 3H), 7.25 – 7.17 (m, 5H), 7.08 (s, 2H), 6.96 – 6.92 (m, 1H), 6.91 (d, J = 2.7 Hz, 1H), 6.68 (d, J = 9.0 Hz, 1H), 6.66 (d, J = 9.1 Hz, 1H), 6.58 (s, 1H), 6.48 (s, 2H), 6.35 (s, 1H), 5.47 (s, 2H), 5.15 (s, 2H), 4.33 (d, J = 4.8 Hz, 2H), 3.80 (s, 2H), 3.54 (t, J = 5.8 Hz, 2H), 3.44 – 3.34 (m, 6H), 3.30 – 3.26 (m, 2H), 3.10 (q, J = 6.5 Hz, 4H), 2.90 (s, 9H), 2.66 (s, 2H), 2.48 (t, J = 6.1 Hz, 2H), 2.46 – 2.41 (m, 2H), 2.15 (t, J = 7.3 Hz, 2H), 2.02 (p, J = 6.5 Hz, 2H), 1.82 (p, J = 7.2 Hz, 2H), 1.74 – 1.66 (m, 2H), 1.58 (p, J = 7.2 Hz, 2H), 1.44 – 1.38 (m, 4H), 1.29 – 1.21 (m, 4H), 1.17 (t, J = 6.7 Hz, 6H), 0.62 (s, 3H), 0.54 (s, 3H).
$^{13}$**C-NMR** (151 MHz, $CDCl_3$/MeOD 2:1): $\delta$ (ppm) = 174.3, 174.2, 173.6, 173.6, 173.6, 171.3, 167.8, 160.3, 156.4, 155.5, 151.7, 150.1, 148.9, 140.8, 138.8, 137.8, 135.8, 131.4, 131.0, 130.0, 129.0, 128.9, 128.8, 128.6, 128.0, 127.8, 126.1, 124.8, 124.5, 117.2, 116.8, 113.8, 113.5, 109.6, 97.9, 68.2, 63.1, 52.0, 48.7, 45.1, 43.5, 42.4, 40.7, 40.4, 39.9, 39.7, 38.3, 35.4, 33.6, 31.9, 31.9, 31.5, 29.5, 29.5, 29.2, 29.1, 26.8, 26.8, 23.4, 12.6, 0.5, -1.5.
**HRMS** (ESI+): m/z calc. for $C_{81}H_{97}ClN_{13}O_{11}Si^{+}$ $[M+H]^{+}$: 1490.6883, found: 1490.6895.

#### 10.2.6 Proof-of-Concept and Design Optimisation

##### Synthetic route for 6-Carboxyfluorescein conjugation precursor (Fluo)

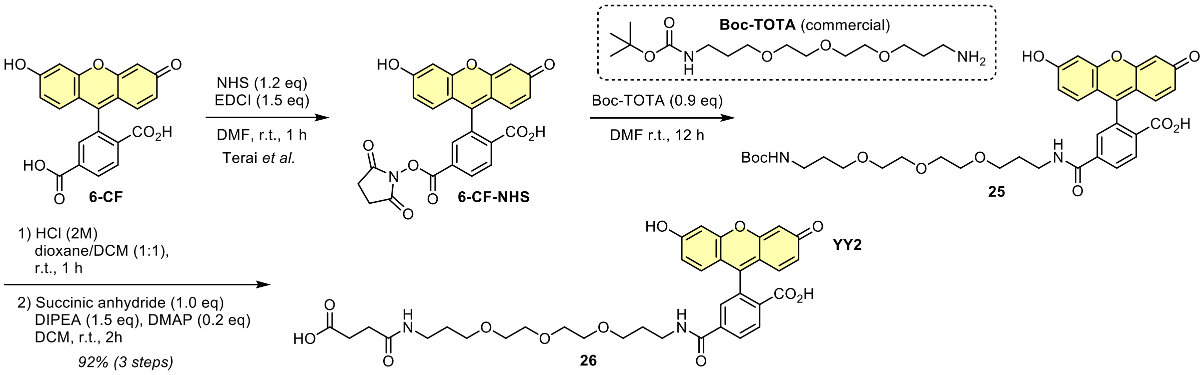

**6-CF-NHS** was prepared according to a previously described procedure (Terai *et al.*[87], first synthetic step for compound 25).

##### Compound 26

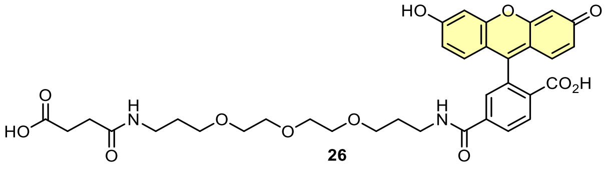

**6-CF-NHS** (51 mg, 0.11 mmol, 1.1 eq) was dissolved in anhydrous DMF (2 mL). **Boc-TOTA** (31 mg, 98 µmol, 1.0 eq) was added and the reaction was stirred at room temperature for 12 h. The volatiles were removed *in vacuo* to afford crude **25** which was used for the next synthetic step without purification.

**25** was deprotected according to General Procedure A (0.03 M) and dissolved in anhydrous DMF (1.5 mL). DIPEA (50 µL, 0.29 mmol, 3.0 eq), 4-(dimethylamino)pyridine (6 mg, 50 µmol, 0.5 eq) and succinic anhydride (10 mg, 98 µmol, 1.0 eq) were added and the reaction mixture was stirred at room temperature for 2 h. The volatiles were removed *in vacuo* and the crude product was purified by *rp*-flash column chromatography (acetonitrile/water, 0.1% formic acid; 5→60% MeCN) affording **26** (61 mg, 90 µmol, 92% over 3 steps) as an orange solid.

**TLC** *Rf* = 0.81 (*rp*, 50% MeCN).
**$^1$H-NMR** (400 MHz, MeOD): *δ* (ppm) = 8.14 (dd, J = 8.0, 1.4 Hz, 1H), 8.08 (dd, J = 8.0, 0.6 Hz, 1H), 7.64 (dd, J = 1.2, 0.7 Hz, 1H), 6.69 (d, J = 2.3 Hz, 2H), 6.61 (d, J = 8.7 Hz, 2H), 6.55 (dd, J = 8.7, 2.4 Hz, 2H), 3.50 (d, J = 7.6 Hz, 8H), 3.44 – 3.37 (m, 6H), 3.19 (t, J = 6.8 Hz, 2H), 2.58 – 2.53 (m, 2H), 2.42 (t, J = 6.8 Hz, 2H), 1.80 (p, J = 6.3 Hz, 2H), 1.67 (p, J = 6.6 Hz, 2H).
**$^{13}$C-NMR** (101 MHz, MeOD): *δ* (ppm) = 176.2, 174.4, 170.6, 167.9, 161.4, 154.7, 154.0, 142.4, 130.4, 130.3, 130.3, 126.1, 123.9, 113.7, 110.9, 103.6, 85.7, 71.3, 71.2, 71.1, 70.9, 70.2, 69.7, 39.1, 37.8, 31.6, 30.3, 30.1.
**HRMS** (ESI+): m/z calc. for $C_{35}H_{39}N_2O_{12}^+$ $[M+H]^+$: 679.2498, found: 679.2484.

##### Synthetic route for CA-Fluo

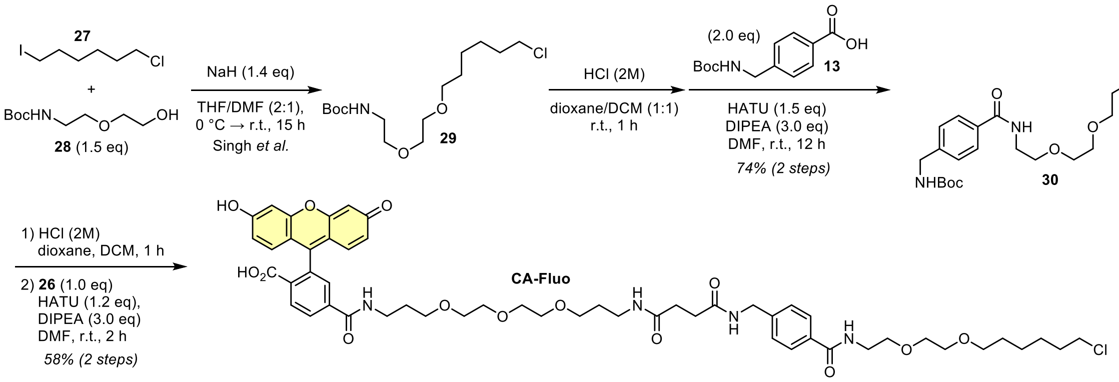

Compound **29** was prepared according to a previously described procedure (Singh *et al.*[88], compound A3).

##### Compound 30

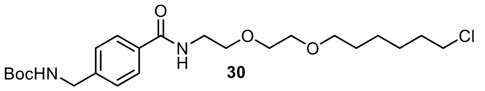

**30** was previously described in a patent by PROMEGA (US9702824[89]). **29** (50 mg, 0.15 mmol, 1.0 eq) was deprotected according to General Procedure A (0.08 M) and coupled with carboxylate **13** (40 mg, 0.15 mmol, 2.0 eq, 0.1 M) following General Procedure B (1.1 eq HATU, 12 h reaction time). The crude product was purified by *np*-flash column chromatography (*iso*-hexanes/ethyl acetate; 20→70% EtOAc) affording **30** (52 mg, 0.11 mmol, 74%) as an off-white solid.

**TLC** *Rf* = 0.70 (*np*, DCM:MeOH 9:1)
**$^1$H-NMR** (400 MHz, $CDCl_3$): *δ* (ppm) = 7.71 (d, J = 8.2 Hz, 2H), 7.29 (d, J = 8.0 Hz, 2H), 6.82 – 6.75 (m, 1H), 5.12 – 5.04 (m, 1H), 4.32 (d, J = 5.8 Hz, 2H), 3.66 – 3.60 (m, 6H), 3.58 – 3.54 (m, 2H), 3.48 (t, J = 6.7 Hz, 2H), 3.43 (t, J = 6.7 Hz, 2H), 1.71 (p, J = 6.7 Hz, 2H), 1.61 – 1.49 (m, 2H), 1.47 – 1.25 (m, 13H)
**$^{13}$C-NMR** (101 MHz, $CDCl_3$): *δ* (ppm) = 167.3, 156.0, 142.8, 133.5, 127.4, 127.4, 79.7, 71.3, 70.3, 70.0, 69.8, 45.1, 44.3, 39.7, 32.5, 29.5, 28.5, 26.7, 25.4
**HRMS** (ESI+): m/z calc. for $C_{23}H_{38}ClN_2O_5^+$ $[M+H]^+$: 457.2464, found: 457.2458

#### CA-Fluo

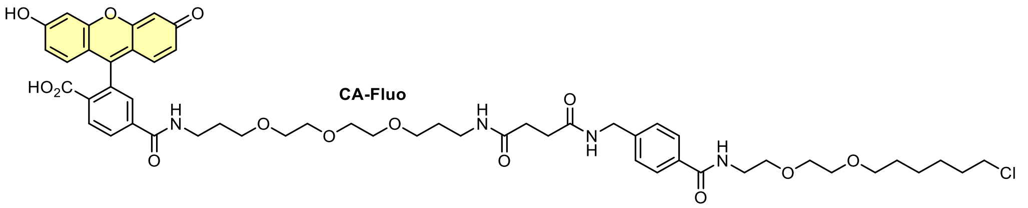

**30** (10 mg, 22 µmol, 1.0 eq) was deprotected according to General Procedure A (0.02 M) and coupled with carboxylate **26** (15 mg, 22 µmol, 1.0 eq, 0.02 M) following General Procedure B (1.1 eq HATU, 12 h reaction time). The crude product was purified by preparative HPLC (acetonitrile/water, 0.1% formic acid; 5→65% MeCN, 20 min) affording **CA-Fluo** (13 mg, 13 µmol, 58% over 2 steps) as an orange solid.

**TLC** *Rf* = 0.49 (*rp*, 50% MeCN); *Rf* = 0.17 (*np*, DCM:MeOH 9:1)
**$^1$H-NMR** (400 MHz, DMSO-d$_6$): $\delta$ (ppm) = 10.16 (s, 2H), 8.67 (t, J = 5.5 Hz, 1H), 8.46 (t, J = 5.6 Hz, 1H), 8.39 (t, J = 6.0 Hz, 1H), 8.16 (dd, J = 8.0, 1.3 Hz, 1H), 8.07 (d, J = 8.1 Hz, 1H), 7.82 – 7.75 (m, 3H), 7.67 (s, 1H), 7.30 (d, J = 8.3 Hz, 2H), 6.69 (d, J = 2.1 Hz, 2H), 6.59 (d, J = 8.7 Hz, 2H), 6.55 (dd, J = 8.7, 2.2 Hz, 2H), 4.29 (d, J = 5.9 Hz, 2H), 3.60 (t, J = 6.6 Hz, 2H), 3.55 – 3.49 (m, 4H), 3.48 – 3.32 (m, 18H), 3.25 (q, J = 6.7 Hz, 2H), 3.05 (q, J = 6.7 Hz, 2H), 2.41 – 2.35 (m, 2H), 2.35 – 2.28 (m, 2H), 1.73 – 1.63 (m, 4H), 1.58 (p, J = 6.6 Hz, 2H), 1.45 (p, J = 6.7 Hz, 2H), 1.39 – 1.21 (m, 4H)
**$^{13}$C-NMR** (101 MHz, DMSO-d$_6$): $\delta$ (ppm) = 171.5, 171.2, 168.1, 166.0, 164.4, 159.8, 152.5, 151.9, 142.9, 140.7, 132.8, 129.4, 129.3, 128.3, 127.2, 126.8, 124.9, 122.3, 112.9, 109.2, 102.3, 83.9, 70.2, 69.7, 69.6, 69.5, 69.5, 69.5, 69.4, 68.9, 68.2, 68.0, 45.4, 41.7, 39.2, 36.9, 35.8, 32.0, 30.8, 30.7, 29.4, 29.1, 29.1, 26.1, 24.9
**HRMS** (ESI+): m/z calc. for $C_{53}H_{66}ClN_4O_{14}^+$ $[M+H]^+$: 1017.426, found: 1017.425

#### Synthetic route for HL2$^O$-Fluo

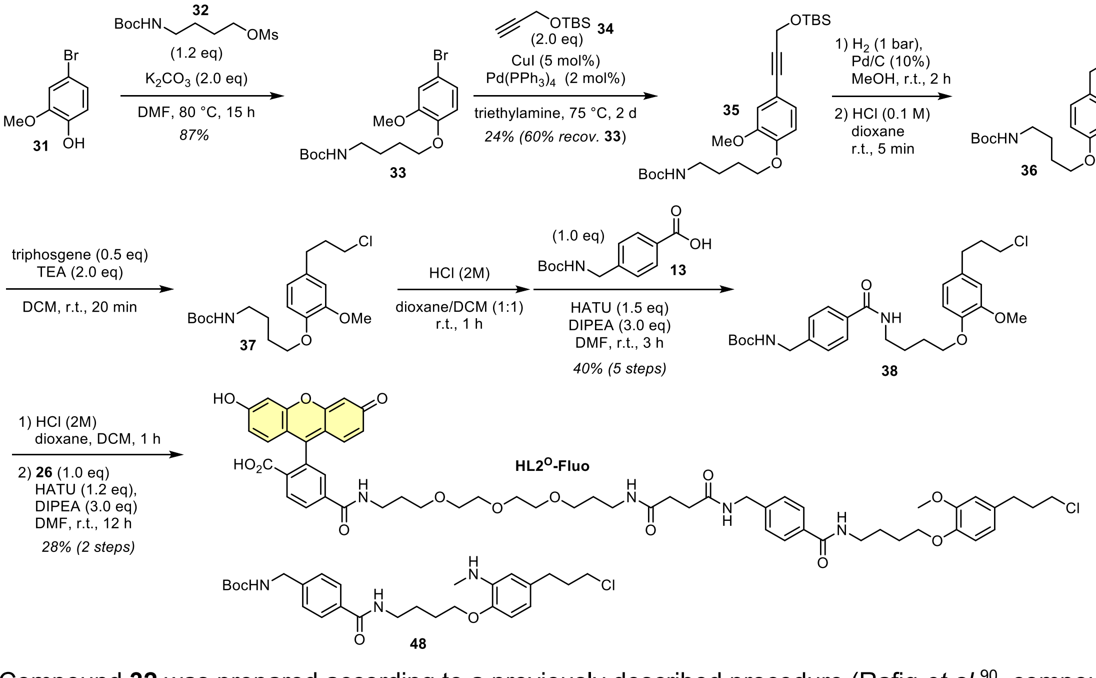

Compound **32** was prepared according to a previously described procedure (Rafiq *et al.*[90], compound 25).

#### Compound 33

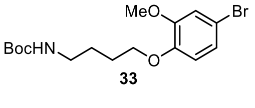

**31** (1.00 g, 4.93 mmol, 1.0 eq) was dissolved in anhydrous DMF (40 mL; *Note*: the mixture becomes very viscous if too concentrated) under nitrogen atmosphere. Potassium carbonate (1.36 g, 9.86 mmol, 2.0 eq) and **32** (1.58 g, 5.92 mmol, 1.2 eq) were added and the reaction mixture heated to 80 °C for 15 h. The solvent was removed *in* vacuo, then a half-saturated solution of sodium bicarbonate (50 mL) was added and the mixture was extracted with ethyl acetate (3 × 50 mL). The organic layer was dried over sodium sulfate, the desiccant was filtered off and the solvent was removed *in vacuo*. The crude product was purified by *np*-flash column chromatography (*iso*-hexanes/ethyl acetate; 5→30% EtOAc) affording **33** (1.60 g, 4.27 mmol, 87%) as a colourless solid.

**TLC** *Rf* = 0.37 (*np*, Hex:EtOAc 4:1)
**$^{1}$H-NMR** (400 MHz, $CDCl_3$): *δ* (ppm) = 7.00 (dd, J = 8.5, 2.3 Hz, 1H), 6.97 (d, J = 2.3 Hz, 1H), 6.72 (d, J = 8.5 Hz, 1H), 4.81 (s, 1H), 3.99 (t, J = 6.3 Hz, 2H), 3.85 (s, 3H), 3.19 (q, J = 6.4 Hz, 2H), 1.91 – 1.81 (m, 2H), 1.67 (p, J = 7.0 Hz, 2H), 1.44 (s, 9H)
**$^{13}$C-NMR** (101 MHz, $CDCl_3$): *δ* (ppm) = 156.2, 150.2, 147.7, 123.5, 115.1, 114.2, 113.0, 79.2, 68.8, 56.2, 40.2, 28.6, 26.9, 26.4
**HRMS** (ESI+): m/z calc. for $C_{16}H_{24}BrNNaO_4^+$ $[M+Na]^+$: 396.0781, found: 396.0785

**Compound 35**

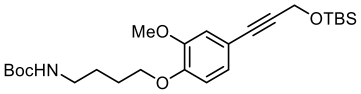

35

**33** (0.55 g, 1.5 mmol, 1.0 eq) was dissolved in triethylamine under nitrogen atmosphere. The solution was degassed by bubbling nitrogen through the mixture for 10 min (ca. half solvent evaporated). Tetrakis(tri-phenylphosphine)palladium(0) (34 mg, 29 µmol, 2 mol%), copper iodide (14 mg, 74 µmol, 5 mol%) and TBS-propargyl alcohol **34** (commercial, 0.60 mL, 2.9 mmol, 2.0 eq) were added and the reaction mixture was heated to 75 °C for 2 d. The volatiles were removed *in vacuo*, the crude product was diluted with water (20 mL) and extracted with ethyl acetate (3 × 20 mL). The combined organic layers were dried over sodium sulfate, the desiccant was filtered off and the solvent was removed *in vacuo*. The crude product was purified by *rp*-flash column chromatography (acetonitrile/water, 0.1% formic acid; 50→100% MeCN) affording **35** (0.17 g, 0.36 mmol, 24%) as a brown oil and recovering the starting material **33** (0.33 g, 0.88 mmol, 60%).

**TLC** *Rf* = 0.74 (*np*, Hex:EtOAc 7:3)
**$^{1}$H-NMR** (400 MHz, $CDCl_3$): *δ* (ppm) = 7.00 (dd, J = 8.2, 1.9 Hz, 1H), 6.93 (d, J = 1.9 Hz, 1H), 6.77 (d, J = 8.3 Hz, 1H), 4.85 (s, 1H), 4.53 (s, 2H), 4.02 (t, J = 6.3 Hz, 2H), 3.84 (s, 3H), 3.19 (q, J = 6.6 Hz, 2H), 1.92 – 1.83 (m, 2H), 1.71 – 1.63 (m, 2H), 1.44 (s, 9H), 0.94 (s, 9H), 0.16 (s, 6H)
**$^{13}$C-NMR** (101 MHz, $CDCl_3$): *δ* (ppm) = 156.2, 148.9, 148.9, 125.0, 115.3, 114.7, 112.4, 86.5, 85.0, 79.1, 68.6, 56.0, 52.5, 40.2, 28.6, 26.9, 26.3, 26.0, 18.5, -4.9
**HRMS** (ESI+): m/z calc. for $C_{25}H_{41}NNaO_5Si^+$ $[M+Na]^+$: 486.2646, found: 486.2642

**Compound 38**

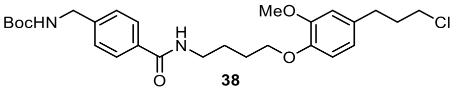

38

**38** was prepared in five synthetic steps without purification of the intermediate products.

**35** (82 mg, 0.18 mmol, 1.0 eq) was dissolved in methanol (3 mL) under nitrogen atmosphere, palladium on charcoal (Pd/C, 21 mg, 18 µmol, 10 mol%) was added and the reaction was stirred under hydrogen atmos-phere (1 atm) for 2 h. The Pd/C was removed by filtration over Celite (washed with methanol (3×) and ethyl acetate (3×)), the filtrate was concentrated under reduce pressure, redissolved in dioxane (0.1 M HCl, 3 mL) and stirred at room temperature for 5 min for selective TBS-deprotection. The crude product was semi-purified by *np*-flash column chromatography (*iso*-hexanes/ethyl acetate; 20→80% EtOAc) to give **36** (40 mg, ca. 64% over 2 steps).

Alcohol **36** was converted to the alkyl chloride **37** adapting a previously described protocol by Ayala *et al.*[91] **36** was dissolved in anhydrous DCM (1 mL) under nitrogen atmosphere and triethylamine (31 µL, 0.23 mmol, 2.0 eq) was added. A solution of triphosgene (17 mg, 57 µmol, 0.5 eq) in anhydrous DCM (0.4 mL) was added dropwise and the reaction mixture was stirred at room temperature for 20 min. The mixture was diluted with a half-saturated solution of sodium bicarbonate (20 mL) and extracted with DCM (3 × 15 mL) to give **37**.

**37** was Boc-deprotected according to General Procedure A (0.11 M) and coupled with carboxylate **13** (29 mg, 0.11 mmol, 1.0 eq, 0.11 M) following General Procedure B (1.1 eq HATU, 3 h reaction time). The crude prod-uct was purified by *np*-flash column chromatography (*iso*-hexanes/ethyl acetate; 20→70% EtOAc) to give **38** (36 mg, 71 µmol, 40% over 5 steps) as a colourless solid.

**TLC** *Rf* = 0.50 (*np*, Hex:EtOAc 1:2)
**$^{1}$H-NMR** (400 MHz, $CDCl_3$): *δ* (ppm) = 7.76 (d, J = 8.1 Hz, 2H), 7.28 (d, J = 8.1 Hz, 2H), 7.10 – 7.05 (m, 1H), 6.79 (d, J = 8.0 Hz, 1H), 6.74 – 6.68 (m, 2H), 5.04 (s, 1H), 4.32 (d, J = 5.7 Hz, 2H), 4.04 (t, J = 5.7 Hz, 2H), 3.70 (s, 3H), 3.56 – 3.48 (m, 4H), 2.71 (t, J = 7.4 Hz, 2H), 2.09 – 2.00 (m, 2H), 1.92 (p, J = 6.2 Hz, 2H), 1.82 (p, J = 6.6 Hz, 2H), 1.45 (s, 9H)
**$^{13}$C-NMR** (101 MHz, $CDCl_3$): *δ* (ppm) = 167.4, 156.0, 149.1, 146.5, 142.5, 133.8, 133.6, 127.4, 127.3, 120.6, 112.6, 112.0, 79.8, 68.7, 55.7, 44.3, 44.3, 39.3, 34.2, 32.4, 28.5, 26.6, 26.1
**HRMS** (ESI+): m/z calc. for $C_{27}H_{38}ClN_2O_5^+$ $[M+H]^+$: 505.2464, found: 505.2458

##### HL2$^{O}$-Fluo

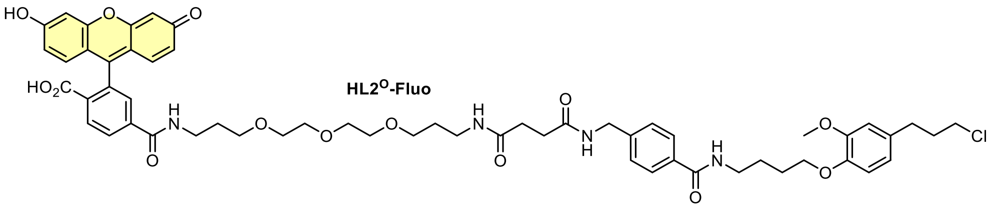

**38** (5.5 mg, 11 µmol, 1.0 eq) was deprotected according to General Procedure A (0.02 M) and coupled with carboxylate **26** (7.6 mg, 11 µmol, 1.0 eq, 0.02 M) following General Procedure B (1.1 eq HATU, 12 h reaction time). The crude product was purified by preparative HPLC (acetonitrile/water, 0.1% formic acid; 10→75% MeCN, 20 min) affording **HL2$^{O}$-Fluo** (3.3 mg, 3.1 µmol, 29% over 2 steps) as an orange solid.

**TLC** *Rf* = 0.41 (*rp*, 50% MeCN); *Rf* = 0.19 (*np*, DCM:MeOH 9:1)
**$^{1}$H-NMR** (800 MHz, DMSO-d$_6$): *δ* (ppm) = 10.18 (s, br, 2H), 8.65 (t, J = 5.6 Hz, 1H), 8.43 (t, J = 5.6 Hz, 1H), 8.38 (t, J = 6.0 Hz, 1H), 8.15 (dd, J = 8.1, 1.1 Hz, 1H), 8.07 (d, J = 8.1 Hz, 1H), 7.81 – 7.76 (m, 3H), 7.66 (s, 1H), 7.30 (d, J = 8.3 Hz, 2H), 6.84 (d, J = 8.2 Hz, 1H), 6.80 (d, J = 1.9 Hz, 1H), 6.68 (d, J = 8.1 Hz, 3H), 6.59 (d, J = 8.7 Hz, 2H), 6.55 (dd, J = 8.7, 2.0 Hz, 2H), 4.29 (d, J = 5.9 Hz, 2H), 3.92 (t, J = 6.4 Hz, 2H), 3.73 (s, 3H), 3.60 (t, J = 6.5 Hz, 2H), 3.46 – 3.44 (m, 4H), 3.44 – 3.40 (m, 4H), 3.38 (t, J = 6.2 Hz, 2H), 3.37 – 3.33 (m, 2H), 3.30 (q, J = 6.8 Hz, 2H), 3.24 (q, J = 6.6 Hz, 2H), 3.05 (q, J = 6.8 Hz, 2H), 2.64 – 2.61 (m, 2H), 2.37 (t, J = 7.2 Hz, 2H), 2.31 (t, J = 7.2 Hz, 2H), 2.00 – 1.96 (m, 2H), 1.73 (dt, J = 14.6, 6.5 Hz, 2H), 1.67 (dp, J = 21.7, 6.7 Hz, 4H), 1.57 (p, J = 6.6 Hz, 2H)
**$^{13}$C-NMR** (201 MHz, DMSO-d$_6$): *δ* (ppm) = 171.5, 171.1, 168.1, 165.9, 164.4, 159.9, 152.4, 151.9, 149.0, 146.4, 142.8, 140.6, 133.2, 133.1, 129.3, 129.3, 128.5, 127.1, 126.8, 125.0, 122.3, 120.1, 113.3, 112.9, 112.4, 109.3, 102.3, 84.4, 69.7, 69.7, 69.5, 69.5, 68.2, 68.0, 68.0, 55.4, 44.8, 41.7, 38.8, 36.9, 35.8, 33.9, 31.8, 30.8, 30.7, 29.3, 29.1, 26.4, 25.9
**HRMS** (ESI+): m/z calc. for $C_{57}H_{66}ClN_4O_{14}^{+}$ $[M+H]^{+}$: 1065.426, found: 1065.424

##### Synthetic route for HL2$^{N}$-Fluo

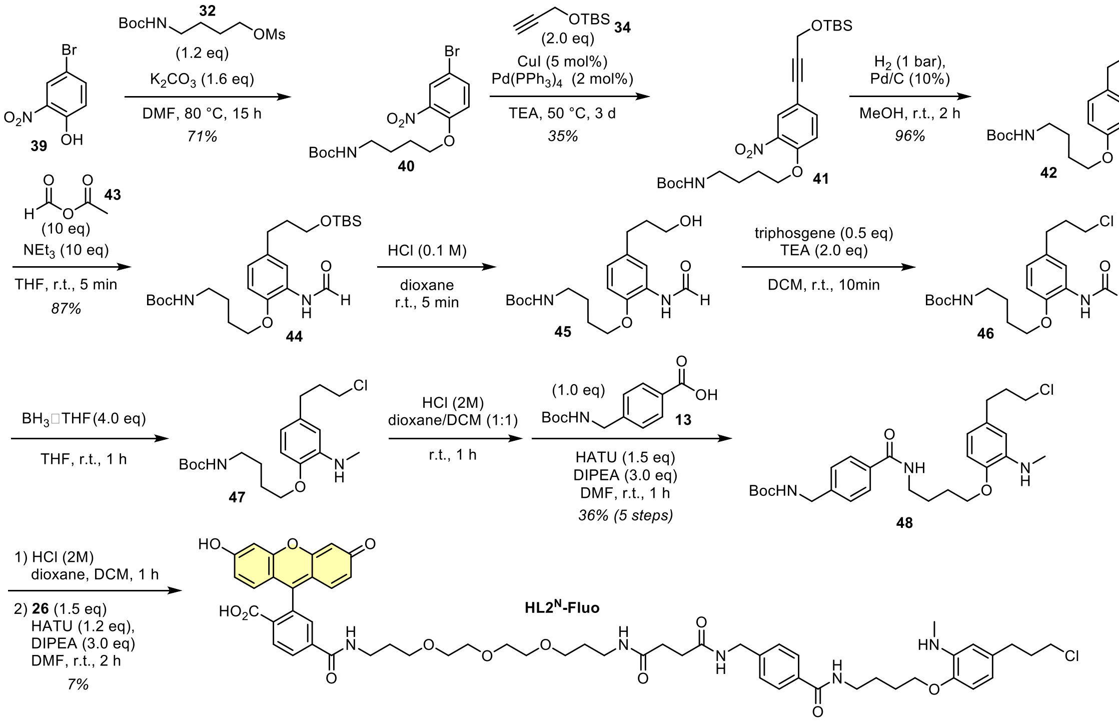

##### Compound 40

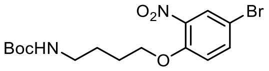

**40**

**39** (1.50 g, 6.88 mmol, 1.0 eq) was dissolved in anhydrous DMF (40 mL; *Note*: the mixture becomes very viscous if too concentrated) under nitrogen atmosphere. Potassium carbonate (1.90 g, 13.8 mmol, 2.0 eq) and mesylate **32** (2.21 g, 8.26 mmol, 1.2 eq) were added and the reaction mixture was heated to 80 °C for 15 h. The solvent was removed *in* vacuo, then a half-saturated solution of sodium bicarbonate (50 mL) was added and the mixture was extracted with ethyl acetate (3 × 50 mL). The organic layer was dried over sodium sulfate, the desiccant was filtered off and the solvent was removed *in vacuo*. The crude product was purified by *np*-flash column chromatography (*iso*-hexanes/ethyl acetate; 5→35% EtOAc) affording **40** (1.26 g, 3.24 mmol, 71%) as a light-yellow solid.

**TLC** *Rf* = 0.18 (*np*, Hex:EtOAc 4:1)
**$^1$H-NMR** (400 MHz, $CDCl_3$): *δ* (ppm) = 7.95 (d, J = 2.5 Hz, 1H), 7.60 (dd, J = 8.9, 2.5 Hz, 1H), 6.96 (d, J = 9.0 Hz, 1H), 4.63 (s, 1H), 4.10 (t, J = 6.1 Hz, 2H), 3.18 (q, J = 6.7 Hz, 2H), 1.90 – 1.81 (m, 2H), 1.73 – 1.64 (m, 2H), 1.43 (s, 9H)
**$^{13}$C-NMR** (101 MHz, $CDCl_3$): *δ* (ppm) = 156.2, 151.6, 140.3, 136.9, 128.4, 116.2, 111.8, 79.3, 69.5, 40.0, 28.5, 26.6, 26.2
**HRMS** (ESI+): m/z calc. for $C_{15}H_{21}BrN_2NaO_5^+$ $[M+Na]^+$: 411.0526, found: 411.0528

**Compound 41**

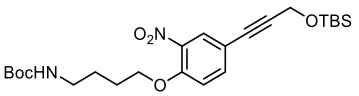

**41**

**40** (1.00 g, 2.57 mmol, 1.0 eq) was dissolved in triethylamine under nitrogen atmosphere. The solution was degassed by bubbling nitrogen through the mixture for 10 min (ca. half solvent evaporated). Tetrakis(triphenylphosphine)palladium(0) (59 mg, 51 µmol, 2 mol%), copper iodide (25 mg, 0.13 mmol, 5 mol%) and TBS-propargyl alcohol **34** (commercial, 1.04 mL, 5.14 mmol, 2.0 eq) were added and the reaction mixture was heated to 50 °C for 3 d. The volatiles were removed *in vacuo*, the crude product was diluted with water (40 mL) and extracted with ethyl acetate (3 × 25 mL). The combined organic layers were dried over sodium sulfate, the desiccant was filtered off and the solvent was removed *in vacuo*. The crude product was purified by *rp*-flash column chromatography (acetonitrile/water, 0.1% formic acid; 50→100% MeCN) affording **41** (0.44 g, 0.91 mmol, 36%) as a brown oil.

**TLC** *Rf* = 0.59 (*np*, Hex:EtOAc 7:3)
**$^1$H-NMR** (400 MHz, $CDCl_3$): *δ* (ppm) = 7.88 (d, J = 2.1 Hz, 1H), 7.55 (dd, J = 8.7, 2.1 Hz, 1H), 6.99 (d, J = 8.8 Hz, 1H), 4.63 (s, 1H), 4.52 (s, 2H), 4.12 (t, J = 6.1 Hz, 2H), 3.19 (q, J = 6.7 Hz, 2H), 1.92 – 1.82 (m, 2H), 1.74 – 1.65 (m, 2H), 1.43 (s, 9H), 0.93 (s, 9H), 0.16 (s, 6H)
**$^{13}$C-NMR** (101 MHz, $CDCl_3$): *δ* (ppm) = 156.2, 152.2, 139.6, 137.2, 128.8, 115.5, 114.4, 88.7, 82.3, 79.4, 69.4, 52.3, 40.1, 28.5, 26.7, 26.2, 26.0, 18.5, -4.9
**HRMS** (ESI+): m/z calc. for $C_{24}H_{38}N_2NaO_6Si^+$ $[M+Na]^+$: 501.2391, found: 501.2387

**Compound 42**

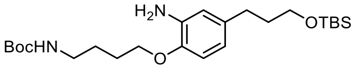

**42**

**41** (0.40 g, 0.84 mmol, 1.0 eq) was dissolved in methanol (3 mL) under nitrogen atmosphere, palladium on charcoal (Pd/C, 50 mg, 42 µmol, 10 mol%) was added and the reaction was stirred under hydrogen atmosphere (1 atm) for 2.5 h. The Pd/C was removed by filtration over Celite (washed with methanol (3×) and ethyl acetate (3×)), the volatiles were removed *in vacuo* giving **42** (0.36 g, 0.80 mmol, 96%) as a colourless oil.

**TLC** *Rf* = 0..52 (*np*, Hex:EtOAc 2:1)
**$^1$H-NMR** (400 MHz, $CDCl_3$): *δ* (ppm) = 6.68 (d, J = 8.2 Hz, 1H), 6.58 (s, 1H), 6.53 (d, J = 8.1 Hz, 1H), 4.58 (s, 1H), 3.98 (t, J = 6.2 Hz, 2H), 3.61 (t, J = 6.4 Hz, 2H), 3.19 (q, J = 6.7 Hz, 2H), 2.56 – 2.50 (m, 2H), 1.87 – 1.73 (m, 4H), 1.72 – 1.62 (m, 2H), 1.44 (s, 9H), 0.90 (s, 9H), 0.05 (s, 6H)
**$^{13}$C-NMR** (101 MHz, $CDCl_3$): *δ* (ppm) = 145.0, 135.2, 118.5, 115.8, 111.6, 79.3, 68.0, 62.6, 40.5, 34.8, 31.6, 28.6, 27.1, 26.8, 26.1, 18.5, -5.1
**HRMS** (ESI+): m/z calc. for $C_{24}H_{45}N_2O_4Si^+$ $[M+H]^+$: 453.3143, found: 453.3137

**Compound 44**

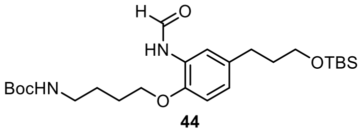

**44**

Adaptation from previously described procedure.[92] First, mixed anhydride **43** was prepared: Formic acid (0.17 mL, 4.5 mmol, 1.5 eq) and acetic anhydride (0.28 mL, 3.0 mmol, 1.0 eq) were heated to 50 °C for 2.5 h to form the mixed anhydride.

In a separate flask, **42** (90 mg, 0.20 mmol, 1.0 eq) was dissolved in anhydrous THF (3 mL), triethylamine (0.28 mL, 2.0 mmol, 10 eq) and **43** (0.16 mL, 2.0 mmol, 10 eq) were added and the reaction was stirred at room temperature for 1 h. The volatiles were removed *in vacuo*, the crude product was diluted with a half-saturated solution of sodium bicarbonate (20 mL) and the mixture was extracted with ethyl acetate (3 × 15 mL). The organic layer was dried over sodium sulfate, the desiccant was filtered off and the solvent was removed *in vacuo*. The crude product was purified by *np*-flash column chromatography (*iso*-hexanes/ethyl acetate; 10→50% EtOAc) affording **44** (83 mg, 0.17 mmol, 87%) as a light-red oil.

*Note: the product might partially TBS-deprotected during silica column chromatography.*

**TLC** *Rf* = 0.32 (*np*, Hex:EtOAc 2:1)
**$^1$H-NMR** (400 MHz, $CDCl_3$), *rotamers observed*: *δ* (ppm) = 8.73 (d, J = 11.7 Hz, 0.2H), 8.50 (d, J = 1.4 Hz, 0.6H), 8.38 (s, 0.5H), 8.25 (d, J = 1.7 Hz, 0.6H), 8.16 (s, 0.1H), 7.68 (d, J = 10.8 Hz, 0.2H), 7.12 – 7.01 (m, 0.4H), 6.95 – 6.89 (m, 0.3H), 6.86 (dd, J = 8.2, 1.8 Hz, 0.7H), 6.81 (d, J = 8.4 Hz, 0.3H), 6.77 (d, J = 8.3 Hz, 0.7H), 4.59 (s, 1H), 4.01 (dt, J = 20.9, 6.3 Hz, 2H), 3.62 (t, J = 6.4 Hz, 2H), 3.31 – 3.13 (m, 2H), 2.64 – 2.58 (m, 2H), 1.89 – 1.75 (m, 4H), 1.67 (dq, J = 12.8, 6.7 Hz, 2H), 1.44 (s, 9H), 0.90 (s, 9H), 0.05 (s, 6H)
**HRMS** (ESI−): m/z calc. for $C_{25}H_{43}N_2O_5Si^-$ $[M-H]^-$: 479.2947, found: 479.2945

**Compound 48**

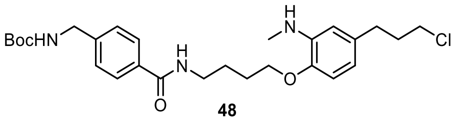

**48**

**48** was prepared in five synthetic steps without purification of the intermediate products.

**44** (53 mg, 0.11 mmol, 1.0 eq) dissolved in dioxane with HCl (0.1 M HCl, 2 mL) and stirred at room temperature for 5 min for selective TBS-deprotection. The volatiles were removed *in vacuo* and the crude product was diluted with a half-saturated solution of sodium bicarbonate (20 mL) and extracted with ethyl acetate (3 × 15 mL) affording alcohol **45** which was used without further purification.

Alcohol **45** was converted to the alkyl chloride **46** adapting a previously described protocol by Ayala *et al.*[91] **45** was dissolved in anhydrous DCM (1 mL) under nitrogen atmosphere and triethylamine (30 µL, 0.22 mmol, 2.0 eq) was added. A solution of triphosgene (16 mg, 55 µmol, 0.5 eq) in anhydrous DCM (0.4 mL) was added dropwise and the reaction mixture was stirred at room temperature for 20 min. The mixture was diluted with a half-saturated solution of sodium bicarbonate (20 mL) and extracted with DCM (3 × 15 mL). The organic layer was dried over sodium sulfate and the solvent was removed *in vacuo* to give **46** which was used without further purification.

**46** was dissolved in anhydrous THF (3 mL) under nitrogen atmosphere and cooled to 0 °C. Borane-tetrahydrofuane complex (1 M in THF, 0.44 mL, 0.44 mmol, 4.0 eq) was added and the reaction mixture was stirred at room temperature for 1 h. Citric acid (0.12 g, 0.55 mmol, 5.0 eq) was dissolved in water (2 mL) and added to the reaction mixture dropwise to quench the excess of the borane. The resulting mixture was diluted with water (20 mL), a saturated solution of bicarbonate (5 mL) and extracted with ethyl acetate (3 × 15 mL). The organic layer was dried over sodium sulfate and the solvent was removed *in vacuo* to give **47** which was used without further purification.

**47** was deprotected according to General Procedure A (0.09 M) and coupled with carboxylate **13** (28 mg, 0.11 mmol, 1.0 eq, 0.09 M) following General Procedure B (1.1 eq HATU, 2 h reaction time). The crude product was purified by flash column chromatography (*iso*-hexanes/ethyl acetate; 20→70% EtOAc) to give **48** (16 mg, 32 µmol, 36% over 5 steps) as a colourless solid.

**TLC** *Rf* = 0.54 (*np*, Hex:EtOAc 1:2); *Rf* = 0.76 (*np*, DCM:MeOH 9:1)
**$^1$H-NMR** (400 MHz, $CDCl_3$): *δ* (ppm) = 7.69 (d, J = 8.2 Hz, 2H), 7.30 (d, J = 8.2 Hz, 2H), 6.66 (d, J = 8.0 Hz, 1H), 6.46 (dd, J = 8.0, 2.0 Hz, 1H), 6.43 (d, J = 2.0 Hz, 1H), 6.35 – 6.27 (m, 1H), 4.99 – 4.90 (m, 1H), 4.34 (d, J = 5.8 Hz, 2H), 4.01 (t, J = 5.9 Hz, 2H), 3.57 – 3.49 (m, 4H), 2.85 (s, 3H), 2.69 (t, J = 7.3 Hz, 2H), 2.10 – 2.02 (m, 2H), 1.89 (dq, J = 10.7, 5.9 Hz, 2H), 1.81 (dt, J = 13.5, 6.2 Hz, 2H), 1.46 (s, 9H)
**$^{13}$C-NMR** (101 MHz, $CDCl_3$): *δ* (ppm) = 167.4, 156.0, 144.6, 142.7, 139.3, 133.8, 133.7, 127.5, 127.3, 116.3, 110.3, 110.1, 79.9, 67.9, 44.6, 44.4, 39.9, 34.5, 32.7, 30.6, 28.5, 26.9, 26.8
**HRMS** (ESI+): m/z calc. for $C_{27}H_{39}ClN_3O_4^+$ $[M+H]^+$: 504.2624, found: 504.2617

##### $HL2^N$-Fluo

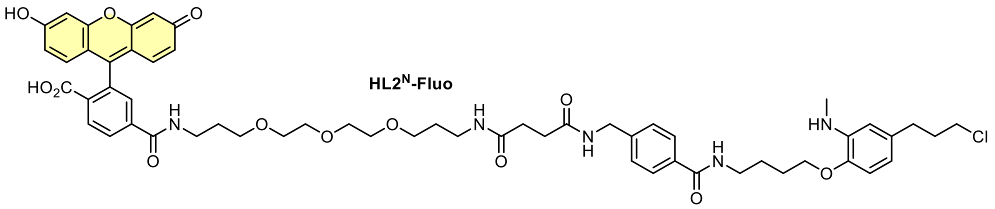

**48** (8.0 mg, 16 µmol, 1.0 eq) was deprotected according to General Procedure A (0.02 M) and coupled with carboxylate **26** (16 mg, 24 µmol, 1.5 eq, 0.04 M) following General Procedure B (1.1 eq HATU, 3 h reaction time). The crude product was purified by preparative HPLC (acetonitrile/water, 0.1% formic acid; 10→50% MeCN, 20 min) affording **$HL2^N$-Fluo** (1.2 mg, 1.2 µmol, 7% over 2 steps) as an orange solid.

**$^1$H-NMR** (400 MHz, DMSO-d$_6$): $\delta$ (ppm) = 8.64 (t, $J$ = 5.4 Hz, 1H), 8.49 – 8.39 (m, 2H), 8.13 (d, $J$ = 8.9 Hz, 1H), 8.07 (d, $J$ = 8.1 Hz, 1H), 7.81 (dd, $J$ = 16.3, 6.9 Hz, 3H), 7.65 (s, 1H), 7.30 (d, $J$ = 8.3 Hz, 2H), 6.67 (d, $J$ = 8.0 Hz, 1H), 6.61 (d, $J$ = 8.8 Hz, 3H), 6.51 (d, $J$ = 8.0 Hz, 2H), 6.33 (dd, $J$ = 7.9, 2.0 Hz, 1H), 6.31 (d, $J$ = 1.9 Hz, 1H), 4.89 – 4.84 (m, 1H), 4.29 (d, $J$ = 5.9 Hz, 2H), 3.93 (t, $J$ = 6.0 Hz, 2H), 3.60 (t, $J$ = 6.5 Hz, 2H), 3.51 – 3.38 (m, 16H), 3.28 – 3.22 (m, 4H), 3.05 (q, $J$ = 6.8 Hz, 2H), 2.70 (d, $J$ = 4.8 Hz, 2H), 2.60 – 2.55 (m, 2H), 2.42 – 2.36 (m, 2H), 2.33 (d, $J$ = 5.8 Hz, 3H), 1.97 (dt, $J$ = 13.7, 6.6 Hz, 2H), 1.73 (dq, $J$ = 19.8, 6.6 Hz, 6H), 1.58 (p, $J$ = 6.6 Hz, 2H)
**HRMS** (ESI+): m/z calc. for $C_{57}H_{67}ClN_5O_{13}^+$ $[M+H]^+$: 1064.442, found: 1064.441.

##### Synthetic route for Cou-$HL2^N$-Fluo

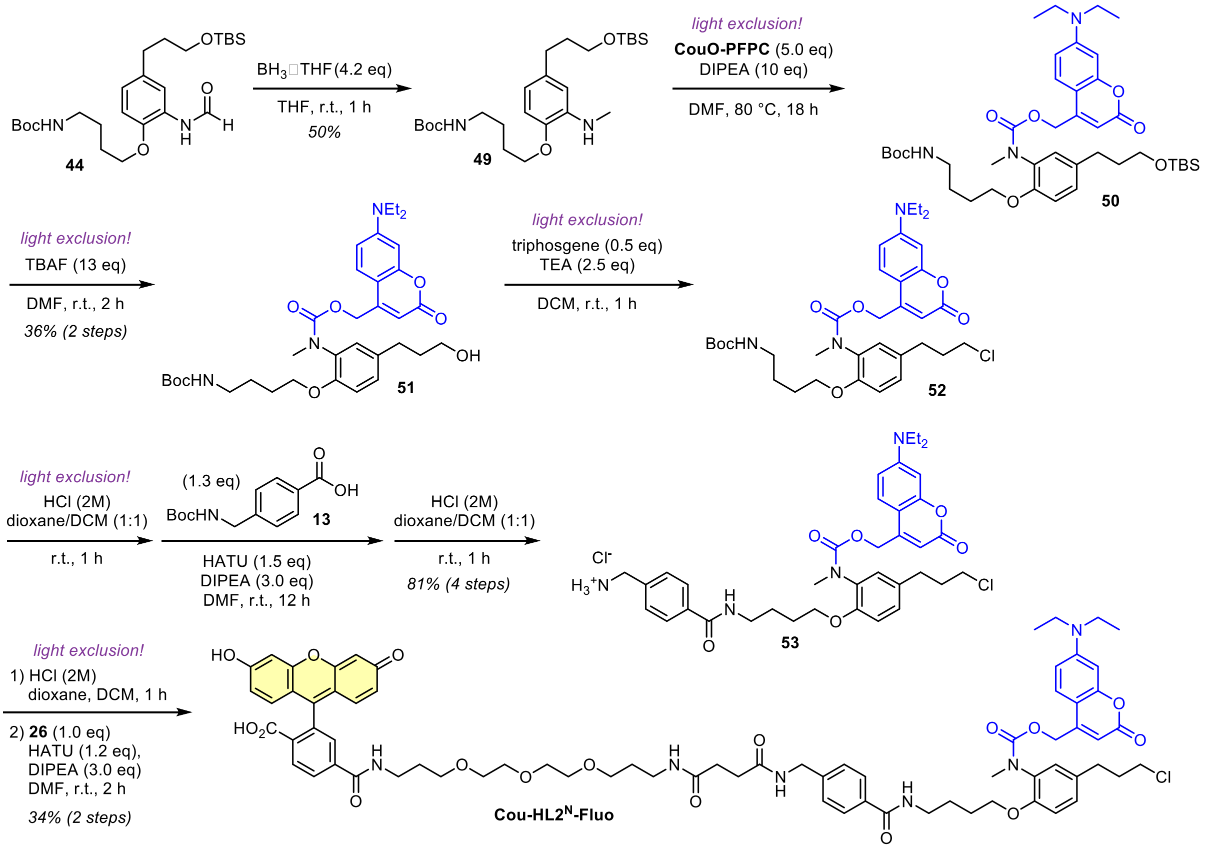

##### Compound 49

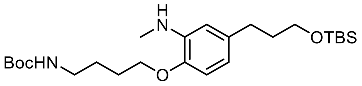

49

**44** (40 mg, 83 µmol, 1.0 eq) was dissolved in anhydrous THF (1.5 mL) under nitrogen atmosphere and cooled to 0 °C. Borane-tetrahydrofuane complex (1 M in THF, 0.35 mL, 0.35 mmol, 4.2 eq) was added and the reaction mixture was stirred at room temperature for 1 h. Citric acid (87 g, 0.42 mmol, 5.0 eq) was dissolved in water (1 mL) and added to the reaction mixture dropwise to quench the excess of the borane. The resulting mixture was diluted with water (20 mL), a saturated solution of bicarbonate (5 mL) and extracted with ethyl acetate (3 × 15 mL). The organic layer was dried over sodium sulfate, the desiccant was filtered off and the solvent was removed *in vacuo*. The crude product was purified by flash column chromatography (*iso*-hexanes/ethyl acetate; 5→30% EtOAc) to give **49** (19 mg, 41 µmol, 50%) as a colourless oil.

**TLC** *Rf* = 0.47 (*np*, Hex:EtOAc 3:1)
**$^1$H-NMR** (400 MHz, $CDCl_3$): *δ* (ppm) = 6.64 (d, J = 7.8 Hz, 1H), 6.48 – 6.43 (m, 2H), 4.58 (s, 1H), 3.97 (t, J = 6.2 Hz, 2H), 3.64 (t, J = 6.4 Hz, 2H), 3.19 (q, J = 6.7 Hz, 2H), 2.86 (s, 3H), 2.62 – 2.54 (m, 2H), 1.86 – 1.77 (m, 4H), 1.70 – 1.61 (m, 2H), 1.44 (s, 9H), 0.91 (s, 9H), 0.06 (s, 6H)
**$^{13}$C-NMR** (101 MHz, $CDCl_3$): *δ* (ppm) = 156.1, 144.5, 139.3, 135.4, 116.0, 110.2, 110.1, 79.3, 67.9, 62.8, 40.5, 34.9, 32.0, 30.6, 28.6, 27.1, 26.8, 26.1, 18.5, -5.1
**HRMS** (ESI+): m/z calc. for $C_{25}H_{47}N_2O_4Si^+$ $[M+H]^+$: 467.3300, found: 467.3296

**Compound 51**

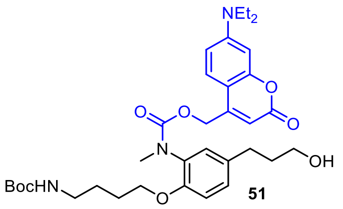

51

*Note: the reaction must be performed under light exclusion.*

**49** (25 mg, 54 µmol, 1.0 eq) was dissolved in anhydrous DMF (1 mL) and transformed to **50** following General Procedure D (0.05 M) which was TBS-deprotected by addition of tetrabutylammonium fluoride (1 M in THF, 0.70 mL, 0.70 mmol, 13 eq) into the reaction mixture. The mixture was diluted with water (20 mL) and extracted with ethyl acetate (4 × 15 mL). The organic layer was washed with brine (30 mL), dried over sodium sulfate and the volatiles were removed *in vacuo*. The crude product was purified by reversed-phase flash column chromatography (acetonitrile/water, 0.1% formic acid; 20→90% MeCN) affording **51** (12 mg, 19 µmol, 36% over 2 steps) as a yellow solid.

**TLC** *Rf* = 0.33 (*np*, Hex:EtOAc 1:2); *Rf* = 0.65 (*np*, DCM:MeOH 9:1)
**HRMS** (ESI+): m/z calc. for $C_{34}H_{48}N_3O_8^+$ $[M+H]^+$: 626.3436, found: 626.3438.

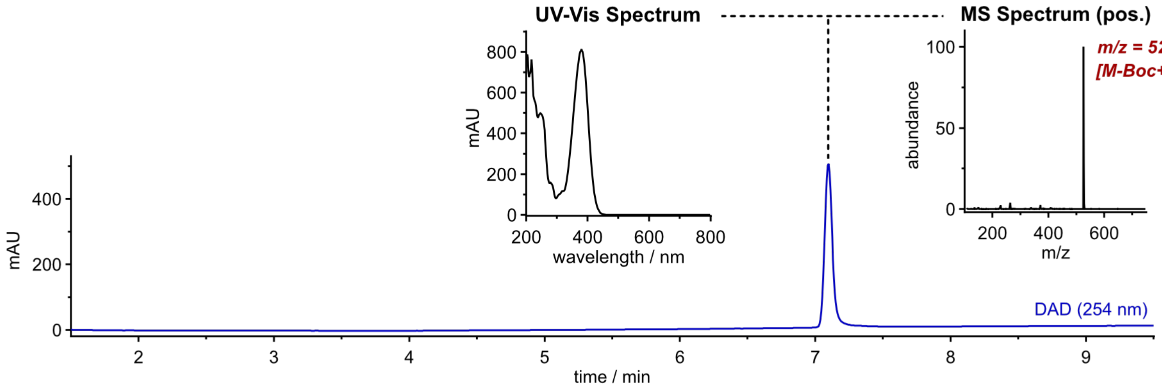

**Compound 53**

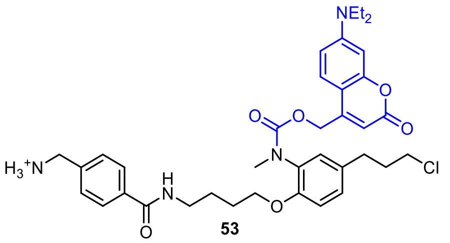

53

*Note: the reaction must be performed under light exclusion.*

Alcohol **51** was converted to the alkyl chloride **52** adapting a previously described protocol by Ayala *et al.*[91]
**51** (12 mg, 19 µmol, 1.0 eq) was dissolved in anhydrous DCM (2 mL) under nitrogen atmosphere and triethylamine (6.7 µL, 48 µmol, 2.5 eq) was added. A solution of triphosgene (2.9 mg, 9.6 µmol, 0.5 eq) in anhydrous DCM (0.1 mL) was added dropwise and the reaction mixture was stirred at room temperature for 20 min. The mixture was diluted with a half-saturated solution of sodium bicarbonate (20 mL) and extracted with DCM (3 × 15 mL). The organic layer was dried over sodium sulfate and the solvent was removed *in vacuo* to give **52** which was used without further purification.

**52** was deprotected according to General Procedure A (0.02 M) and coupled with carboxylate **13** (6.4 mg, 25 µmol, 1.3 eq, 0.02 M) following General Procedure B (1.1 eq HATU, 2 h reaction time). The crude product was purified by *np*-flash column chromatography (*iso*-hexanes/ethyl acetate; 30→80% EtOAcwhich was directly converted to **53** following General Procedure A (0.02 M) affording the product (11 mg, 15 µmol, 81% over 4 steps) as a colourless solid.

**LRMS** (ESI+): m/z calc. for $C_{37}H_{46}ClN_4O_6^+$ $[M+H]^+$: 677.3, found: 677.3.

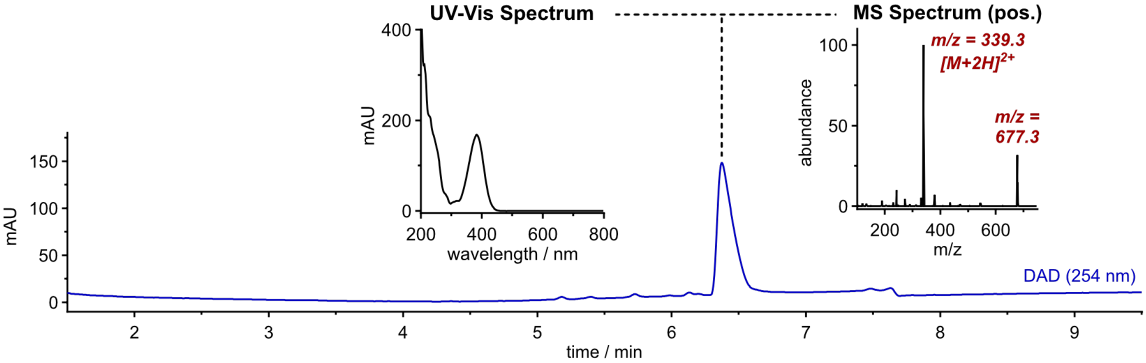

#### HL2$^N$-Fluo

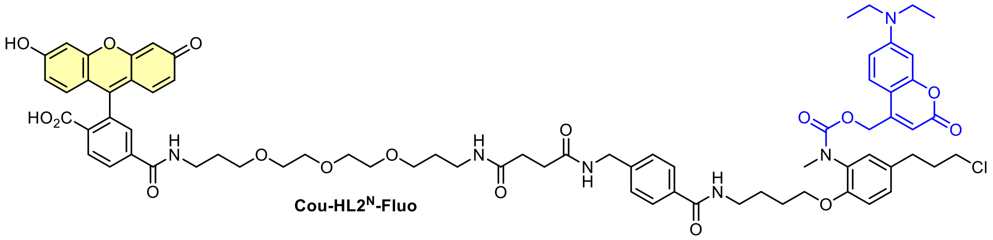

*Note: the reaction must be performed under light exclusion.*

**53** (11 mg, 15 µmol, 1.0 eq) coupled with carboxylate **2** (11 mg, 15 µmol, 1.0 eq, 0.03 M) following General Procedure B (1.1 eq HATU, 12 h reaction time). The crude product was purified by preparative HPLC (acetonitrile/water, 0.1% formic acid; 20→85% MeCN, 20 min) affording **Cou-HL2$^N$-Fluo** (7.1 mg, 5.3 µmol, 34% over 2 steps) as an orange solid.

**TLC** *Rf* = 0.19 (*rp*, 50% MeCN); *Rf* = 0.27 (*np*, DCM:MeOH 9:1)

**$^1$H-NMR** (800 MHz, DMSO-d$_6$): *δ* (ppm) = 10.21 (s, 2H), 8.66 (t, J = 5.5 Hz, 1H), 8.42 (t, J = 5.3 Hz, 1H), 8.38 (t, J = 5.7 Hz, 1H), 8.16 (d, J = 8.1 Hz, 1H), 8.07 (d, J = 8.1 Hz, 1H), 7.80 (t, J = 5.4 Hz, 1H), 7.76 (d, J = 8.1 Hz, 2H), 7.67 (s, 1H), 7.30 (dd, J = 12.9, 8.6 Hz, 3H), 7.15 – 7.09 (m, 2H), 7.03 (d, J = 8.3 Hz, 1H), 6.69 (s, 2H), 6.62 – 6.57 (m, 3H), 6.56 (d, J = 7.1 Hz, 2H), 6.50 (s, 1H), 5.67 (s, 1H), 5.34 – 5.13 (m, 2H), 4.29 (d, J = 5.6 Hz, 2H), 4.01 (t, J = 6.2 Hz, 2H), 3.59 (t, J = 6.4 Hz, 2H), 3.49 – 3.45 (m, 4H), 3.45 – 3.34 (m, 12H), 3.30 – 3.27 (m, 2H), 3.27 – 3.23 (m, 2H), 3.12 (s, 3H), 3.06 (q, J = 6.6 Hz, 2H), 2.67 – 2.62 (m, 2H), 2.38 (t, J = 7.2 Hz, 2H), 2.32 (t, J = 7.2 Hz, 2H), 1.98 (p, J = 6.6 Hz, 2H), 1.74 (s, 2H), 1.69 (p, J = 6.6 Hz, 2H), 1.66 – 1.61 (m, 2H), 1.58 (p, J = 6.6 Hz, 2H), 1.11 (t, J = 6.9 Hz, 6H)

**$^{13}$C-NMR** (201 MHz, DMSO-d$_6$): *δ* (ppm) = 171.5, 171.1, 168.1, 165.9, 164.4, 160.6, 159.9, 155.7, 154.5, 152.5, 152.2, 151.9, 151.4, 150.3, 142.7, 140.6, 133.1, 132.9, 130.6, 129.3, 129.3, 128.6, 128.6, 127.1, 126.8, 125.2, 125.0, 122.3, 113.0, 113.0, 112.9, 109.3, 108.5, 105.1, 104.1, 102.3, 96.8, 82.8, 69.7, 69.7, 69.5, 69.5, 68.2, 68.0, 67.6, 62.1, 44.7, 44.0, 41.7, 38.8, 37.3, 36.9, 35.8, 33.6, 31.1, 30.8, 30.7, 29.3, 29.1, 26.2, 25.8, 12.3

**HRMS** (ESI+): m/z calc. for $C_{72}H_{82}ClN_6O_{17}^+$ $[M+H]^+$: 1337.542, found: 1337.539.

#### Synthetic route for HL3-Fluo

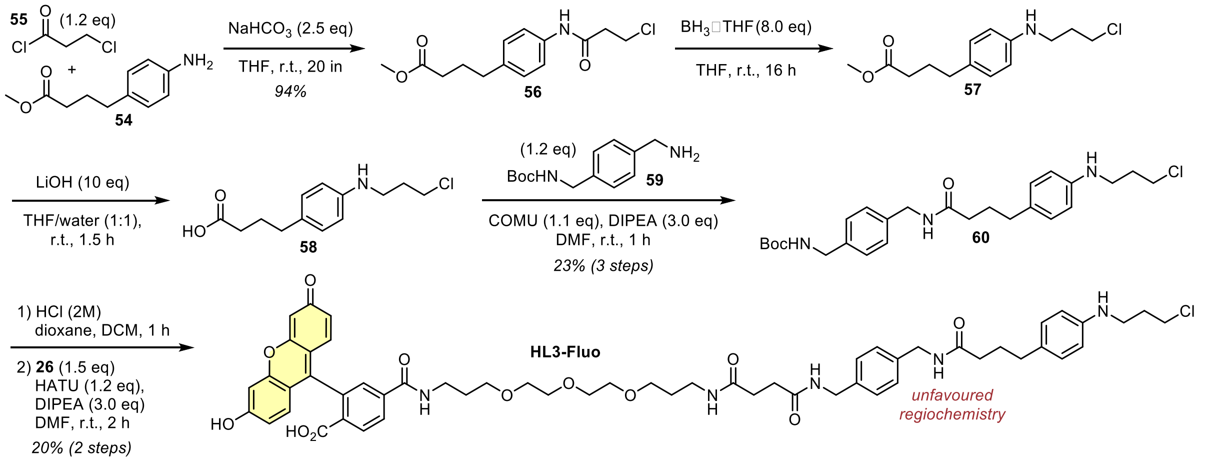

#### Compound 56

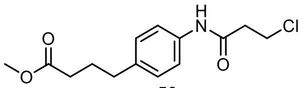

56

Methyl 4-(4-aminophenyl)butanoate (**54**, 50 mg, 0.26 mmol, 1.0 eq) was dissolved in anhydrous THF. Sodium bicarbonate (54 mg, 0.65 mmol, 2.5 eq) and 3-chloropropionyl chloride (**55**, 30 µL, 0.31 mmol, 1.2 eq) were added and the reaction mixture was stirred at room temperature for 20 min. The mixture was diluted with a half-saturated solution of sodium bicarbonate (20 mL) and extracted with ethyl acetate (3 × 15 mL). The organic layer was washed with brine (30 mL), dried over sodium sulfate and the solvent was removed *in vacuo* affording **56** (69 mg, 0.24 mmol, 94%) as a colourless solid.

**TLC** *Rf* = 0.25 (*np*, Hex:EtOAc 3:1)
**$^1$H-NMR** (400 MHz, $CDCl_3$): *δ* (ppm) = 7.43 (d, J = 8.4 Hz, 2H), 7.28 (s, 1H), 7.14 (d, J = 8.4 Hz, 2H), 3.88 (t, J = 6.4 Hz, 2H), 3.66 (s, 3H), 2.80 (t, J = 6.4 Hz, 2H), 2.62 (t, J = 7.6 Hz, 2H), 2.32 (t, J = 7.5 Hz, 2H), 1.93 (p, J = 7.5 Hz, 2H).
**$^{13}$C-NMR** (101 MHz, $CDCl_3$): *δ* (ppm) = 174.1, 167.7, 138.0, 135.5, 129.2, 120.3, 51.7, 40.7, 40.1, 34.6, 33.4, 26.6.
**HRMS** (ESI+): m/z calc. for $C_{14}H_{19}ClNO_3^+$ $[M+H]^+$: 284.1048, found: 284.1051

#### Compound 60

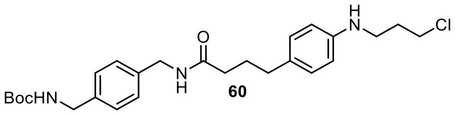

**56** (50 mg, 0.18 µmol, 1.0 eq) was dissolved in anhydrous THF (1 mL) under nitrogen atmosphere and cooled to 0 °C. Borane-tetrahydrofuane complex (1 M in THF, 1.4 mL, 1.4 mmol, 8.0 eq) was added and the reaction mixture was stirred at room temperature for 16 h. Citric acid (370 g, 1.8 mmol, 10 eq) was dissolved in water (4 mL) and added to the reaction mixture dropwise to quench the excess of the borane. The resulting mixture was diluted with water (20 mL) and extracted with ethyl acetate (3 × 15 mL). The organic layer was dried over sodium sulfate, the desiccant was filtered off and the solvent was removed *in vacuo*. The crude product was semi-purified by *np*-flash column chromatography (*iso*-hexanes/ethyl acetate; 0→40% EtOAc) to give **57** (15 mg, ≤56 µmol) which was used without further purification.

Crude **57** (15 mg) was dissolved in THF (2 mL) and water (2 mL), lithium hydroxide (23 mg, 0.56 mmol, 10 eq) was added and the reaction mixture was stirred at room temperature for 1.5 h. The mixture was acidified with hydrochloric acid (2 M, 0.33 mL, 0.67 mmol, 12 eq), the volatiles were removed under reduced pressure and the crude product was dried under high vacuum for 14 h affording **58** which was used without further purification.

Carboxylate **58** was dissolved in anhydrous DMF (0.03 M) and coupled with amine **59** (16 mg, 67 µmol, 1.2 eq) following General Procedure B (1.2 eq COMU, 2 h reaction time). The Crude product was purified by flash column chromatography (*iso*-hexanes/ethyl acetate; 20→80% EtOAc) affording **60** (19 mg, 40 µmol, 23% over 3 steps) as a colourless solid.

**TLC** *Rf* = 0.35 (*np*, Hex:EtOAc 1:1)
**$^1$H-NMR** (400 MHz, $CDCl_3$): *δ* (ppm) = 7.22 (s, 4H), 6.97 (d, J = 8.4 Hz, 2H), 6.56 (d, J = 8.4 Hz, 2H), 5.73 (s, 1H), 4.87 (s, 1H), 4.39 (d, J = 5.7 Hz, 2H), 4.28 (d, J = 5.7 Hz, 2H), 3.65 (t, J = 6.3 Hz, 2H), 3.31 (t, J = 6.6 Hz, 2H), 2.54 (t, J = 7.4 Hz, 2H), 2.19 (t, J = 7.5 Hz, 2H), 2.06 (p, J = 6.5 Hz, 2H), 1.92 (p, J = 7.5 Hz, 2H), 1.45 (s, 9H).
**$^{13}$C-NMR** (101 MHz, $CDCl_3$): *δ* (ppm) = 172.9, 156.0, 145.9, 138.4, 137.6, 130.8, 129.5, 128.2, 127.9, 113.3, 79.7, 44.4, 43.4, 42.8, 41.5, 36.0, 34.4, 32.0, 28.5, 27.6.
**LRMS** (ESI+): m/z calc. for $C_{26}H_{37}ClN_3O_3^+$ [M+H]$^+$: 474.3, found: 474.3.

#### HL3-Fluo

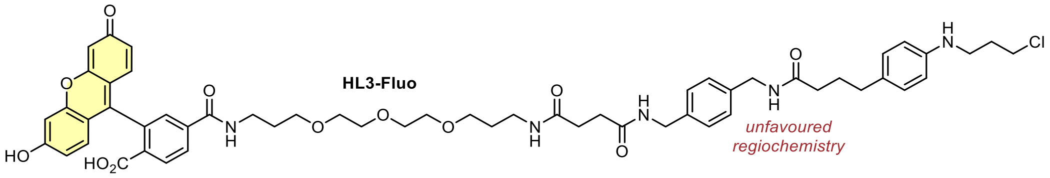

**60** (5 mg, 11 µmol, 1.0 eq) was deprotected according to General Procedure A (0.02 M) and coupled with carboxylate **26** (11 mg, 16 µmol, 1.5 eq, 0.03 M) following General Procedure B (1.2 eq COMU, 3 h reaction time). The crude product was purified by preparative HPLC (acetonitrile/water, 0.1% formic acid; 10→65% MeCN, 20 min) affording **HL3-Fluo** (2.1 mg, 2.1 µmol, 20% over 2 steps) as an orange solid.

**$^1$H-NMR** (400 MHz, DMSO-$d_6$): *δ* (ppm) = 8.66 (t, *J* = 5.5 Hz, 1H), 8.30 (t, *J* = 6.1 Hz, 1H), 8.26 (t, *J* = 5.8 Hz, 1H), 8.15 (dd, *J* = 8.0, 1.2 Hz, 1H), 8.07 (d, *J* = 8.1 Hz, 1H), 7.79 (t, *J* = 5.5 Hz, 1H), 7.66 (s, 1H), 7.16 (s, 4H), 6.88 (d, *J* = 8.4 Hz, 2H), 6.67 (s, 2H), 6.59 (d, *J* = 8.7 Hz, 2H), 6.54 (dd, *J* = 8.7, 2.1 Hz, 2H), 6.48 (d, *J* = 8.5 Hz, 2H), 5.45 (t, *J* = 5.4 Hz, 1H), 4.20 (dd, *J* = 5.8, 2.0 Hz, 4H), 3.73 (t, *J* = 6.5 Hz, 2H), 3.48 – 3.39 (m, 10H), 3.27 – 3.21 (m, 2H), 3.10 (q, *J* = 6.4 Hz, 2H), 3.04 (q, *J* = 6.8 Hz, 2H), 2.42 – 2.36 (m, 3H), 2.31 (dt, *J* = 10.3, 5.1 Hz, 5H), 2.13 – 2.06 (m, 2H), 1.95 (p, *J* = 6.6 Hz, 2H), 1.70 (tt, *J* = 13.2, 7.1 Hz, 4H), 1.57 (p, *J* = 6.6 Hz, 2H).
**HRMS** (ESI+): m/z calc. for $C_{56}H_{65}ClN_5O_{12}^+$ [M+H]$^+$: 1034.431, found: 1034.434.

#### Synthetic route for Cou-HL3-Fluo

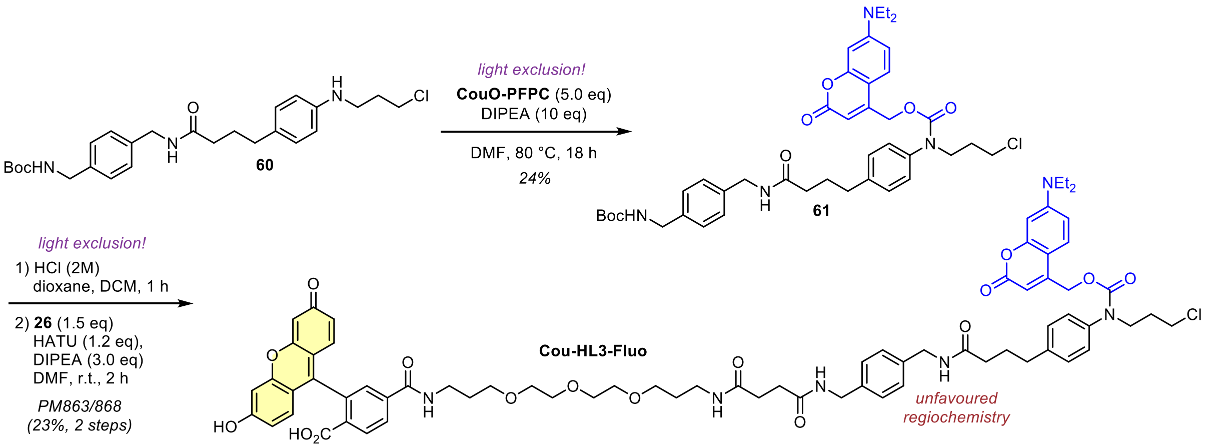

#### Compound 61

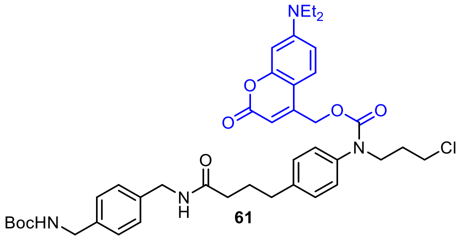

*Note: the reaction must be performed under light exclusion.*

**60** (12 mg, 25 µmol, 1.0 eq) was dissolved in anhydrous DMF (1 mL) transformed to **61** following General Procedure D. The product was semi-purified by *rp*-flash column chromatography (acetonitrile/water, 0.1% formic acid; 20→80% MeCN) affording **61** (4.5 mg, 6.0 µmol, 24%) as a yellow solid.

**TLC** *Rf* = 15 (*np*, Hex:EtOAc 1:1)
**HRMS** (ESI+): m/z calc. for $C_{41}H_{52}ClN_4O_7^+$ [M+H]$^+$: 747.3519, found: 747.3536

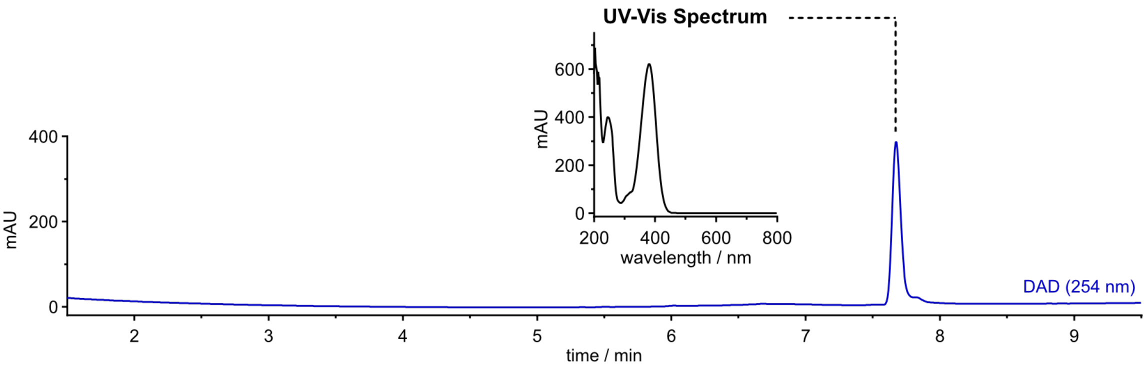

#### Cou-HL3-Fluo

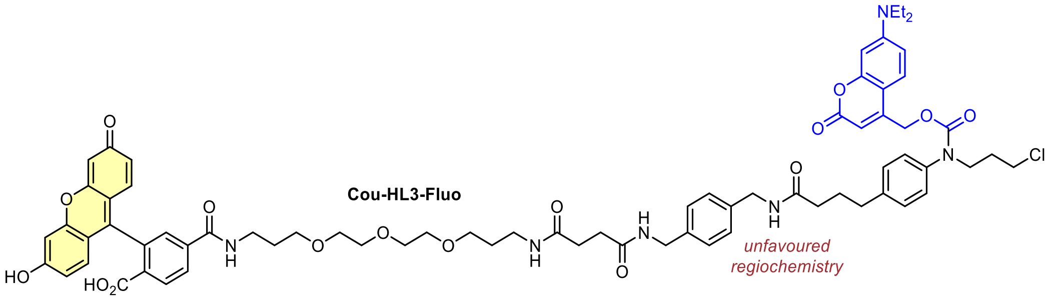

*<u>Note</u>: the reaction must be performed under light exclusion.*

**61** (4.5 mg, 6.0 µmol, 1.0 eq) was deprotected according to General Procedure A (0.02 M) and coupled with carboxylate **26** (6.2 mg, 9.0 µmol, 1.5 eq, 0.03 M) following General Procedure B (1.0 eq HATU, 1 h reaction time). The crude product was purified by preparative HPLC (acetonitrile/water, 0.1% formic acid; 20→70% MeCN, 20 min) affording **Cou-HL3-Fluo** (1.9 mg, 1.4 µmol, 23% over 2 steps) as an orange solid.

**$^1$H-NMR** (400 MHz, DMSO-d$_6$): $\delta$ (ppm) = 8.65 (t, *J* = 5.5 Hz, 1H), 8.31 (t, *J* = 5.7 Hz, 2H), 8.13 (d, *J* = 8.0 Hz, 1H), 8.06 (d, *J* = 8.1 Hz, 1H), 7.79 (t, *J* = 5.5 Hz, 1H), 7.65 (s, 1H), 7.37 (s, 1H), 7.25 (t, *J* = 6.4 Hz, 4H), 7.17 (s, 4H), 6.64 (s, 3H), 6.59 (d, *J* = 8.7 Hz, 3H), 6.52 (d, *J* = 7.2 Hz, 3H), 5.26 (s, 2H), 4.21 (dd, *J* = 8.8, 6.0 Hz, 4H), 3.77 (s, 2H), 3.64 (t, *J* = 6.4 Hz, 2H), 3.43 (ddt, *J* = 14.1, 7.3, 2.9 Hz, 16H), 3.04 (q, *J* = 6.7 Hz, 2H), 2.61 – 2.55 (m, 4H), 2.31 (dt, *J* = 10.2, 4.3 Hz, 4H), 2.16 (t, *J* = 7.3 Hz, 2H), 1.92 (p, *J* = 6.5 Hz, 2H), 1.82 (p, *J* = 7.5 Hz, 2H), 1.69 (q, *J* = 6.6 Hz, 2H), 1.57 (p, *J* = 6.7 Hz, 2H), 1.10 (t, *J* = 7.0 Hz, 6H).
**HRMS** (ESI+): m/z calc. for $C_{71}H_{80}ClN_6O_{16}^+$ $[M+H]^+$: 1307.531, found: 1307.533.

#### Initial synthetic route for CHalo building block (improved procedure above)

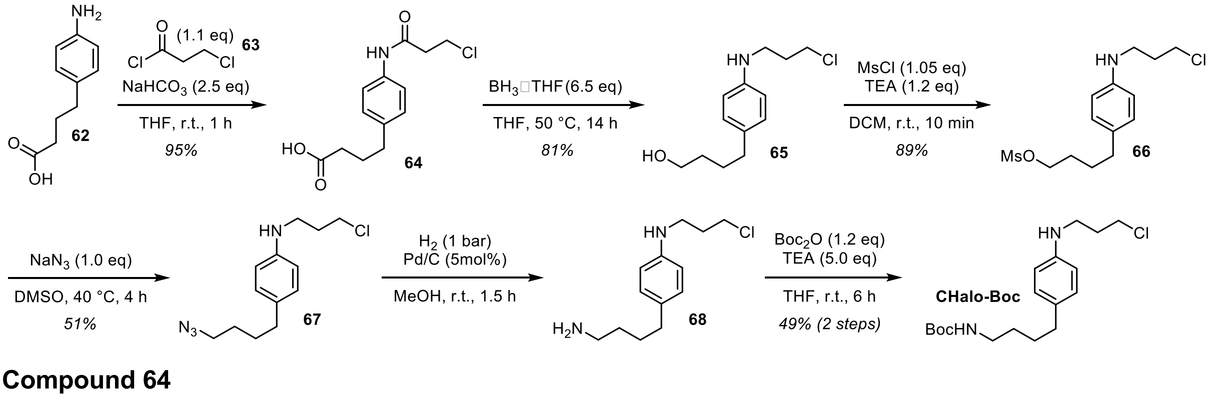

#### Compound 64

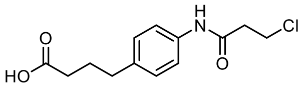

64

4-Aminobenzenebutanoic acid (**62**, 1.50 g, 7.95 mmol, 1.0 eq) was dissolved in anhydrous THF (30 mL). Sodium hydrogen carbonate (1.67 g, 19.9 mmol, 2.5 eq) and 3-chloropropionyl chloride (**63**, 0.835 mL, 8.75 mmol, 1.1 eq) were added and the reaction mixture was stirred at room temperature for 1 h. The solvent was removed *in vacuo* and the residue was diluted with water (15 mL) and hydrochloric acid (2 M, 20 mL, 5.0 eq) and extracted with ethyl acetate (3 × 25 mL). The organic layer was washed with brine (30 mL), dried over sodium sulfate, the desiccant was filtered off and the solvent was removed *in vacuo* affording **64** (2.03 g, 7.53 mmol, 95%) as a colourless solid.

**$^1$H-NMR** (600 MHz, DMSO-d$_6$): *δ* (ppm) = 10.00 (s, 1H), 7.50 (d, J = 8.5 Hz, 2H), 7.11 (d, J = 8.5 Hz, 2H), 3.87 (t, J = 6.3 Hz, 2H), 2.79 (t, J = 6.3 Hz, 2H), 2.56 – 2.51 (m, 2H), 2.19 (t, J = 7.4 Hz, 2H), 1.76 (p, J = 7.5 Hz, 2H).
**$^{13}$C-NMR** (151 MHz, DMSO-d$_6$): *δ* (ppm) = 174.4, 167.8, 136.9, 136.6, 128.6, 119.2, 41.0, 39.2, 33.9, 33.1, 26.4.
**HRMS** (ESI+): m/z calc. for $C_{13}H_{16}ClNNaO_3^+$ [M+Na]$^+$: 202.0711, found: 292.0711.

**Compound 65**

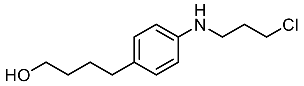

**65**

**64** (2.00 g, 7.41 mmol, 1.0 eq) was dissolved in anhydrous THF (10 mL) under nitrogen atmosphere and borane tetrahydrofurane complex (1 M in THF, 48.2 mL, 48.2 mmol, 6.5 eq) was added. The reaction mixture was heated to 50 °C for 14 h. Methanol (ca. 10 mL) was added slowly until no gas development was observed upon addition any more to quench the excess of the borane and the volatiles were removed *in vacuo*. The crude product was purified by flash column chromatography (DCM/MeOH; 0→2% MeOH) to give **65** (1.46 g, 6.04 mmol, 81%) as a light-yellow oil.

**TLC** *Rf* = 0.18 (*np*, DCM:MeOH 99:1)
**$^1$H-NMR** (400 MHz, MeOD): *δ* (ppm) = 6.95 (d, J = 8.4 Hz, 2H), 6.59 (d, J = 8.4 Hz, 2H), 3.67 (t, J = 6.4 Hz, 2H), 3.55 (t, J = 6.4 Hz, 2H), 3.23 (t, J = 6.7 Hz, 2H), 2.50 (t, J = 7.3 Hz, 2H), 2.03 (p, J = 6.6 Hz, 2H), 1.65 – 1.48 (m, 4H).
**$^{13}$C-NMR** (101 MHz, MeOD): *δ* (ppm) = 148.0, 132.4, 130.0, 114.4, 62.9, 43.6, 42.4, 35.8, 33.3, 33.2, 29.2.
**HRMS** (ESI+): m/z calc. for $C_{13}H_{21}ClNO^+$ [M+H]$^+$: 242.1306, found: 242.1310.

**Compound 66**

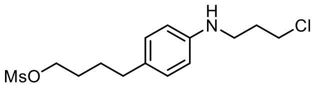

**66**

**65** (0.17 g, 0.70 mmol, 1.0 eq) was dissolved in anhydrous DCM (4 mL) under nitrogen atmosphere, triethylamine (0.17 mL, 0.84 mmol, 1.2 eq) was added and the reaction was cooled to 0 °C. Methanesulfonyl chloride (57 µL, 0.74 mmol, 1.05 eq) was added, the reaction was allowed to warm to room temperature, stirred for 10 min and then diluted with ethyl acetate (15 mL). The organic layer was washed with water (3 × 10 mL), dried over sodium sulfate and the solvent was removed *in vacuo* affording **66** (0.20 g, 0.62 mmol, 89%) as a colourless oil.

**TLC** *Rf* = 0.71 (*np*, DCM:MeOH 99:1)
**$^1$H-NMR** (400 MHz, CDCl$_3$): *δ* (ppm) = 6.99 (d, J = 8.4 Hz, 2H), 6.60 (d, J = 8.4 Hz, 2H), 4.22 (t, J = 6.3 Hz, 2H), 3.65 (t, J = 6.3 Hz, 2H), 3.32 (t, J = 6.6 Hz, 2H), 2.98 (s, 3H), 2.55 (t, J = 7.3 Hz, 2H), 2.07 (p, J = 6.5 Hz, 2H), 1.81 – 1.73 (m, 2H), 1.73 – 1.63 (m, 2H).
**LRMS** (ESI+): m/z calc. for $C_{14}H_{23}ClNO_3S^+$ [M+H]$^+$: 320.1, found: 320.1.

**Compound 67**

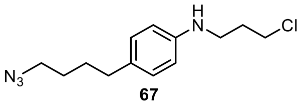

**67**

**66** (1.69 g, 5.28 mmol, 1.0 eq) was dissolved in anhydrous DMSO (6 mL) under nitrogen atmosphere, sodium azide (0.34 g, 5.28 mmol, 1.0 eq) was added and the reaction mixture was heated to 40 °C for 4 h. The mixture was diluted with water (15 mL), brine (40 mL) and extracted with ethyl acetate (40 mL). The organic layer was washed with a 1:1 mixture of water and brine (30 mL), dried over sodium sulfate and the solvent was removed *in vacuo*. The crude product was purified by flash column chromatography (*iso*-hexanes /ethyl acetate; 0→10% EtOAc) to give **67** (0.72 g, 2.7 mmol, 51%) as a colourless oil.

*Note: azide substitution of the alkyl chloride is observed as a side reaction.*

**TLC** *Rf* = 0.71 (*np*, Hex:EtOAc 4:1)
**$^1$H-NMR** (400 MHz, CDCl$_3$): *δ* (ppm) = 7.01 (d, J = 8.1 Hz, 2H), 6.63 (d, J = 8.1 Hz, 2H), 3.65 (t, J = 6.2 Hz, 2H), 3.33 (t, J = 6.6 Hz, 2H), 3.27 (t, J = 6.1 Hz, 2H), 2.54 (t, J = 6.6 Hz, 2H), 2.08 (p, J = 6.3 Hz, 2H), 1.70 – 1.57 (m, 4H).

**$^{13}$C-NMR** (101 MHz, $CDCl_3$): *δ* (ppm) = 145.4, 131.8, 129.4, 113.7, 51.5, 42.7, 41.8, 34.6, 31.9, 28.9, 28.5.
**HRMS** (ESI+): m/z calc. for $C_{13}H_{20}ClN_4^+$ $[M+H]^+$: 267.1371, found: 267.1376.

##### Compound CHalo-Boc

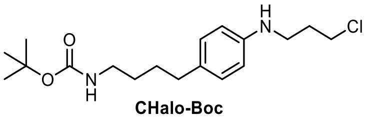

**CHalo-Boc**

*Note*: here we report the initial synthetic procedure towards **CHalo-Boc**, an optimised preparation is described above.

**67** (50 mg, 0.19 mmol, 1.0 eq) was dissolved in methanol (2 mL) under nitrogen atmosphere, palladium on charcoal (Pd/C, 11 mg, 9.4 µmol, 5 mol%) was added and the reaction was stirred under hydrogen atmosphere (1 atm) for 1.5 h. The Pd/C was removed by filtration over Celite (washed with methanol (4×)) and the filtrate was concentrated under reduce pressure. The intermediate product **68** was suspended in anhydrous THF (2.5 mL), triethylamine (0.13 mL, 0.94 mmol, 5.0 eq) and di-*tert*-butyl dicarbonate (51 mg, 0.22 mmol, 1.2 eq) were added and the reaction mixture was stirred at room temperature for 6 h. The mixture was diluted with water (15 mL) and extracted with DCM (3 × 10 mL). The organic layer was dried over sodium sulfate and the solvent was removed *in vacuo*. The crude product was purified by flash column chromatography (*iso*-hexanes/ethyl acetate; 0→30% EtOAc) to give **CHalo-Boc** (31 mg, 91 µmol, 49% over 2 steps) as a colourless oil.

**TLC** *Rf* = 0.34 (*np*, Hex:EtOAc 9:1)
**$^1$H-NMR** (600 MHz, $CDCl_3$): *δ* (ppm) = 6.99 (d, J = 8.4 Hz, 2H), 6.56 (d, J = 8.4 Hz, 2H), 4.52 (s, 1H), 3.65 (t, J = 6.3 Hz, 2H), 3.31 (t, J = 6.6 Hz, 2H), 3.12 (q, J = 6.7 Hz, 2H), 2.51 (t, J = 7.4 Hz, 2H), 2.06 (p, J = 6.5 Hz, 2H), 1.63 – 1.53 (m, 2H), 1.53 – 1.46 (m, 2H), 1.44 (s, 9H).
**$^{13}$C-NMR** (151 MHz, $CDCl_3$): *δ* (ppm) = 156.1, 146.0, 131.4, 129.3, 113.0, 79.1, 42.8, 41.3, 40.6, 34.6, 32.1, 29.7, 29.0, 28.5.
**HRMS** (ESI+): m/z calc. for $C_{18}H_{29}ClN_2NaO_2^+$ $[M+Na]^+$: 363.1810, found: 363.1812.

##### CHalo-Fluo

**18**

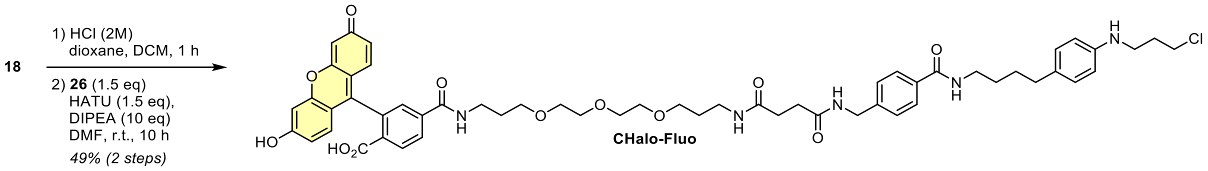

**CHalo-Fluo**

**18** (3 mg, 6 µmol, 1.0 eq) was deprotected according to General Procedure A (0.02 M) and coupled with carboxylate **26** (7 mg, 10 µmol, 1.5 eq, 0.05 M) following General Procedure B (1.0 eq HATU, 10 h reaction time). The crude product was purified by preparative HPLC (acetonitrile/water, 0.1% formic acid; 10→60% MeCN, 20 min) affording **CHalo-Fluo** (3.2 mg, 4.1 µmol, 49% over 2 steps) as an orange solid.

**$^1$H-NMR** (800 MHz, DMSO-$d_6$): *δ* (ppm) = 10.16 (s, 2H), 8.66 (t, J = 5.4 Hz, 1H), 8.38 (t, J = 5.5 Hz, 2H), 8.16 (d, J = 8.1 Hz, 1H), 8.07 (d, J = 8.1 Hz, 1H), 7.79 (t, J = 5.3 Hz, 1H), 7.76 (d, J = 8.1 Hz, 2H), 7.66 (s, 1H), 7.29 (d, J = 8.0 Hz, 2H), 6.89 (d, J = 8.2 Hz, 2H), 6.69 (d, J = 1.8 Hz, 2H), 6.58 (d, J = 8.7 Hz, 2H), 6.55 (dd, J = 8.7, 1.9 Hz, 2H), 6.48 (d, J = 8.2 Hz, 2H), 5.42 (s, 1H), 4.30 – 4.26 (m, 2H), 3.72 (t, J = 6.5 Hz, 2H), 3.47 – 3.44 (m, 4H), 3.44 – 3.40 (m, 4H), 3.38 (t, J = 6.2 Hz, 2H), 3.36 – 3.34 (m, 2H), 3.26 – 3.22 (m, 4H), 3.09 (q, J = 6.2 Hz, 2H), 3.05 (q, J = 6.6 Hz, 2H), 2.43 (t, J = 6.9 Hz, 2H), 2.37 (t, J = 7.2 Hz, 2H), 2.31 (t, J = 7.2 Hz, 2H), 1.95 (p, J = 6.5 Hz, 2H), 1.68 (p, J = 6.5 Hz, 2H), 1.57 (p, J = 6.6 Hz, 2H), 1.54 – 1.46 (m, 4H).
**$^{13}$C-NMR** (201 MHz, DMSO-$d_6$): *δ* (ppm) = 171.5, 171.2, 168.1, 165.8, 164.4, 159.6, 152.7, 151.8, 146.8, 142.7, 140.8, 133.1, 129.4, 129.3, 129.2, 128.8, 128.1, 127.1, 126.8, 124.9, 122.2, 112.7, 112.1, 109.2, 102.2, 83.3, 69.7, 69.7, 69.5, 69.5, 68.2, 68.0, 43.4, 41.7, 40.1, 39.0, 36.9, 35.8, 34.0, 31.7, 30.8, 30.7, 29.4, 29.1, 28.9, 28.8.
**HRMS** (ESI+): m/z calc. for $C_{56}H_{65}ClN_5O_{12}^+$ $[M+H]^+$: 1034.431, found: 1034.431.

#### Cou-CHalo-Fluo

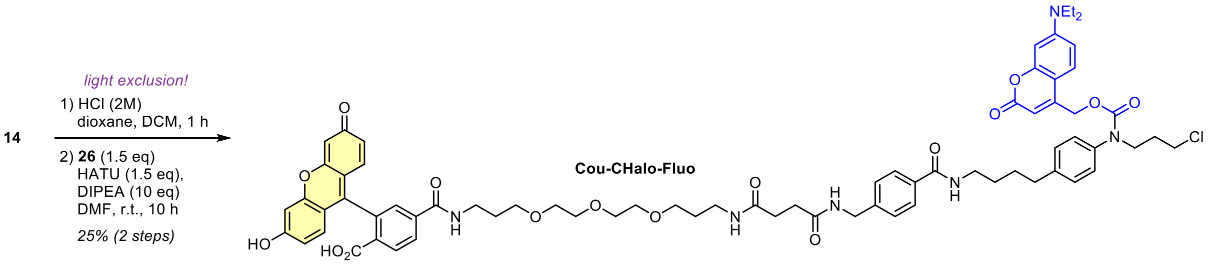

*<u>Note</u>: the reaction must be performed under light exclusion.*

**14** (3 mg, 4 µmol, 1.0 eq) was deprotected according to General Procedure A (0.02 M) and coupled with carboxylate **26** (4 mg, 6 µmol, 1.5 eq, 0.05 M) following General Procedure B (1.0 eq HATU, 10 h reaction time). The crude product was purified by preparative HPLC (acetonitrile/water, 0.1% formic acid; 10→75% MeCN, 20 min) affording **Cou-CHalo-Fluo** (1.3 mg, 1.0 µmol, 25% over 2 steps) as an orange solid.

**$^1$H-NMR** (400 MHz, DMSO-d$_6$): $\delta$ (ppm) = 10.17 (s, 2H), 8.67 (t, *J* = 5.5 Hz, 1H), 8.40 (q, *J* = 5.9 Hz, 2H), 8.16 (d, *J* = 8.1 Hz, 1H), 8.07 (d, *J* = 8.0 Hz, 1H), 7.78 (dd, *J* = 15.0, 6.8 Hz, 3H), 7.66 (s, 1H), 7.43 – 7.34 (m, 1H), 7.29 (d, *J* = 8.2 Hz, 2H), 7.25 (s, 4H), 6.69 (d, *J* = 1.9 Hz, 2H), 6.65 (d, *J* = 8.6 Hz, 1H), 6.60 – 6.55 (m, 4H), 6.55 – 6.49 (m, 2H), 5.25 (s, 2H), 4.28 (d, *J* = 5.8 Hz, 2H), 3.80 – 3.73 (m, 2H), 3.64 (t, *J* = 6.4 Hz, 2H), 3.42 (tt, *J* = 11.9, 5.6 Hz, 16H), 3.26 (dq, *J* = 13.3, 6.6 Hz, 4H), 3.05 (q, *J* = 6.7 Hz, 2H), 2.64 – 2.59 (m, 2H), 2.40 – 2.34 (m, 2H), 2.32 (d, *J* = 6.3 Hz, 2H), 1.91 (p, *J* = 6.5 Hz, 2H), 1.68 (p, *J* = 6.4 Hz, 2H), 1.57 (dt, *J* = 12.6, 6.2 Hz, 6H), 1.10 (t, *J* = 6.9 Hz, 6H).
**HRMS** (ESI+): m/z calc. for $C_{71}H_{80}ClN_6O_{16}^+$ $[M+H]^+$: 1307.531, found: 1307.533.

### 11 References

(1) Merrill, R. A.; Song, J.; Kephart, R. A.; Klomp, A. J.; Noack, C. E.; Strack, S. A Robust and Economical Pulse-Chase Protocol to Measure the Turnover of HaloTag Fusion Proteins. *J. Biol. Chem.* **2019**, *294* (44), 16164–16171. https://doi.org/10.1074/jbc.RA119.010596.

(2) Los, G. V.; Encell, L. P.; McDougall, M. G.; Hartzell, D. D.; Karassina, N.; Zimprich, C.; Wood, M. G.; Learish, R.; Ohana, R. F.; Urh, M.; Simpson, D.; Mendez, J.; Zimmerman, K.; Otto, P.; Vidugiris, G.; Zhu, J.; Darzins, A.; Klaubert, D. H.; Bulleit, R. F.; Wood, K. V. HaloTag: A Novel Protein Labeling Technology for Cell Imaging and Protein Analysis. *ACS Chem. Biol.* **2008**, *3* (6), 373–382. https://doi.org/10.1021/cb800025k.

(3) Chyan, W.; Raines, R. T. Enzyme-Activated Fluorogenic Probes for Live-Cell and in Vivo Imaging. *ACS Chem. Biol.* **2018**, *13* (7), 1810–1823. https://doi.org/10.1021/acschembio.8b00371.

(4) Egyed, A.; Németh, K.; Molnár, T. Á.; Kállay, M.; Kele, P.; Bojtár, M. Turning Red without Feeling Embarrassed─Xanthenium-Based Photocages for Red-Light-Activated Phototherapeutics. *J. Am. Chem. Soc.* **2023**. https://doi.org/10.1021/jacs.2c11499.

(5) Weinstain, R.; Slanina, T.; Kand, D.; Klán, P. Visible-to-NIR-Light Activated Release: From Small Molecules to Nanomaterials. *Chem. Rev.* **2020**, *120* (24), 13135–13272. https://doi.org/10.1021/acs.chemrev.0c00663.

(6) Messina, M. S.; Quargnali, G.; Chang, C. J. Activity-Based Sensing for Chemistry-Enabled Biology: Illuminating Principles, Probes, and Prospects for Boronate Reagents for Studying Hydrogen Peroxide. *ACS Bio Med Chem Au* **2022**, *2* (6), 548–564. https://doi.org/10.1021/acsbiomedchemau.2c00052.

(7) Fosnacht, K. G.; Hammers, M. D.; Earp, M. S.; Gilbert, A. K.; Pluth, M. D. A Cell Trappable Methyl Rhodol-Based Fluorescent Probe for Hydrogen Sulfide Detection. *Chem. Asian J.* **2022**, *17* (16), e202200426. https://doi.org/10.1002/asia.202200426.

(8) Shields, B. C.; Yan, H.; Lim, S. S. X.; Burwell, S. C. V.; Cammarata, C. M.; Fleming, E. A.; Yousefzadeh, S. A.; Goldenshtein, V. Z.; Kahuno, E. W.; Vagadia, P. P.; Loughran, M. H.; Zhiquan, L.; McDonnell, M. E.; Scalabrino, M. L.; Thapa, M.; Hawley, T. M.; Field, G. D.; Hull, C.; Schiltz, G. E.; Glickfeld, L. L.; Reitz, A. B.; Tadross, M. R. DART.2: Bidirectional Synaptic Pharmacology with Thousandfold Cellular Specificity. *Nat. Methods* **2024**, *21* (7), 1288–1297. https://doi.org/10.1038/s41592-024-02292-9.

(9) Wang, L.; Tran, M.; D'Este, E.; Roberti, J.; Koch, B.; Xue, L.; Johnsson, K. A General Strategy to Develop Cell Permeable and Fluorogenic Probes for Multicolour Nanoscopy. *Nat. Chem.* **2019**, 1–8. https://doi.org/10.1038/s41557-019-0371-1.

(10) Lukinavičius, G.; Umezawa, K.; Olivier, N.; Honigmann, A.; Yang, G.; Plass, T.; Mueller, V.; Reymond, L.; Corrêa Jr, I. R.; Luo, Z.-G.; Schultz, C.; Lemke, E. A.; Heppenstall, P.; Eggeling, C.; Manley, S.; Johnsson, K. A Near-Infrared Fluorophore for Live-Cell Super-Resolution Microscopy of Cellular Proteins. *Nat. Chem.* **2013**, *5* (2), 132–139. https://doi.org/10.1038/nchem.1546.

(11) Schmitt, C.; Mauker, P.; Vepřek, N. A.; Gierse, C.; Meiring, J. C. M.; Kuch, J.; Akhmanova, A.; Dehmelt, L.; Thorn-Seshold, O. A Photocaged Microtubule-Stabilising Epothilone Allows Spatiotemporal Control of Cytoskeletal Dynamics. *Angew. Chem. Int. Ed.* **2024**, *63* (43), e202410169. https://doi.org/10.1002/anie.202410169.

(12) Wilhelm, J.; Kühn, S.; Tarnawski, M.; Gotthard, G.; Tünnermann, J.; Tänzer, T.; Karpenko, J.; Mertes, N.; Xue, L.; Uhrig, U.; Reinstein, J.; Hiblot, J.; Johnsson, K. Kinetic and Structural Characterization of the Self-Labeling Protein Tags HaloTag7, SNAP-Tag, and CLIP-Tag. *Biochemistry* **2021**, *60* (33), 2560–2575. https://doi.org/10.1021/acs.biochem.1c00258.

(13) Russo, M.; Janeková, H.; Meier, D.; Generali, M.; Štacko, P. Light in a Heartbeat: Bond Scission by a Single Photon above 800 Nm. *J. Am. Chem. Soc.* **2024**, *146* (12), 8417–8424. https://doi.org/10.1021/jacs.3c14197.

(14) Gorka, A. P.; Yamamoto, T.; Zhu, J.; Schnermann, M. J. Cyanine Photocages Enable Spatial Control of Inducible Cre-Mediated Recombination. *ChemBioChem* **2018**, *19* (12), 1239–1243. https://doi.org/10.1002/cbic.201800061.

(15) Yan, X.; Zhao, J.-H.; Wang, Q.; Wang, W.; Ding, Y.; Zhou, Y.; Chen, G.; Du, J.; Huang, W.; Chu, L. Silicon Rhodamine-Catalyzed Near-Infrared Light-Induced Photodecaging of Ortho-Nitrobenzyl Groups In Vitro and In Vivo. *J. Am. Chem. Soc.* **2025**, *147* (24), 20957–20966. https://doi.org/10.1021/jacs.5c04942.

(16) Zeisel, L.; Felber, J. G.; Scholzen, K. C.; Poczka, L.; Cheff, D.; Maier, M. S.; Cheng, Q.; Shen, M.; Hall, M. D.; Arnér, E. S. J.; Thorn-Seshold, J.; Thorn-Seshold, O. Selective Cellular Probes for Mammalian Thioredoxin Reductase TrxR1: Rational Design of RX1, a Modular 1,2-Thiaselenane Redox Probe. *Chem* **2022**, *8* (5), 1493–1517. https://doi.org/10.1016/j.chempr.2022.03.010.

(17) Zeisel, L.; Felber, J. G.; Scholzen, K. C.; Schmitt, C.; Wiegand, A. J.; Komissarov, L.; Arnér, E. S. J.; Thorn-Seshold, O. Piperazine-Fused Cyclic Disulfides Unlock High-Performance Bioreductive Probes of Thioredoxins and Bifunctional Reagents for Thiol Redox Biology. *J. Am. Chem. Soc.* **2024**, *146* (8), 5204–5214. https://doi.org/10.1021/jacs.3c11153.

(18) Papot, S.; Tranoy, I.; Tillequin, F.; Florent, J.-C.; Gesson, J.-P. Design of Selectively Activated Anticancer Prodrugs: Elimination and Cyclization Strategies. *Curr. Med. Chem. Anticancer Agents* **2002**, *2* (2), 155–185. https://doi.org/10.2174/1568011023354173.

(19) Jasim, S. Development of a Dehalogenase-Based Protein Fusion Tag Capable of Rapid, Selective and Covalent Attachment to Customizable Ligands. *AACE Clin. Case Rep.* **2021**, *7* (1), 1.
(20) Los, G. V.; Encell, L. P.; McDougall, M. G.; Hartzell, D. D.; Karassina, N.; Zimprich, C.; Wood, M. G.; Learish, R.; Ohana, R. F.; Urh, M.; Simpson, D.; Mendez, J.; Zimmerman, K.; Otto, P.; Vidugiris, G.; Zhu, J.; Darzins, A.; Klaubert, D. H.; Bulleit, R. F.; Wood, K. V. HaloTag: A Novel Protein Labeling Technology for Cell Imaging and Protein Analysis. *ACS Chem. Biol.* **2008**, *3* (6), 373–382. https://doi.org/10.1021/cb800025k.
(21) Sun, X.; Zhang, A.; Baker, B.; Sun, L.; Howard, A.; Buswell, J.; Maurel, D.; Masharina, A.; Johnsson, K.; Noren, C. J.; Xu, M.-Q.; Corrêa Jr., I. R. Development of SNAP-Tag Fluorogenic Probes for Wash-Free Fluorescence Imaging. *ChemBioChem* **2011**, *12* (14), 2217–2226. https://doi.org/10.1002/cbic.201100173.
(22) Gronemeyer, T.; Chidley, C.; Juillerat, A.; Heinis, C.; Johnsson, K. Directed Evolution of O6-Alkylguanine-DNA Alkyltransferase for Applications in Protein Labeling. *Protein Eng. Des. Sel.* **2006**, *19* (7), 309–316. https://doi.org/10.1093/protein/gzl014.
(23) Gautier, A.; Juillerat, A.; Heinis, C.; Corrêa, I. R.; Kindermann, M.; Beaufils, F.; Johnsson, K. An Engineered Protein Tag for Multiprotein Labeling in Living Cells. *Chem. Biol.* **2008**, *15* (2), 128–136. https://doi.org/10.1016/j.chembiol.2008.01.007.
(24) Porzberg, N.; Gries, K.; Johnsson, K. Exploiting Covalent Chemical Labeling with Self-Labeling Proteins. *Annu. Rev. Biochem.* **2025**, *94* (Volume 94, 2025), 29–58. https://doi.org/10.1146/annurev-biochem-030222-121016.
(25) Deo, C.; Abdelfattah, A. S.; Bhargava, H. K.; Berro, A. J.; Falco, N.; Farrants, H.; Moeyaert, B.; Chupanova, M.; Lavis, L. D.; Schreiter, E. R. The HaloTag as a General Scaffold for Far-Red Tunable Chemigenetic Indicators. *Nat. Chem. Biol.* **2021**, *17* (6), 718–723. https://doi.org/10.1038/s41589-021-00775-w.
(26) Grimm, J. B.; Tkachuk, A. N.; Xie, L.; Choi, H.; Mohar, B.; Falco, N.; Schaefer, K.; Patel, R.; Zheng, Q.; Liu, Z.; Lippincott-Schwartz, J.; Brown, T. A.; Lavis, L. D. A General Method to Optimize and Functionalize Red-Shifted Rhodamine Dyes. *Nat. Methods* **2020**, *17* (8), 815–821. https://doi.org/10.1038/s41592-020-0909-6.
(27) Schwenzer, N.; Teiwes, N. K.; Kohl, T.; Pohl, C.; Giller, M. J.; Lehnart, S. E.; Steinem, C. CaV1.3 Channel Clusters Characterized by Live-Cell and Isolated Plasma Membrane Nanoscopy. *Commun. Biol.* **2024**, *7* (1), 1–11. https://doi.org/10.1038/s42003-024-06313-3.
(28) Roed, S. N.; Wismann, P.; Underwood, C. R.; Kulahin, N.; Iversen, H.; Cappelen, K. A.; Schäffer, L.; Lehtonen, J.; Hecksher-Soerensen, J.; Secher, A.; Mathiesen, J. M.; Bräuner-Osborne, H.; Whistler, J. L.; Knudsen, S. M.; Waldhoer, M. Real-Time Trafficking and Signaling of the Glucagon-like Peptide-1 Receptor. *Mol. Cell. Endocrinol.* **2014**, *382* (2), 938–949. https://doi.org/10.1016/j.mce.2013.11.010.
(29) Komatsubara, A. T.; Goto, Y.; Kondo, Y.; Matsuda, M.; Aoki, K. Single-Cell Quantification of the Concentrations and Dissociation Constants of Endogenous Proteins. *J. Biol. Chem.* **2019**, *294* (15), 6062–6072. https://doi.org/10.1074/jbc.RA119.007685.
(30) Kohl, J.; Ng, J.; Cachero, S.; Ciabatti, E.; Dolan, M.-J.; Sutcliffe, B.; Tozer, A.; Ruehle, S.; Krueger, D.; Frechter, S.; Branco, T.; Tripodi, M.; Jefferis, G. S. X. E. Ultrafast Tissue Staining with Chemical Tags. *Proc. Natl. Acad. Sci. U.S.A.* **2014**, *111* (36), E3805–E3814. https://doi.org/10.1073/pnas.1411087111.
(31) Zimmermann, M.; Cal, R.; Janett, E.; Hoffmann, V.; Bochet, C. G.; Constable, E.; Beaufils, F.; Wymann, M. P. Cell-Permeant and Photocleavable Chemical Inducer of Dimerization. *Angew. Chem. Int. Ed.* **2014**, *53* (18), 4717–4720. https://doi.org/10.1002/anie.201310969.
(32) Erhart, D.; Zimmermann, M.; Jacques, O.; Wittwer, M. B.; Ernst, B.; Constable, E.; Zvelebil, M.; Beaufils, F.; Wymann, M. P. Chemical Development of Intracellular Protein Heterodimerizers. *Chem. Biol.* **2013**, *20* (4), 549–557. https://doi.org/10.1016/j.chembiol.2013.03.010.
(33) Ballister, E. R.; Aonbangkhen, C.; Mayo, A. M.; Lampson, M. A.; Chenoweth, D. M. Localized Light-Induced Protein Dimerization in Living Cells Using a Photocaged Dimerizer. *Nat. Commun.* **2014**, *5* (1), 5475. https://doi.org/10.1038/ncomms6475.
(34) Deo, C.; Sheu, S.-H.; Seo, J.; Clapham, D. E.; Lavis, L. D. Isomeric Tuning Yields Bright and Targetable Red Ca2+ Indicators. *J. Am. Chem. Soc.* **2019**, *141* (35), 13734–13738. https://doi.org/10.1021/jacs.9b06092.
(35) Neklesa, T. K.; Tae, H. S.; Schneekloth, A. R.; Stulberg, M. J.; Corson, T. W.; Sundberg, T. B.; Raina, K.; Holley, S. A.; Crews, C. M. Small-Molecule Hydrophobic Tagging–Induced Degradation of HaloTag Fusion Proteins. *Nat. Chem. Biol.* **2011**, *7* (8), 538–543. https://doi.org/10.1038/nchembio.597.
(36) Shields, B. C.; Kahuno, E.; Kim, C.; Apostolides, P. F.; Brown, J.; Lindo, S.; Mensh, B. D.; Dudman, J. T.; Lavis, L. D.; Tadross, M. R. Deconstructing Behavioral Neuropharmacology with Cellular Specificity. *Science* **2017**, *356* (6333), eaaj2161. https://doi.org/10.1126/science.aaj2161.
(37) Lin, H.-Y.; Haegele, J. A.; Disare, M. T.; Lin, Q.; Aye, Y. A Generalizable Platform for Interrogating Target- and Signal-Specific Consequences of Electrophilic Modifications in Redox-Dependent Cell Signaling. *J. Am. Chem. Soc.* **2015**, *137* (19), 6232–6244. https://doi.org/10.1021/ja5132648.
(38) Wilhelm, J.; Kühn, S.; Tarnawski, M.; Gotthard, G.; Tünnermann, J.; Tänzer, T.; Karpenko, J.; Mertes, N.; Xue, L.; Uhrig, U.; Reinstein, J.; Hiblot, J.; Johnsson, K. Kinetic and Structural Characterization of

the Self-Labeling Protein Tags HaloTag7, SNAP-Tag, and CLIP-Tag. *Biochemistry* **2021**, *60* (33), 2560–2575. https://doi.org/10.1021/acs.biochem.1c00258.

(39) Ohana, R. F.; Encell, L. P.; Zhao, K.; Simpson, D.; Slater, M. R.; Urh, M.; Wood, K. V. HaloTag7: A Genetically Engineered Tag That Enhances Bacterial Expression of Soluble Proteins and Improves Protein Purification. *Protein Expression Purif.* **2009**, *68* (1), 110–120. https://doi.org/10.1016/j.pep.2009.05.010.

(40) R. Correa, I.; Baker, B.; Zhang, A.; Sun, L.; R. Provost, C.; Lukinavicius, G.; Reymond, L.; Johnsson, K.; Xu, M.-Q. Substrates for Improved Live-Cell Fluorescence Labeling of SNAP-Tag. *Curr. Pharm. Des.* **2013**, *19* (30), 5414–5420.

(41) Kompa, J.; Bruins, J.; Glogger, M.; Wilhelm, J.; Frei, M. S.; Tarnawski, M.; D'Este, E.; Heilemann, M.; Hiblot, J.; Johnsson, K. Exchangeable HaloTag Ligands for Super-Resolution Fluorescence Microscopy. *J. Am. Chem. Soc.* **2023**, *145* (5), 3075–3083. https://doi.org/10.1021/jacs.2c11969.

(42) Lincoln, R.; Bossi, M. L.; Remmel, M.; D'Este, E.; Butkevich, A. N.; Hell, S. W. A General Design of Caging-Group-Free Photoactivatable Fluorophores for Live-Cell Nanoscopy. *Nat. Chem.* **2022**, *14* (9), 1013–1020. https://doi.org/10.1038/s41557-022-00995-0.

(43) Lippert, A. R.; Van de Bittner, G. C.; Chang, C. J. Boronate Oxidation as a Bioorthogonal Reaction Approach for Studying the Chemistry of Hydrogen Peroxide in Living Systems. *Acc. Chem. Res.* **2011**, *44* (9), 793–804. https://doi.org/10.1021/ar200126t.

(44) Albers, A. E.; Dickinson, B. C.; Miller, E. W.; Chang, C. J. A Red-Emitting Naphthofluorescein-Based Fluorescent Probe for Selective Detection of Hydrogen Peroxide in Living Cells. *Bioorg. Med. Chem. Lett.* **2008**, *18* (22), 5948–5950. https://doi.org/10.1016/j.bmcl.2008.08.035.

(45) Gruber, T. D.; Krishnamurthy, C.; Grimm, J. B.; Tadross, M. R.; Wysocki, L. M.; Gartner, Z. J.; Lavis, L. D. Cell-Specific Chemical Delivery Using a Selective Nitroreductase–Nitroaryl Pair. *ACS Chem. Biol.* **2018**, *13* (10), 2888–2896. https://doi.org/10.1021/acschembio.8b00524.

(46) Li, L.; Ge, J.; Wu, H.; Xu, Q.-H.; Yao, S. Q. Organelle-Specific Detection of Phosphatase Activities with Two-Photon Fluorogenic Probes in Cells and Tissues. *J. Am. Chem. Soc.* **2012**, *134* (29), 12157–12167. https://doi.org/10.1021/ja3036256.

(47) Ito, H.; Kawamata, Y.; Kamiya, M.; Tsuda-Sakurai, K.; Tanaka, S.; Ueno, T.; Komatsu, T.; Hanaoka, K.; Okabe, S.; Miura, M.; Urano, Y. Red-Shifted Fluorogenic Substrate for Detection of lacZ-Positive Cells in Living Tissue with Single-Cell Resolution. *Angew. Chem. Int. Ed.* **2018**, *57* (48), 15702–15706. https://doi.org/10.1002/anie.201808670.

(48) Tian, L.; Yang, Y.; Wysocki, L. M.; Arnold, A. C.; Hu, A.; Ravichandran, B.; Sternson, S. M.; Looger, L. L.; Lavis, L. D. Selective Esterase–Ester Pair for Targeting Small Molecules with Cellular Specificity. *Proc. Natl. Acad. Sci. U.S.A.* **2012**, *109* (13), 4756–4761. https://doi.org/10.1073/pnas.1111943109.

(49) Turnbull, J. L.; Benlian, B. R.; Golden, R. P.; Miller, E. W. Phosphonofluoresceins: Synthesis, Spectroscopy, and Applications. *J. Am. Chem. Soc.* **2021**, *143* (16), 6194–6201. https://doi.org/10.1021/jacs.1c01139.

(50) Dickinson, B. C.; Peltier, J.; Stone, D.; Schaffer, D. V.; Chang, C. J. Nox2 Redox Signaling Maintains Essential Cell Populations in the Brain. *Nat. Chem. Biol.* **2011**, *7* (2), 106–112. https://doi.org/10.1038/nchembio.497.

(51) Mauker, P.; Dessen-Weissenhorn, L.; Zecha, C.; Vepřek, N. A.; Brandmeier, J. I.; Beckmann, D.; Kitowski, A.; Kernmayr, T.; Thorn-Seshold, J.; Kerschensteiner, M.; Thorn-Seshold, O. Cellularly-Retained Fluorogenic Probes for Sensitive Cell-Resolved Bioactivity Imaging. bioRxiv April 20, 2025, p 2025.04.17.649302. https://doi.org/10.1101/2025.04.17.649302.

(52) Huang, Z.; Terpetschnig, E.; You, W.; Haugland, R. P. 2-(2′-Phosphoryloxyphenyl)-4(3*H*)-Quinazolinone Derivatives as Fluorogenic Precipitating Substrates of Phosphatases. *Anal.Biochem.* **1992**, *207* (1), 32–39. https://doi.org/10.1016/0003-2697(92)90495-S.

(53) Doura, T.; Kamiya, M.; Obata, F.; Yamaguchi, Y.; Hiyama, T. Y.; Matsuda, T.; Fukamizu, A.; Noda, M.; Miura, M.; Urano, Y. Detection of LacZ-Positive Cells in Living Tissue with Single-Cell Resolution. *Angew. Chem. Int. Ed.* **2016**, *55* (33), 9620–9624. https://doi.org/10.1002/anie.201603328.

(54) Sun, H.; Johnson, D. R.; Finch, R. A.; Sartorelli, A. C.; Miller, D. W.; Elmquist, W. F. Transport of Fluorescein in MDCKII-MRP1 Transfected Cells and Mrp1-Knockout Mice. *Biochem. Biophys. Res. Commun.* **2001**, *284* (4), 863–869. https://doi.org/10.1006/bbrc.2001.5062.

(55) Iwashita, H.; Castillo, E.; Messina, M. S.; Swanson, R. A.; Chang, C. J. A Tandem Activity-Based Sensing and Labeling Strategy Enables Imaging of Transcellular Hydrogen Peroxide Signaling. *Proc. Natl. Acad. Sci. U.S.A.* **2021**, *118* (9), e2018513118. https://doi.org/10.1073/pnas.2018513118.

(56) Papot, S.; Tranoy, I.; Tillequin, F.; Florent, J. C.; Gesson, J. P. Design of Selectively Activated Anticancer Prodrugs: Elimination and Cyclization Strategies. *Current Medicinal Chemistry -Anti-Cancer Agents* **2002**, *2* (2), 155–185. https://doi.org/10.2174/1568011023354173.

(57) Huppertz, M.-C.; Wilhelm, J.; Grenier, V.; Schneider, M. W.; Falt, T.; Porzberg, N.; Hausmann, D.; Hoffmann, D. C.; Hai, L.; Tarnawski, M.; Pino, G.; Slanchev, K.; Kolb, I.; Acuna, C.; Fenk, L. M.; Baier, H.; Hiblot, J.; Johnsson, K. Recording Physiological History of Cells with Chemical Labeling. *Science* **2024**, *383* (6685), 890–897. https://doi.org/10.1126/science.adg0812.

(58) Broichhagen, J.; Trumpp, M.; Gatin-Fraudet, B.; Bruckmann, K.; Burdzinski, W.; Roßmann, K.; Levitz, J.; Knaus, P.; Jatzlau, J. Developing HaloTag and SNAP-Tag Chemical Inducers of Dimerization to

Probe Receptor Oligomerization and Downstream Signaling. *Angewandte Chemie International Edition* **2025**, *n/a* (n/a), e202506830. https://doi.org/10.1002/anie.202506830.
(59) Banala, S.; Arnold, A.; Johnsson, K. Caged Substrates for Protein Labeling and Immobilization. *ChemBioChem* **2008**, *9* (1), 38–41. https://doi.org/10.1002/cbic.200700472.
(60) Caldwell, S. E.; Demyan, I. R.; Falcone, G. N.; Parikh, A.; Lohmueller, J.; Deiters, A. Conditional Control of Benzylguanine Reaction with the Self-Labeling SNAP-Tag Protein. *Bioconjugate Chem.* **2025**, *36* (3), 540–548. https://doi.org/10.1021/acs.bioconjchem.5c00002.
(61) Kühn, S.; Nasufovic, V.; Wilhelm, J.; Kompa, J.; de Lange, E. M. F.; Lin, Y.-H.; Egoldt, C.; Fischer, J.; Lennoi, A.; Tarnawski, M.; Reinstein, J.; Vlijm, R.; Hiblot, J.; Johnsson, K. SNAP-Tag2 for Faster and Brighter Protein Labeling. *Nature Chemical Biology* **2025**. https://doi.org/10.1038/s41589-025-01942-z.
(62) Chen, X.; Wu, Y.-W. Tunable and Photoswitchable Chemically Induced Dimerization for Chemo-Optogenetic Control of Protein and Organelle Positioning. *Angew. Chem. Int. Ed.* **2018**, *57* (23), 6796–6799. https://doi.org/10.1002/anie.201800140.
(63) Aonbangkhen, C.; Zhang, H.; Wu, D. Z.; Lampson, M. A.; Chenoweth, D. M. Reversible Control of Protein Localization in Living Cells Using a Photocaged-Photocleavable Chemical Dimerizer. *J. Am. Chem. Soc.* **2018**, *140* (38), 11926–11930. https://doi.org/10.1021/jacs.8b07753.
(64) Zhang, H.; Aonbangkhen, C.; Tarasovetc, E. V.; Ballister, E. R.; Chenoweth, D. M.; Lampson, M. A. Optogenetic Control of Kinetochore Function. *Nat. Chem. Biol.* **2017**, *13* (10), 1096–1101. https://doi.org/10.1038/nchembio.2456.
(65) Chen, X.; Venkatachalapathy, M.; Kamps, D.; Weigel, S.; Kumar, R.; Orlich, M.; Garrecht, R.; Hirtz, M.; Niemeyer, C. M.; Wu, Y.-W.; Dehmelt, L. “Molecular Activity Painting”: Switch-like, Light-Controlled Perturbations inside Living Cells. *Angew. Chem. Int. Ed.* **2017**, *56* (21), 5916–5920. https://doi.org/10.1002/anie.201611432.
(66) Escudero, D. Revising Intramolecular Photoinduced Electron Transfer (PET) from First-Principles. *Acc. Chem. Res.* **2016**, *49* (9), 1816–1824. https://doi.org/10.1021/acs.accounts.6b00299.
(67) Würth, C.; Grabolle, M.; Pauli, J.; Spieles, M.; Resch-Genger, U. Relative and Absolute Determination of Fluorescence Quantum Yields of Transparent Samples. *Nat. Protoc.* **2013**, *8* (8), 1535–1550. https://doi.org/10.1038/nprot.2013.087.
(68) Zhang, X.-F.; Zhang, J.; Liu, L. Fluorescence Properties of Twenty Fluorescein Derivatives: Lifetime, Quantum Yield, Absorption and Emission Spectra. *J. Fluoresc.* **2014**, *24* (3), 819–826. https://doi.org/10.1007/s10895-014-1356-5.
(69) Dessolin, J.; Schuler, M.; Quinart, A.; De Giorgi, F.; Ghosez, L.; Ichas, F. Selective Targeting of Synthetic Antioxidants to Mitochondria: Towards a Mitochondrial Medicine for Neurodegenerative Diseases? *Eur. J. Pharmacol.* **2002**, *447* (2), 155–161. https://doi.org/10.1016/S0014-2999(02)01839-3.
(70) Huppertz, M.-C.; Wilhelm, J.; Grenier, V.; Schneider, M. W.; Falt, T.; Porzberg, N.; Hausmann, D.; Hoffmann, D. C.; Hai, L.; Tarnawski, M.; Pino, G.; Slanchev, K.; Kolb, I.; Acuna, C.; Fenk, L. M.; Baier, H.; Hiblot, J.; Johnsson, K. Recording Physiological History of Cells with Chemical Labeling. *Science* **2024**, *383* (6685), 890–897. https://doi.org/10.1126/science.adg0812.
(71) Critchfield, F. E.; Gibson, J. A. Jr.; Hall, J. L. Dielectric Constant for the Dioxane—Water System from 20 to 35°. *J. Am. Chem. Soc.* **1953**, *75* (8), 1991–1992. https://doi.org/10.1021/ja01104a506.
(72) Noordstra, I.; Liu, Q.; Nijenhuis, W.; Hua, S.; Jiang, K.; Baars, M.; Remmelzwaal, S.; Martin, M.; Kapitein, L. C.; Akhmanova, A. Control of Apico–Basal Epithelial Polarity by the Microtubule Minus-End-Binding Protein CAMSAP3 and Spectraplakin ACF7. *J. Cell Sci.* **2016**, *129* (22), 4278–4288. https://doi.org/10.1242/jcs.194878.
(73) Nijenhuis, W.; van Grinsven, M. M. P.; Kapitein, L. C. An Optimized Toolbox for the Optogenetic Control of Intracellular Transport. *J. Cell Biol.* **2020**, *219* (4), e201907149. https://doi.org/10.1083/jcb.201907149.
(74) Jansen, K. I.; Iwanski, M. K.; Burute, M.; Kapitein, L. C. A Live-Cell Marker to Visualize the Dynamics of Stable Microtubules throughout the Cell Cycle. *J. Cell Biol.* **2023**, *222* (5), e202106105. https://doi.org/10.1083/jcb.202106105.
(75) Pasolli, M.; Meiring, J. C. M.; Conboy, J. P.; Koenderink, G. H.; Akhmanova, A. Optogenetic and Chemical Genetic Tools for Rapid Repositioning of Vimentin Intermediate Filaments. *J. Cell Biol.* **2025**, *224* (9), e202504004. https://doi.org/10.1083/jcb.202504004.
(76) Meiring, J. C. M.; Grigoriev, I.; Nijenhuis, W.; Kapitein, L. C.; Akhmanova, A. Opto-Katanin, an Optogenetic Tool for Localized, Microtubule Disassembly. *Curr. Biol.* **2022**, *32* (21), 4660-4674.e6. https://doi.org/10.1016/j.cub.2022.09.010.
(77) Stepanova, T.; Slemmer, J.; Hoogenraad, C. C.; Lansbergen, G.; Dortland, B.; Zeeuw, C. I. D.; Grosveld, F.; Cappellen, G. van; Akhmanova, A.; Galjart, N. Visualization of Microtubule Growth in Cultured Neurons via the Use of EB3-GFP (End-Binding Protein 3-Green Fluorescent Protein). *J. Neurosci.* **2003**, *23* (7), 2655–2664. https://doi.org/10.1523/JNEUROSCI.23-07-02655.2003.
(78) Farías, G. G.; Fréal, A.; Tortosa, E.; Stucchi, R.; Pan, X.; Portegies, S.; Will, L.; Altelaar, M.; Hoogenraad, C. C. Feedback-Driven Mechanisms between Microtubules and the Endoplasmic Reticulum Instruct Neuronal Polarity. *Neuron* **2019**, *102* (1), 184-201.e8. https://doi.org/10.1016/j.neuron.2019.01.030.

(79) Shlevkov, E.; Kramer, T.; Schapansky, J.; LaVoie, M. J.; Schwarz, T. L. Miro Phosphorylation Sites Regulate Parkin Recruitment and Mitochondrial Motility. *Proc. Natl. Acad. Sci. U.S.A.* **2016**, *113* (41), E6097–E6106. https://doi.org/10.1073/pnas.1612283113.

(80) Harbauer, A. B.; Hees, J. T.; Wanderoy, S.; Segura, I.; Gibbs, W.; Cheng, Y.; Ordonez, M.; Cai, Z.; Cartoni, R.; Ashrafi, G.; Wang, C.; Perocchi, F.; He, Z.; Schwarz, T. L. Neuronal Mitochondria Transport *Pink1* mRNA via Synaptojanin 2 to Support Local Mitophagy. *Neuron* **2022**, *110* (9), 1516-1531.e9. https://doi.org/10.1016/j.neuron.2022.01.035.

(81) Hees, J. T.; Wanderoy, S.; Lindner, J.; Helms, M.; Murali Mahadevan, H.; Harbauer, A. B. Insulin Signalling Regulates Pink1 mRNA Localization via Modulation of AMPK Activity to Support PINK1 Function in Neurons. *Nat. Metab.* **2024**, *6* (3), 514–530. https://doi.org/10.1038/s42255-024-01007-w.

(82) Nguyen, H. P.; Stewart, S.; Kukwikila, M. N.; Jones, S. F.; Offenbartl-Stiegert, D.; Mao, S.; Balasubramanian, S.; Beck, S.; Howorka, S. A Photo-Responsive Small-Molecule Approach for the Opto-Epigenetic Modulation of DNA Methylation. *Angew. Chem. Int. Ed.* **2019**, *58* (20), 6620–6624. https://doi.org/10.1002/anie.201901139.

(83) Grimm, J. B.; Brown, T. A.; Tkachuk, A. N.; Lavis, L. D. General Synthetic Method for Si-Fluoresceins and Si-Rhodamines. *ACS Cent. Sci.* **2017**, *3* (9), 975–985. https://doi.org/10.1021/acscentsci.7b00247.

(84) Dwight, S. J.; Levin, S. Scalable Regioselective Synthesis of Rhodamine Dyes. *Org. Lett.* **2016**, *18* (20), 5316–5319. https://doi.org/10.1021/acs.orglett.6b02635.

(85) Zhou, Y.; Bi, T.; Wang, R.; Liang, P.; Lai, J.; Luo, Q.; Wang, H.; Shen, H.; Liu, Z.; Yang, S.; Ren, W. Optimization of the Procedure for Gram Scale Synthesis of Silicon Rhodamine and Its Application in Labeling BRD4 Kinase Inhibitor. *J. Anal. Test.* **2024**, *8* (4), 569–574. https://doi.org/10.1007/s41664-024-00309-y.

(86) Zhao, T.; Zhang, J.; Tang, M.; Ma, L. Z.; Lei, X. Development of an Effective Fluorescence Probe for Discovery of Aminopeptidase Inhibitors to Suppress Biofilm Formation. *J. Antibiot.* **2019**, *72* (6), 461–468. https://doi.org/10.1038/s41429-019-0166-z.

(87) Terai, T.; Kohno, M.; Boncompain, G.; Sugiyama, S.; Saito, N.; Fujikake, R.; Ueno, T.; Komatsu, T.; Hanaoka, K.; Okabe, T.; Urano, Y.; Perez, F.; Nagano, T. Artificial Ligands of Streptavidin (ALiS): Discovery, Characterization, and Application for Reversible Control of Intracellular Protein Transport. *J. Am. Chem. Soc.* **2015**, *137* (33), 10464–10467. https://doi.org/10.1021/jacs.5b05672.

(88) Singh, V.; Wang, S.; Kool, E. T. Genetically Encoded Multispectral Labeling of Proteins with Polyfluorophores on a DNA Backbone. *J. Am. Chem. Soc.* **2013**, *135* (16), 6184–6191. https://doi.org/10.1021/ja4004393.

(89) Cong, M.; Corona, C.; Mcdougall, M. G.; Zimprich, C. pH Sensors. US9702824B2, July 11, 2017.

(90) Taj, R.; Sorensen, J. L. Synthesis of Actinomycetes Natural Products JBIR-94, JBIR-125, and Related Analogues. *Tetrahedron Lett.* **2015**, *56* (51), 7108–7111. https://doi.org/10.1016/j.tetlet.2015.11.020.

(91) Ayala, C. E.; Villalpando, A.; Nguyen, A. L.; McCandless, G. T.; Kartika, R. Chlorination of Aliphatic Primary Alcohols via Triphosgene–Triethylamine Activation. *Org. Lett.* **2012**, *14* (14), 3676–3679. https://doi.org/10.1021/ol301520d.

(92) Nakagawa, Y.; Irie, K.; Masuda, A.; Ohigashi, H. Synthesis, Conformation and PKC Isozyme Surrogate Binding of New Lactone Analogues of Benzolactam-V8s. *Tetrahedron* **2002**, *58* (11), 2101–2115. https://doi.org/10.1016/S0040-4020(02)00099-6.

### 12 NMR spectra

| | |
|---|---|
| Compound 3 | 75 |
| Compound 4 | 76 |
| CHalo-Boc | 77 |
| Cou-CHalo-Boc | 78 |
| CHalo-SiR | 80 |
| Compound 8 | 82 |
| Bn-CHalo-SiR | 83 |
| Cou-CHalo-SiR | 84 |
| Compound 11 | 86 |
| Leu-CHalo-SiR | 88 |
| Compound 72 | 89 |
| Compound 73 | 89 |
| Compound 76 | 90 |
| Compound 77 | 91 |
| Compound 14 | 92 |
| Compound 17 | 94 |
| Cou-CHalo-BG | 95 |
| Compound 18 | 96 |
| Compound 20 | 97 |
| CHalo-BG | 98 |
| Compound 22 | 99 |
| Compound 24 | 101 |
| Cou-CHalo-SiR-BG | 103 |
| Compound 26 | 105 |
| Compound 30 | 106 |
| CA-Fluo | 107 |
| Compound 33 | 108 |
| Compound 35 | 109 |
| Compound 38 | 110 |
| $HL2^{O}$-Fluo | 111 |
| Compound 40 | 113 |
| Compound 41 | 114 |
| Compound 42 | 115 |
| Compound 44 | 116 |
| Compound 48 | 117 |
| $HL2^{N}$-Fluo | 118 |
| Compound 49 | 119 |
| Cou-$HL2^{N}$-Fluo | 120 |
| Compound 56 | 122 |
| Compound 60 | 123 |
| HL3-Fluo | 124 |
| Cou-HL3-Fluo | 124 |
| Compound 64 | 125 |
| Compound 65 | 127 |
| Compound 66 | 128 |
| Compound 67 | 129 |
| CHalo-Fluo | 130 |
| Cou-CHalo-Fluo | 131 |

**Compound 3**

**$^1$H-NMR**

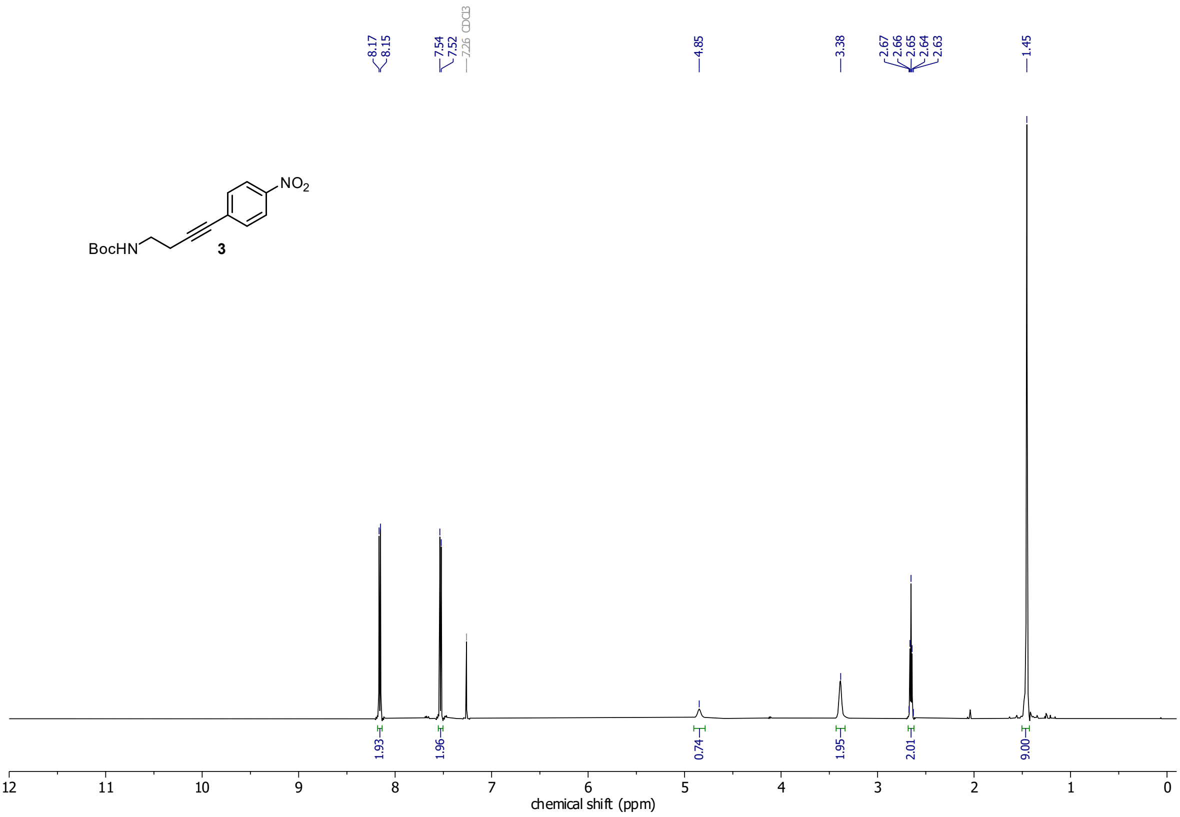

**$^{13}$C-NMR**

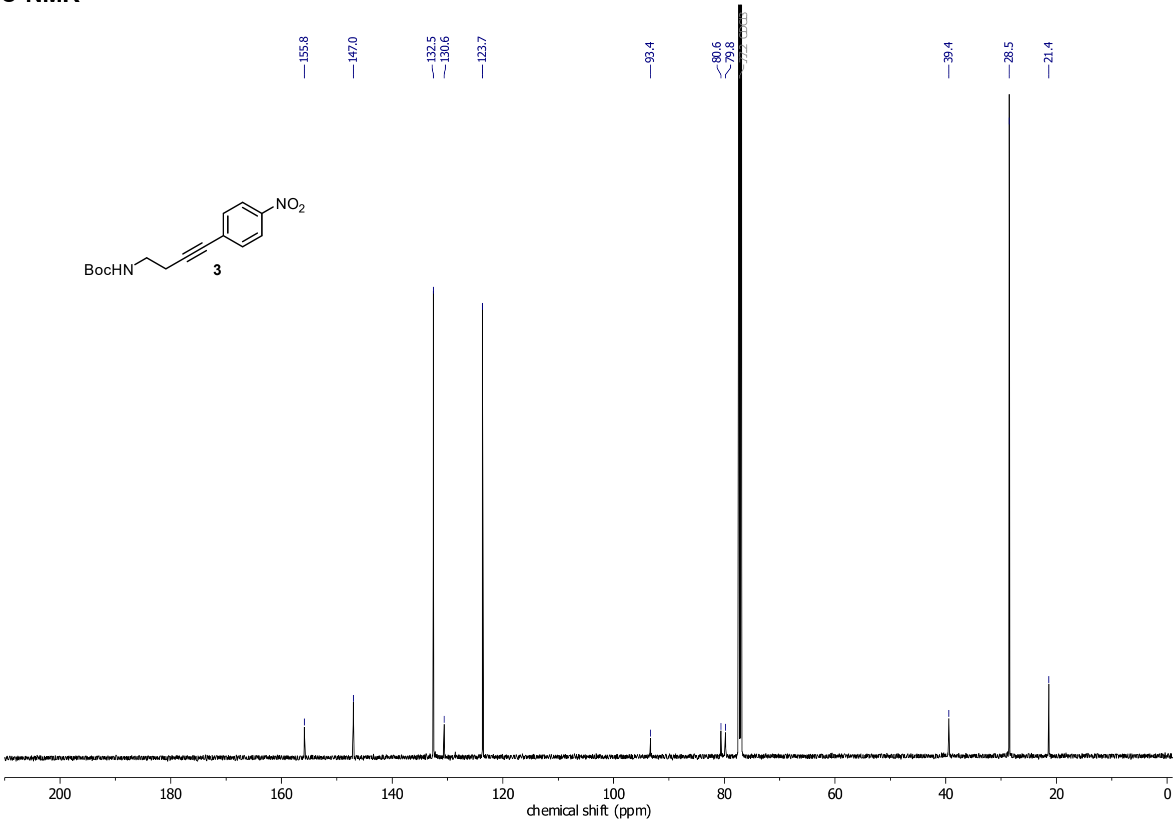

#### Compound 4

##### $^{1}$H-NMR

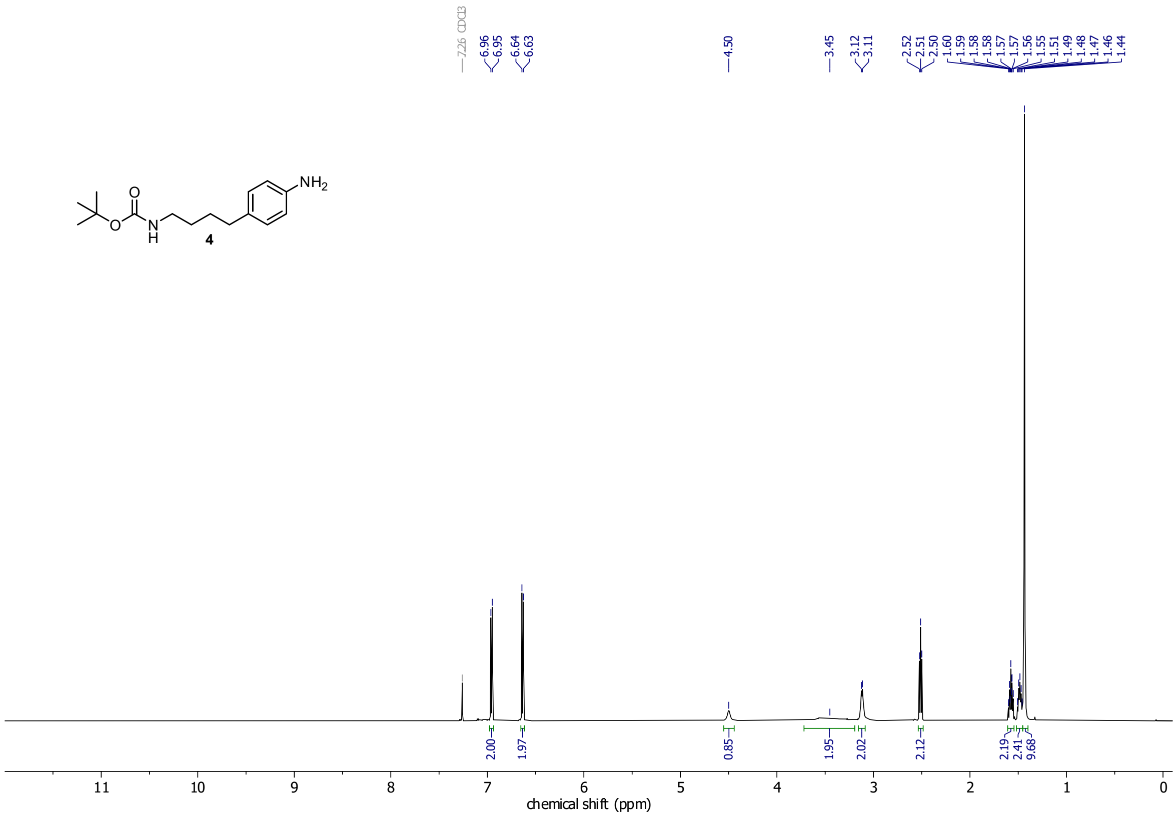

##### $^{13}$C-NMR

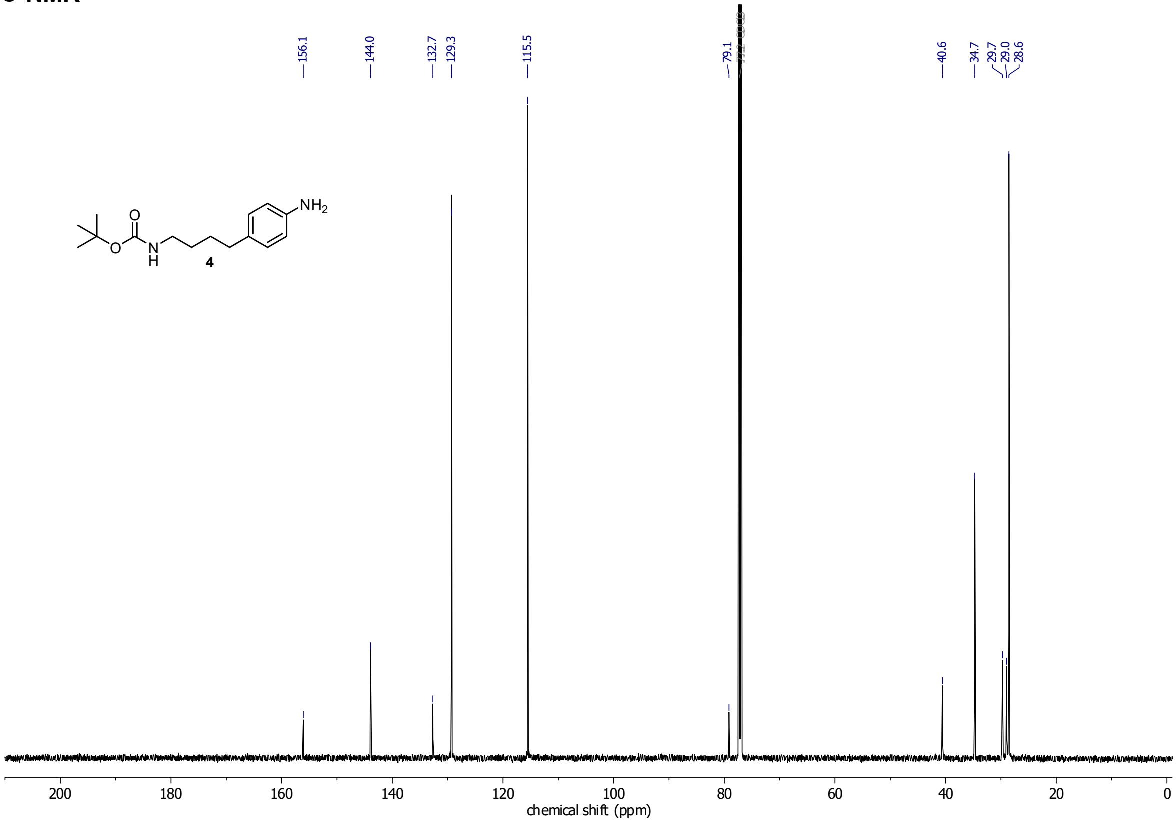

**CHalo-Boc**

**$^1$H-NMR**

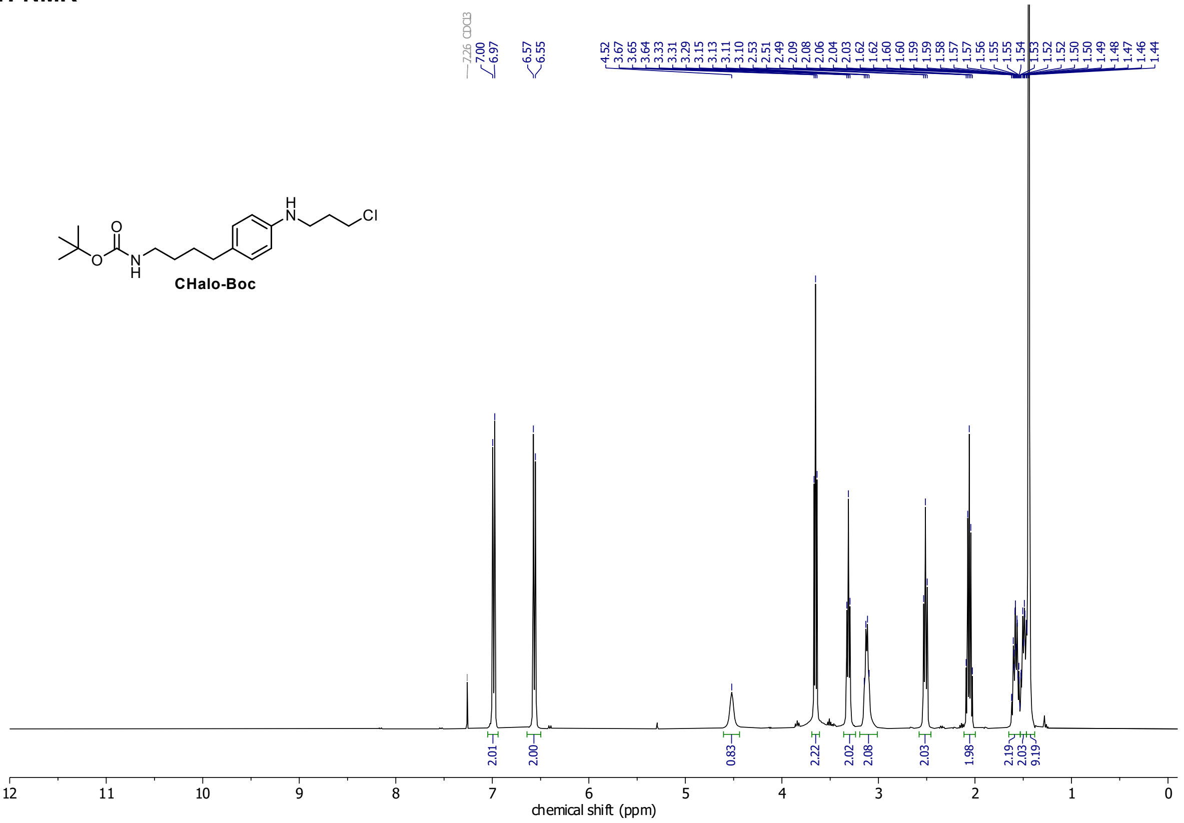

**$^{13}$C-NMR**

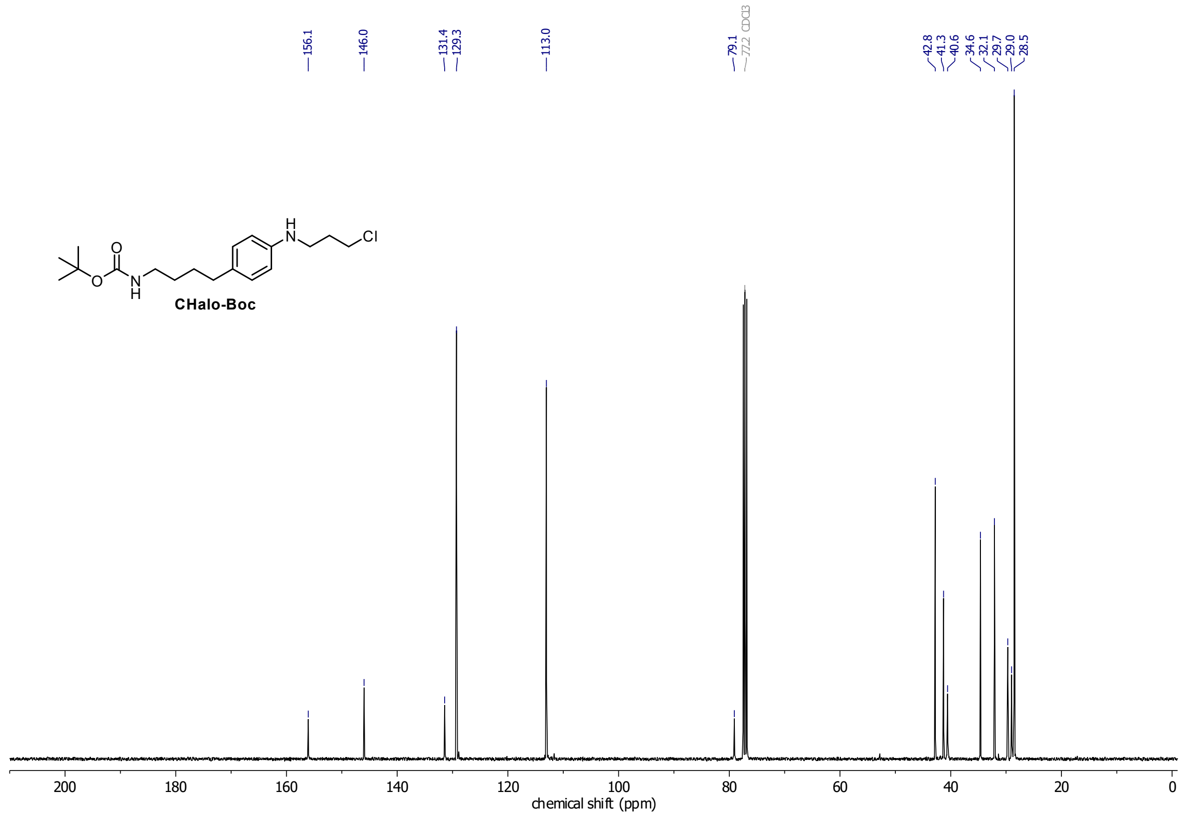

**Cou-CHalo-Boc**

**$^{1}$H-NMR**

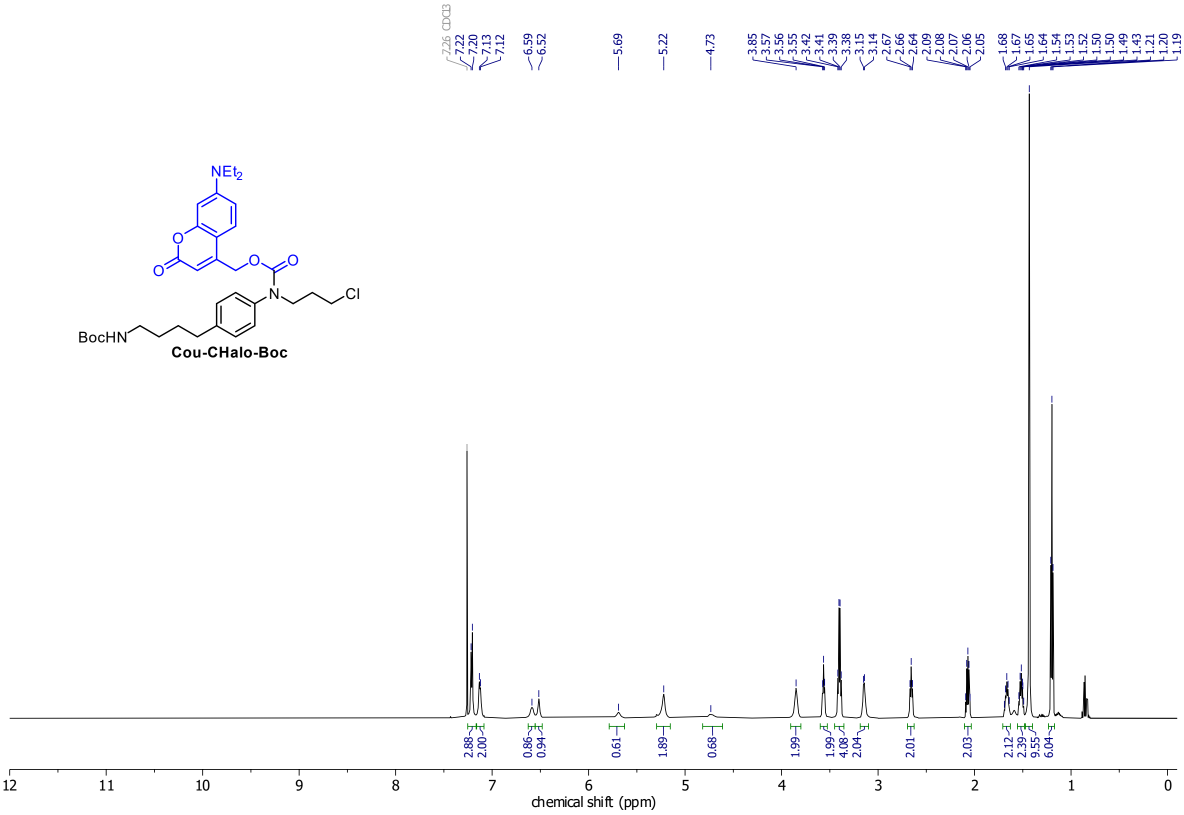

**$^{13}$C-NMR**

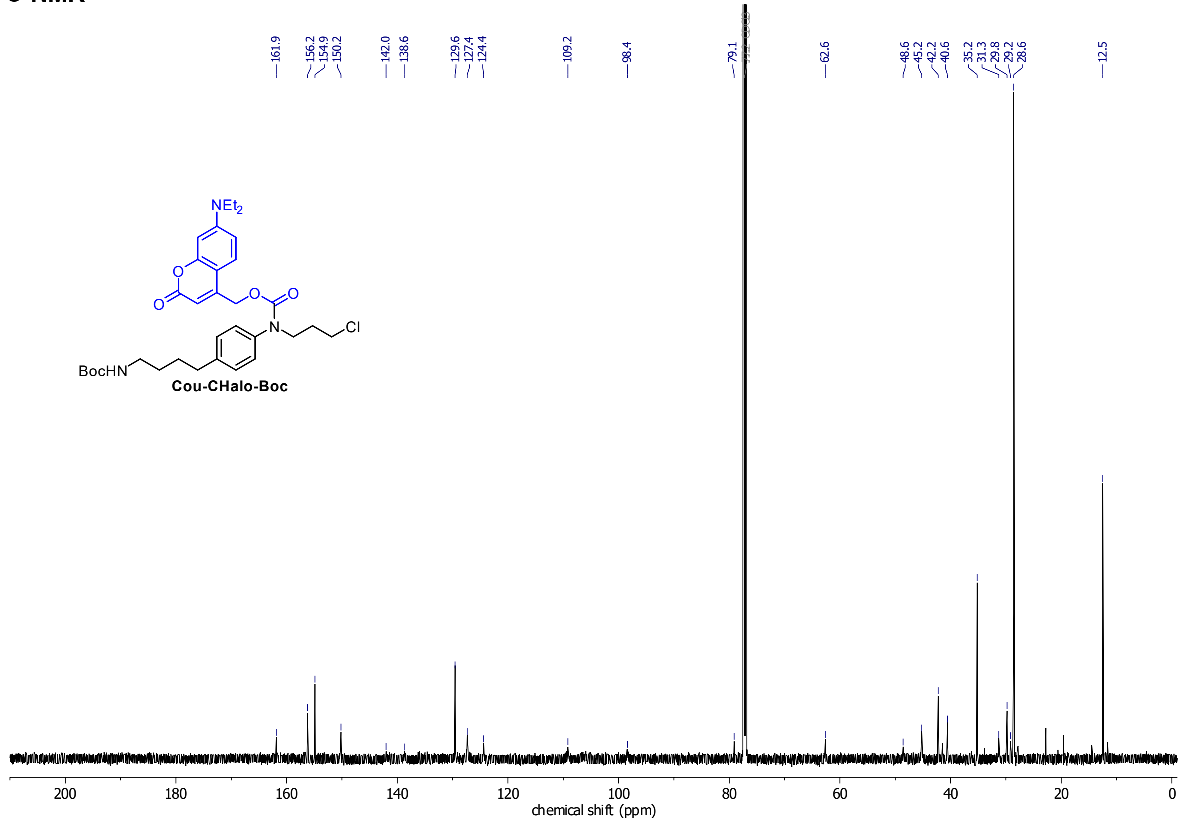

**HSQC**

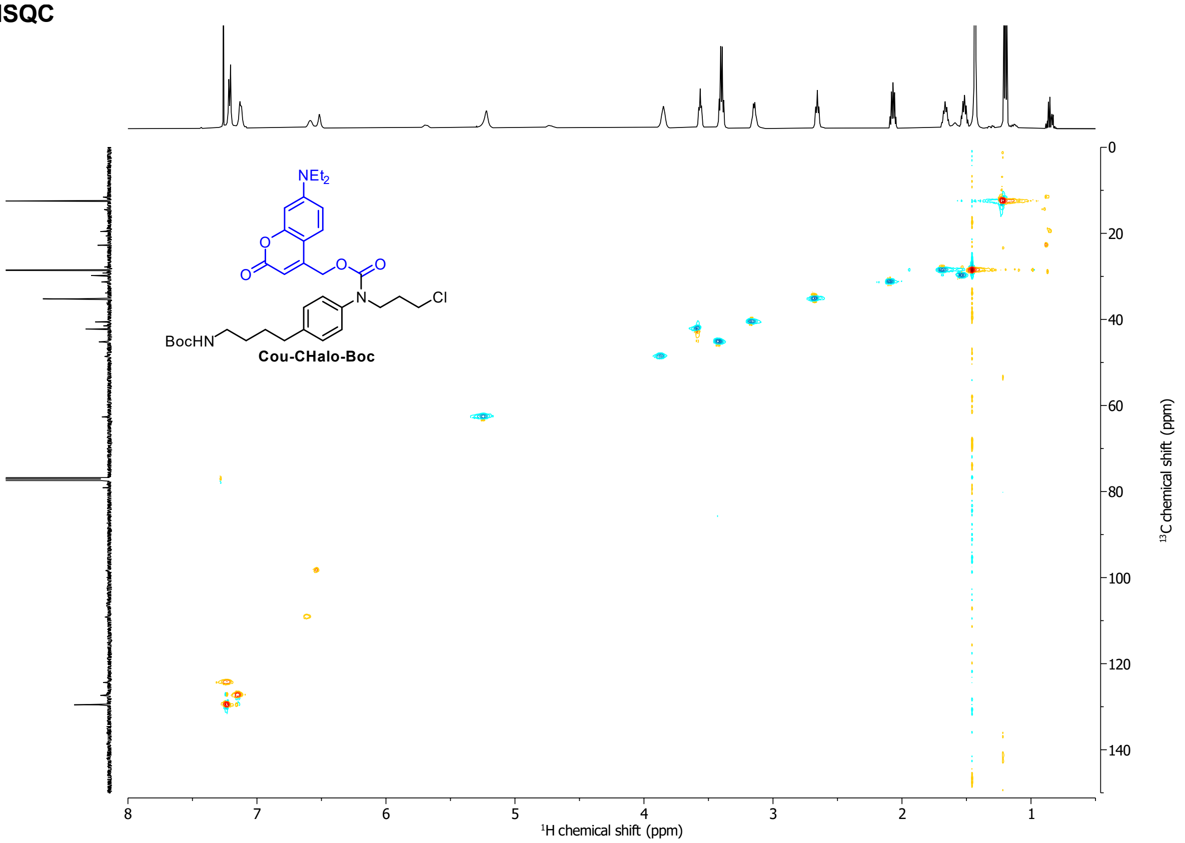
NEt2
O
O
O
O
N
Cl
BocHN
Cou-CHalo-Boc
0
20
40
60
80
100
120
140
13C chemical shift (ppm)
8
7
6
5
4
3
2
1
1H chemical shift (ppm)

#### CHalo-SiR

##### 1H-NMR

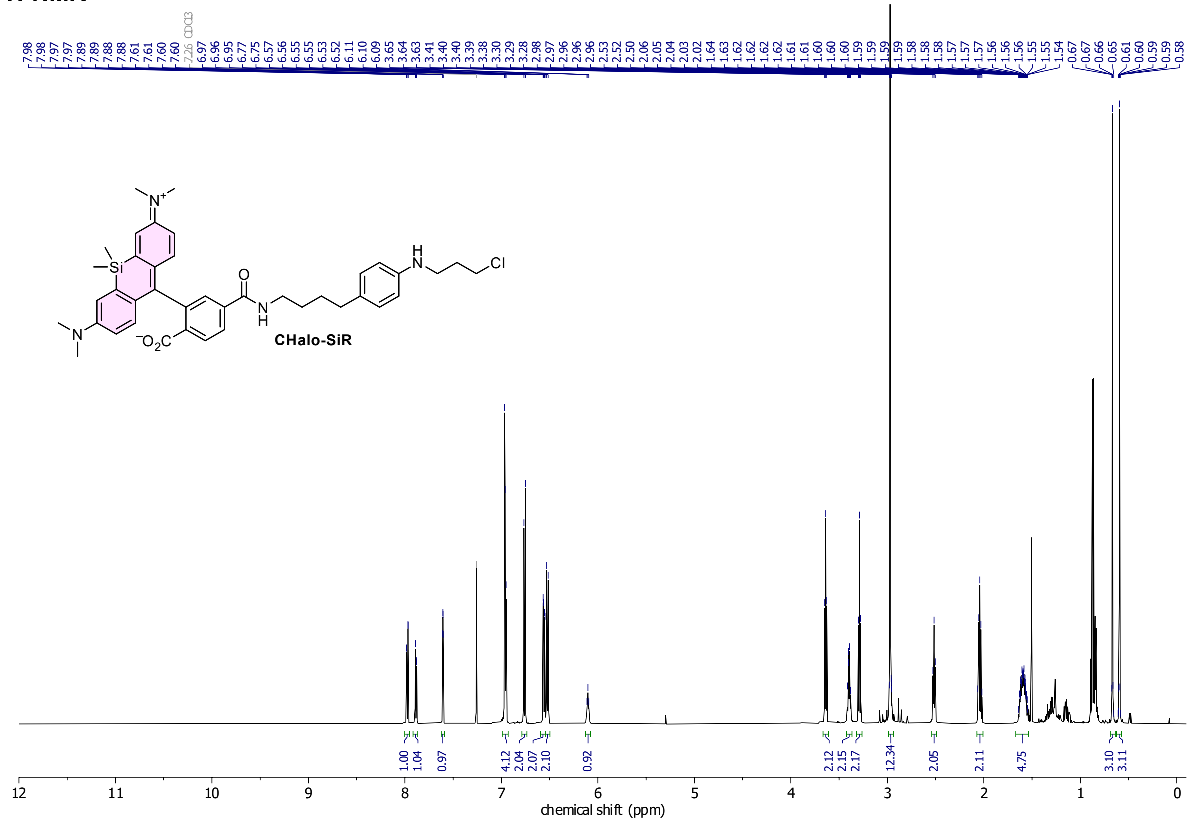

##### 13C-NMR

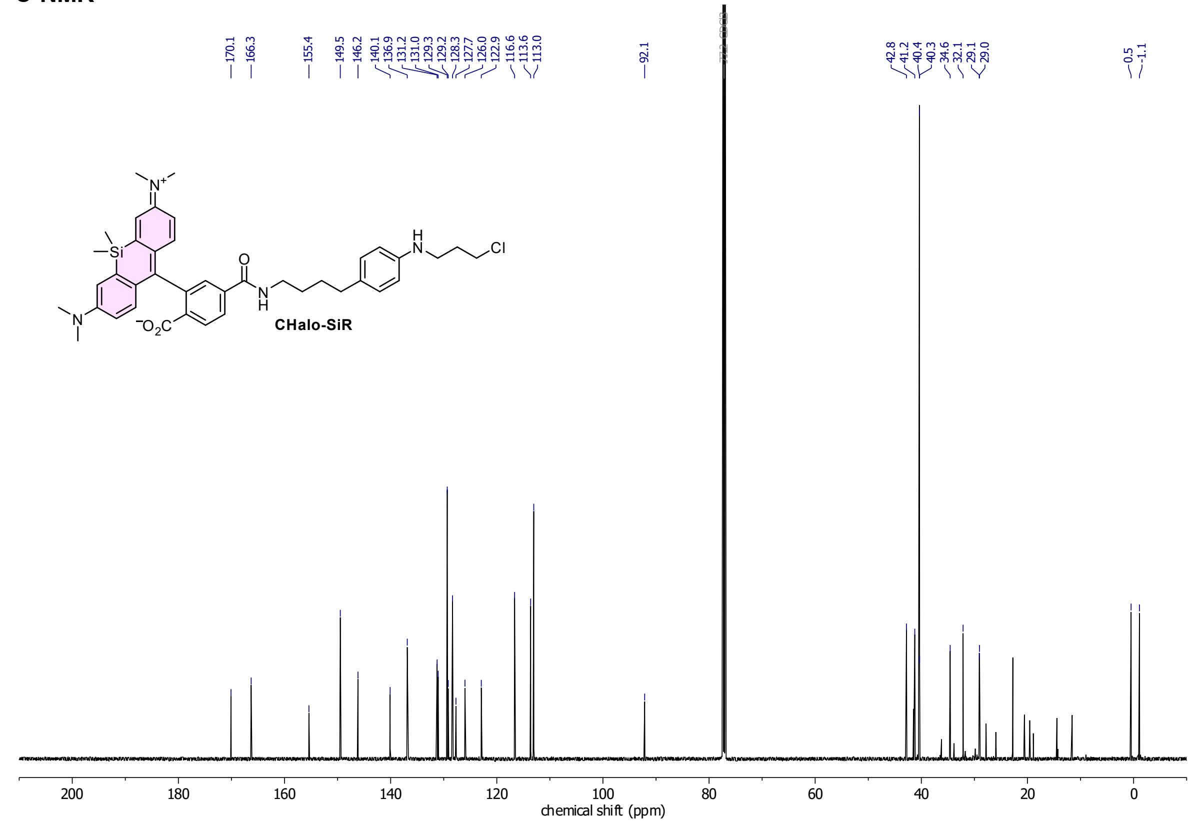

*minor H-grease impurities

#### HSQC

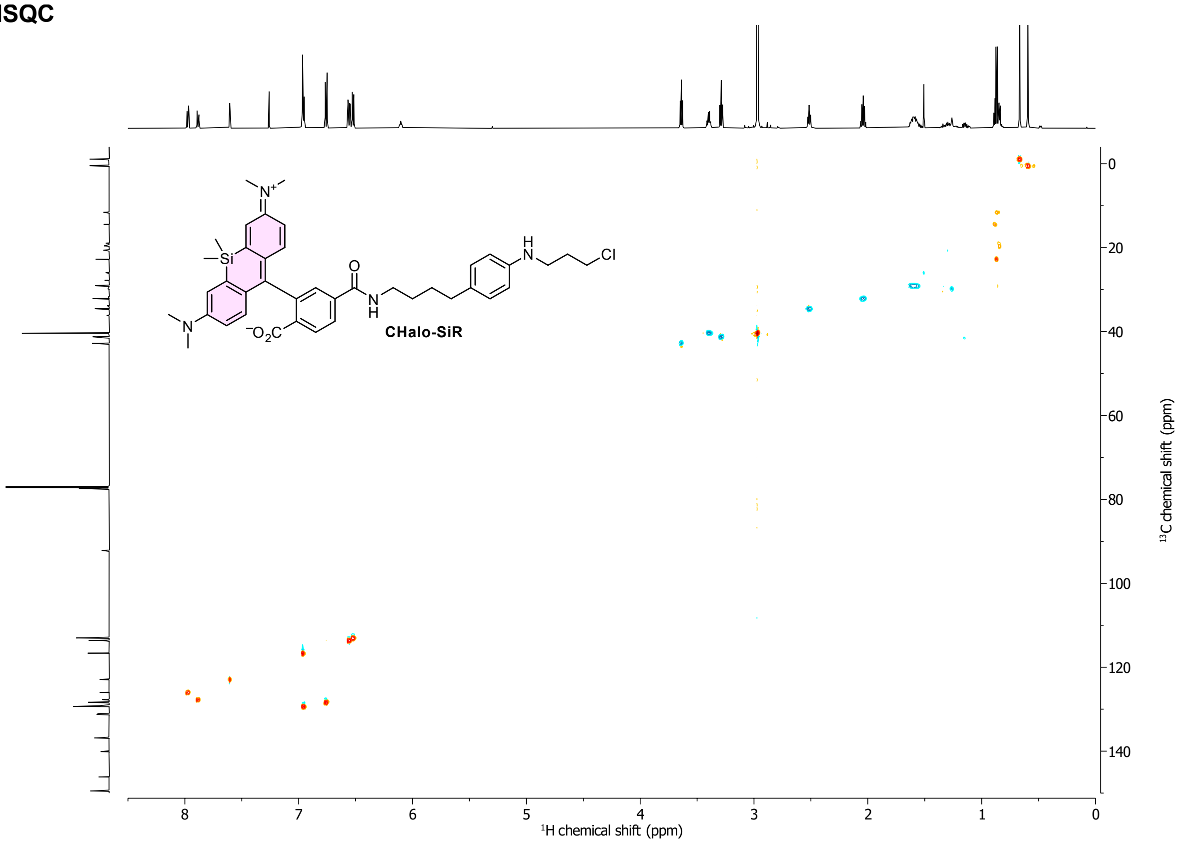

CHalo-SiR
$^{1}$H chemical shift (ppm)
$^{13}$C chemical shift (ppm)

#### HMBC

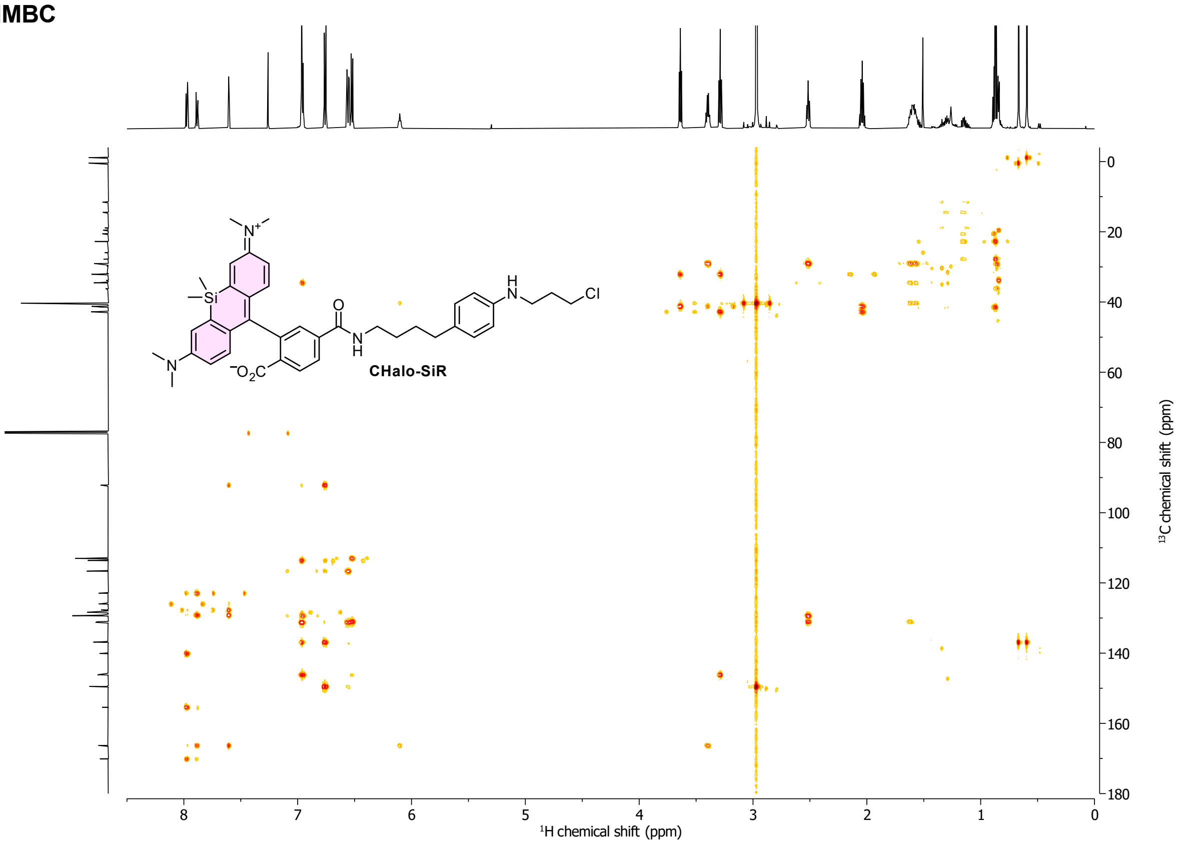

CHalo-SiR
$^{1}$H chemical shift (ppm)
$^{13}$C chemical shift (ppm)

#### Compound 8

##### $^{1}$H-NMR

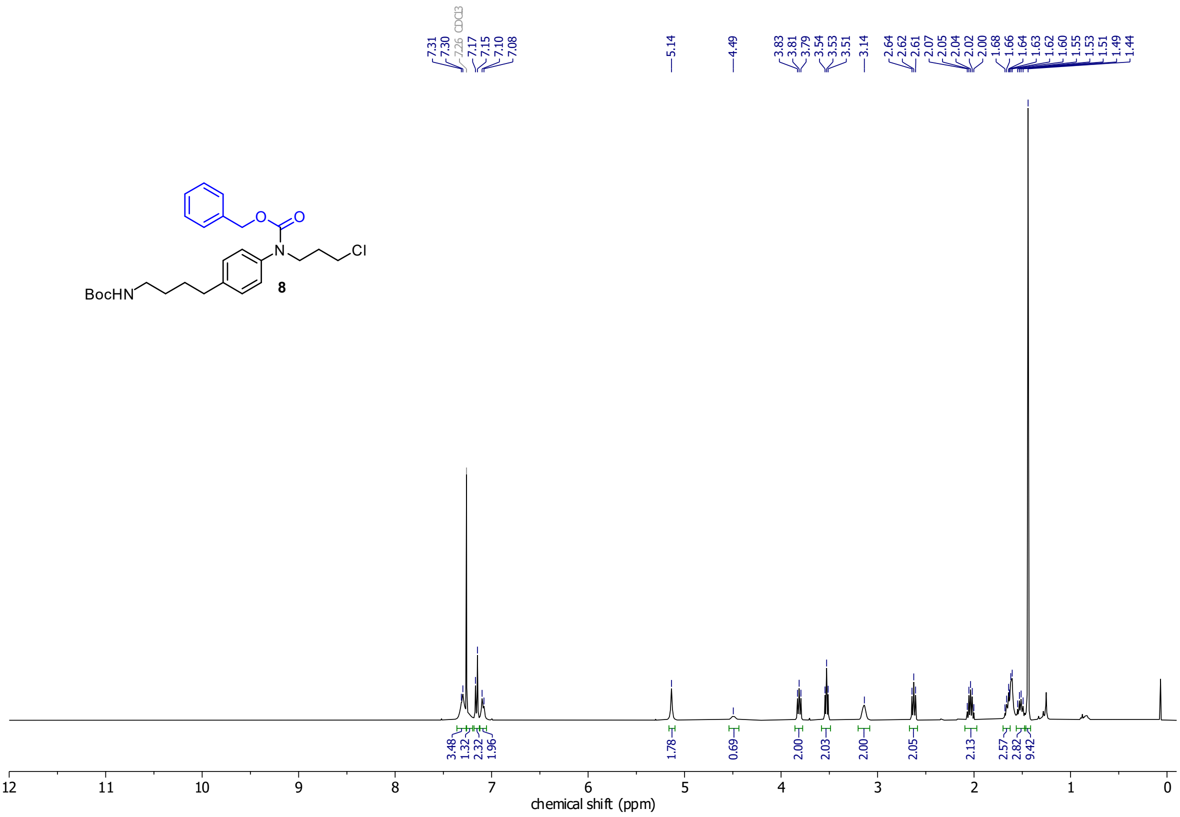

##### $^{13}$C-NMR

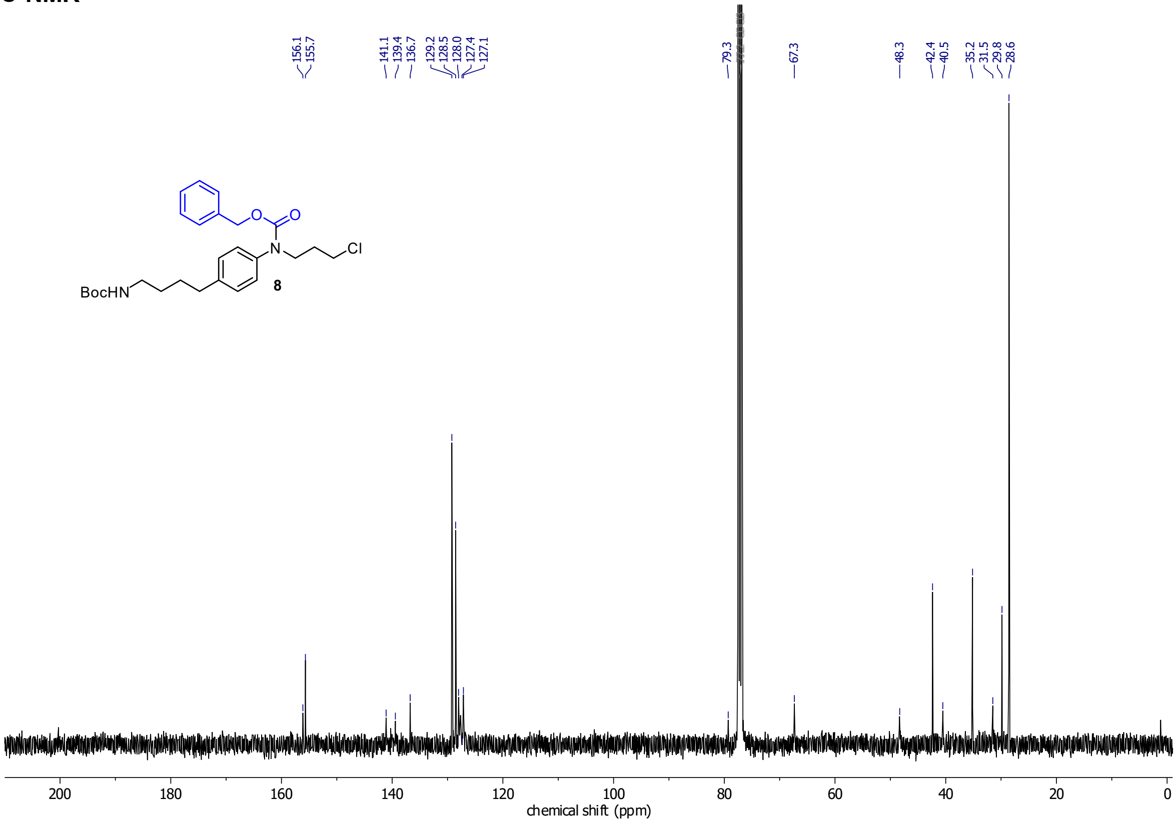

#### Bn-CHalo-SiR

##### $^{1}$H-NMR

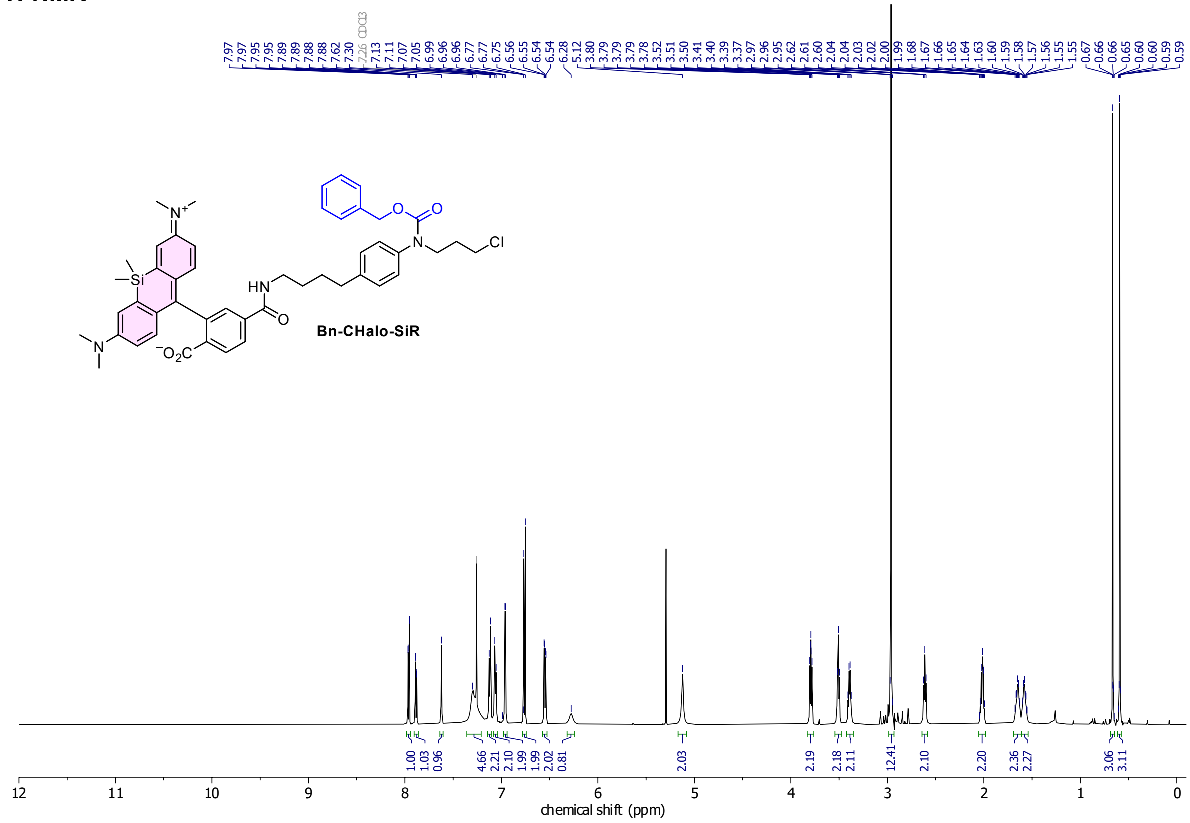

##### $^{13}$C-NMR

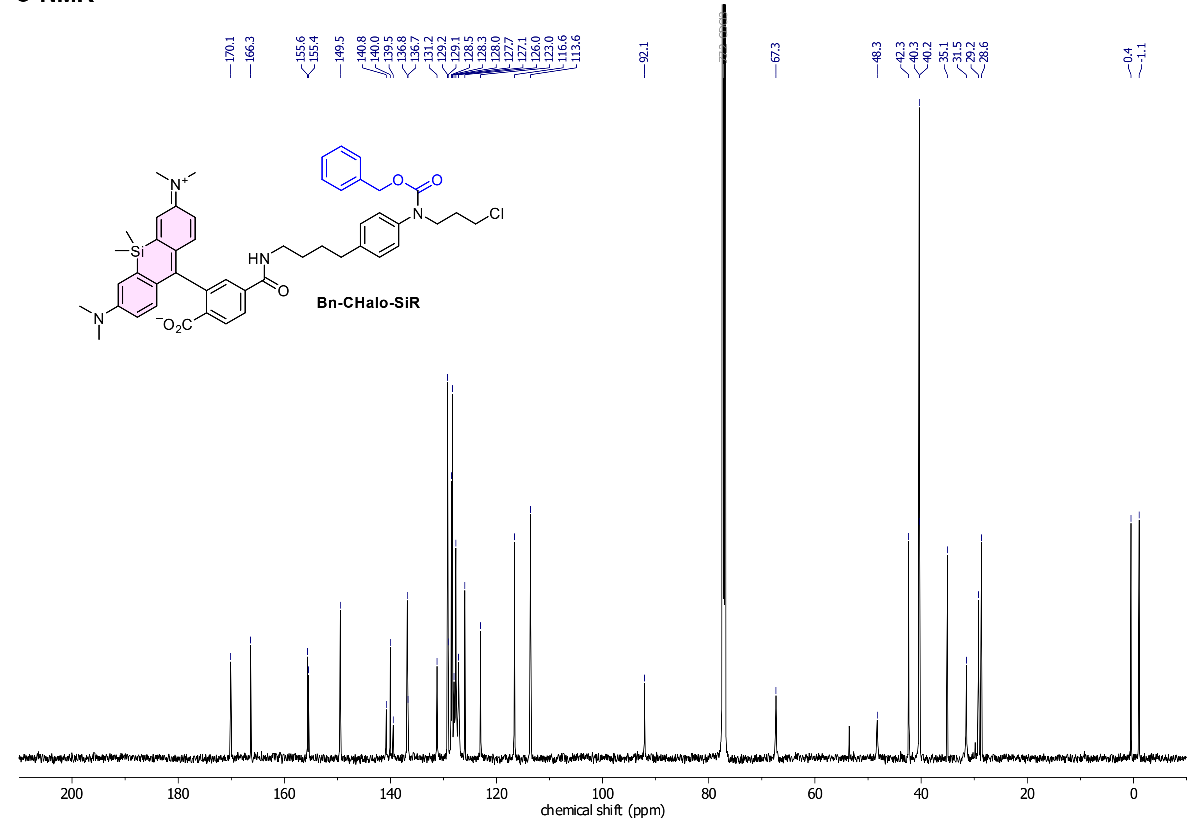

**Cou-CHalo-SiR**

**$^{1}$H-NMR**

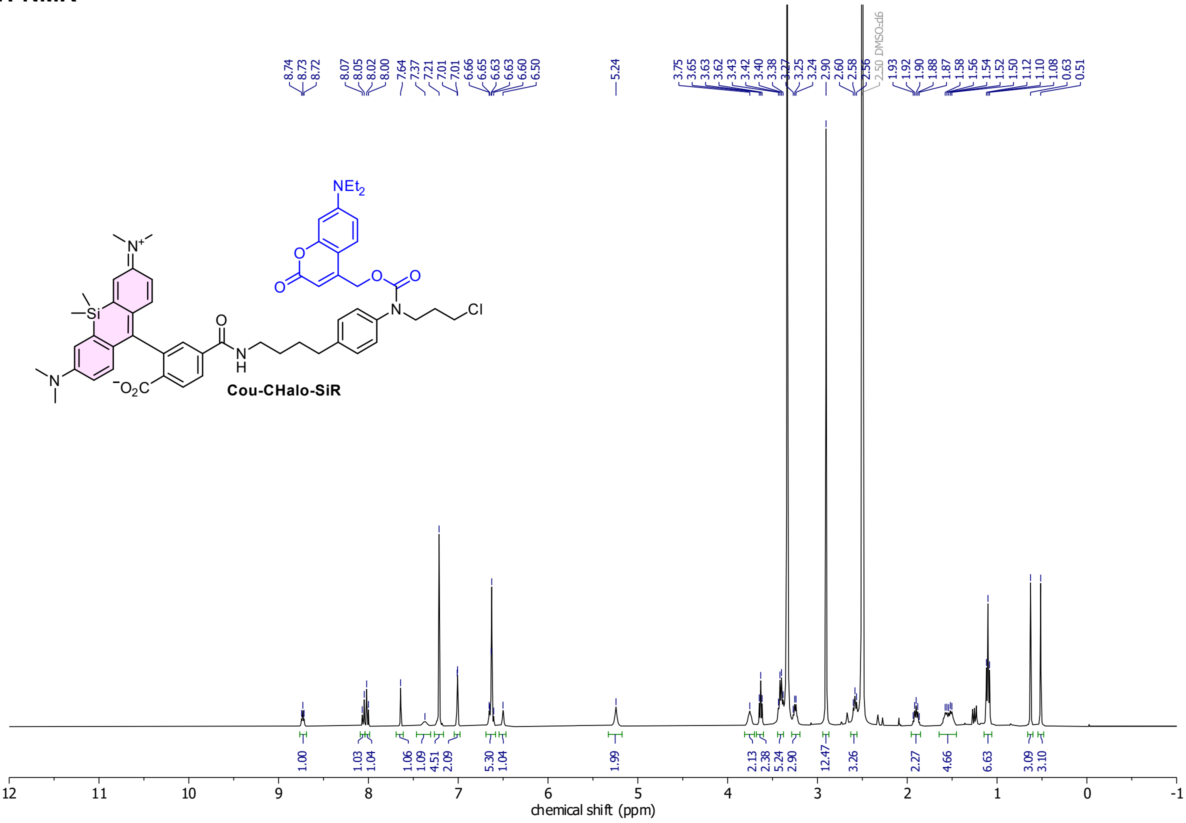

**$^{13}$C-NMR**

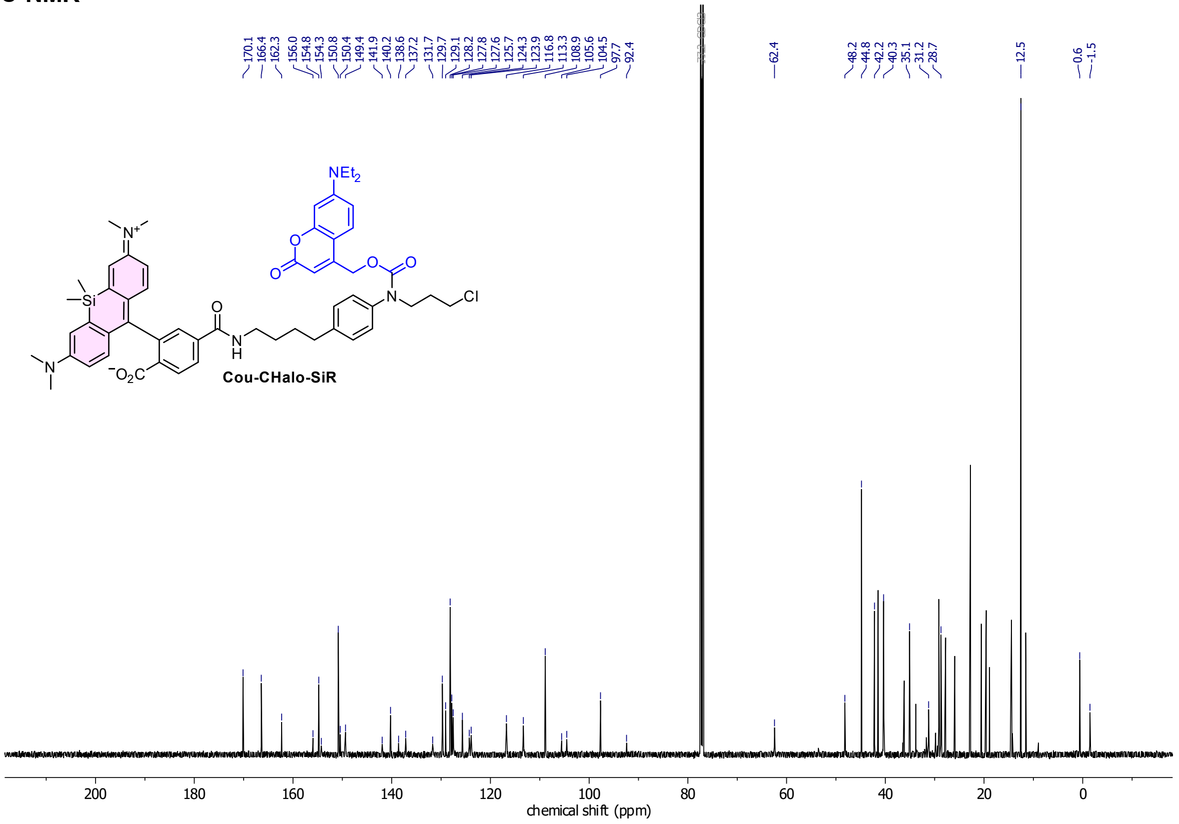

**minor H-grease impurities in $^{13}$C-spectrum*

#### HSQC

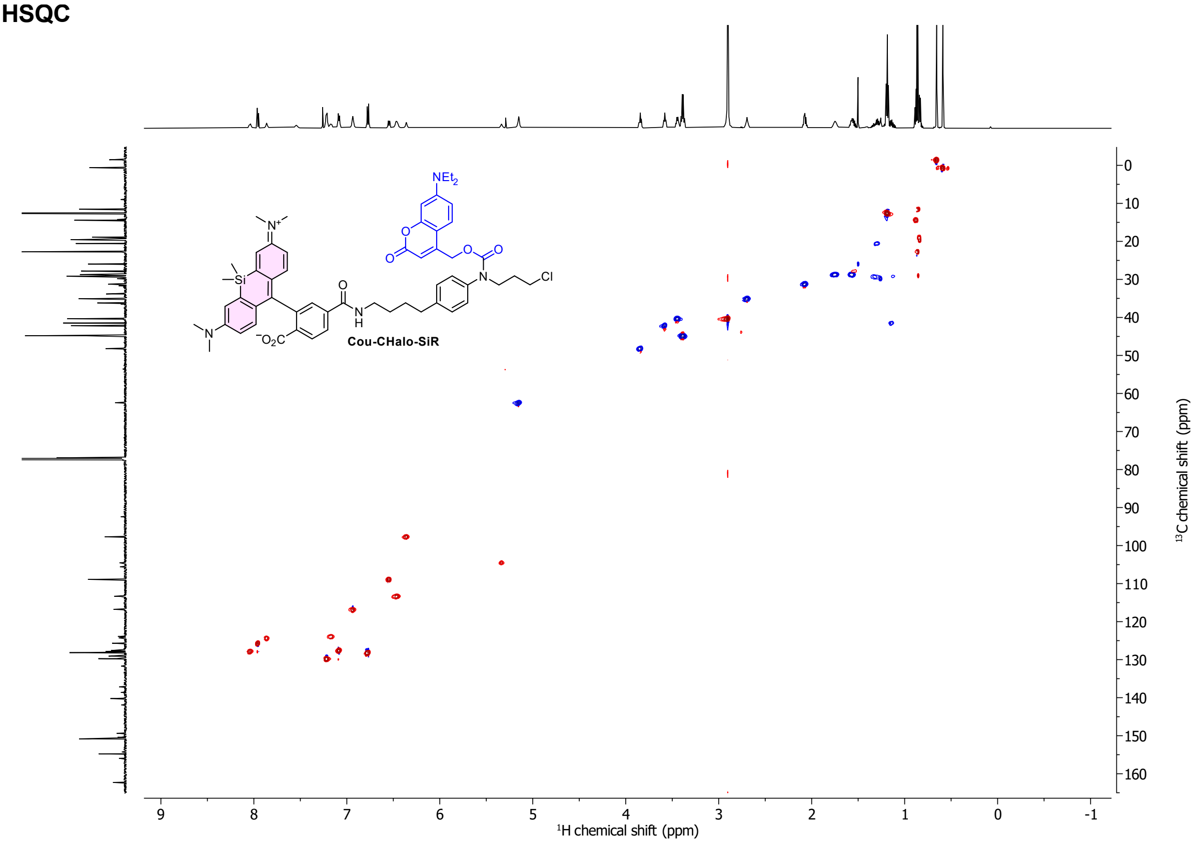

NEt2
Cl
Cou-CHalo-SiR
¹H chemical shift (ppm)
¹³C chemical shift (ppm)
9
8
7
6
5
4
3
2
1
0
-1
0
10
20
30
40
50
60
70
80
90
100
110
120
130
140
150
160

#### HMBC

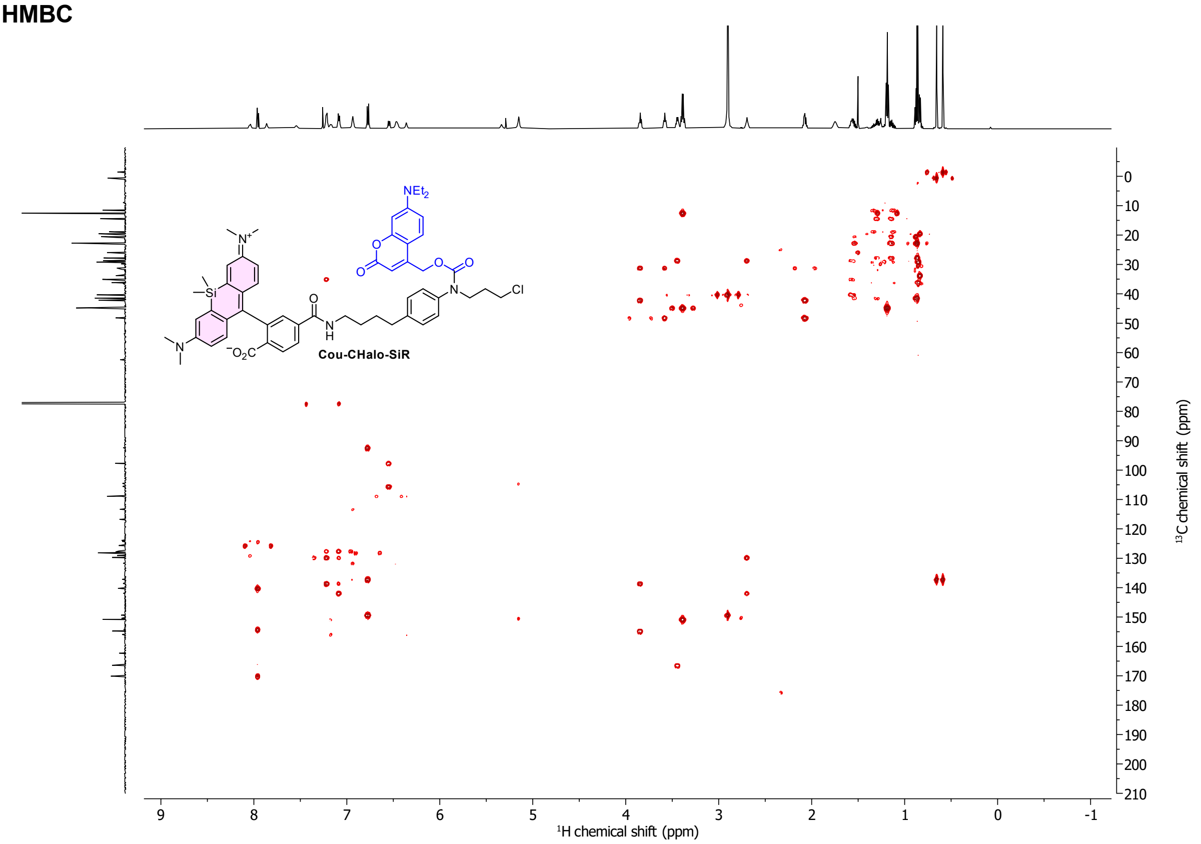

NEt2
Cl
Cou-CHalo-SiR
¹H chemical shift (ppm)
¹³C chemical shift (ppm)
9
8
7
6
5
4
3
2
1
0
-1
0
10
20
30
40
50
60
70
80
90
100
110
120
130
140
150
160
170
180
190
200
210

**Compound 11**

**$^{1}$H-NMR**

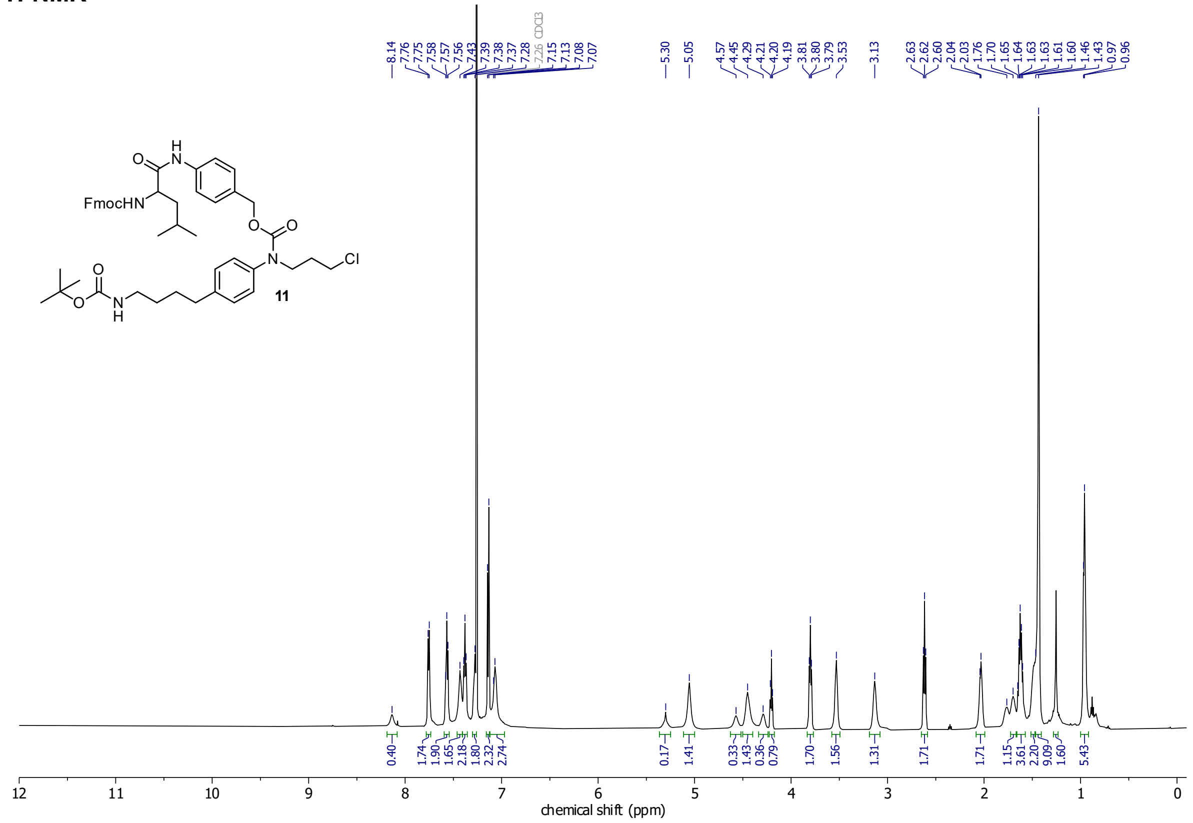

**$^{13}$C-NMR**

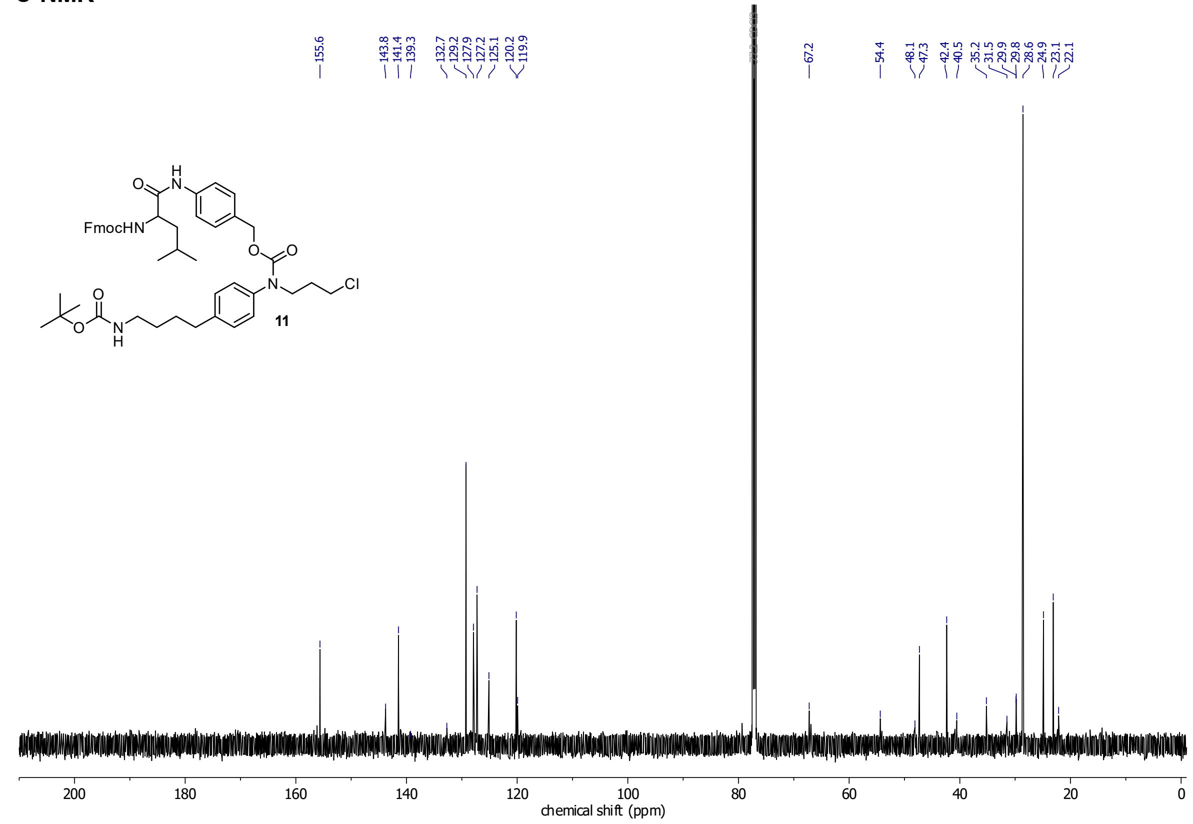

**HSQC**

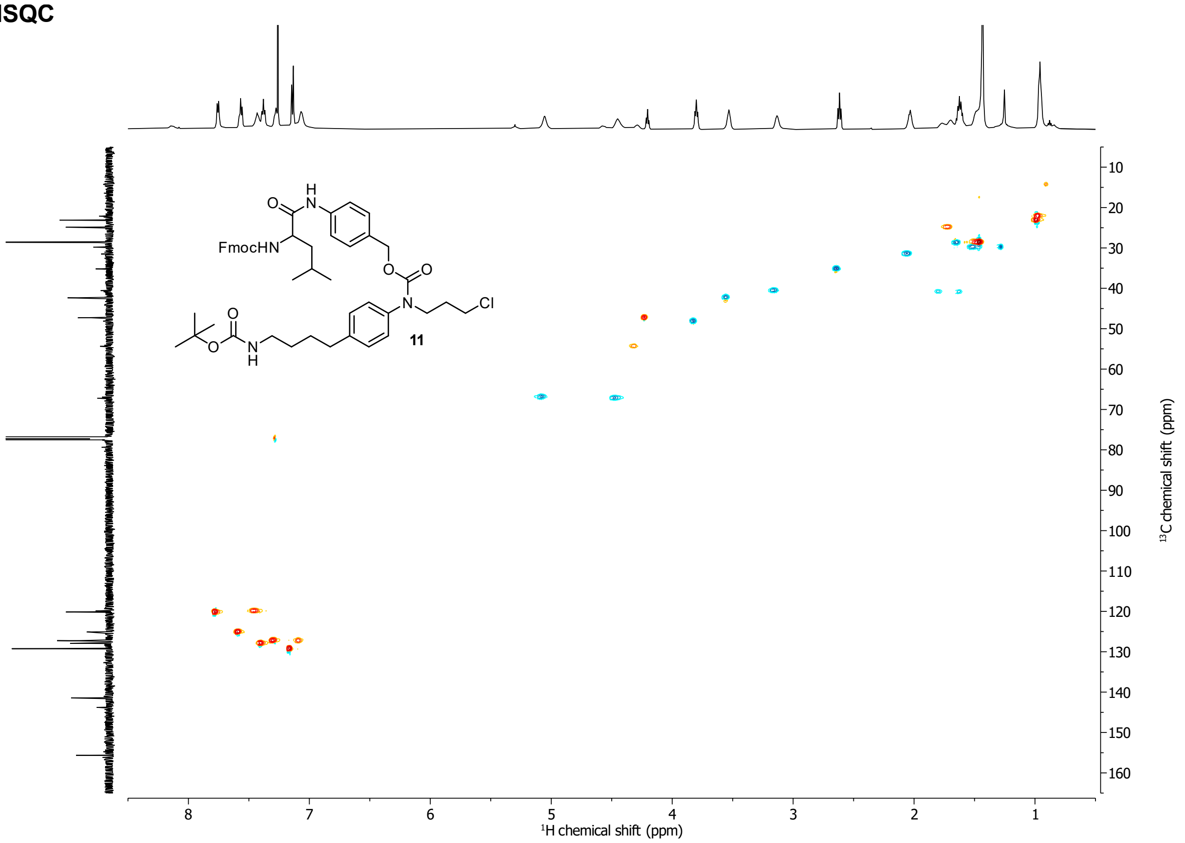

FmocHN
11
Cl
¹H chemical shift (ppm)
¹³C chemical shift (ppm)

**HMBC**

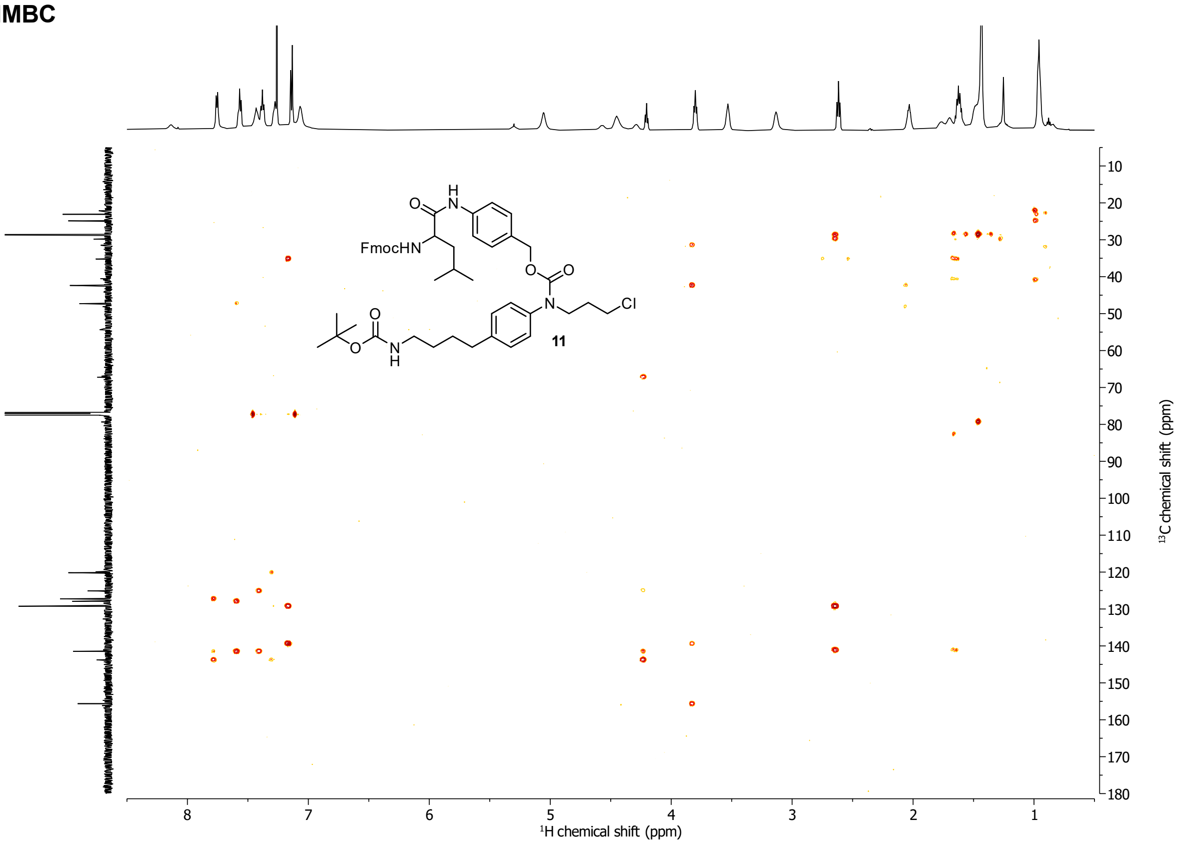

FmocHN
11
Cl
¹H chemical shift (ppm)
¹³C chemical shift (ppm)

**Leu-CHalo-SiR**
**$^1$H-NMR**

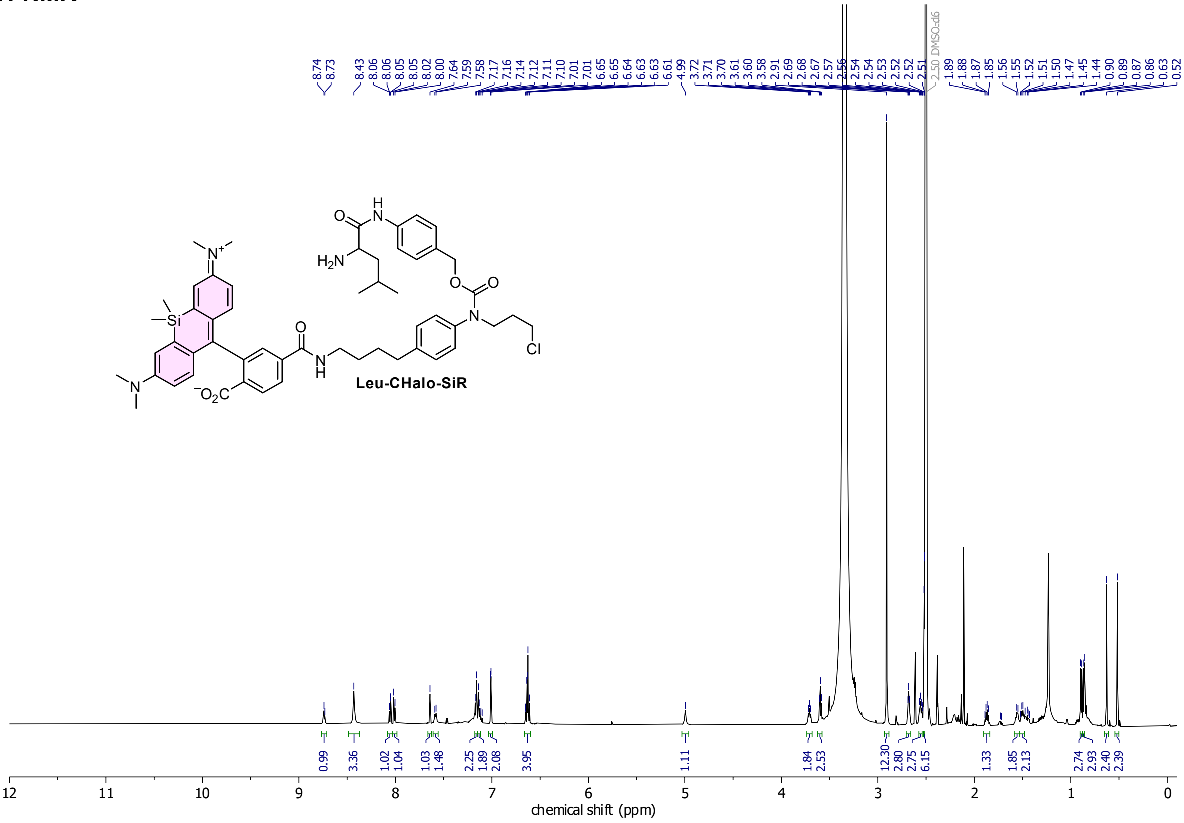

#### Compound 72
**$^1$H-NMR**

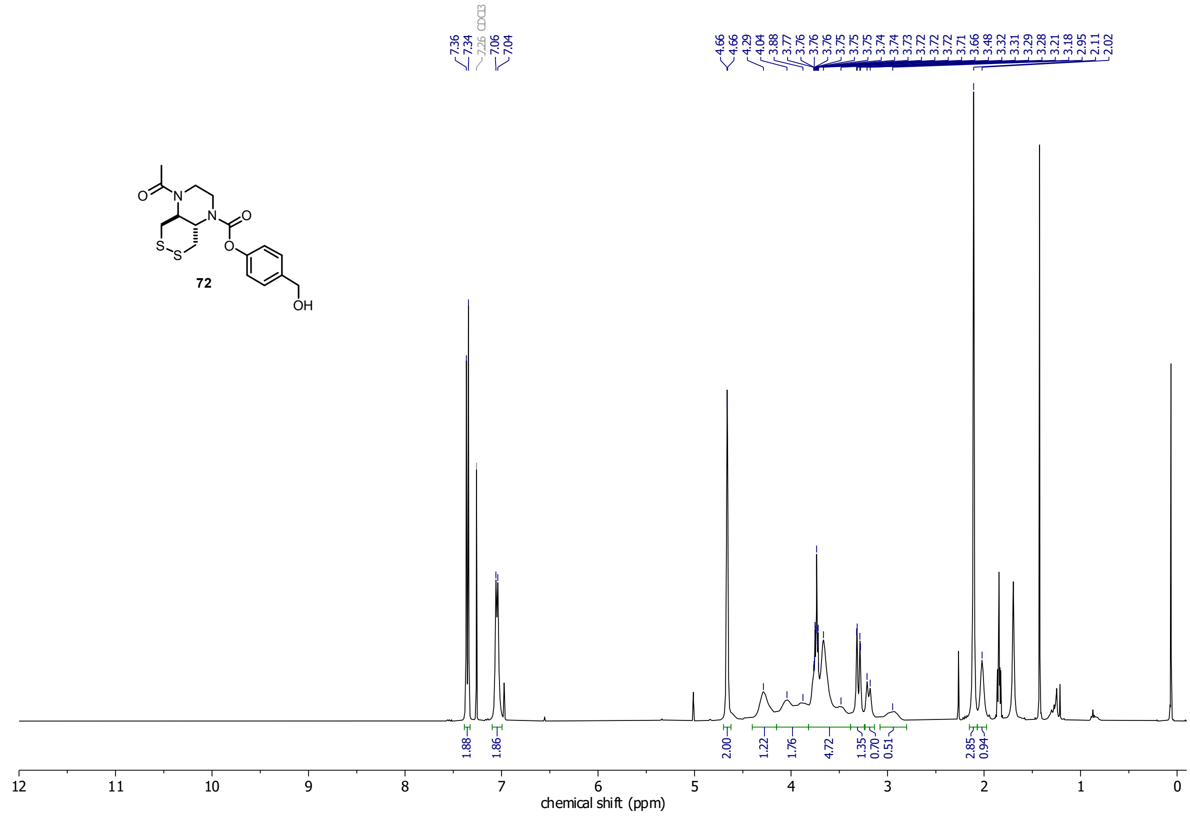

*minor silicon grease impurity*

#### Compound 73
**$^1$H-NMR**

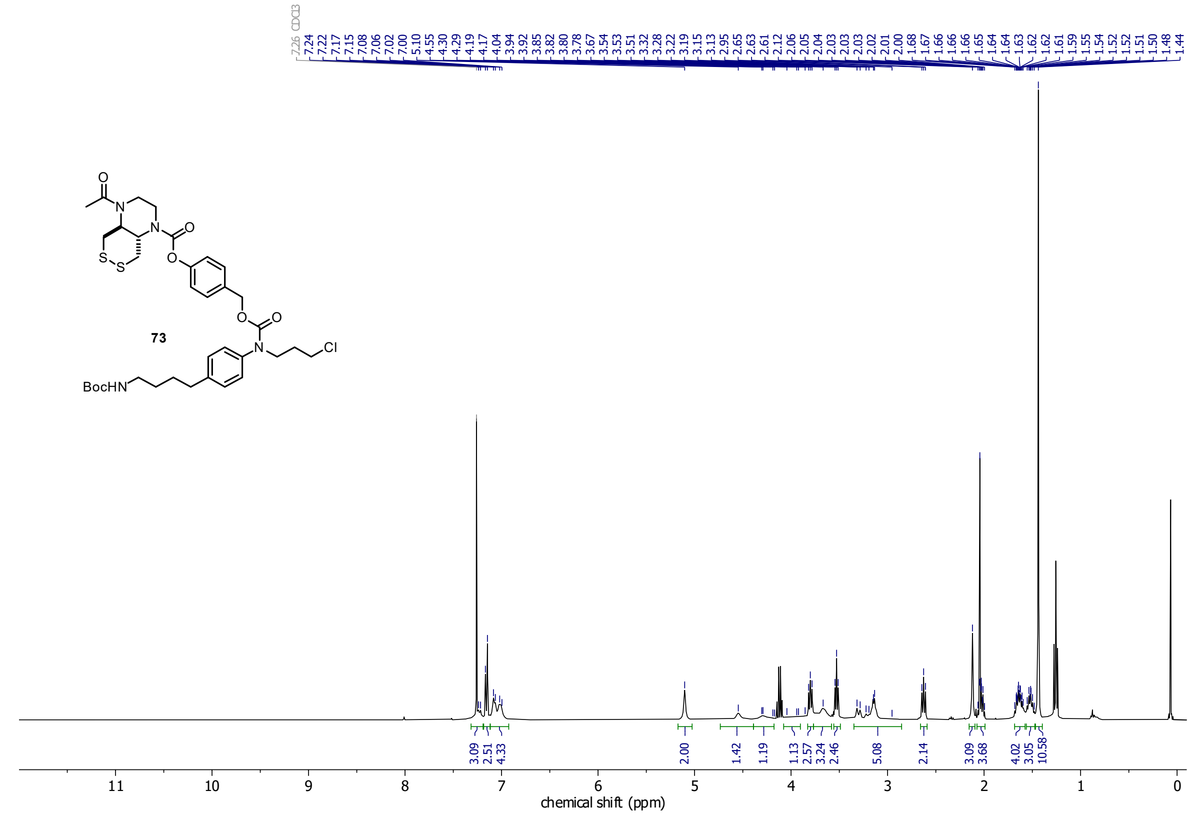

*ethyl acetate and silicon grease impurities*

#### Compound 76

##### $^1$H-NMR

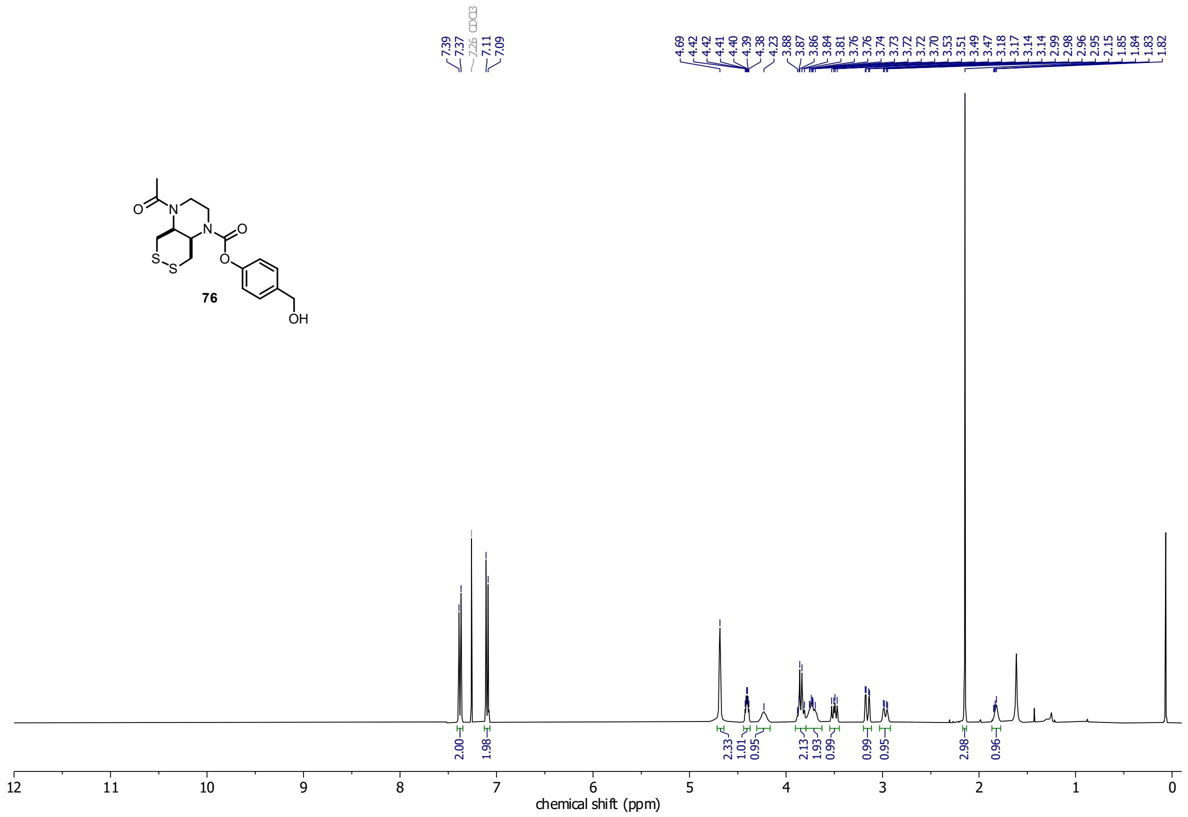

##### $^{13}$C-NMR

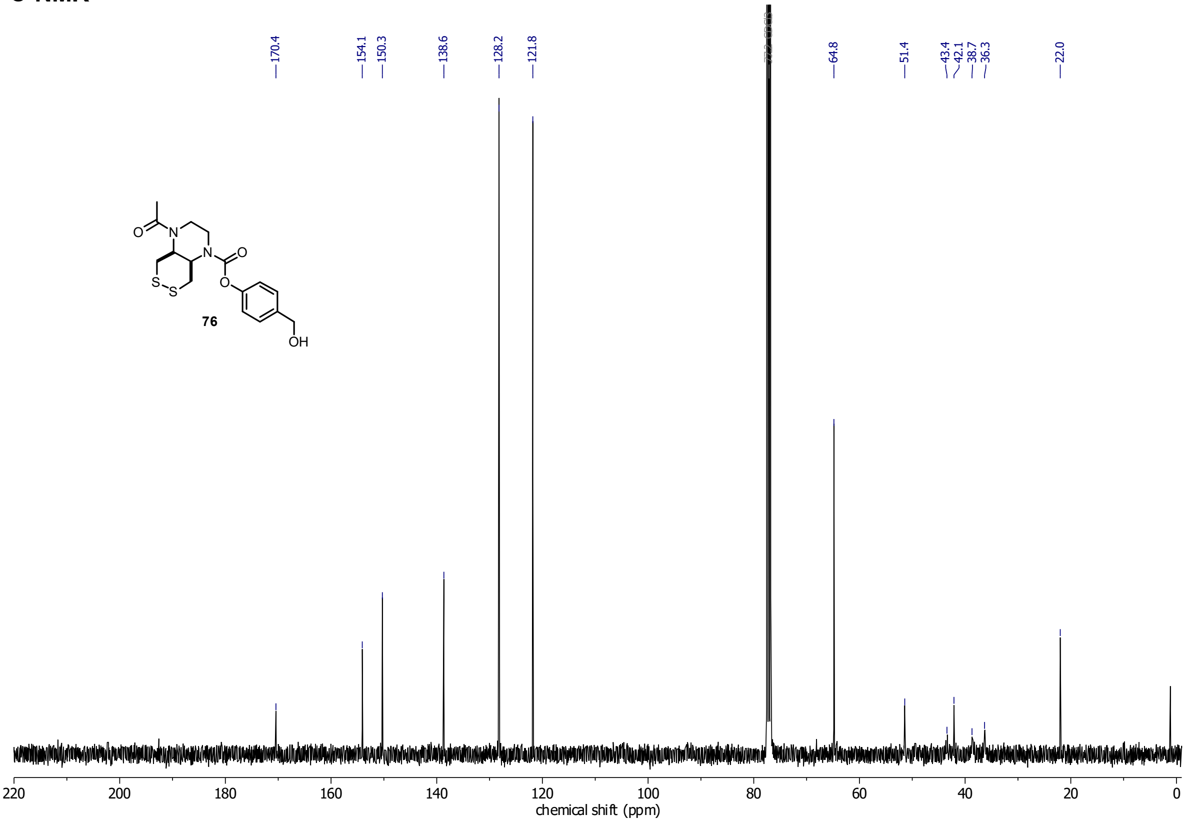

**minor silicon grease impurity*

**Compound 77**
**$^1$H-NMR**

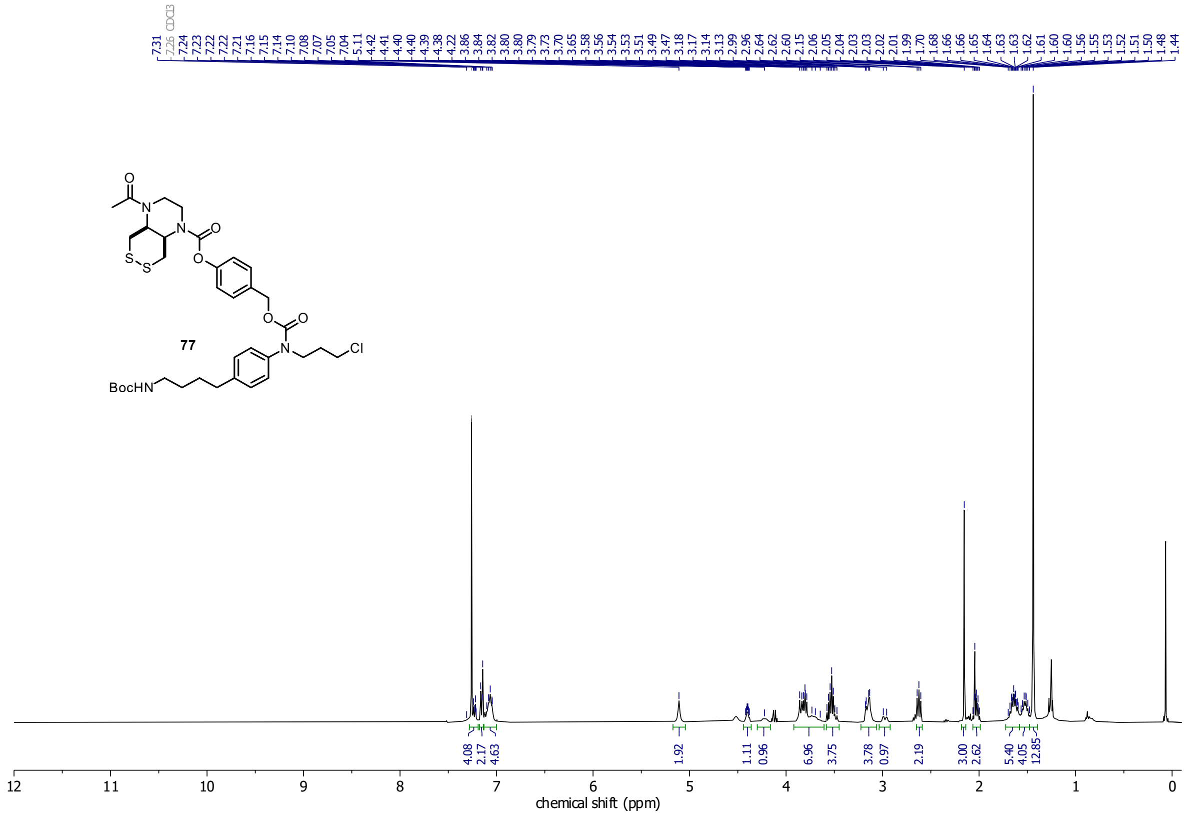

**ethyl acetate and silicon grease impurities*

**Compound 14**

**$^{1}$H-NMR**

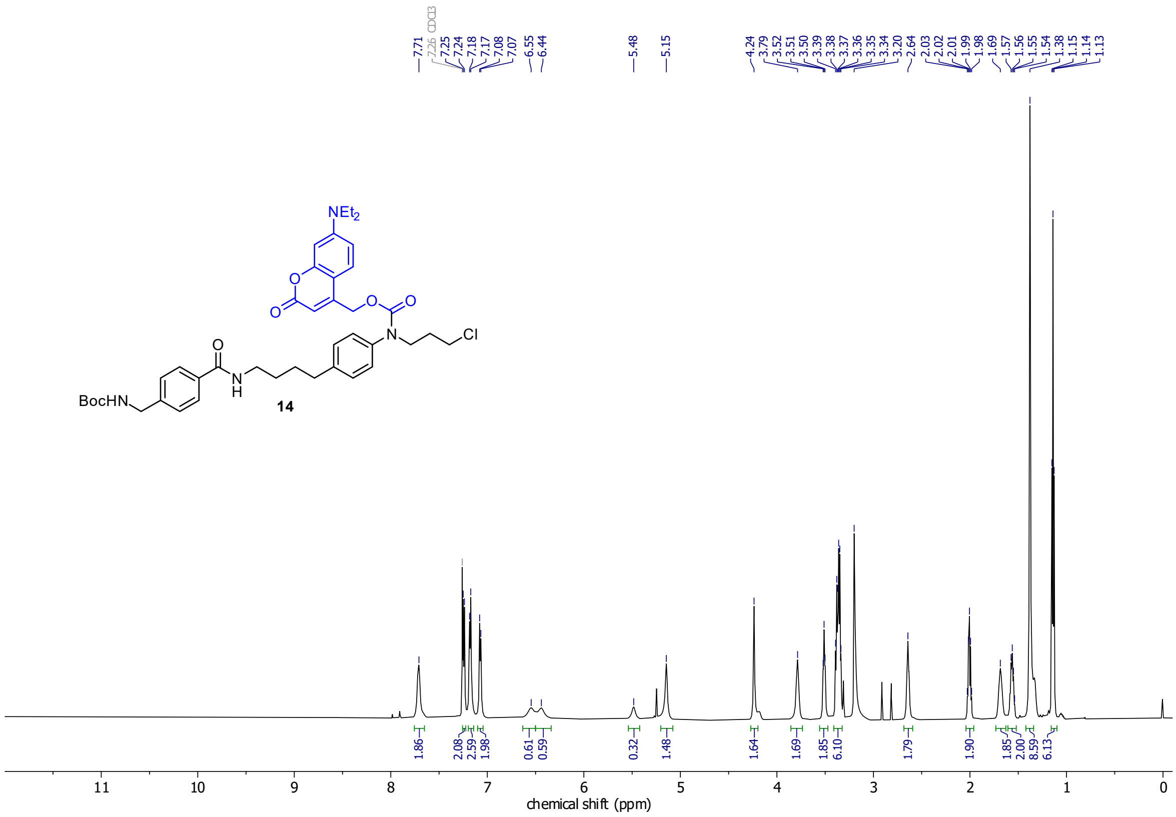

**$^{13}$C-NMR**

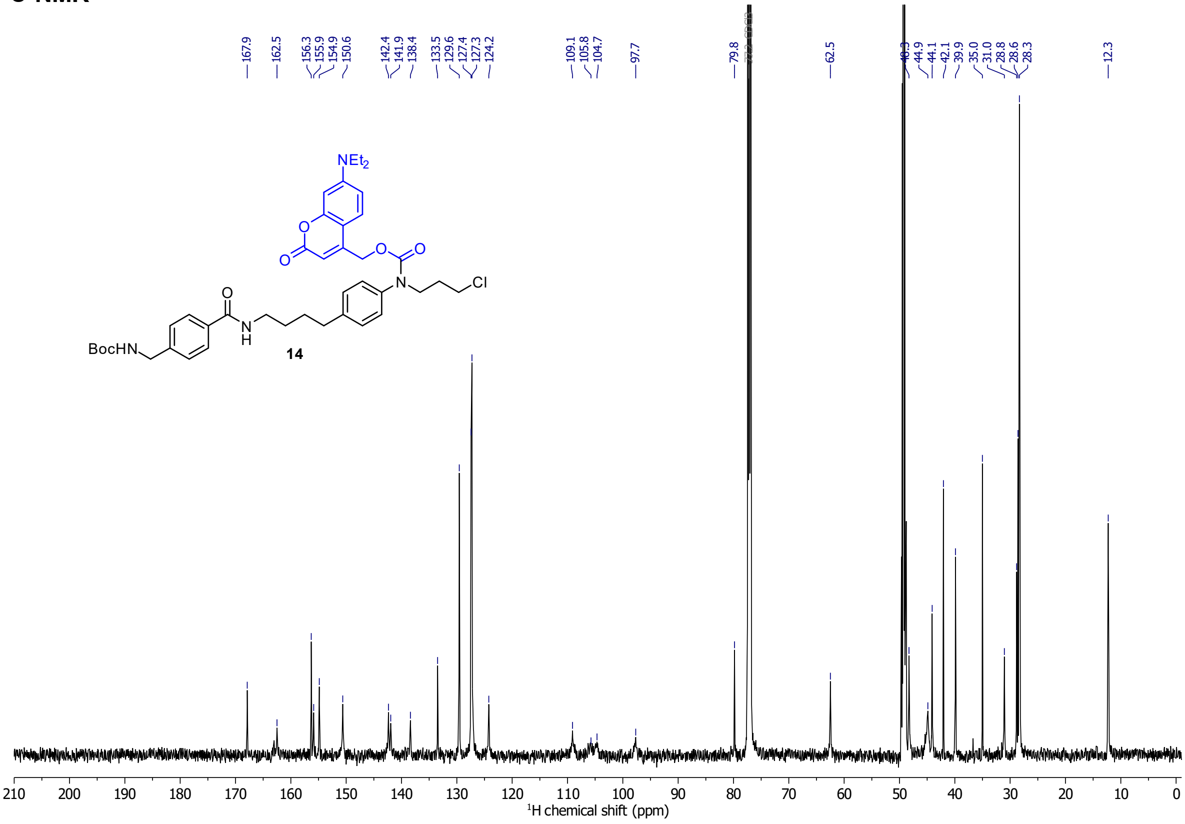

**HSQC**

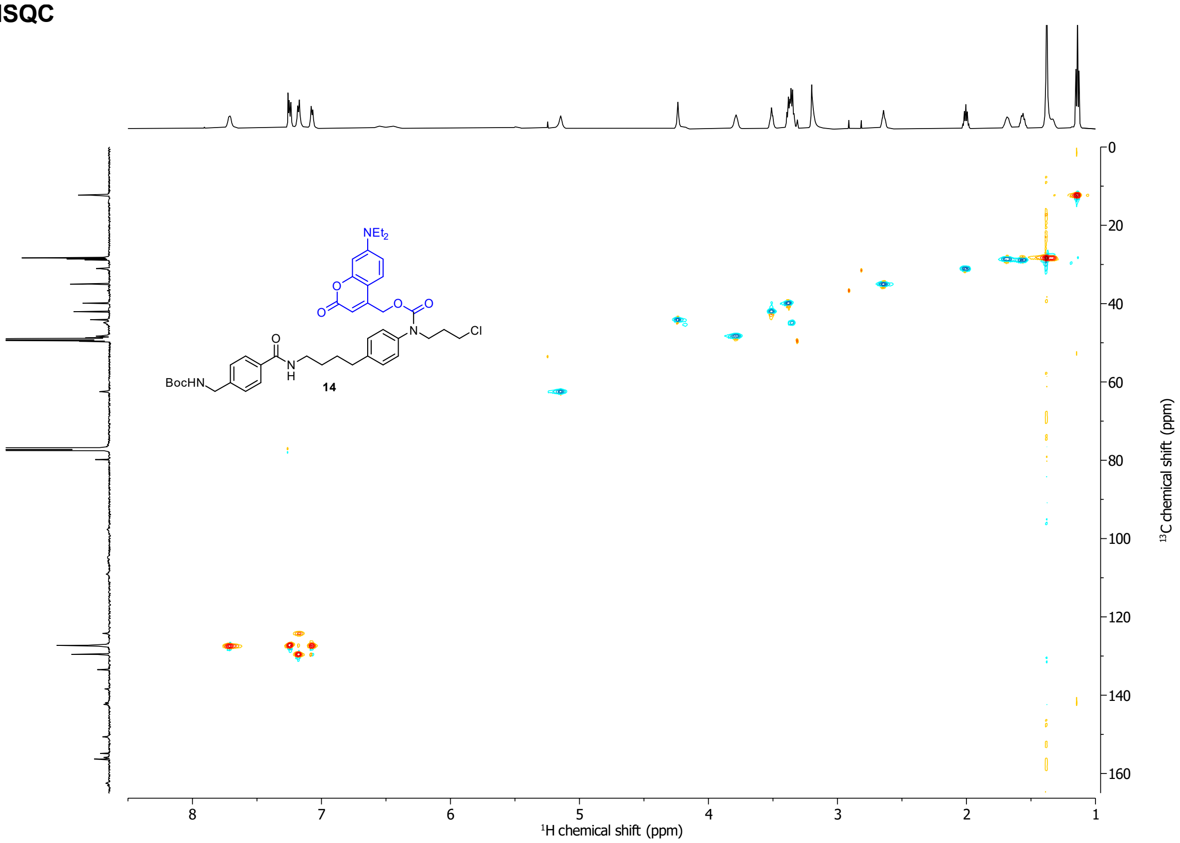

NEt2
O
O
O
O
N
Cl
O
N
H
BocHN
14
0
20
40
60
80
100
120
140
160
8
7
6
5
4
3
2
1
1H chemical shift (ppm)
13C chemical shift (ppm)

**HMBC**

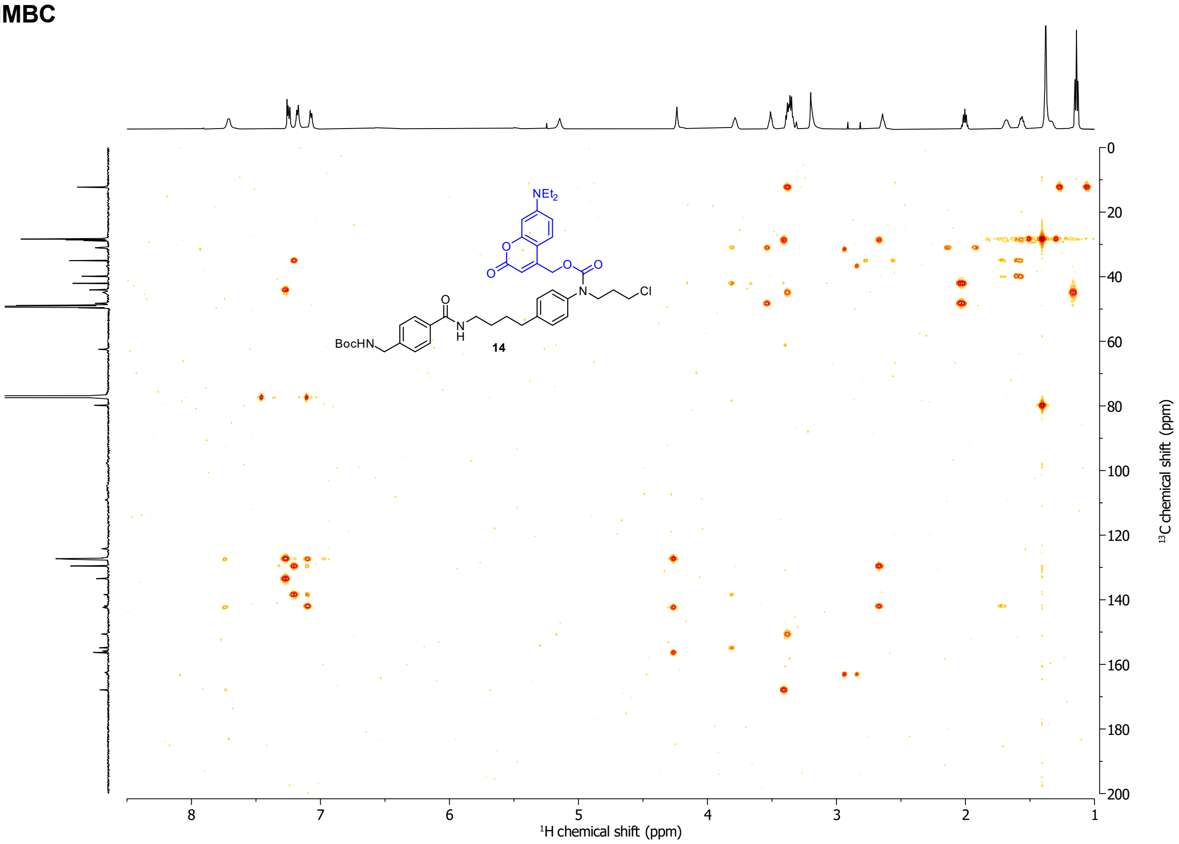

NEt2
O
O
O
O
N
Cl
O
N
H
BocHN
14
0
20
40
60
80
100
120
140
160
180
200
8
7
6
5
4
3
2
1
1H chemical shift (ppm)
13C chemical shift (ppm)

#### Compound 17

##### $^1$H-NMR

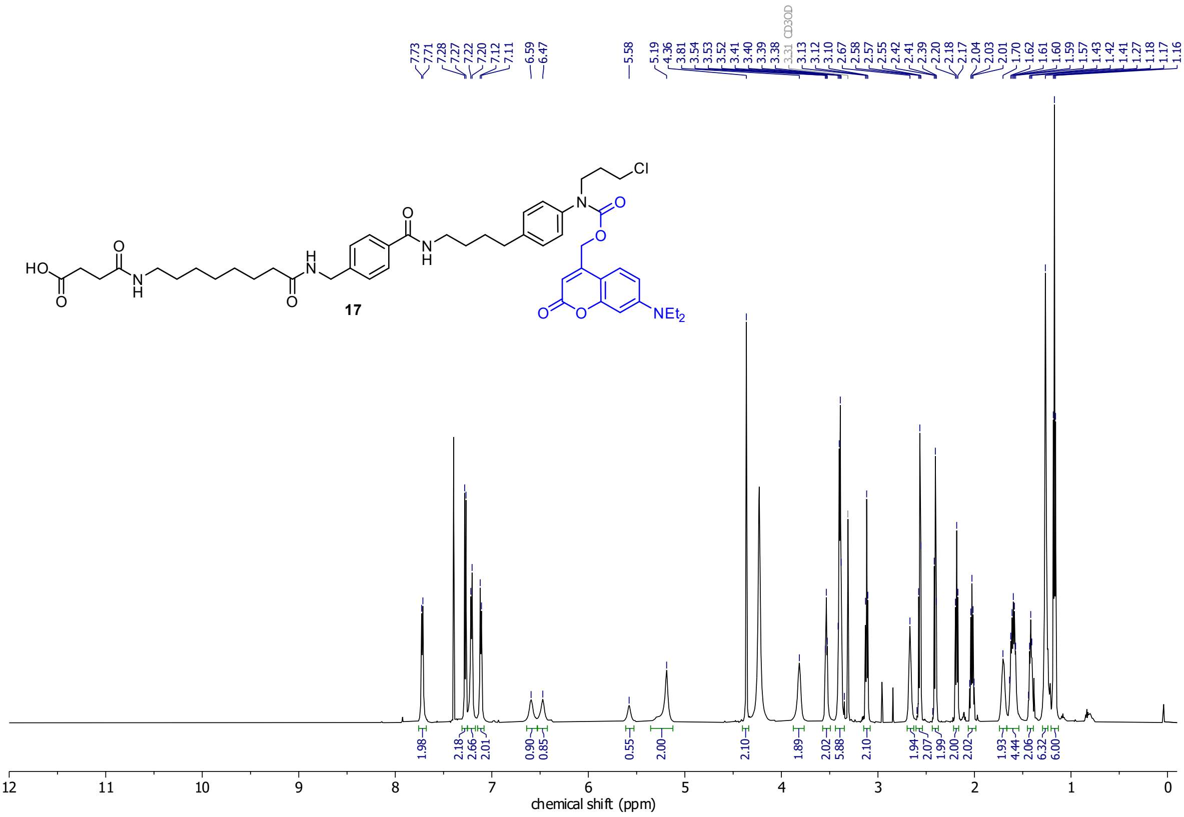

##### $^{13}$C-NMR

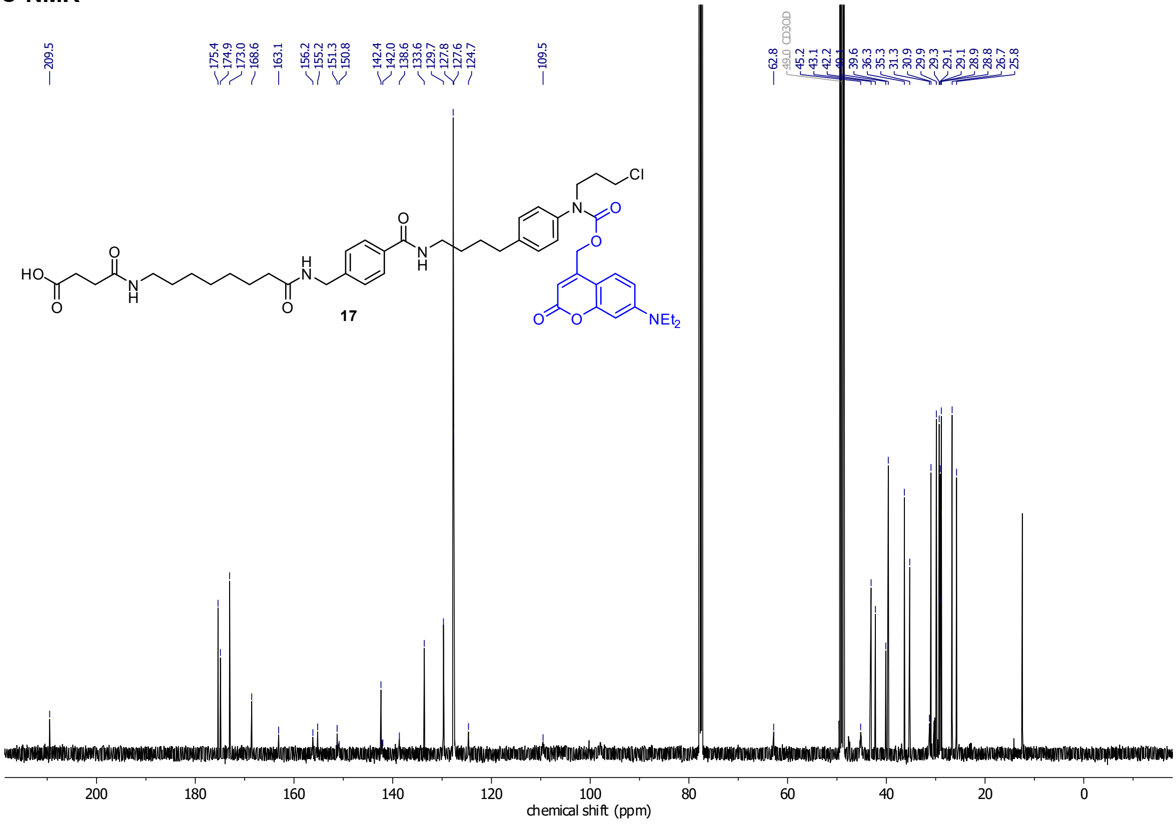

#### Cou-CHalo-BG

##### $^{1}$H-NMR

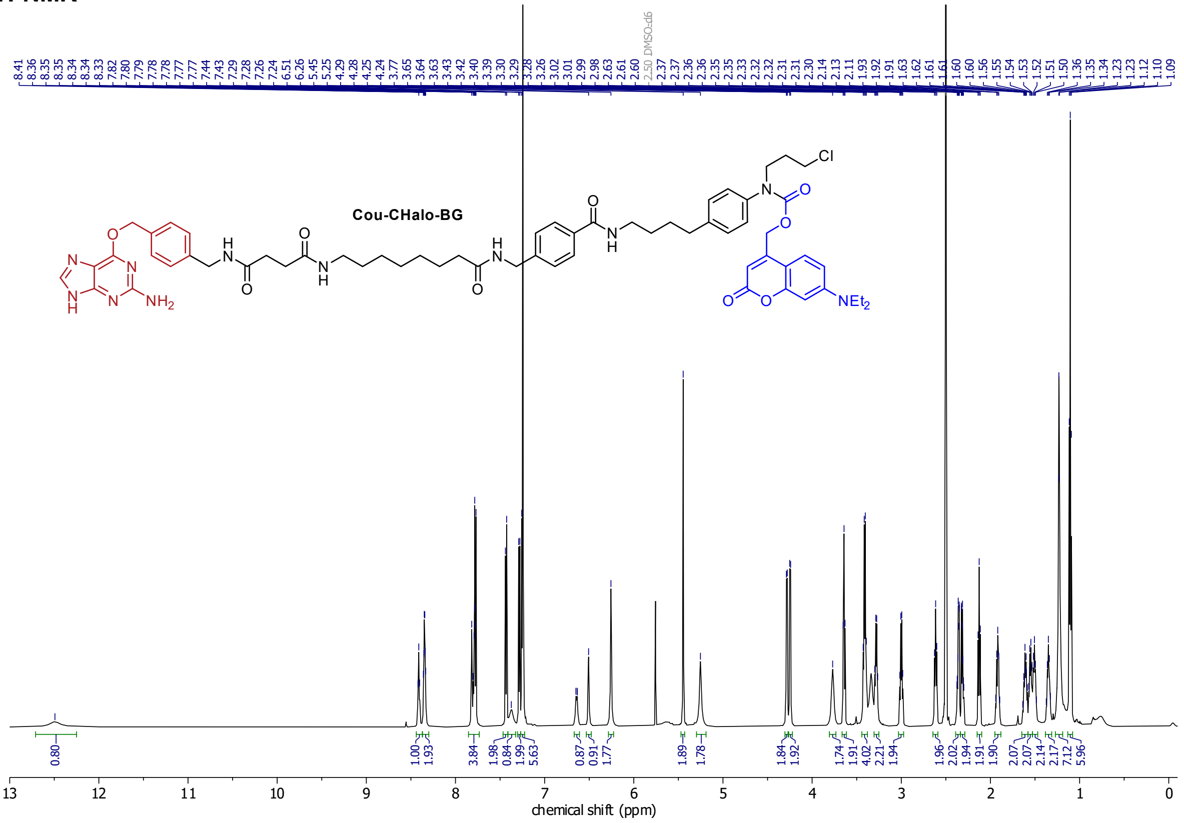

##### $^{13}$C-NMR

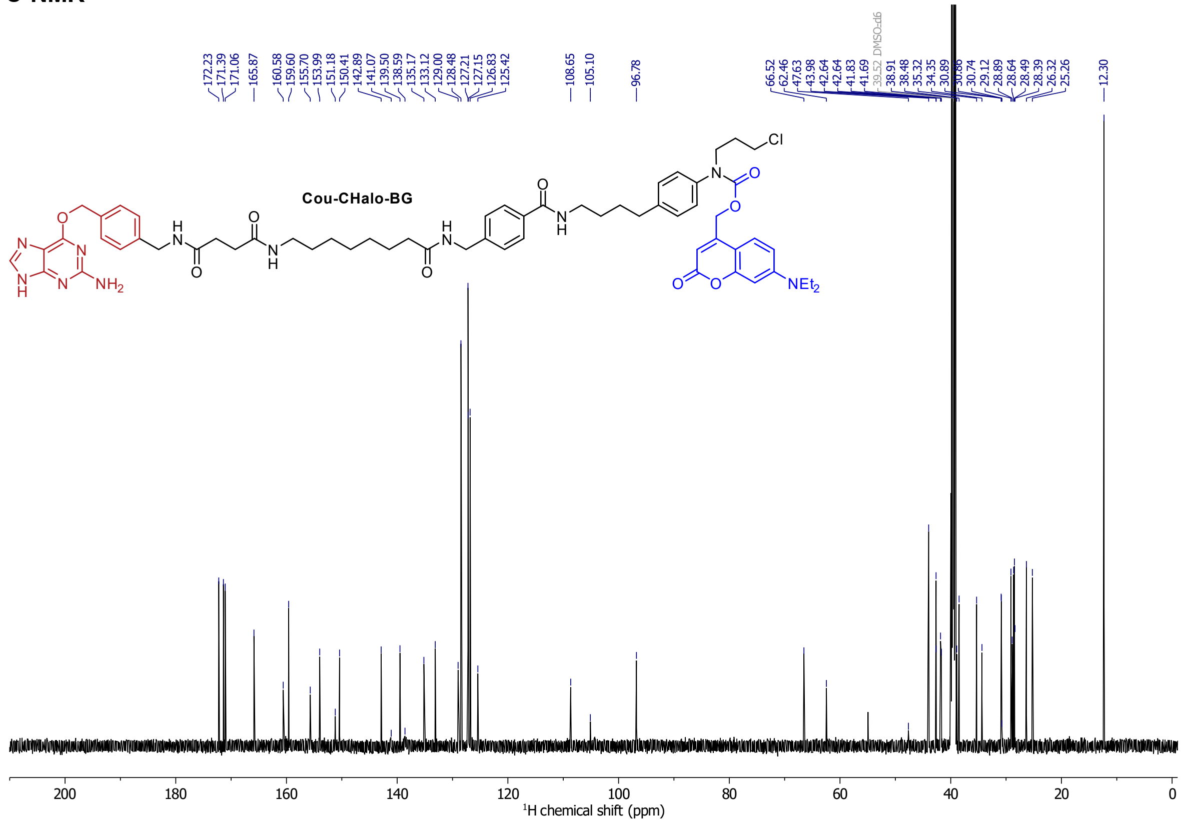

#### Compound 18

##### 1H-NMR

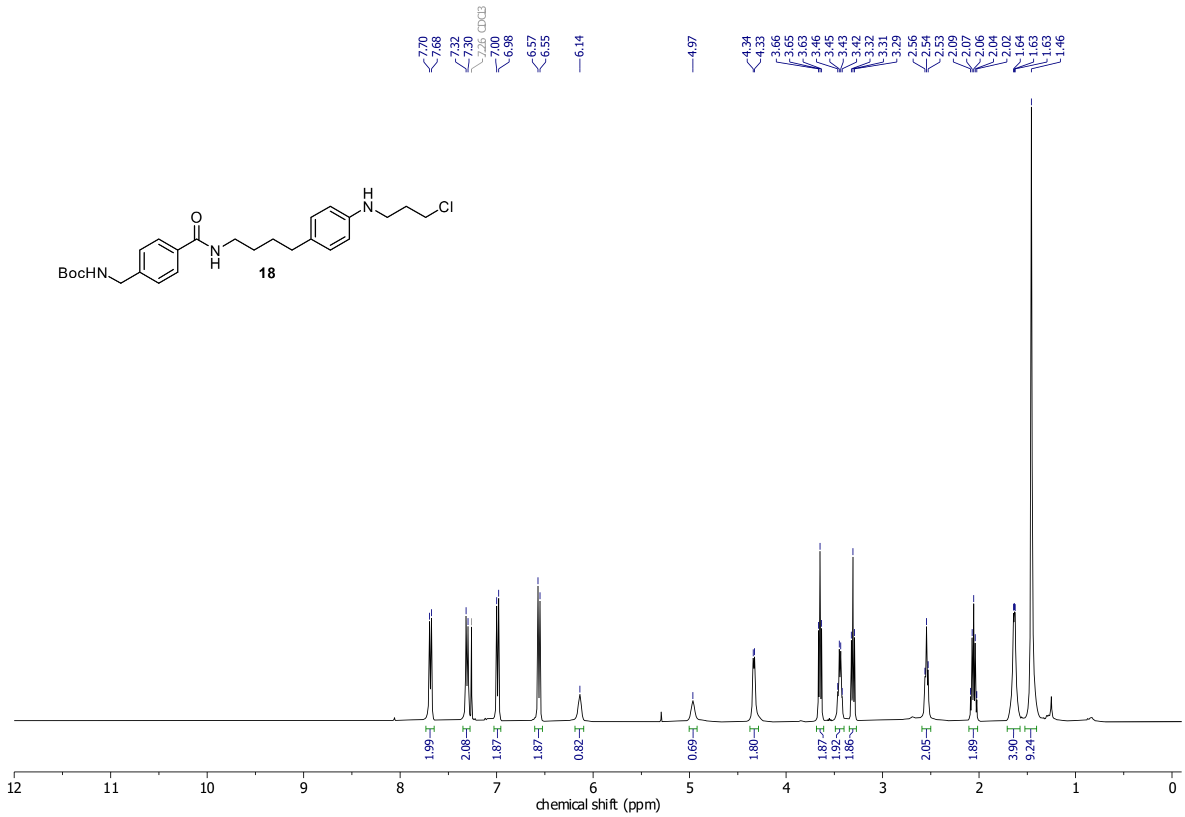

##### 13C-NMR

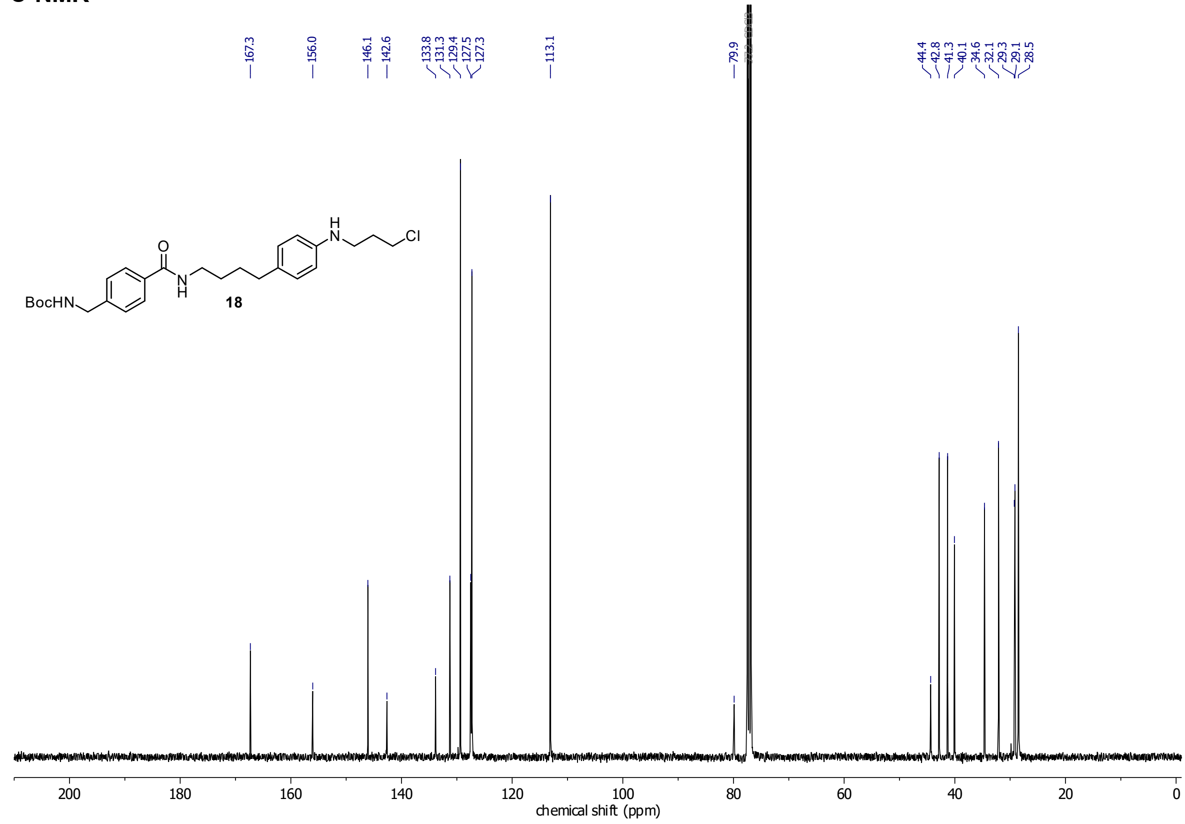

#### Compound 20

##### $^1$H-NMR

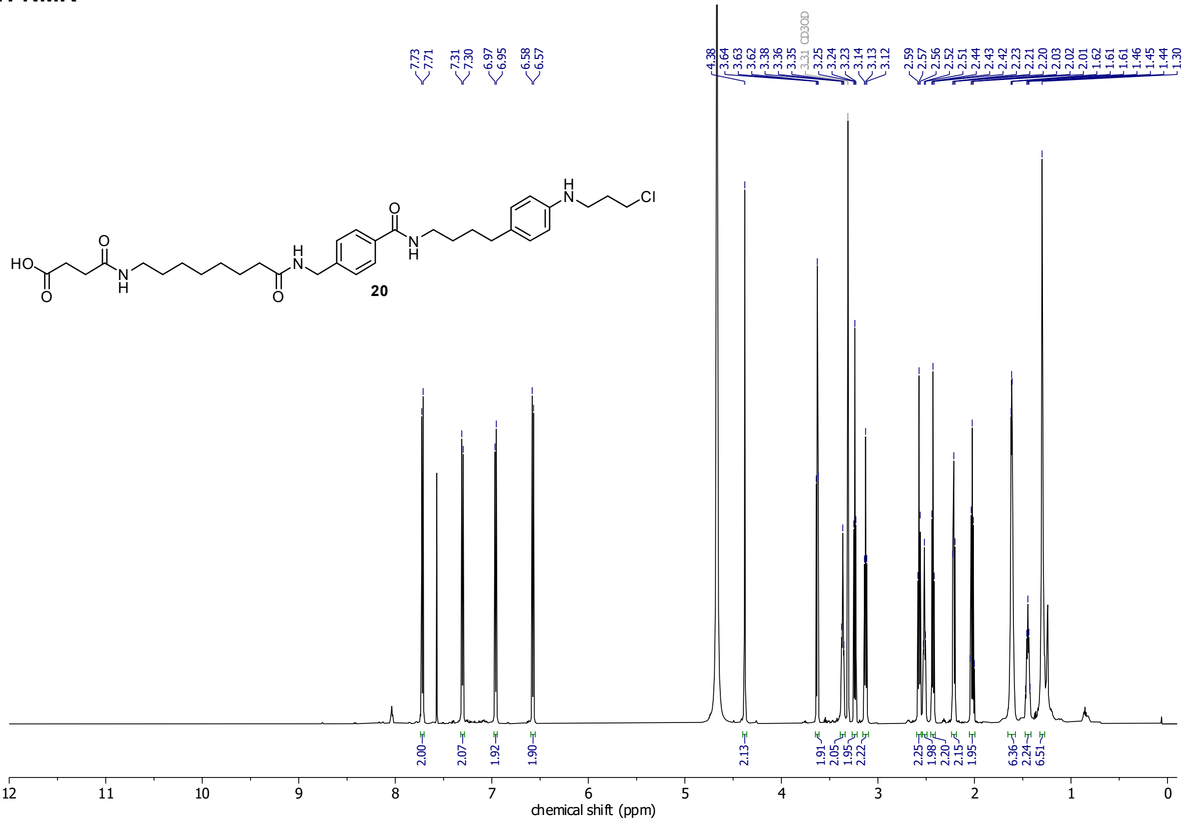

##### $^{13}$C-NMR

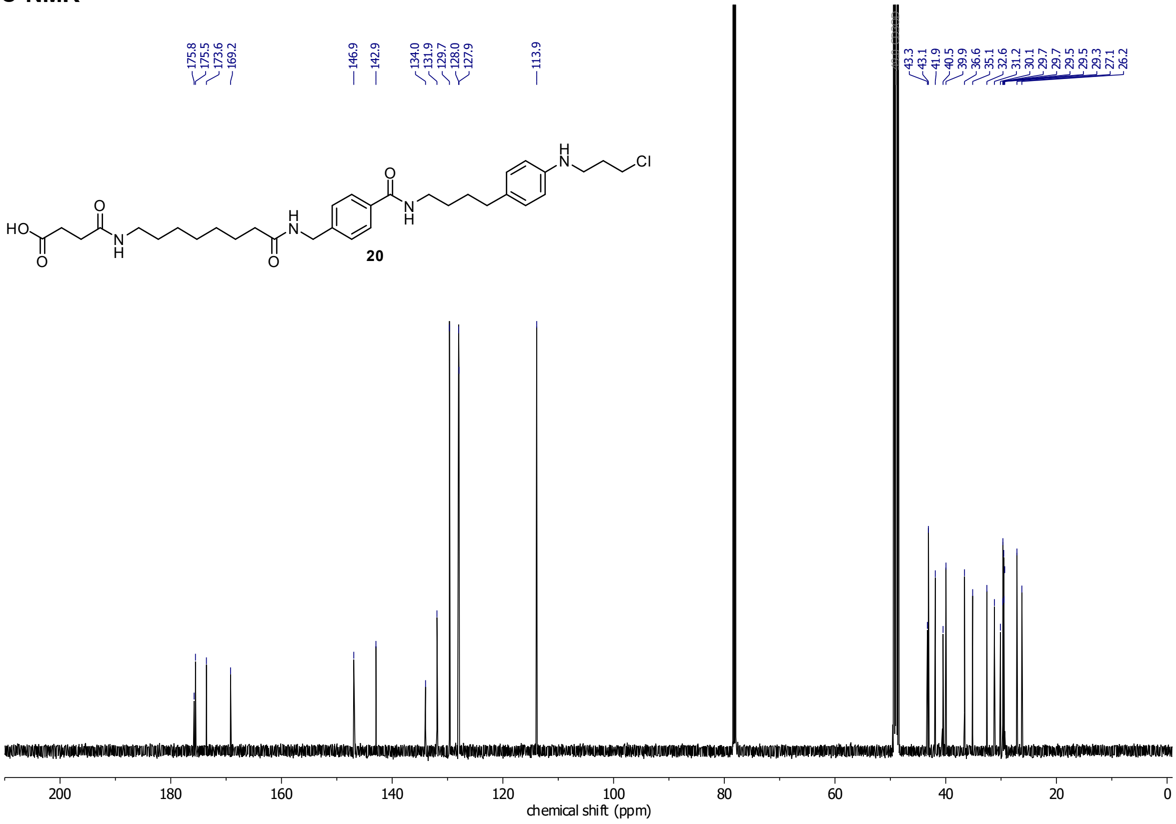

#### CHalo-BG

##### $^{1}$H-NMR

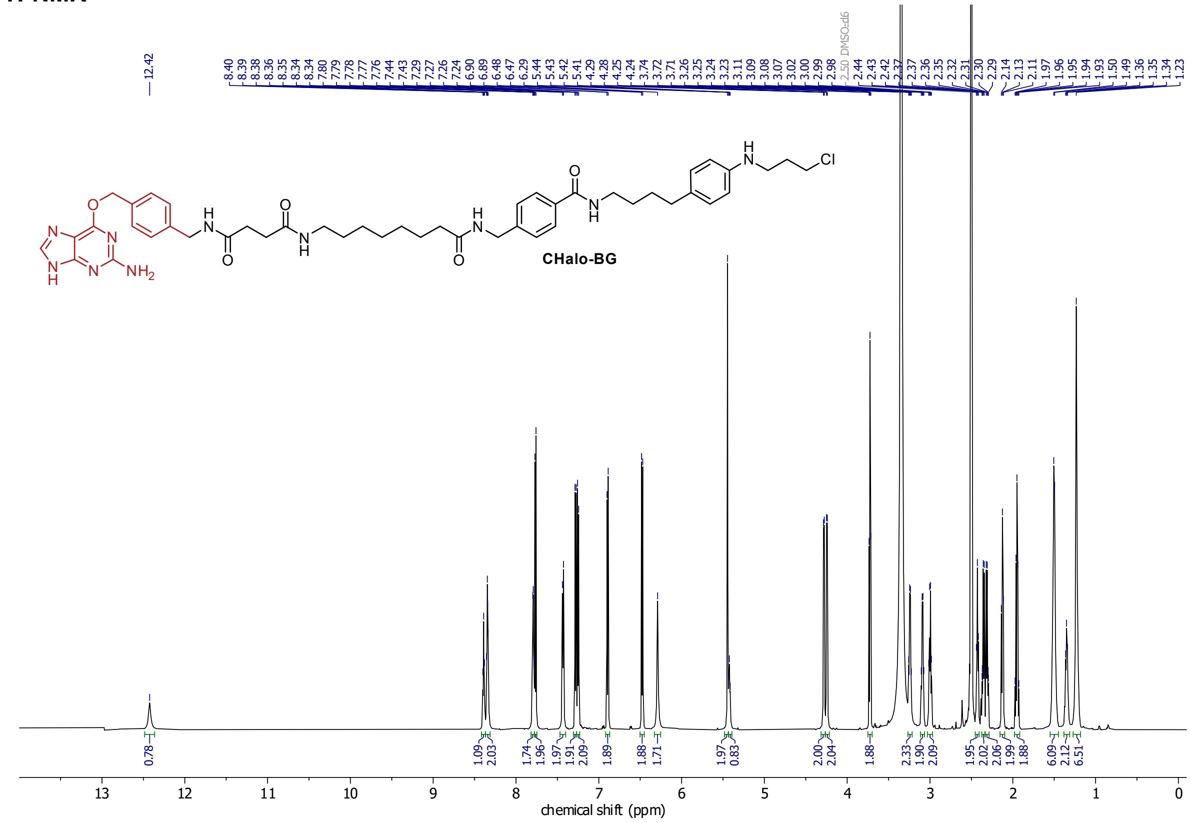

##### $^{13}$C-NMR

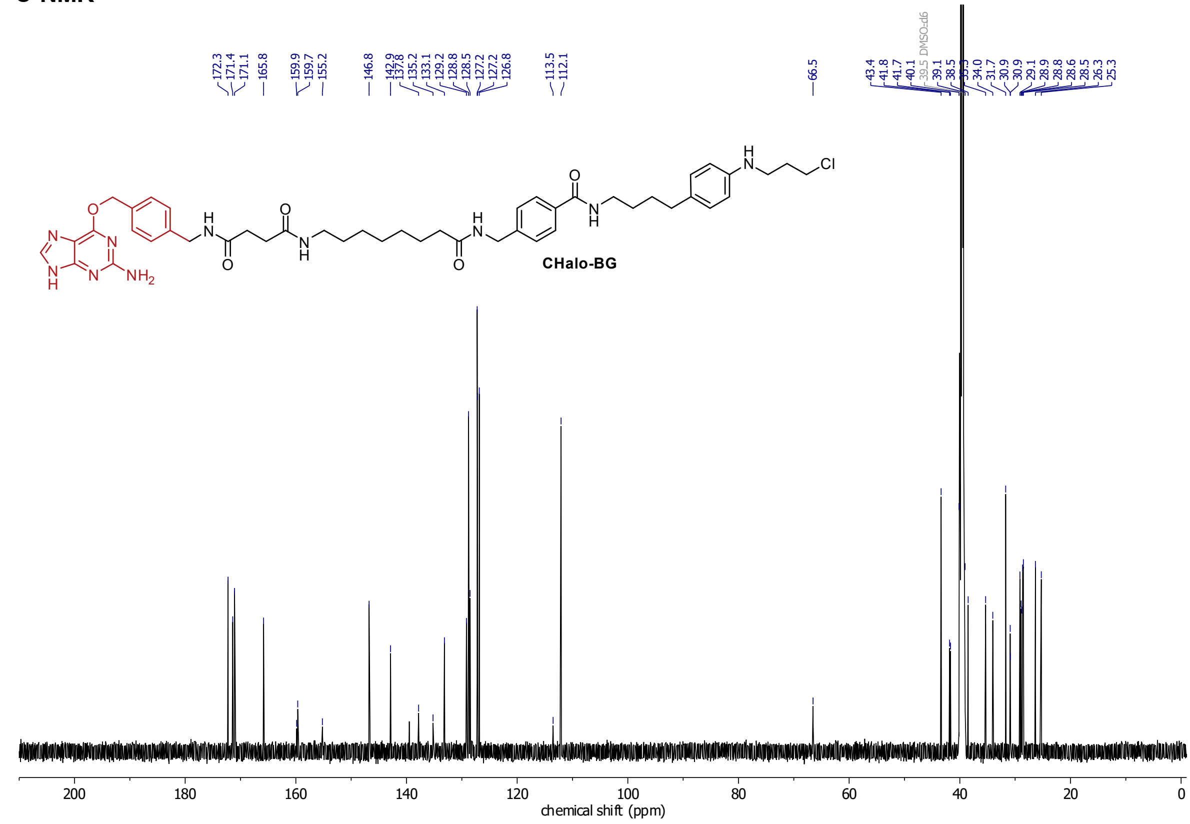

**Compound 22**

**$^{1}$H-NMR**

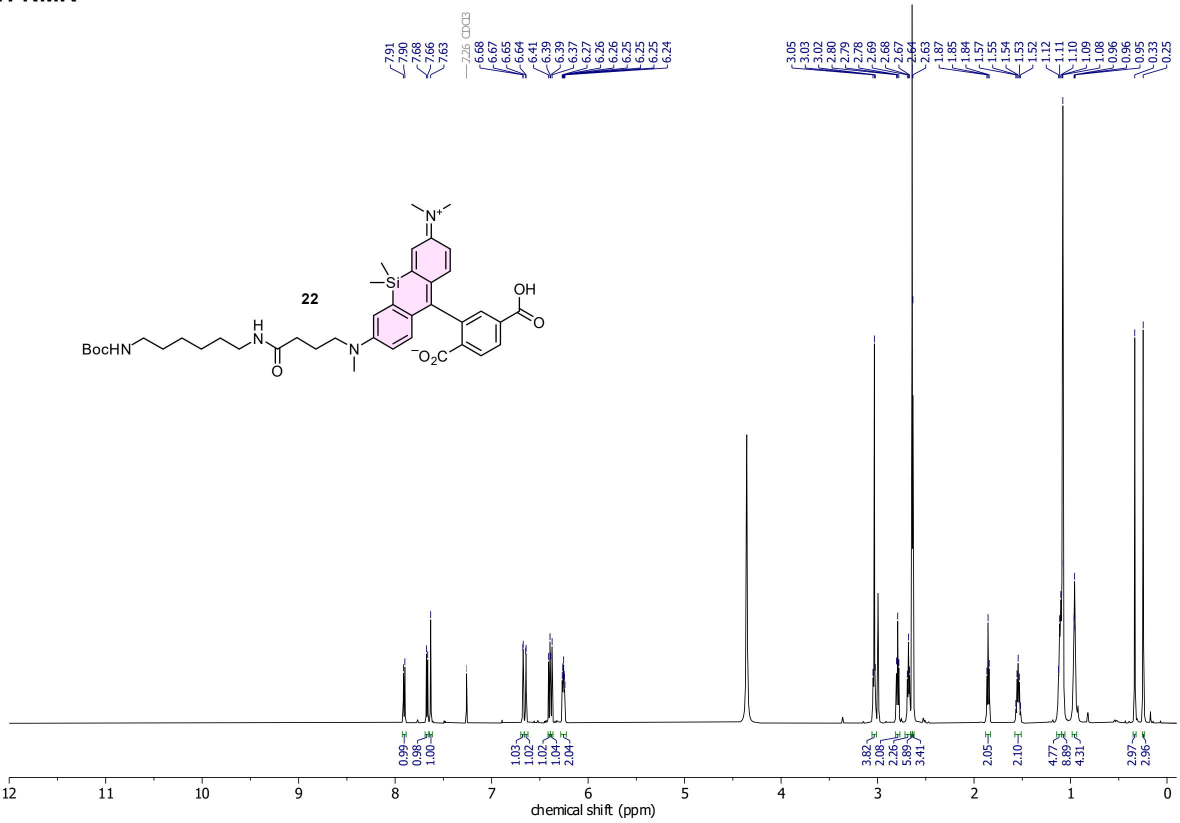

**$^{13}$C-NMR**

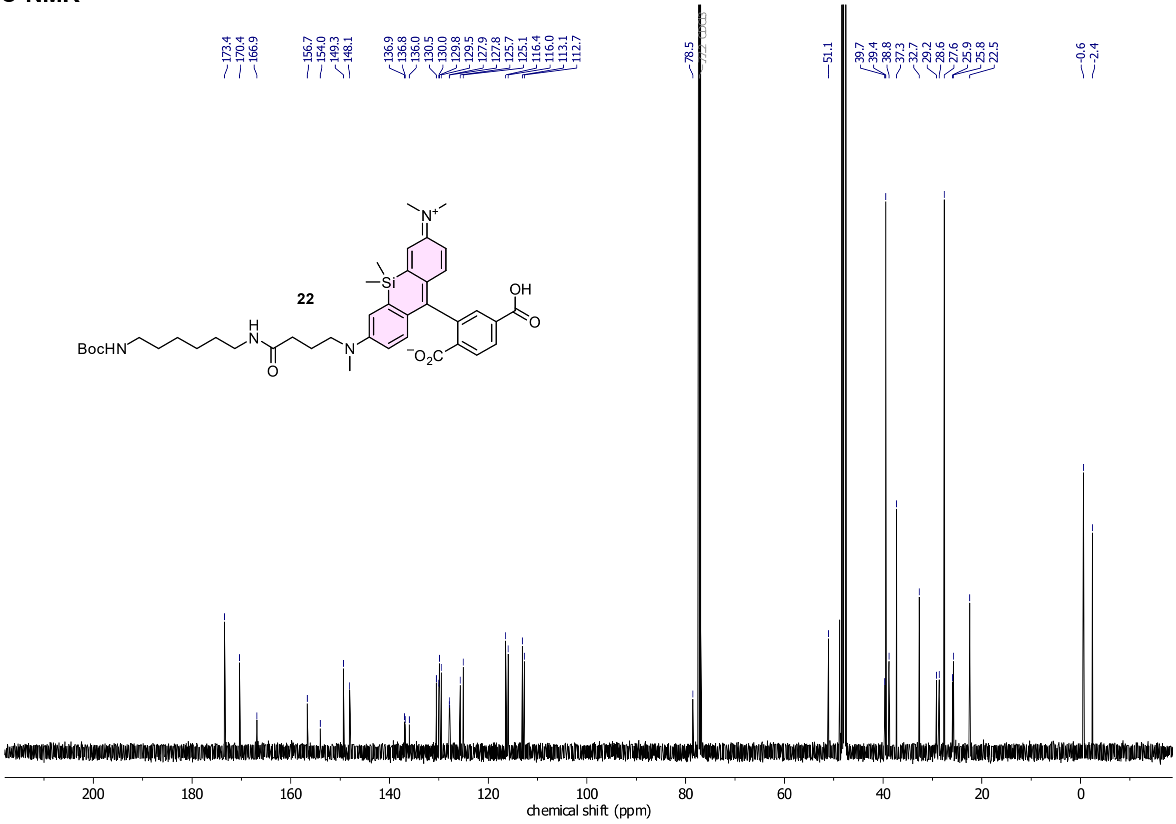

**HSQC**

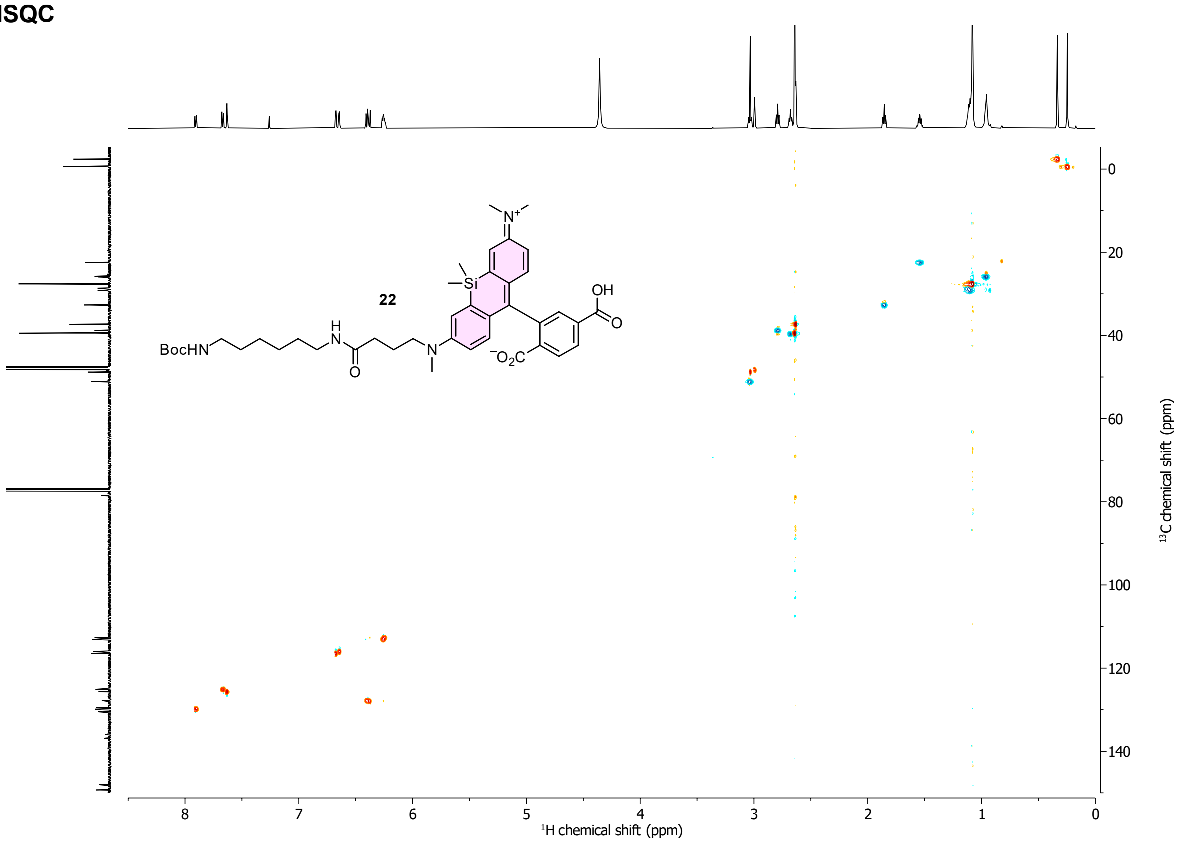

22
BocHN
H
N
O
N
Si
N+
OH
O
-O2C
1H chemical shift (ppm)
13C chemical shift (ppm)

**HMBC**

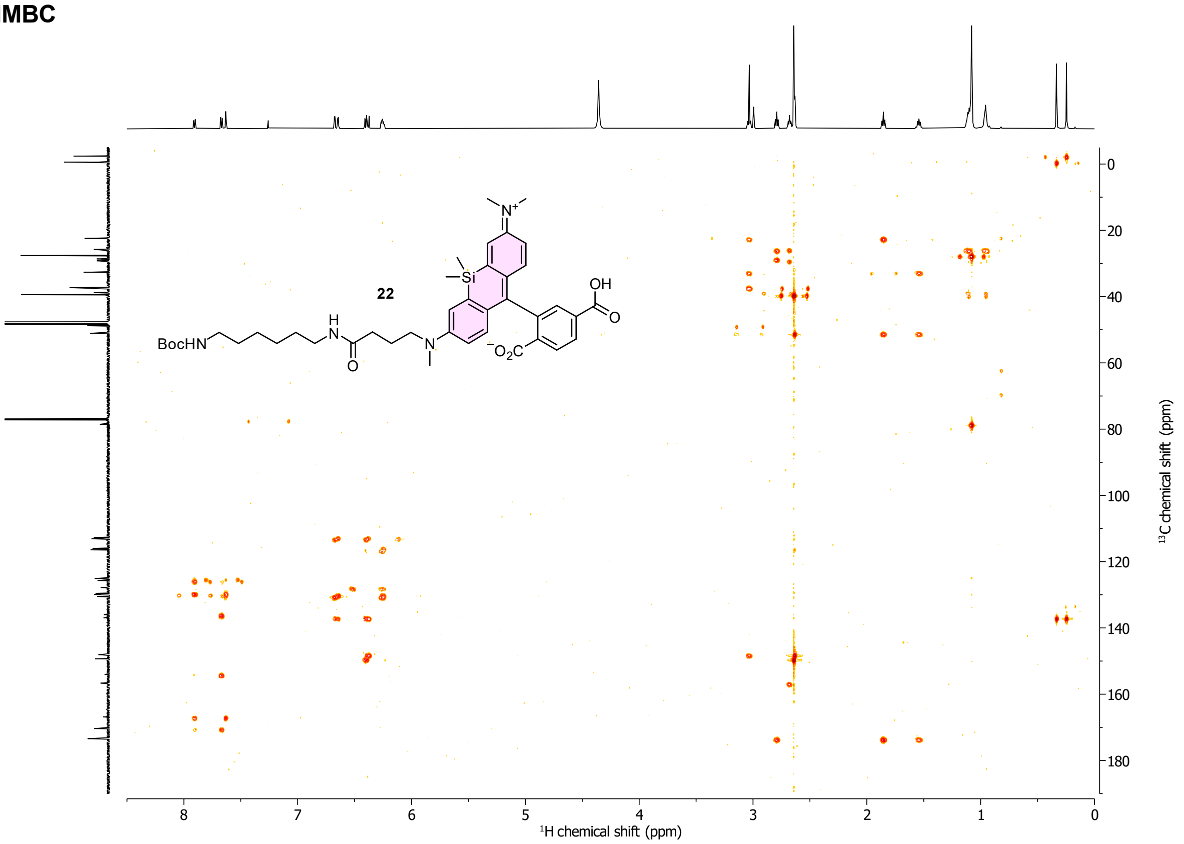

22
BocHN
H
N
O
N
Si
N+
OH
O
-O2C
1H chemical shift (ppm)
13C chemical shift (ppm)

#### Compound 24

##### $^{1}$H-NMR

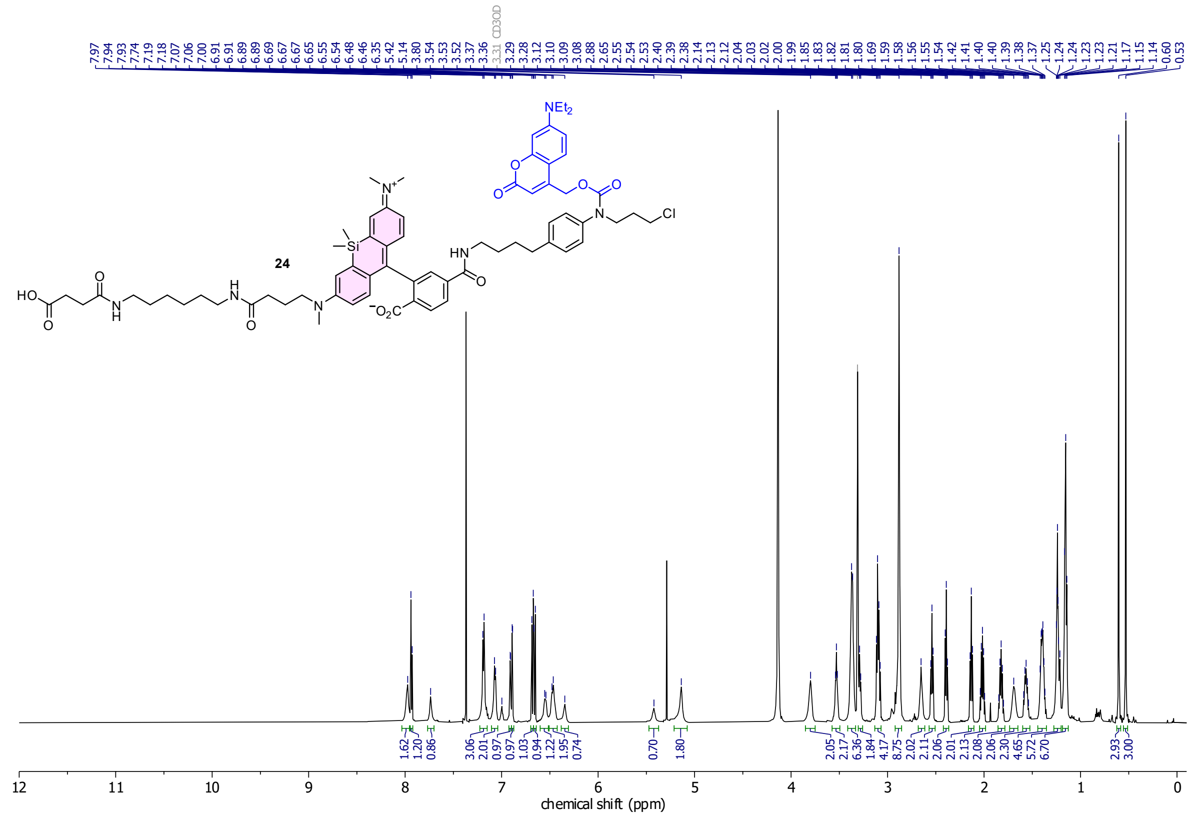

##### $^{13}$C-NMR

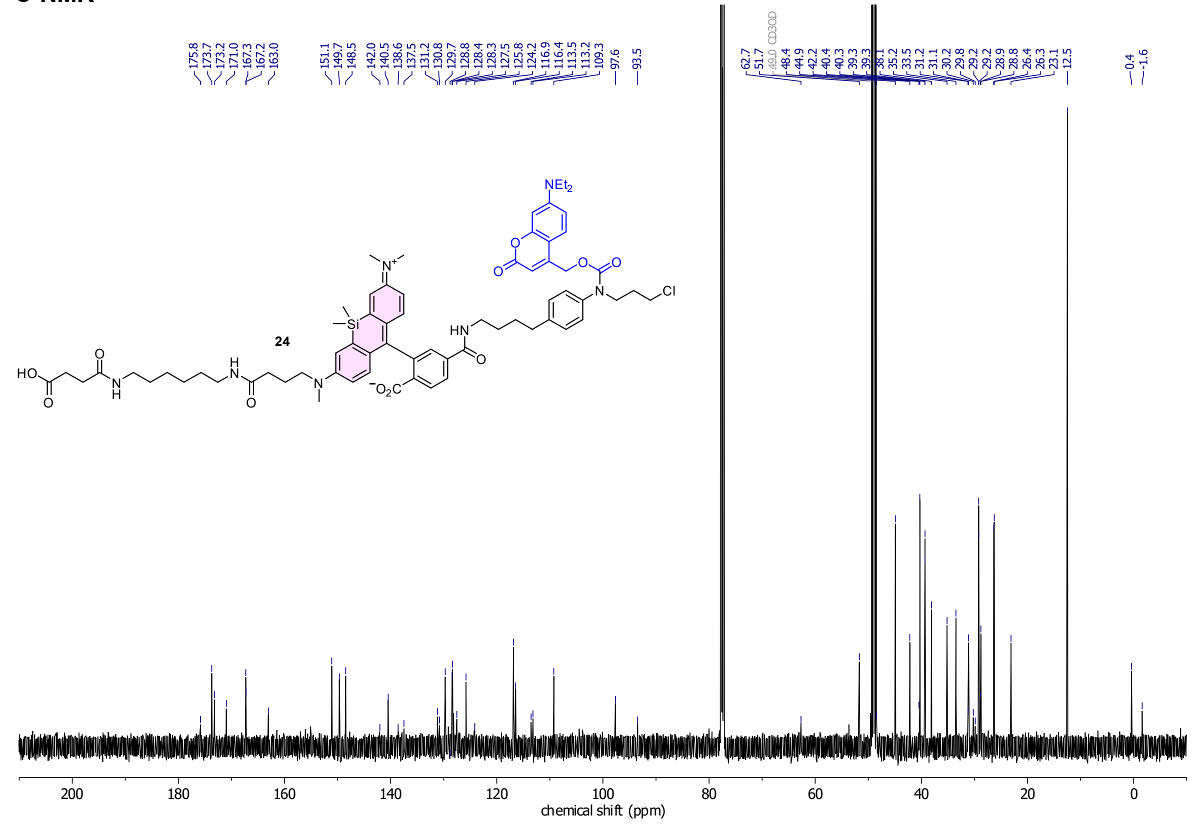

**HSQC**

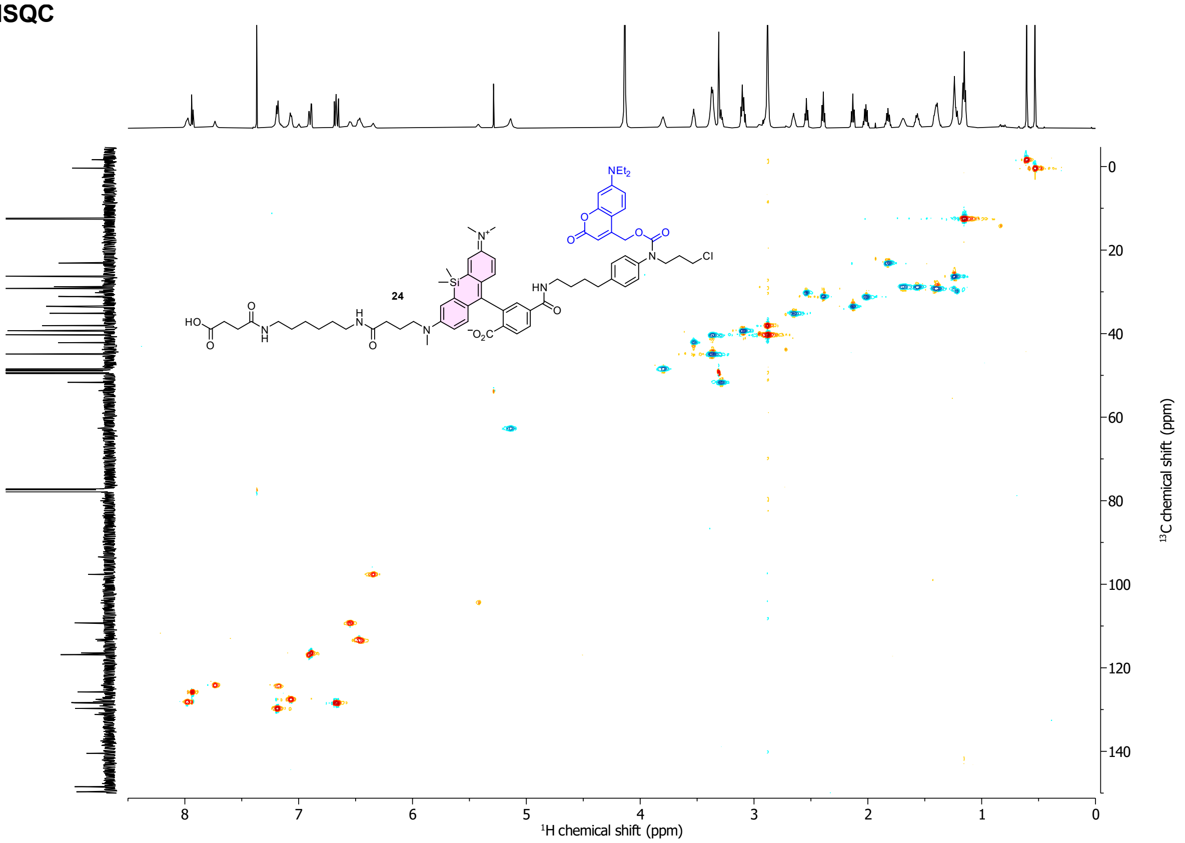

$^{1}$H chemical shift (ppm)
$^{13}$C chemical shift (ppm)
24
NEt$_2$
HN
Cl
HO
$^{-}$O$_2$C
Si

**HMBC**

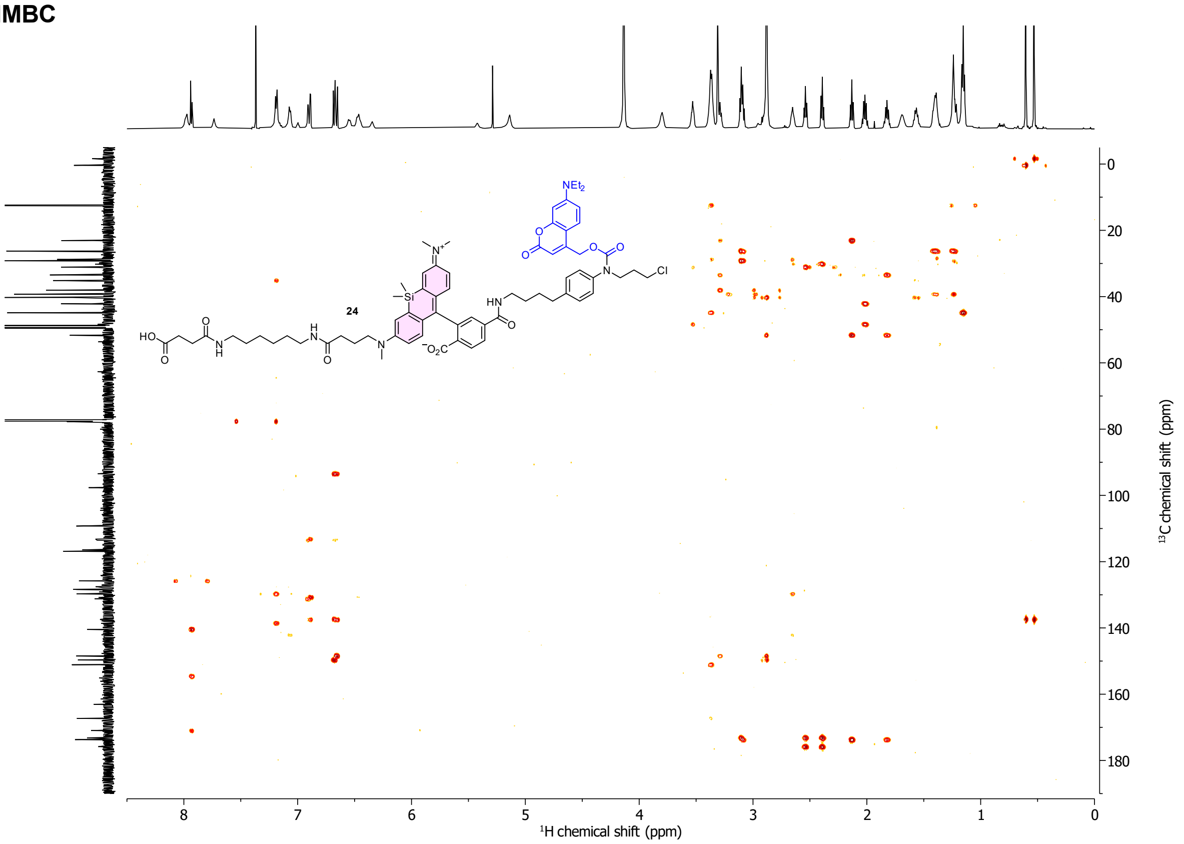

$^{1}$H chemical shift (ppm)
$^{13}$C chemical shift (ppm)
24
NEt$_2$
HN
Cl
HO
$^{-}$O$_2$C
Si

#### Cou-CHalo-SiR-BG

##### $^{1}$H-NMR

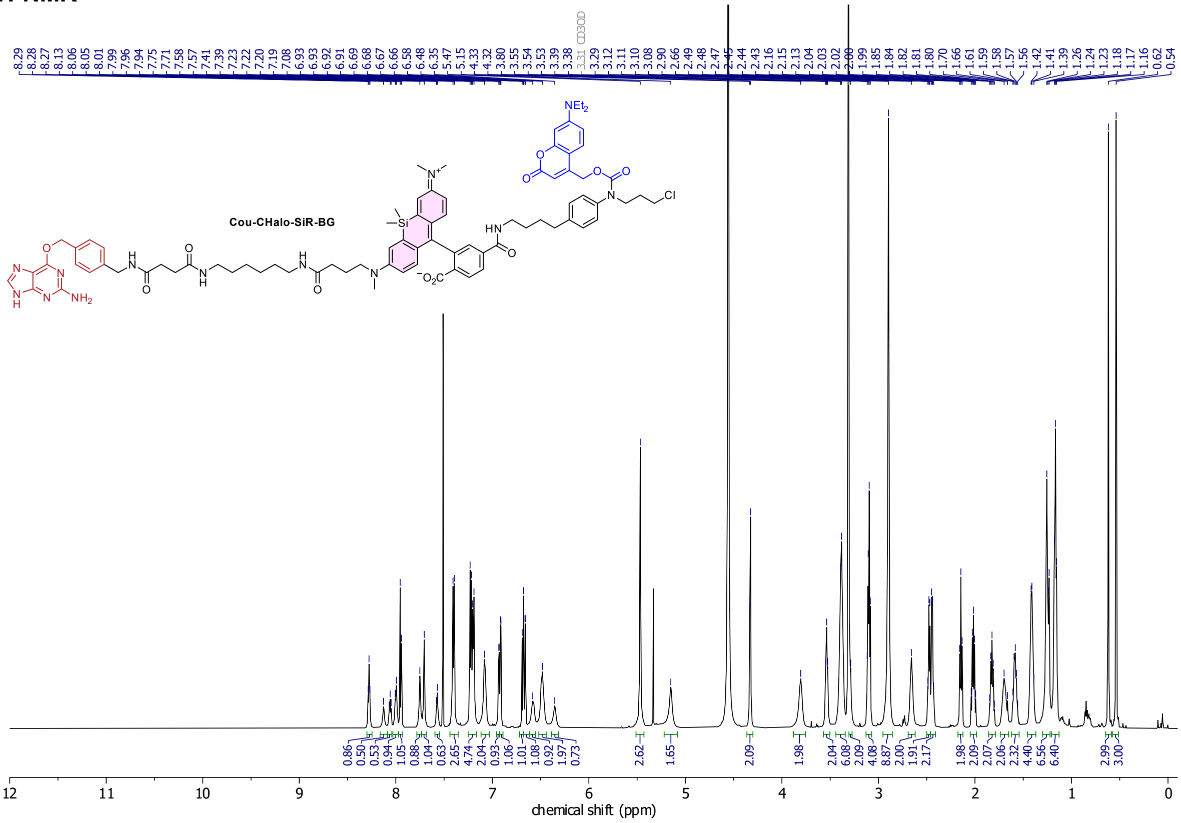

##### $^{13}$C-NMR

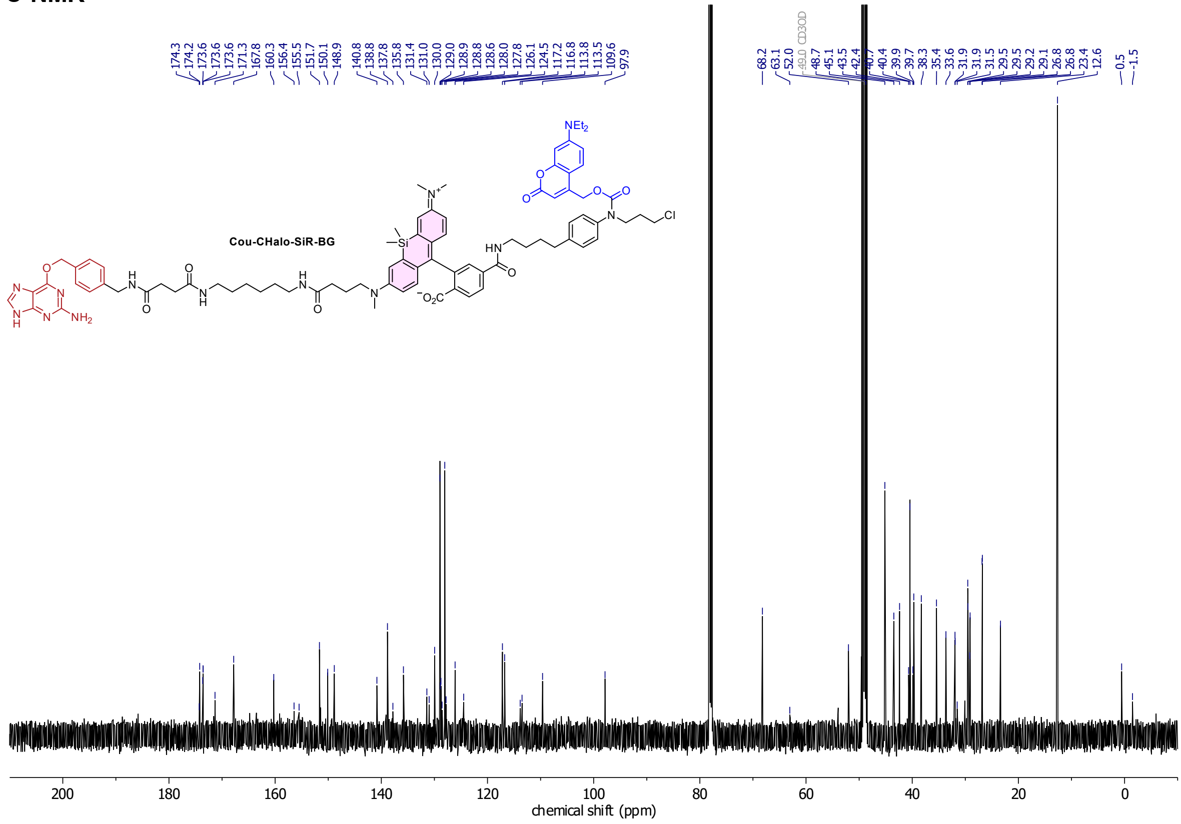

#### HSQC

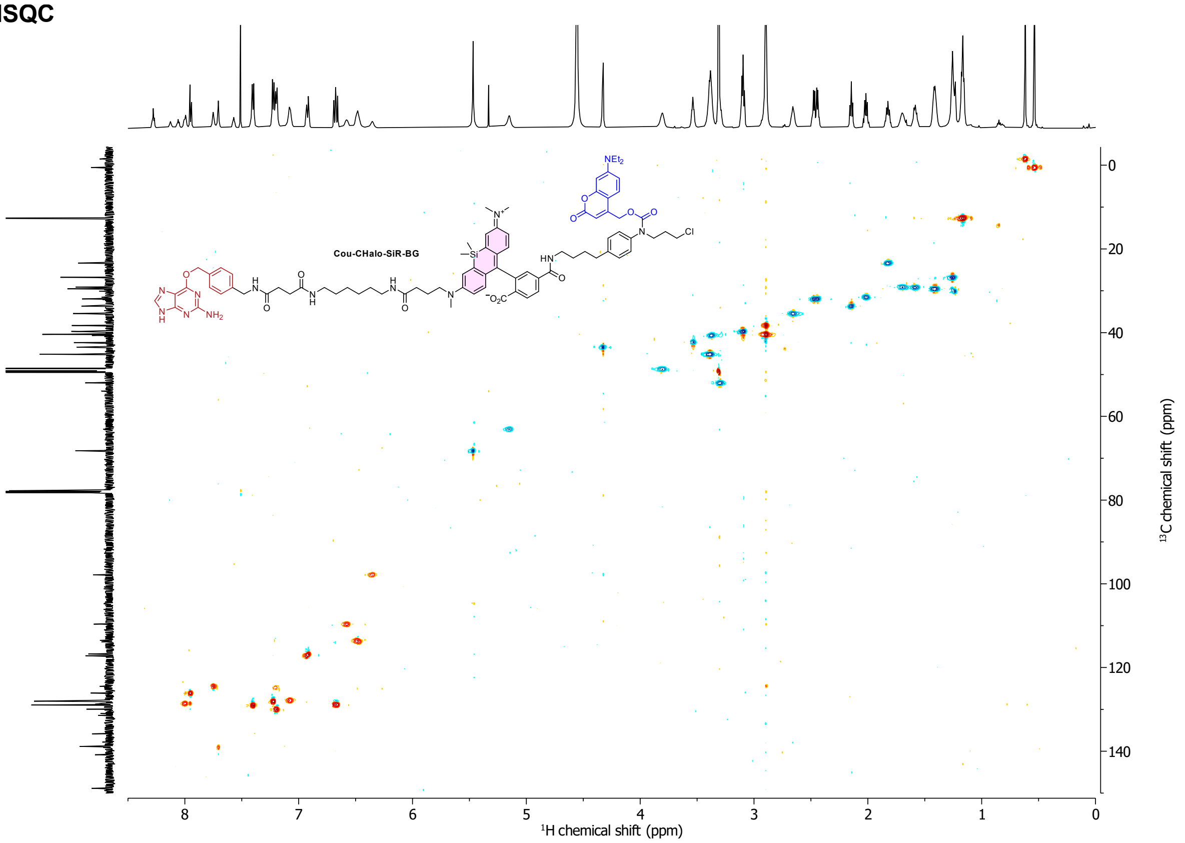

Cou-CHalo-SiR-BG
¹H chemical shift (ppm)
¹³C chemical shift (ppm)

#### HMBC

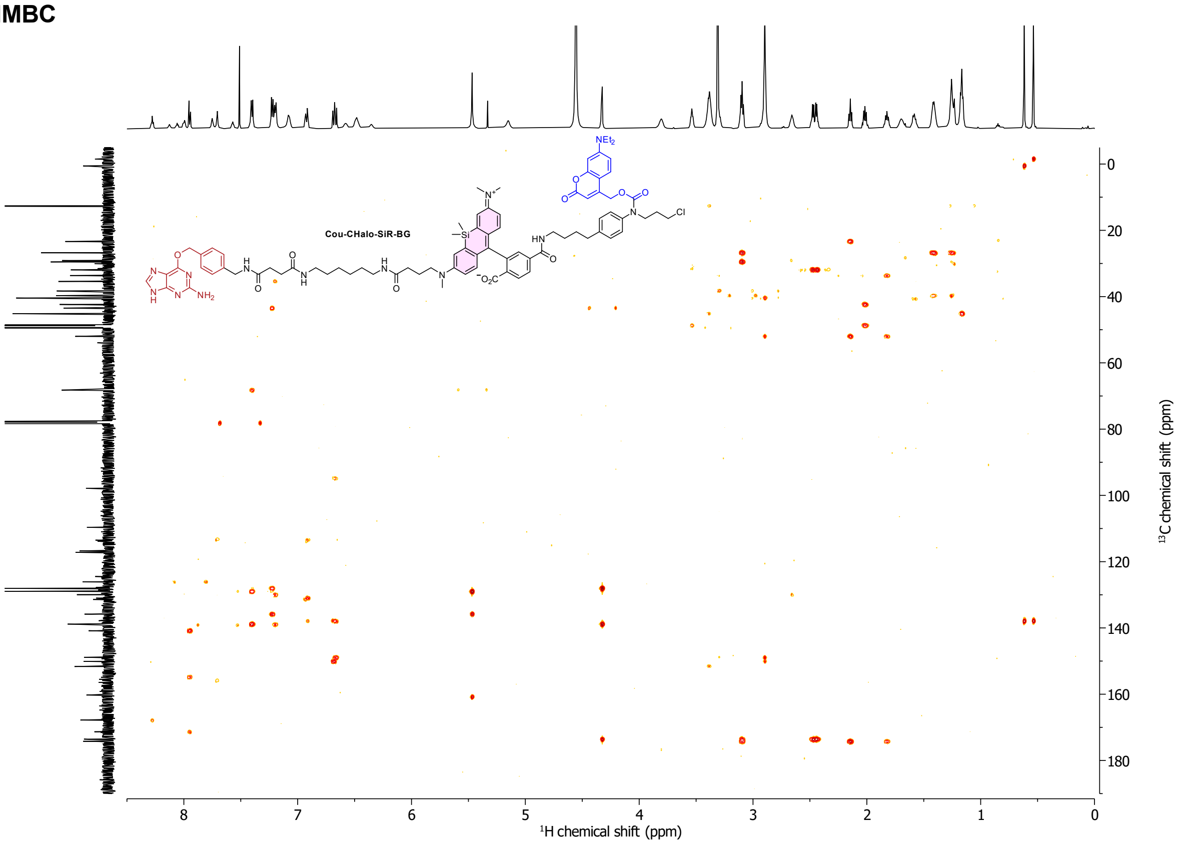

Cou-CHalo-SiR-BG
¹H chemical shift (ppm)
¹³C chemical shift (ppm)

**Compound 26**

**$^1$H-NMR**

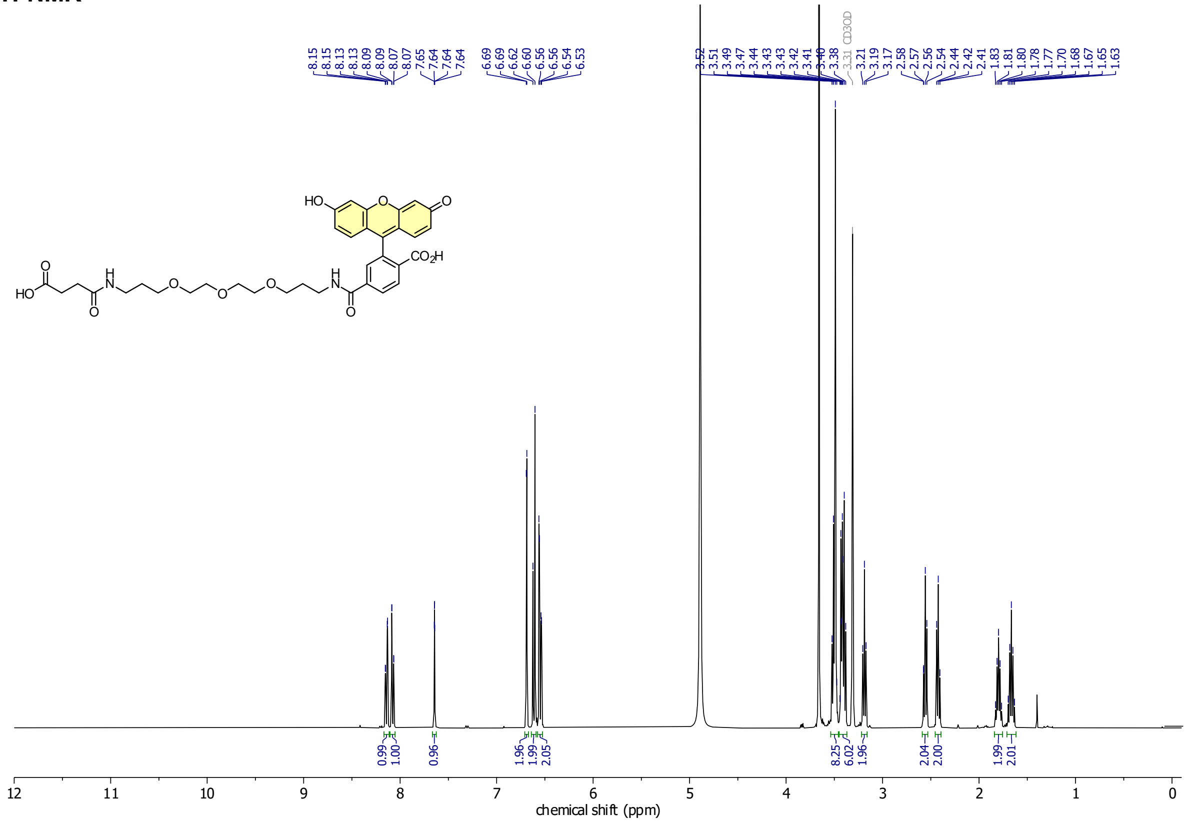

**$^{13}$C-NMR**

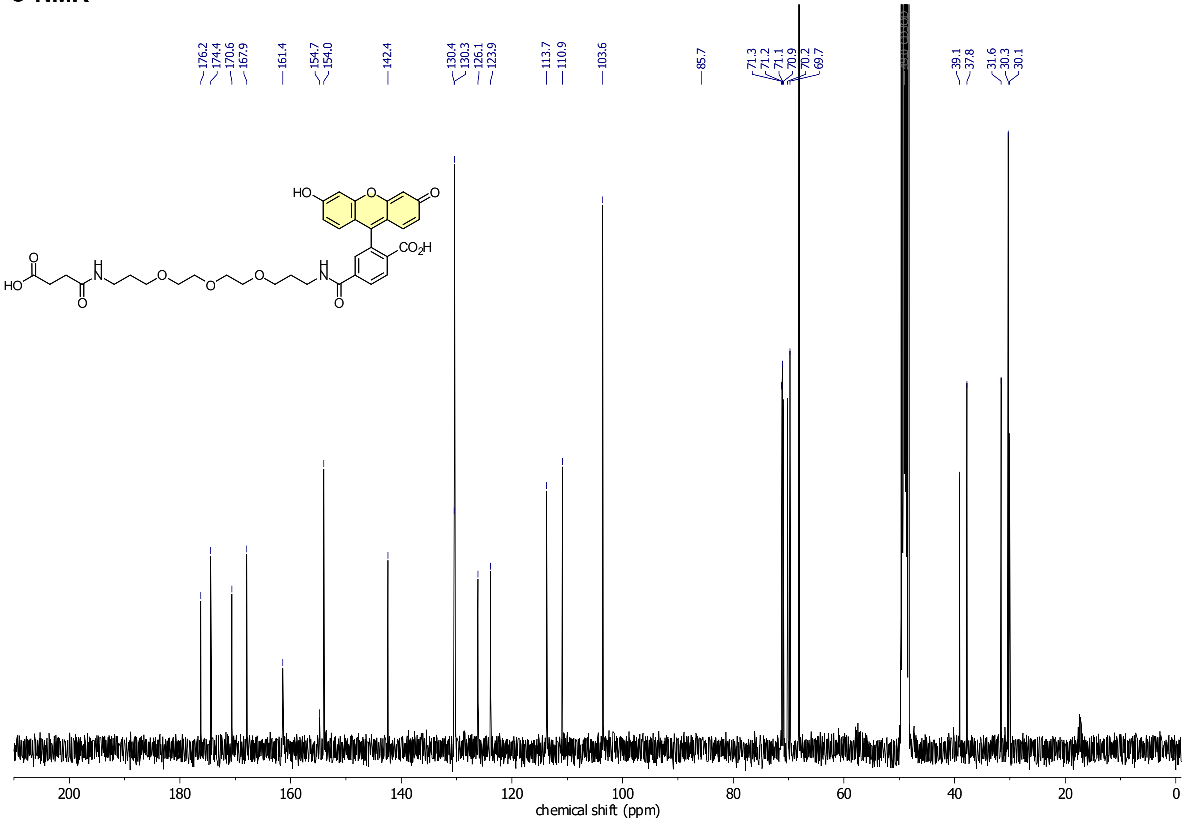

**Compound 30**

**$^{1}$H-NMR**

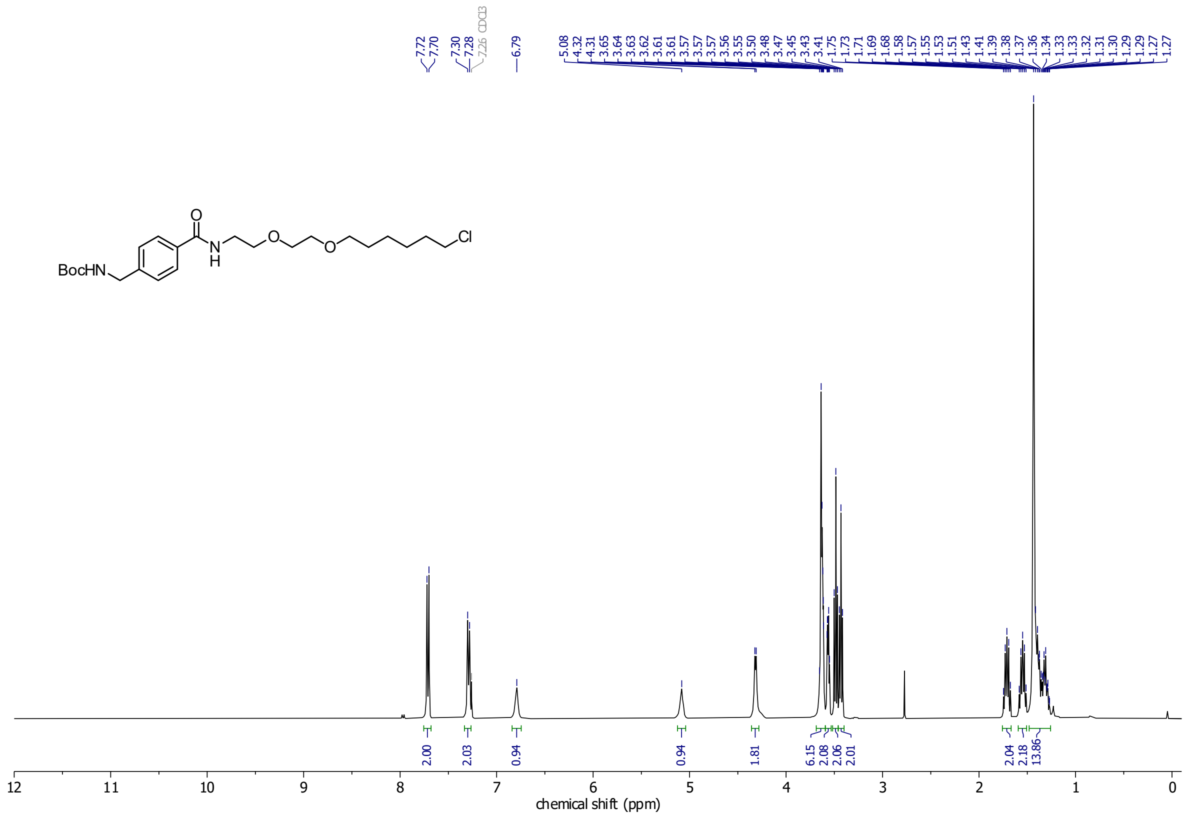

**$^{13}$C-NMR**

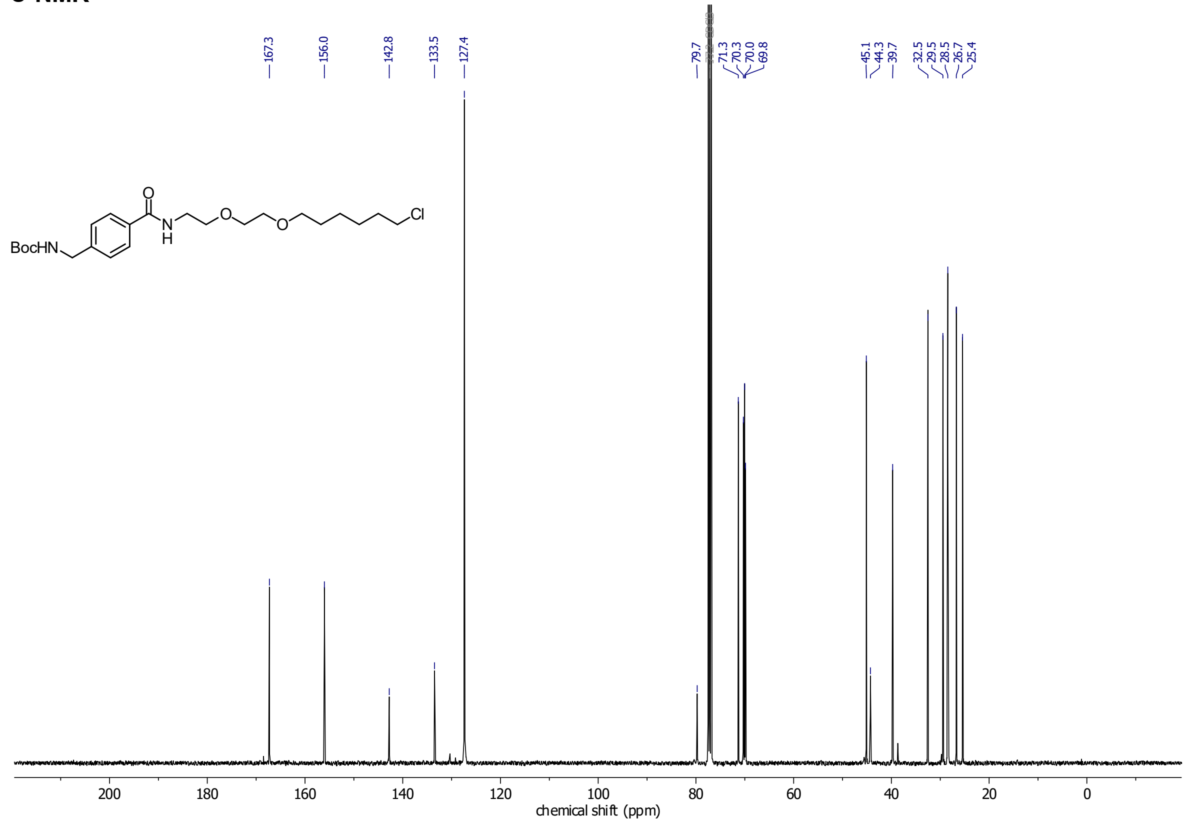

#### CA-Fluo

##### 1H-NMR

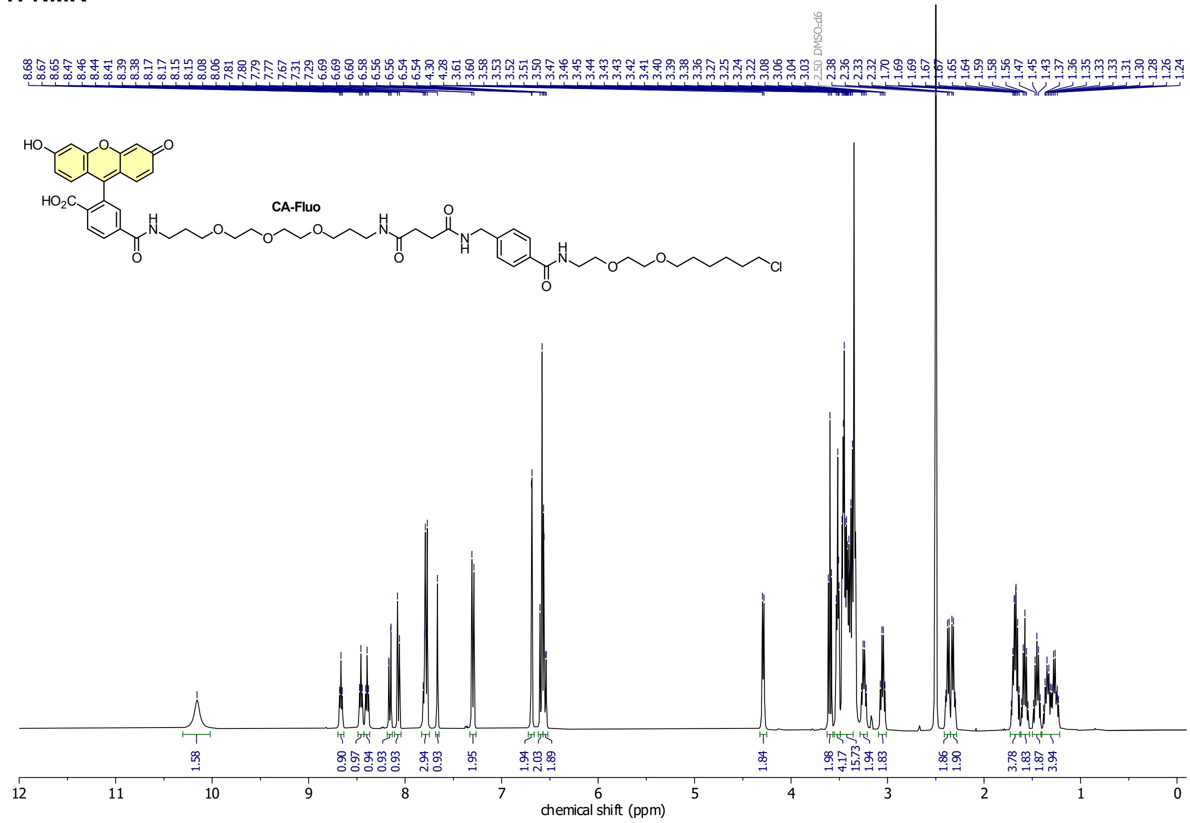

##### 13C-NMR

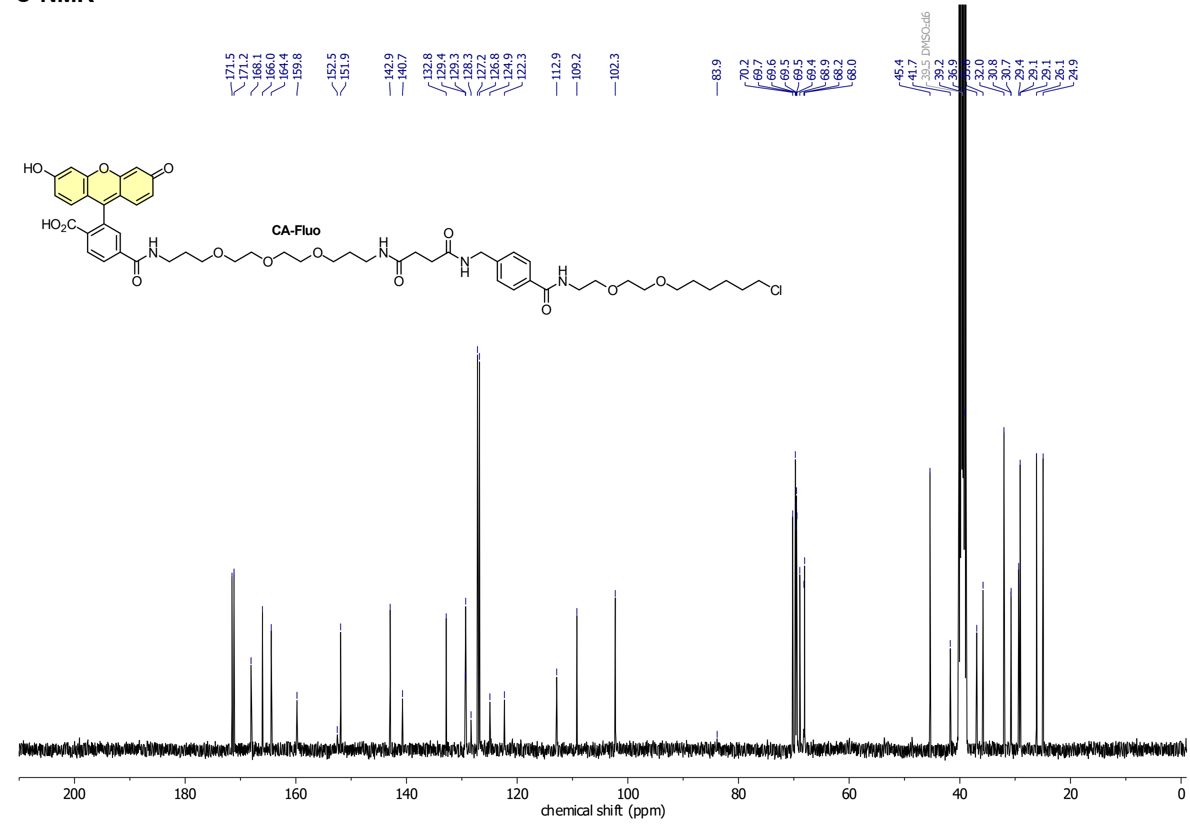

**Compound 33**

**$^{1}$H-NMR**

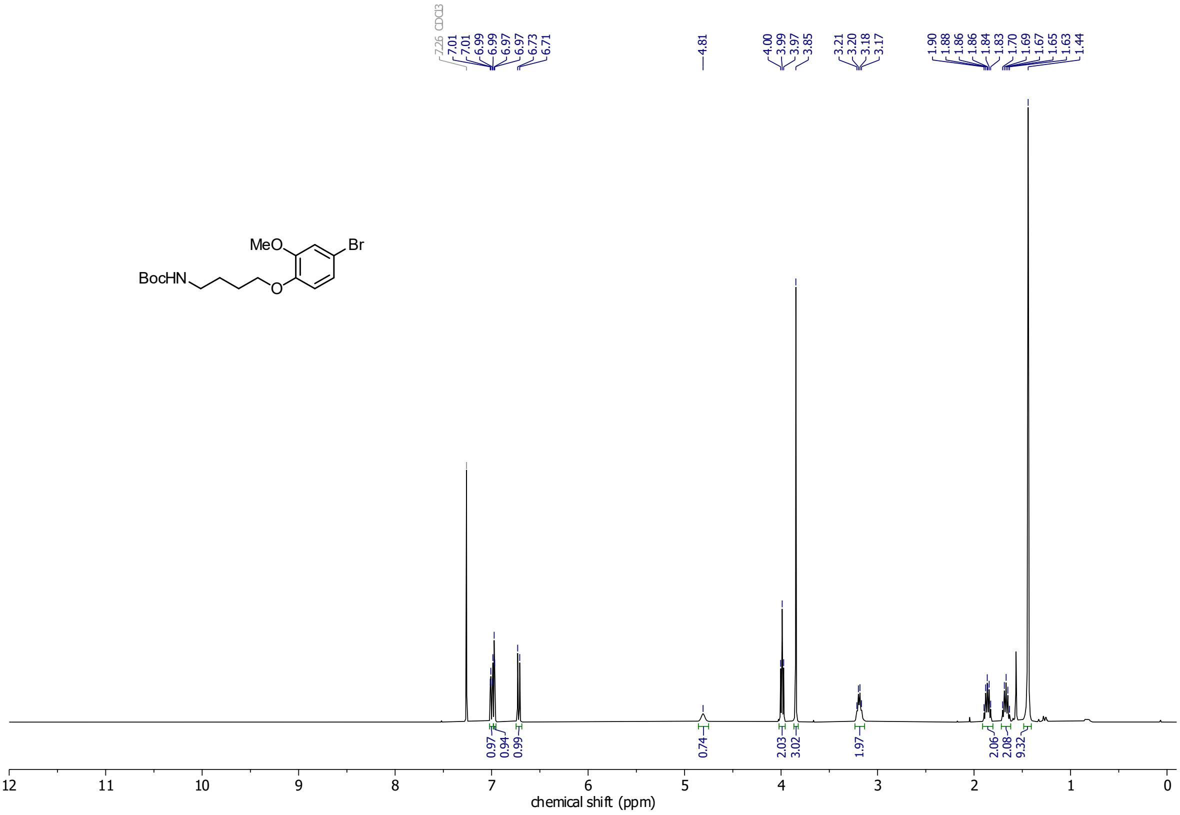

**$^{13}$C-NMR**

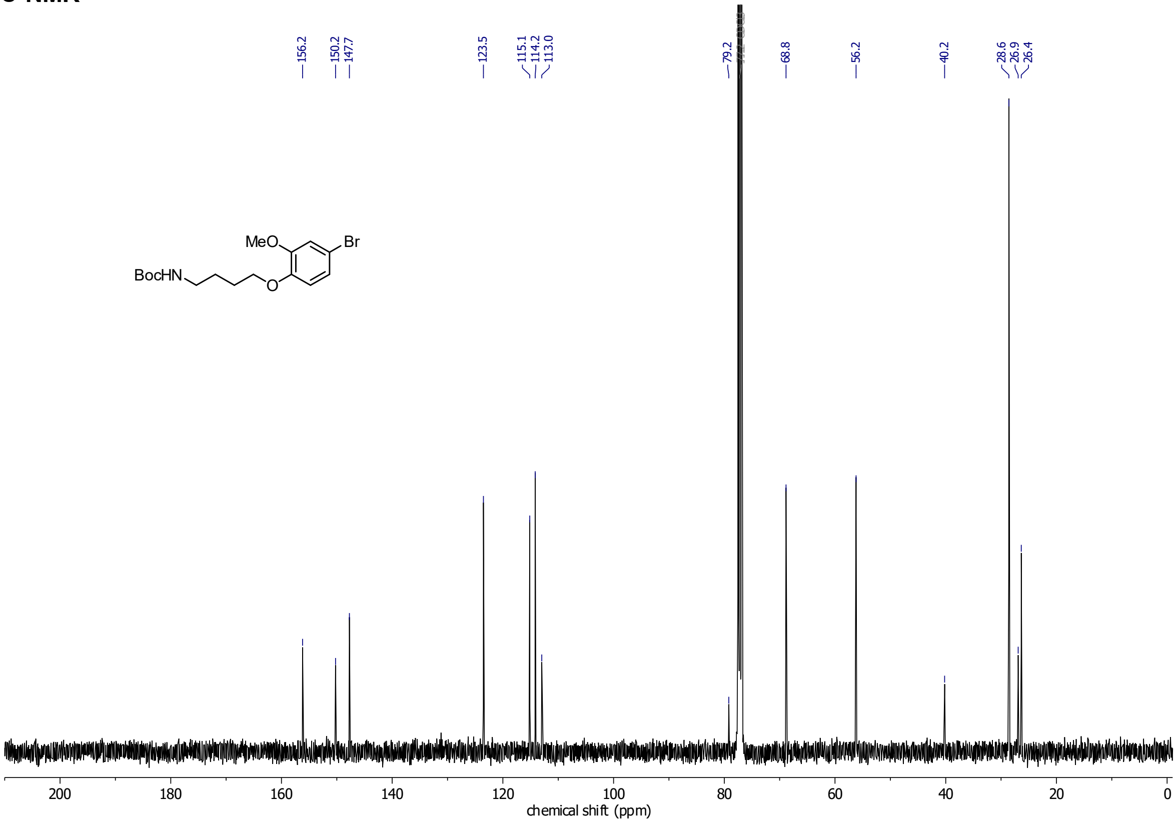

**Compound 35**

**$^1$H-NMR**

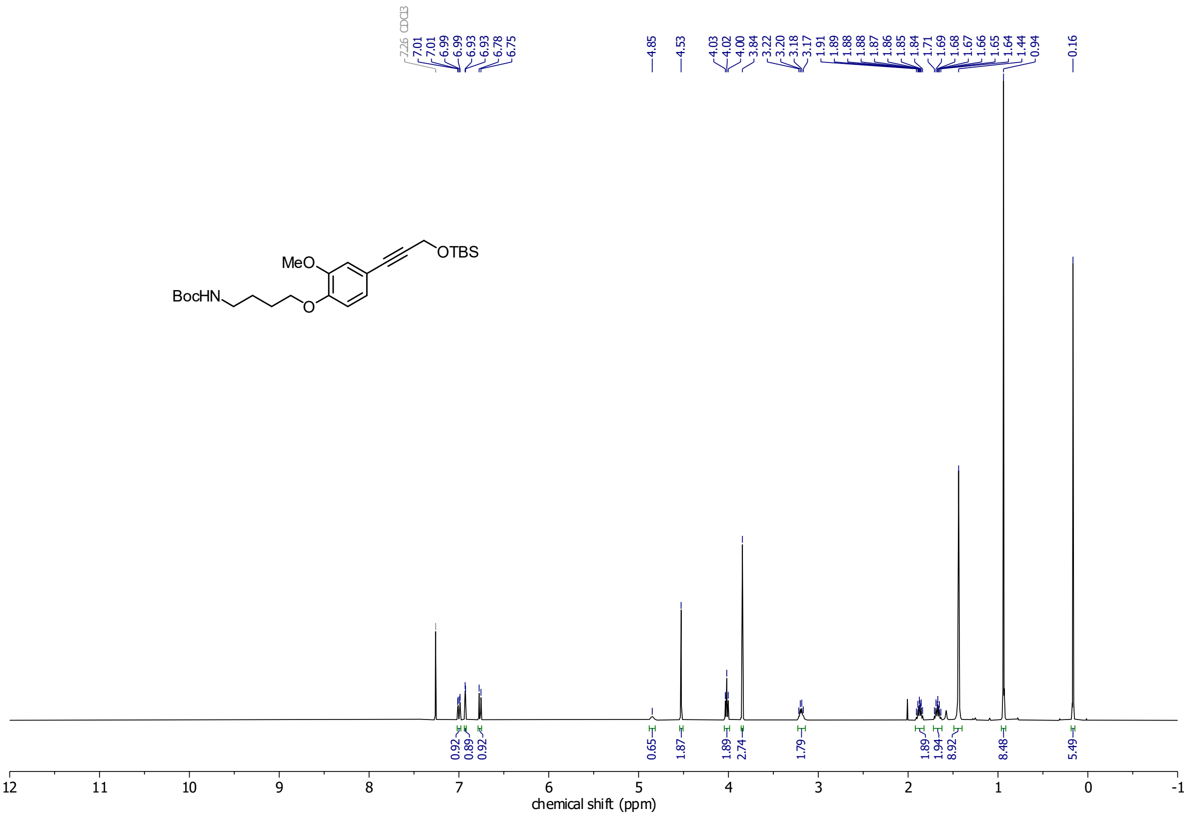

**$^{13}$C-NMR**

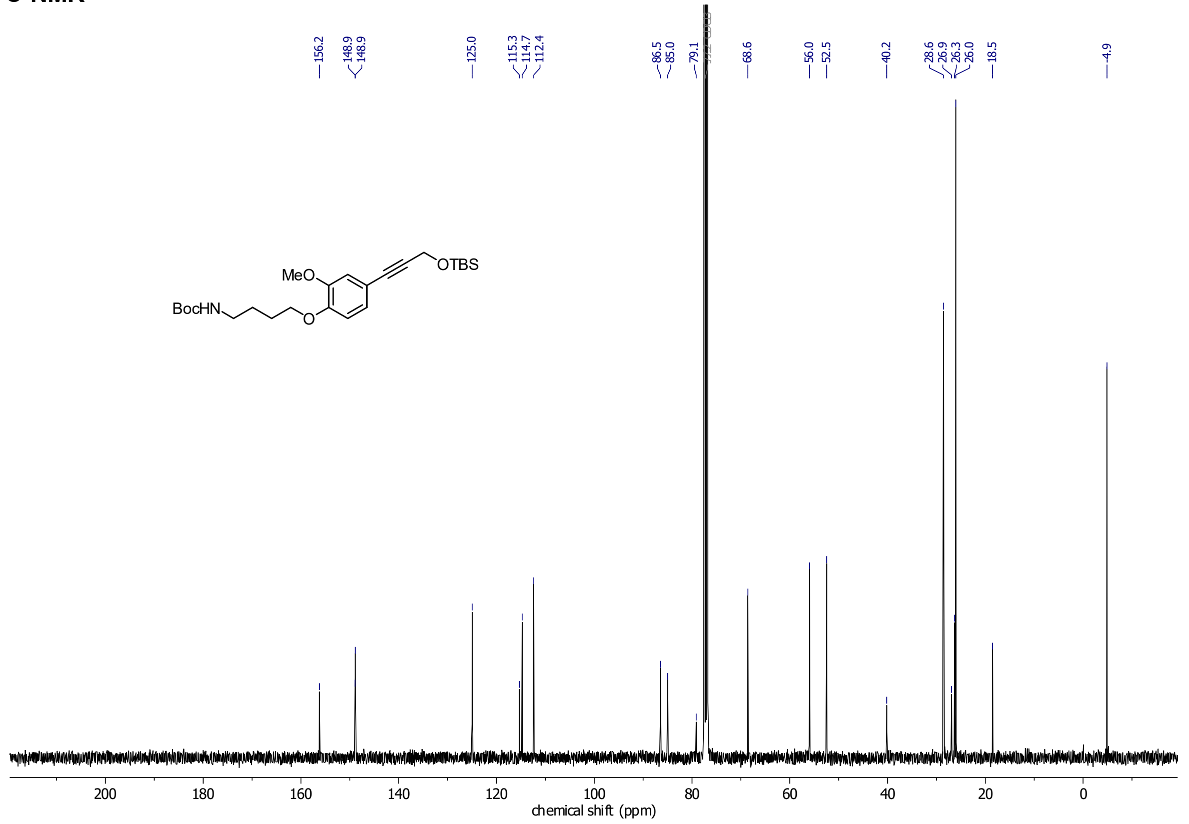

#### Compound 38

##### $^{1}$H-NMR

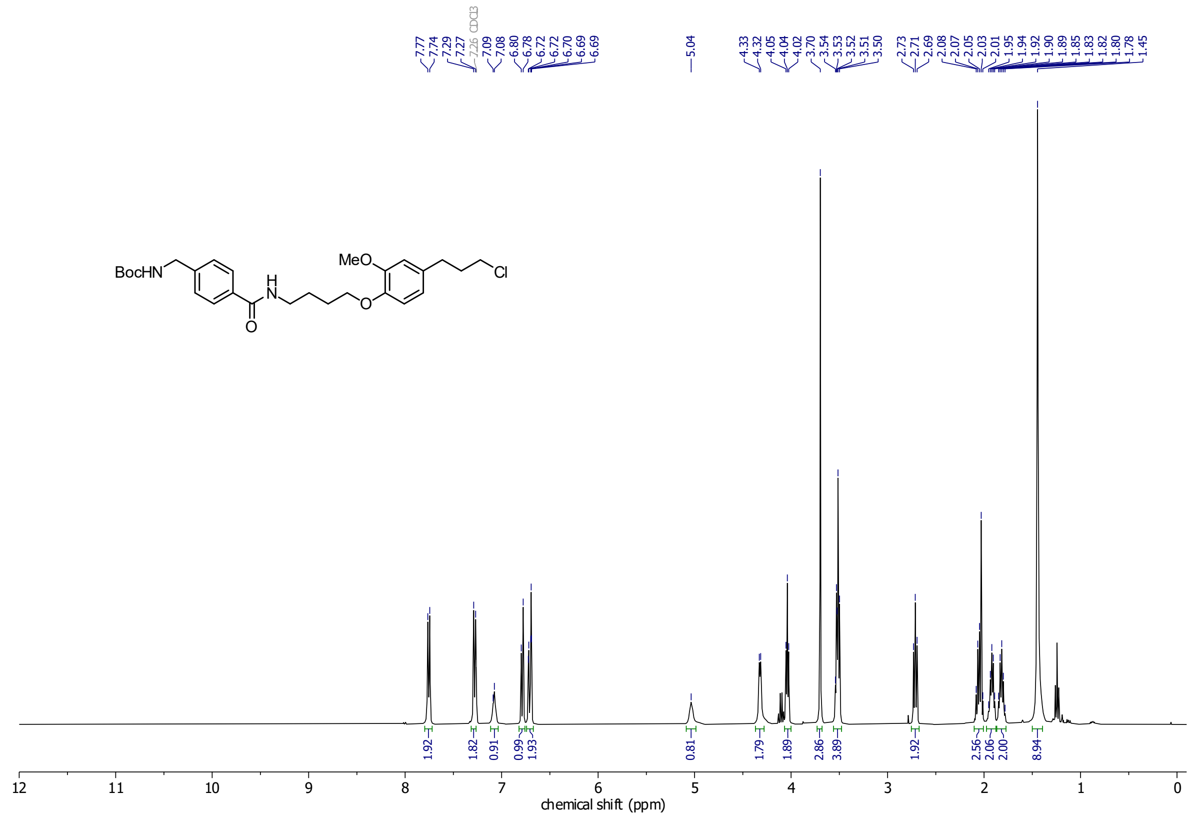

##### $^{13}$C-NMR

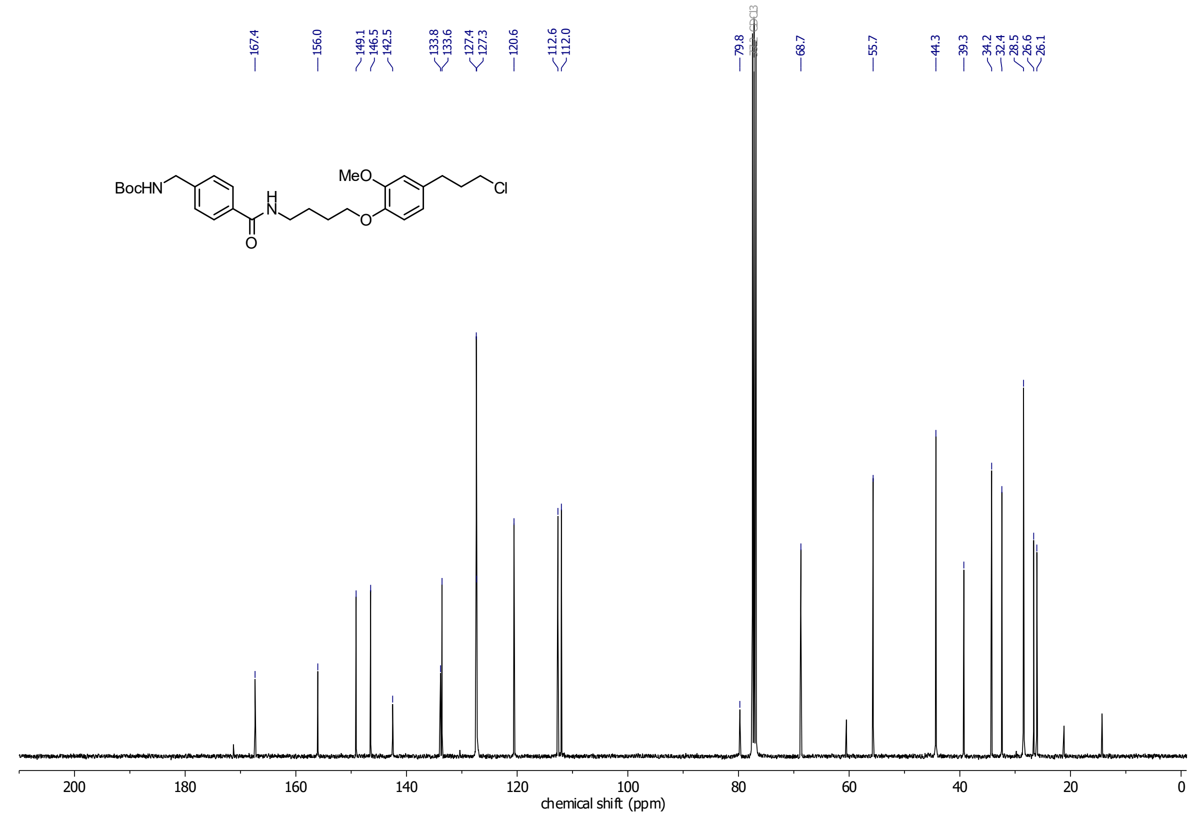

#### HL2$^{O}$-Fluo

##### $^{1}$H-NMR

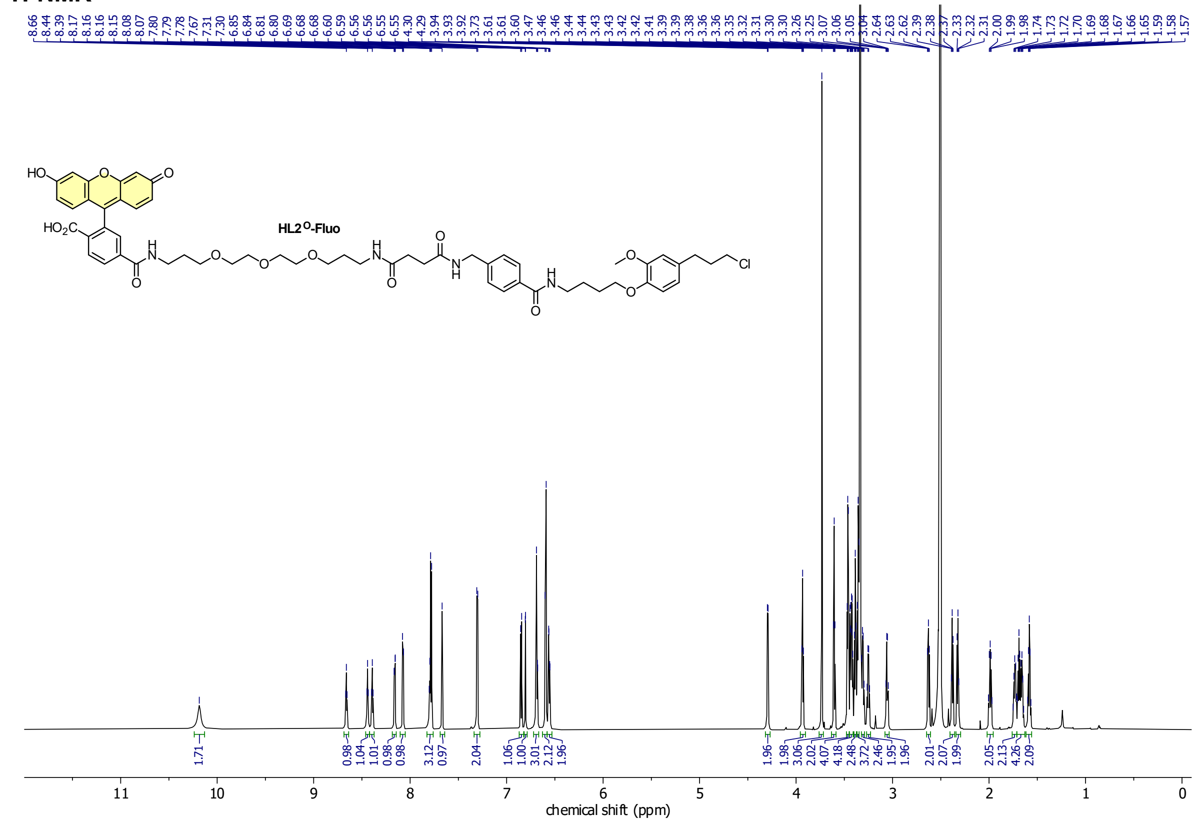

##### $^{13}$C-NMR

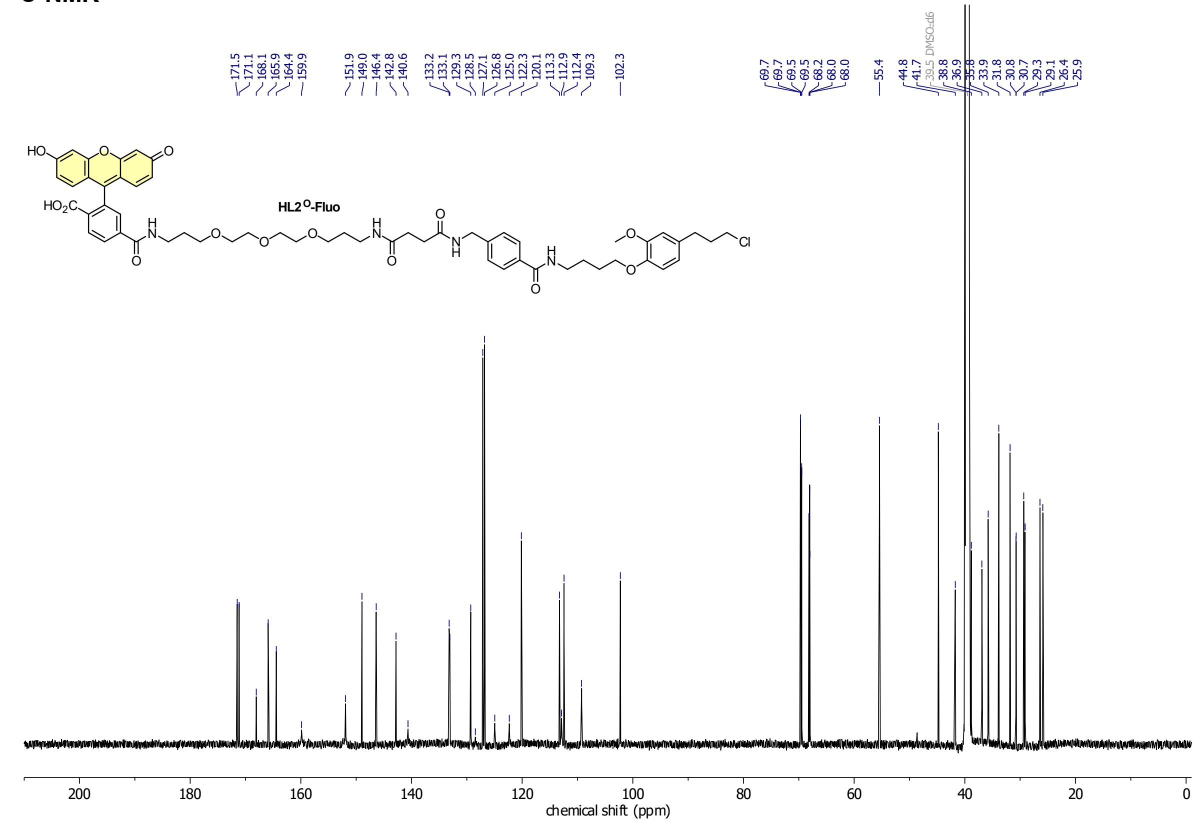

**HSQC**

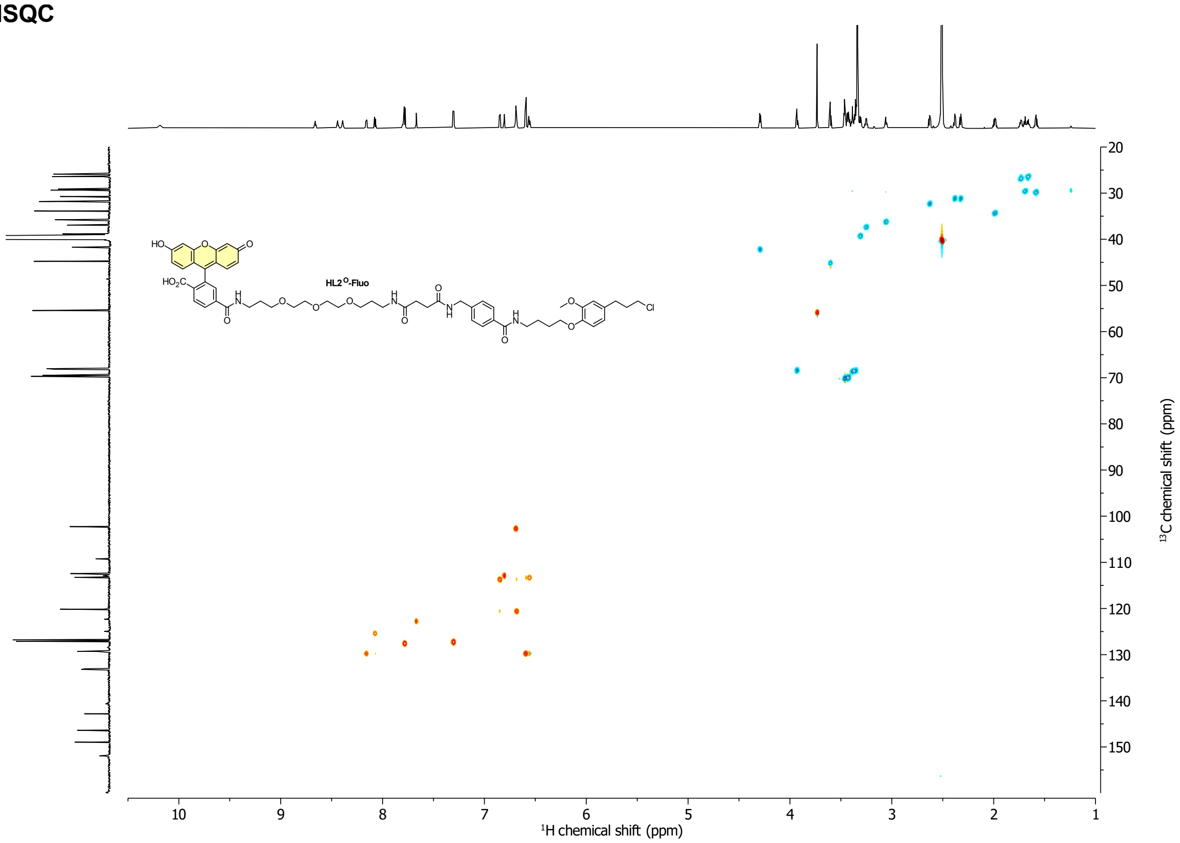

HL2[O]-Fluo
¹H chemical shift (ppm)
¹³C chemical shift (ppm)

**HMBC**

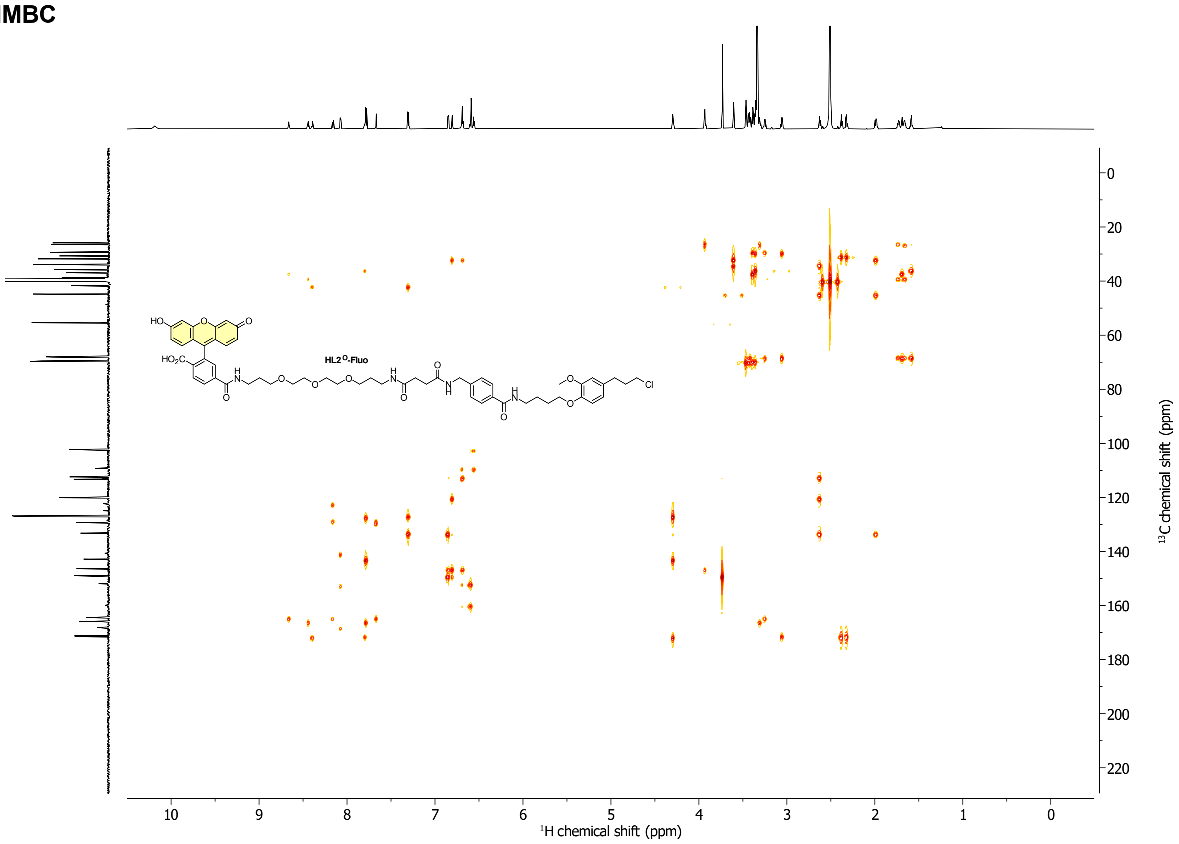

HL2[O]-Fluo
¹H chemical shift (ppm)
¹³C chemical shift (ppm)

**Compound 40**

**[1]H-NMR**

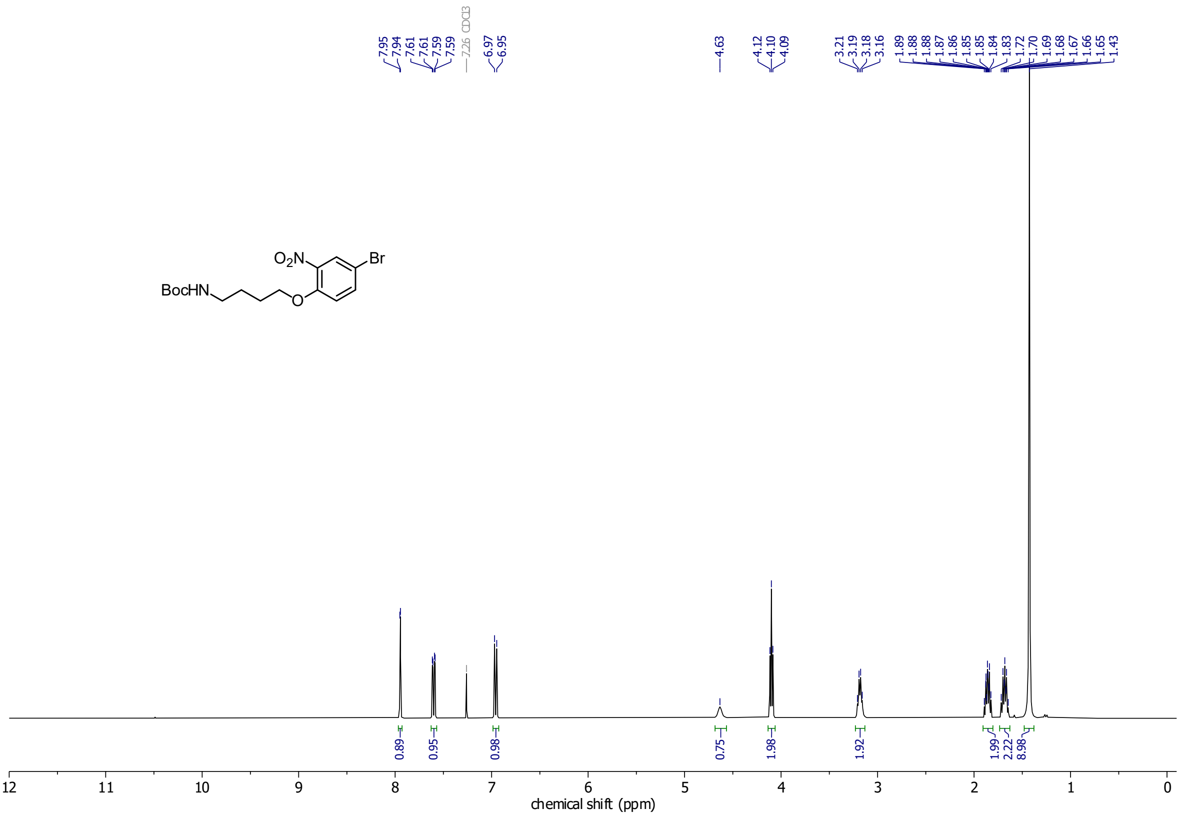

**[13]C-NMR**

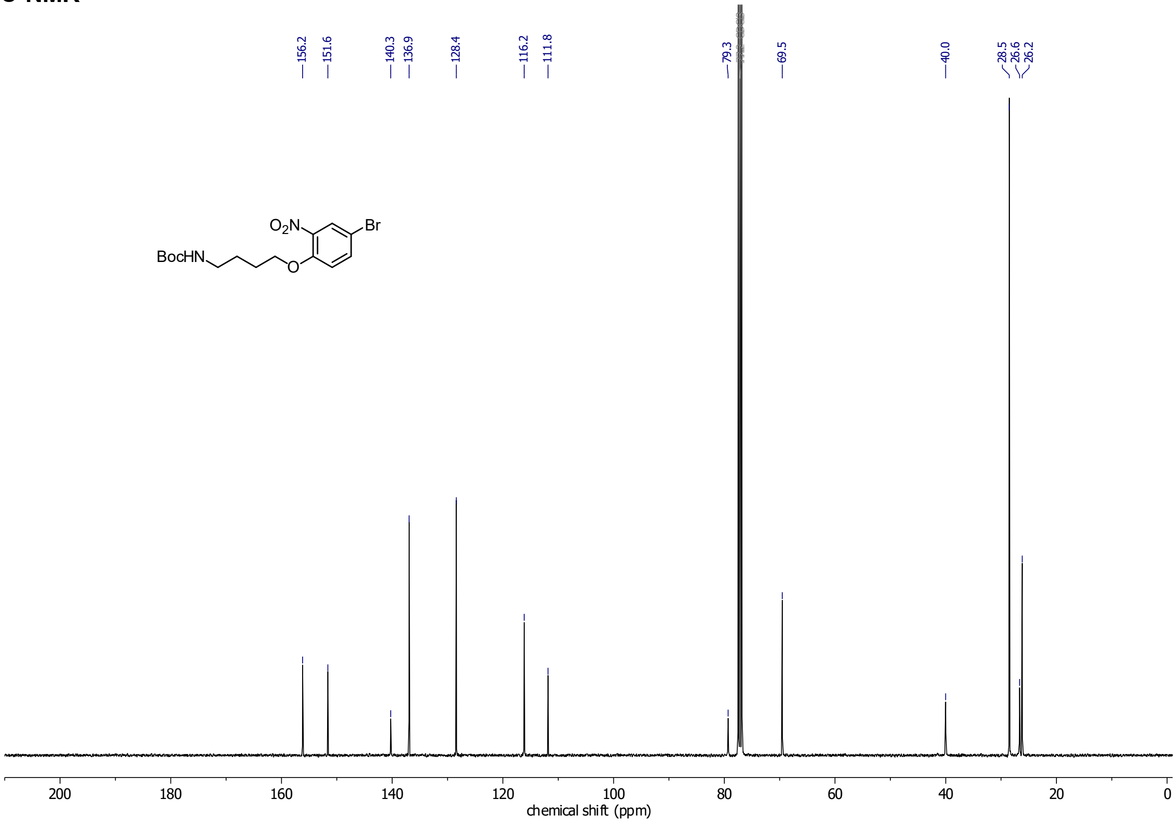

**Compound 41**

**$^{1}$H-NMR**

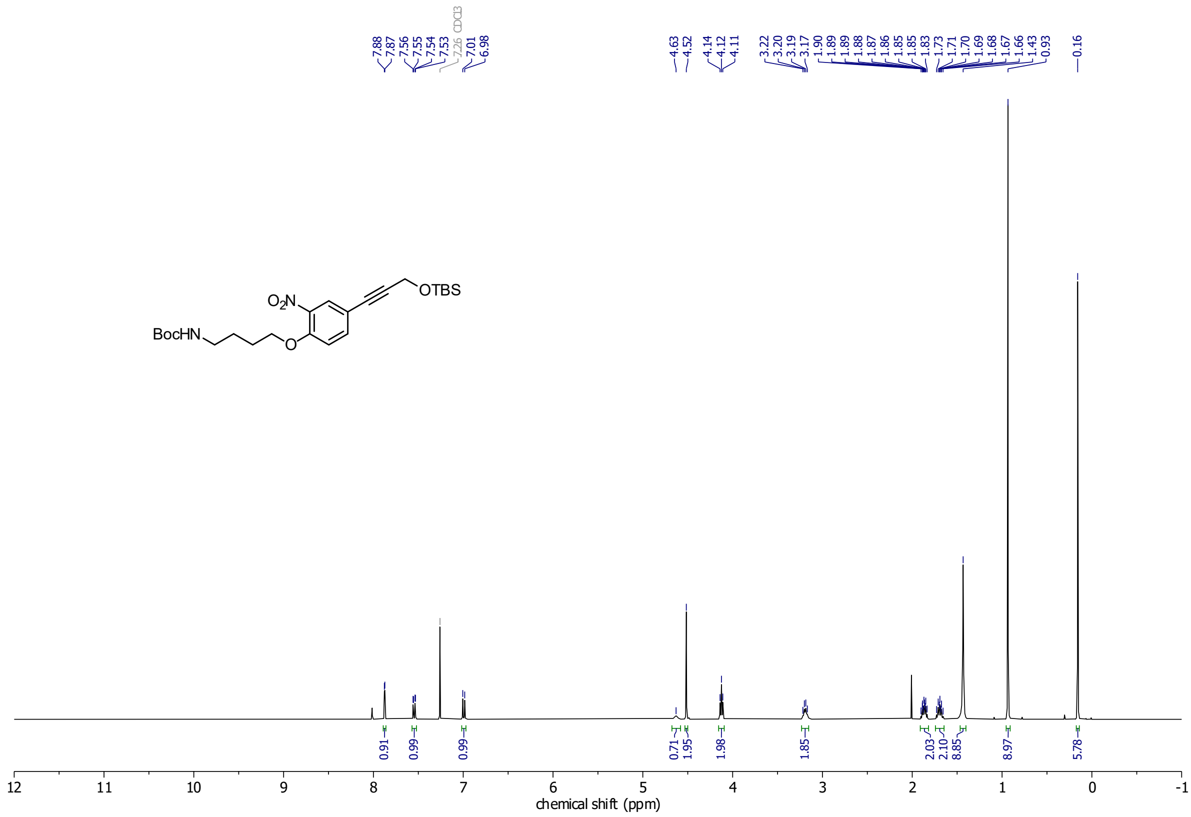

**$^{13}$C-NMR**

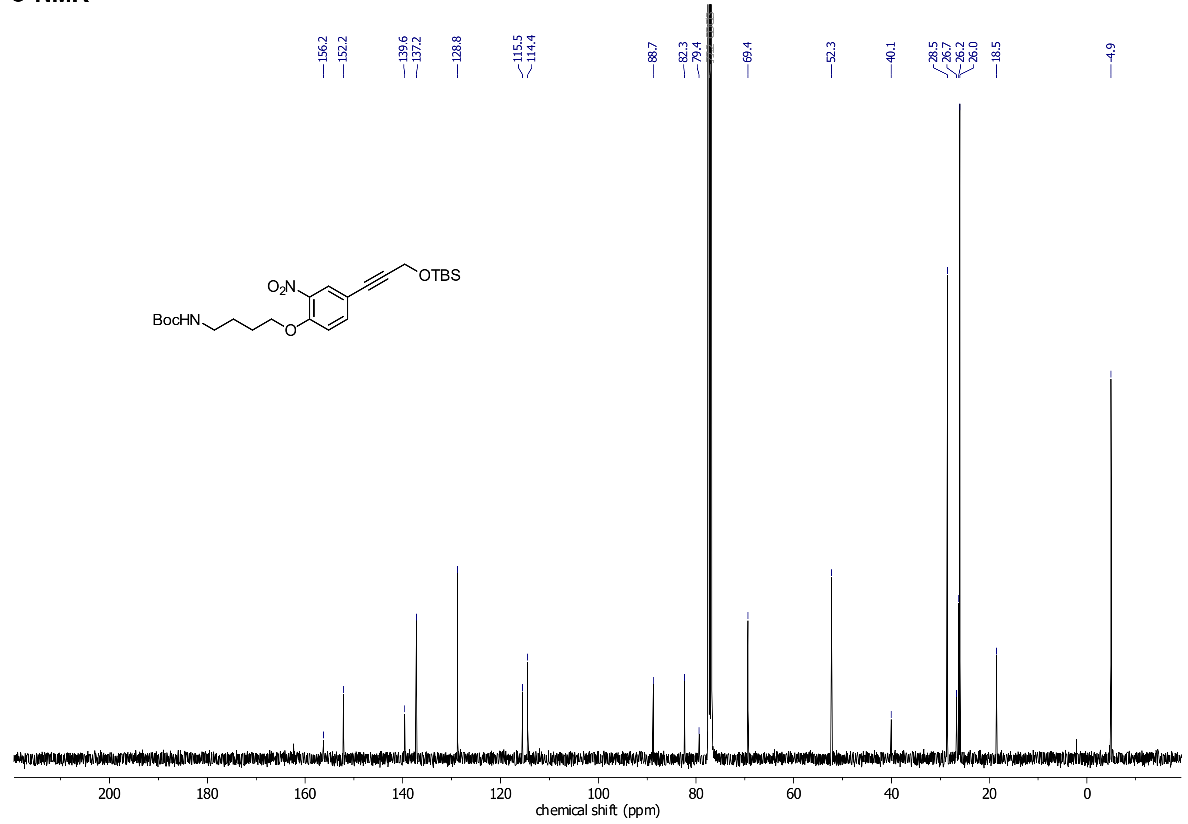

**Compound 42**
**$^1$H-NMR**

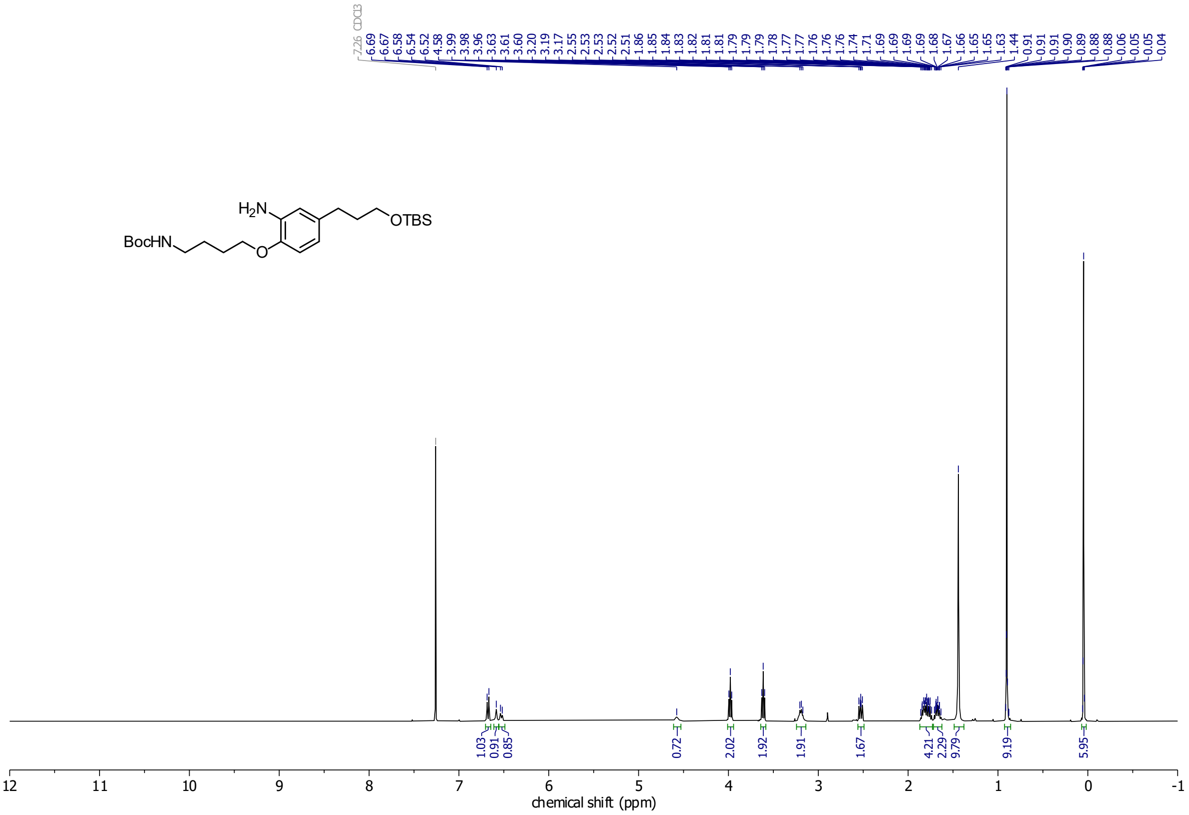

**$^{13}$C-NMR**

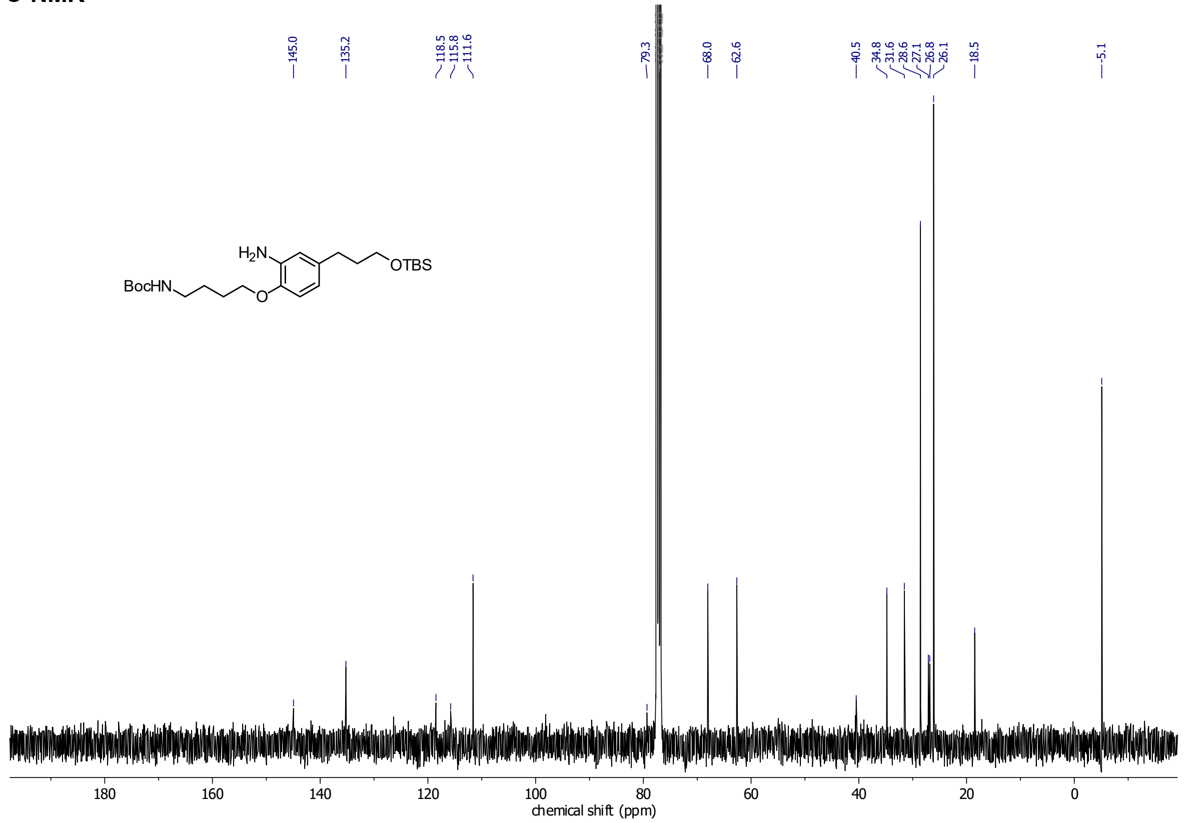

**Compound 44**
**[1]H-NMR**

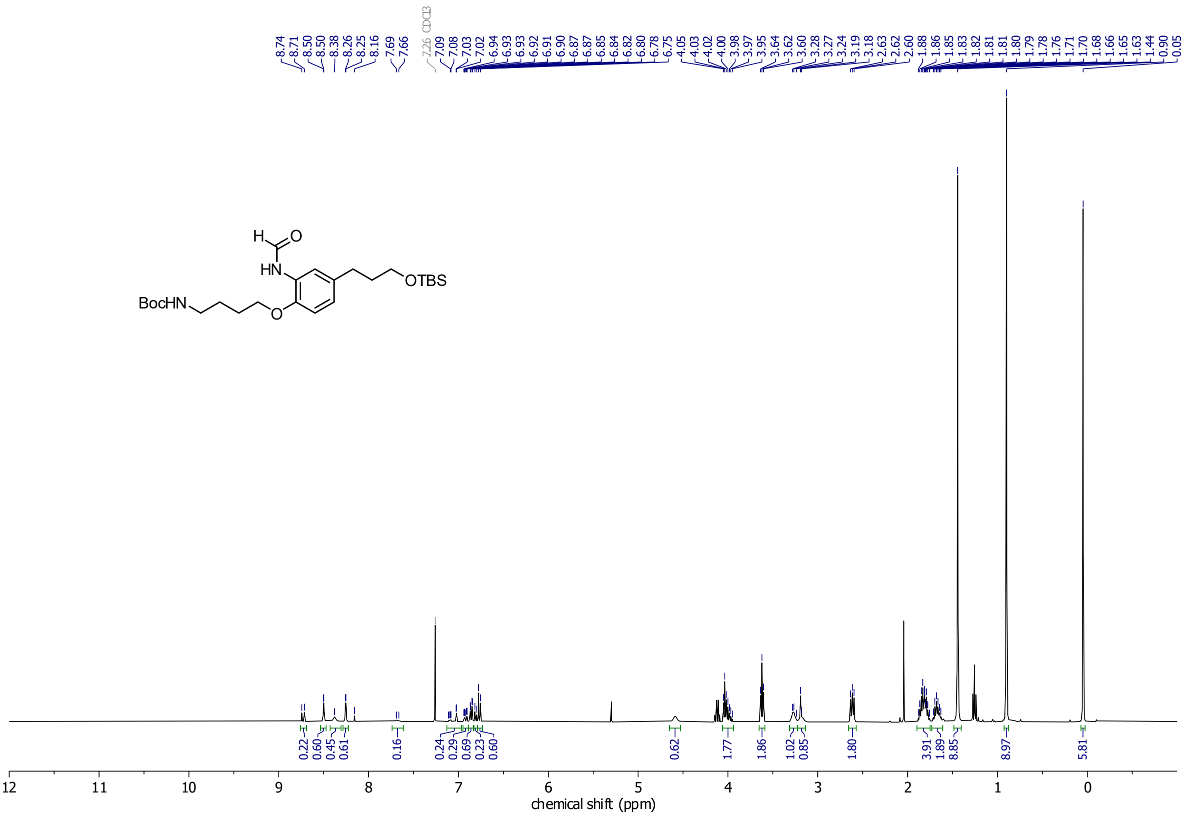

**Compound 48**

**$^1$H-NMR**

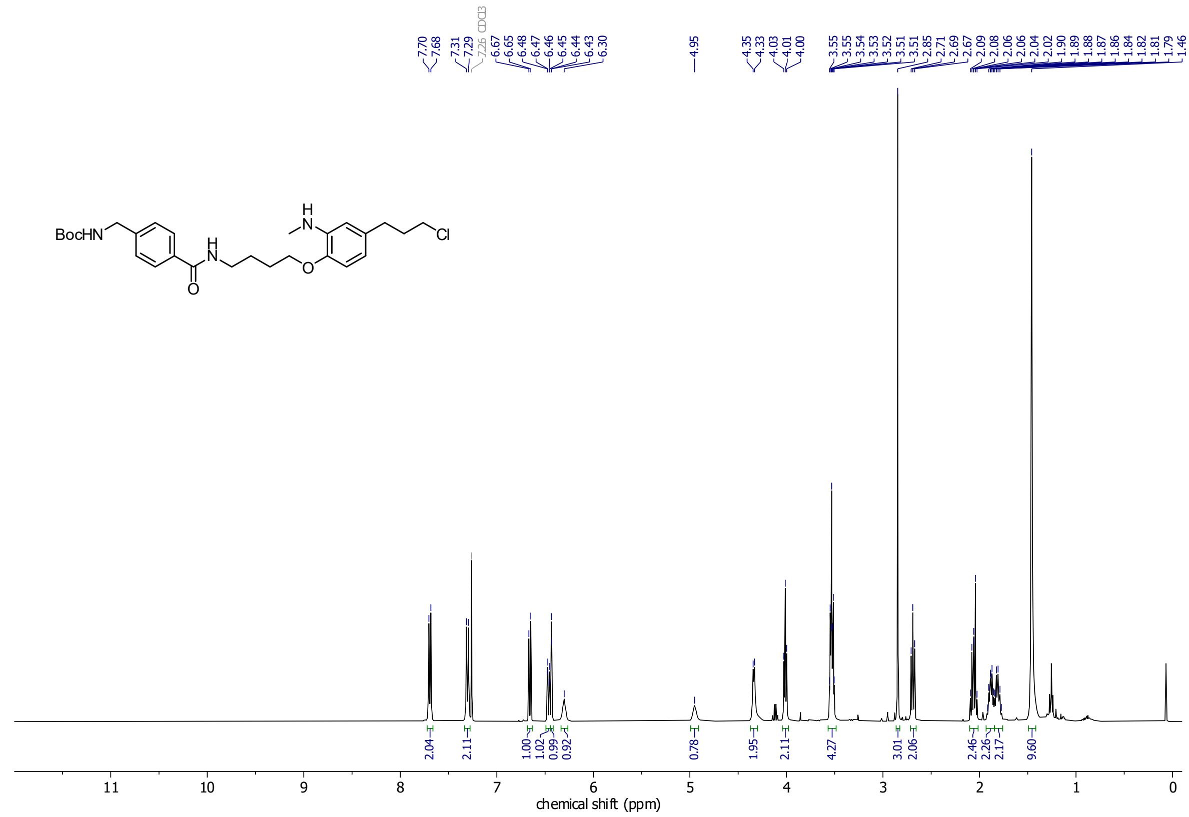

**$^{13}$C-NMR**

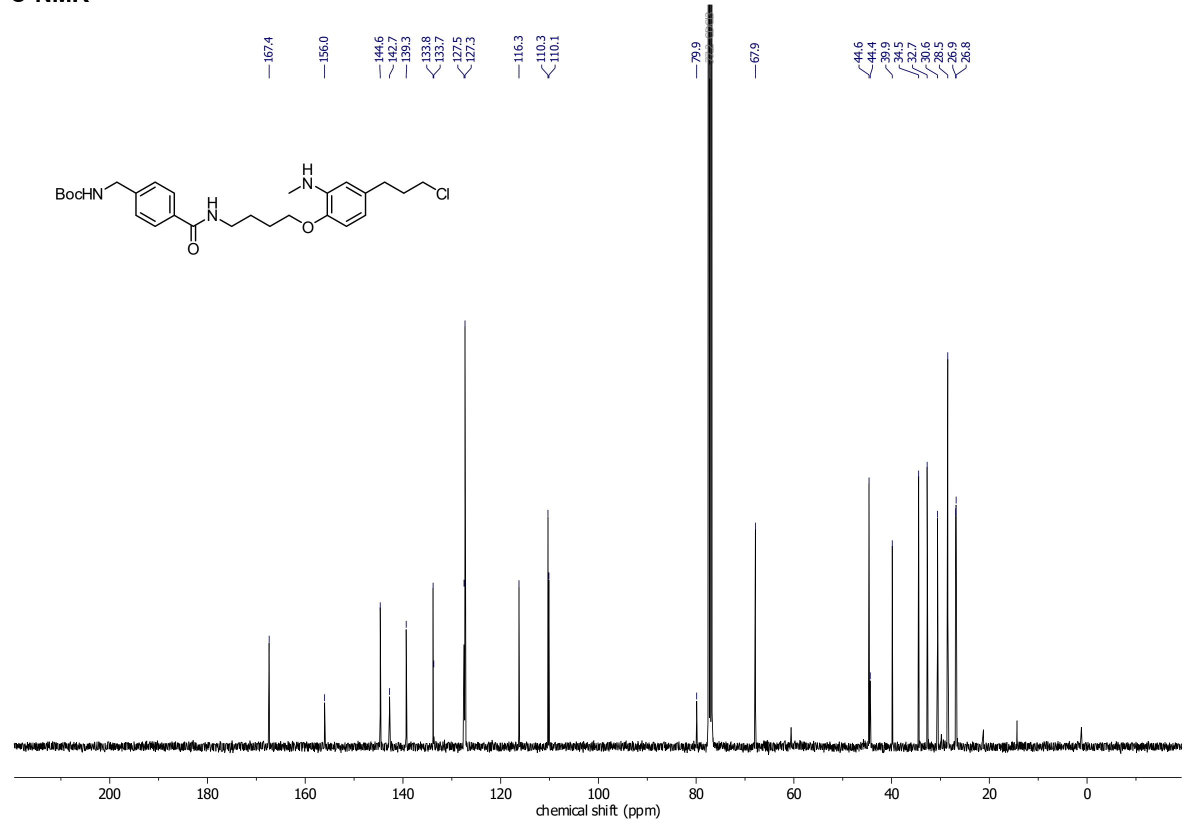

**HL2^N-Fluo**

**¹H-NMR**

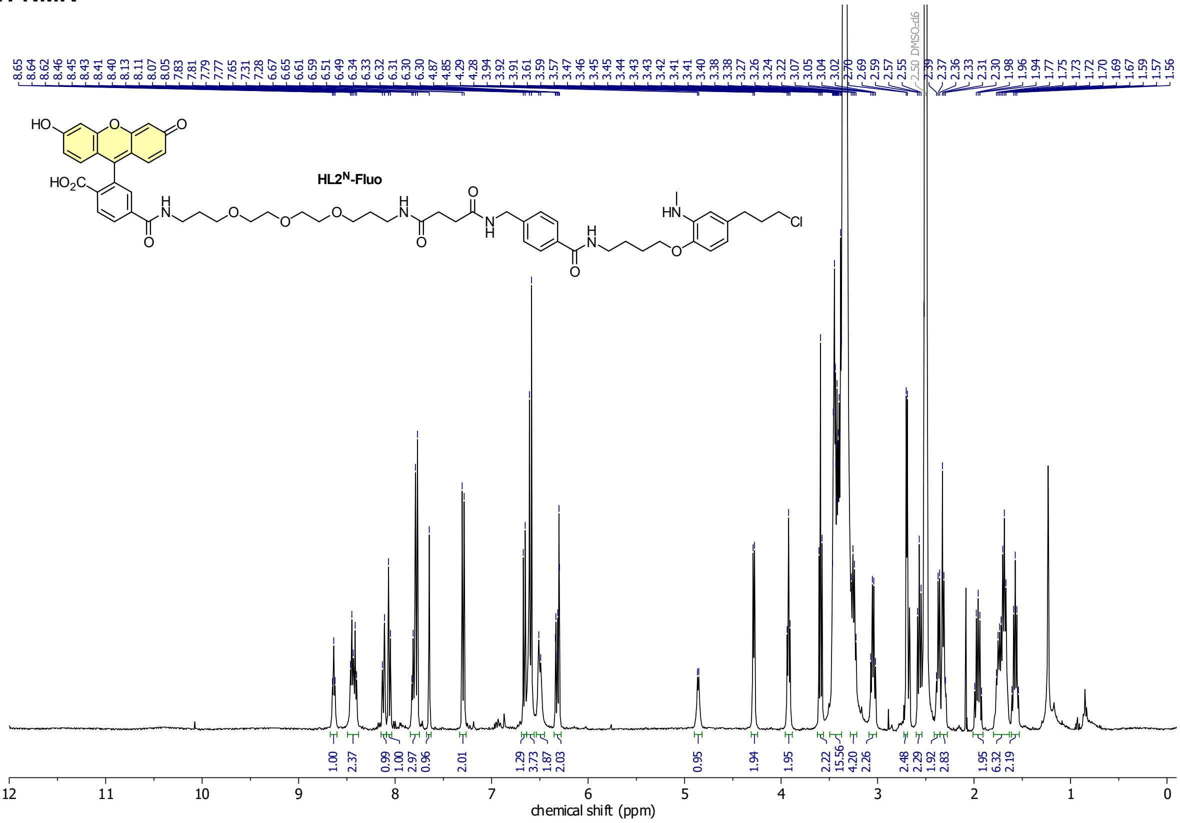

**Compound 49**

**$^1$H-NMR**

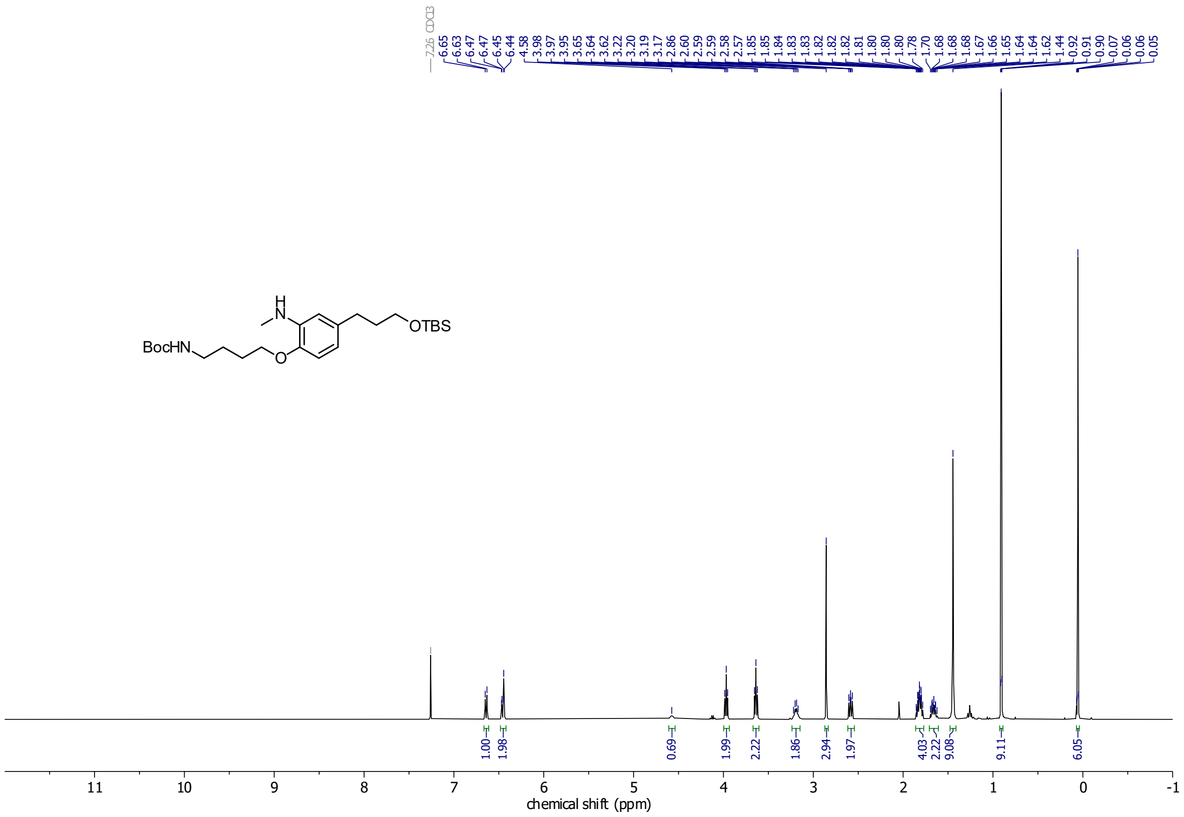

**$^{13}$C-NMR**

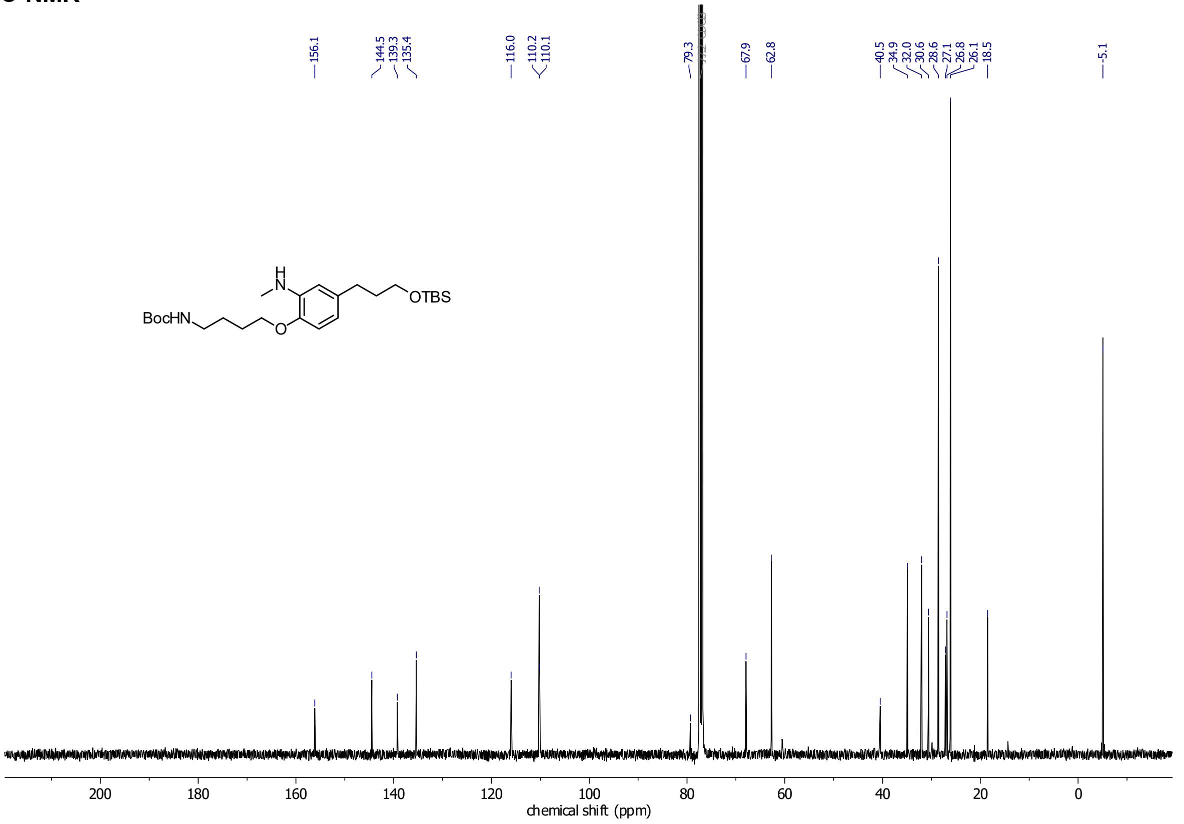

#### Cou-HL2$^{N}$-Fluo

##### $^{1}$H-NMR

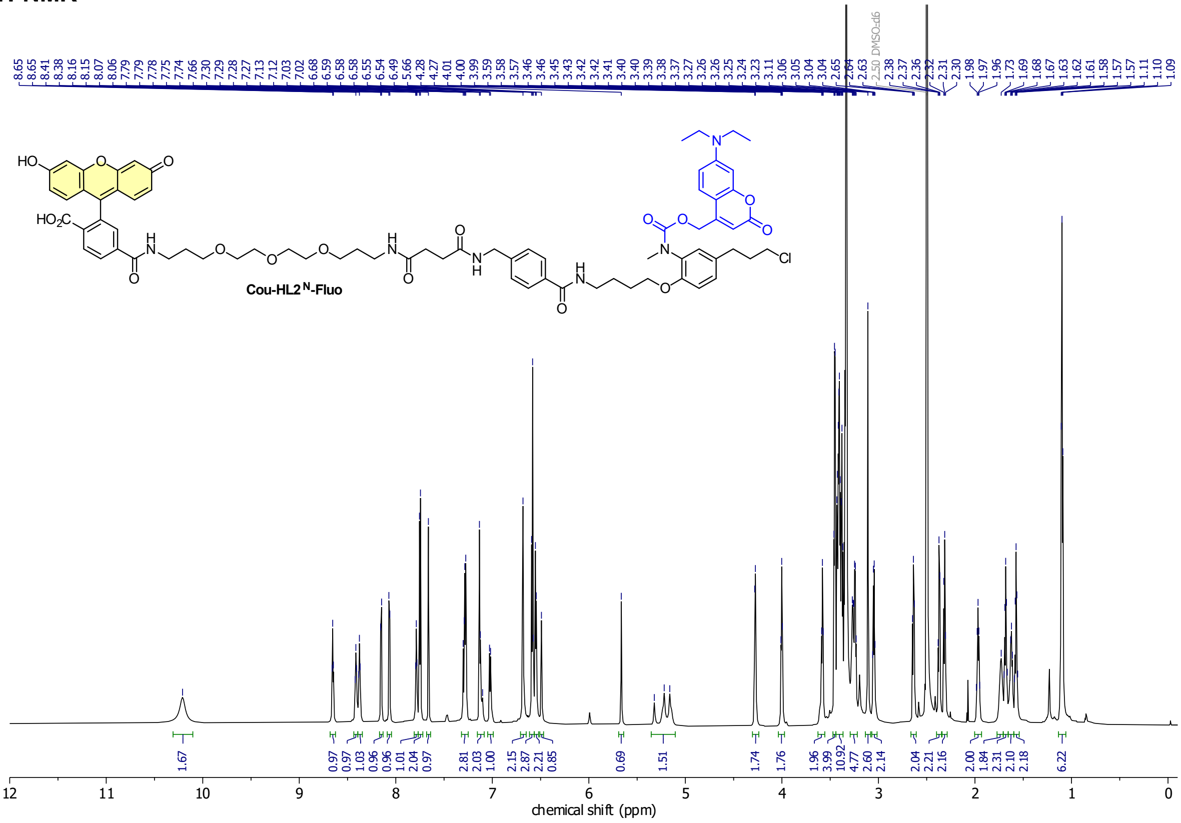

##### $^{13}$C-NMR

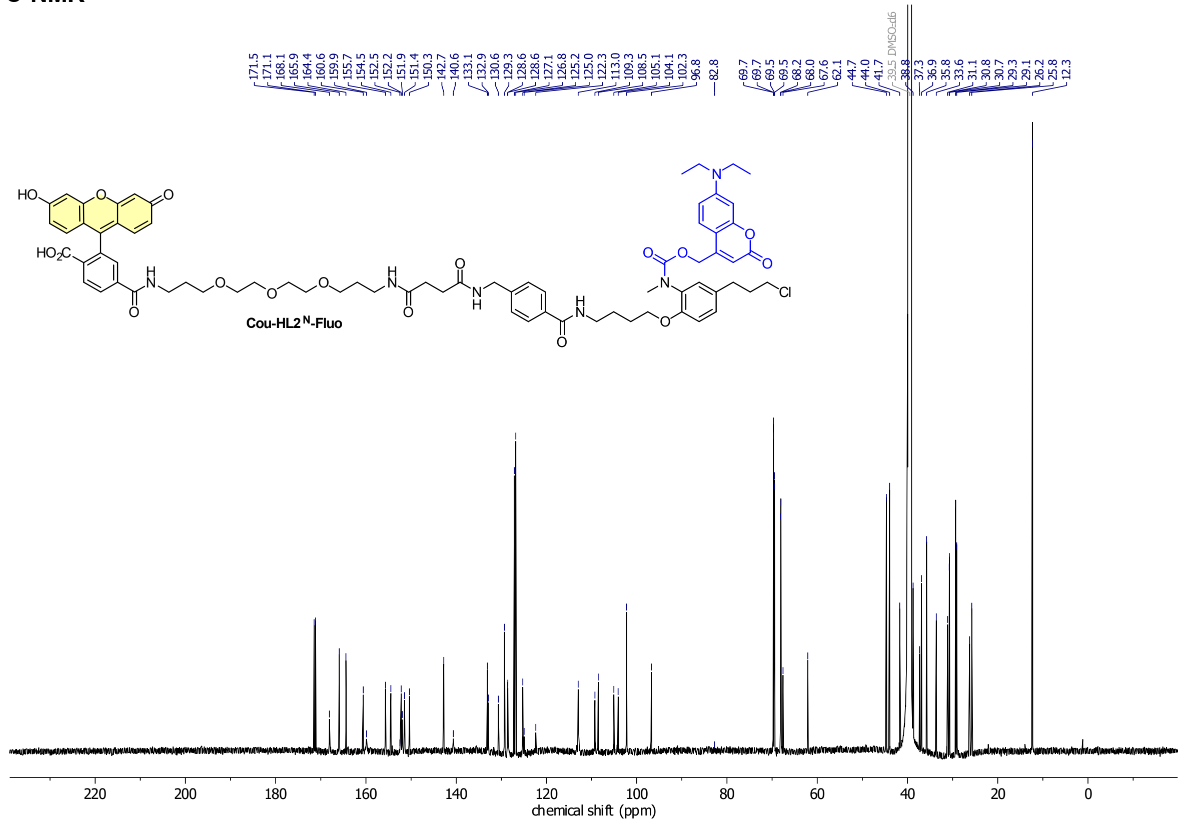

#### HSQC

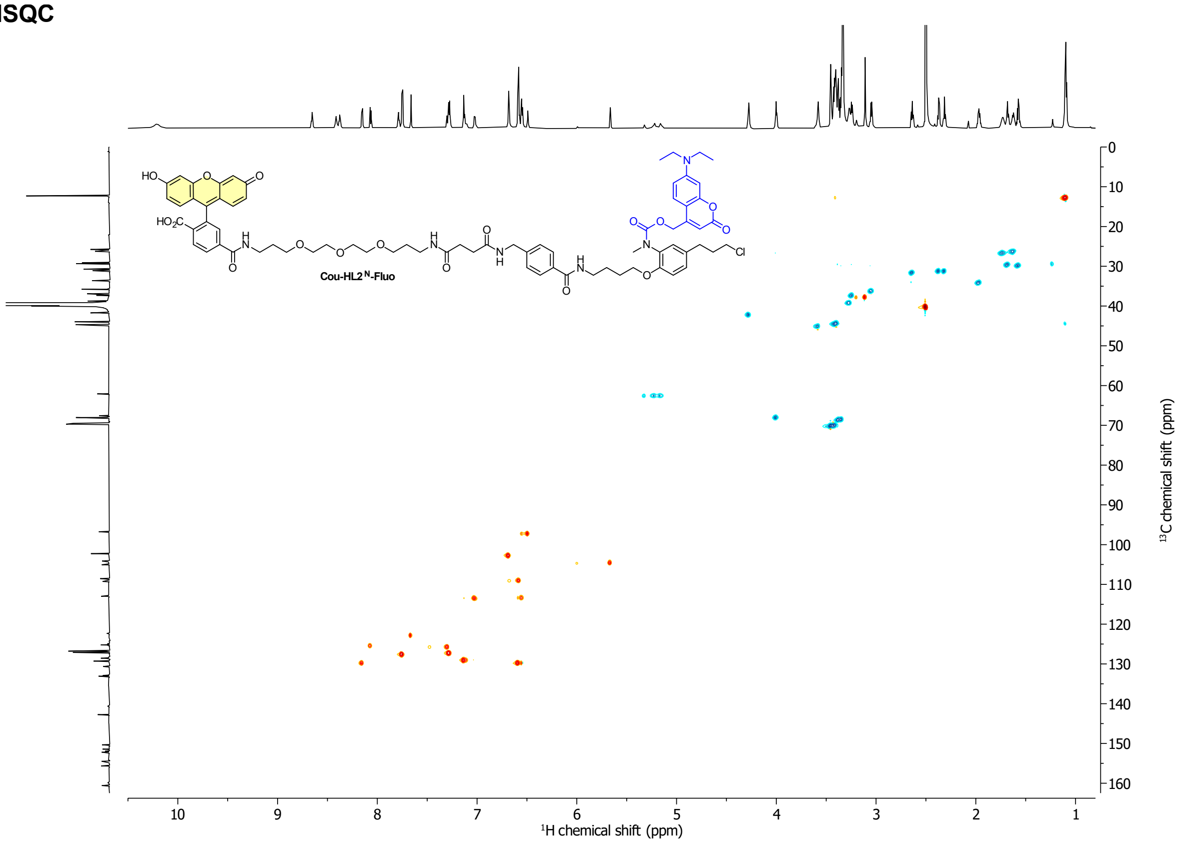

Cou-HL2N-Fluo
¹H chemical shift (ppm)
¹³C chemical shift (ppm)

#### HMBC

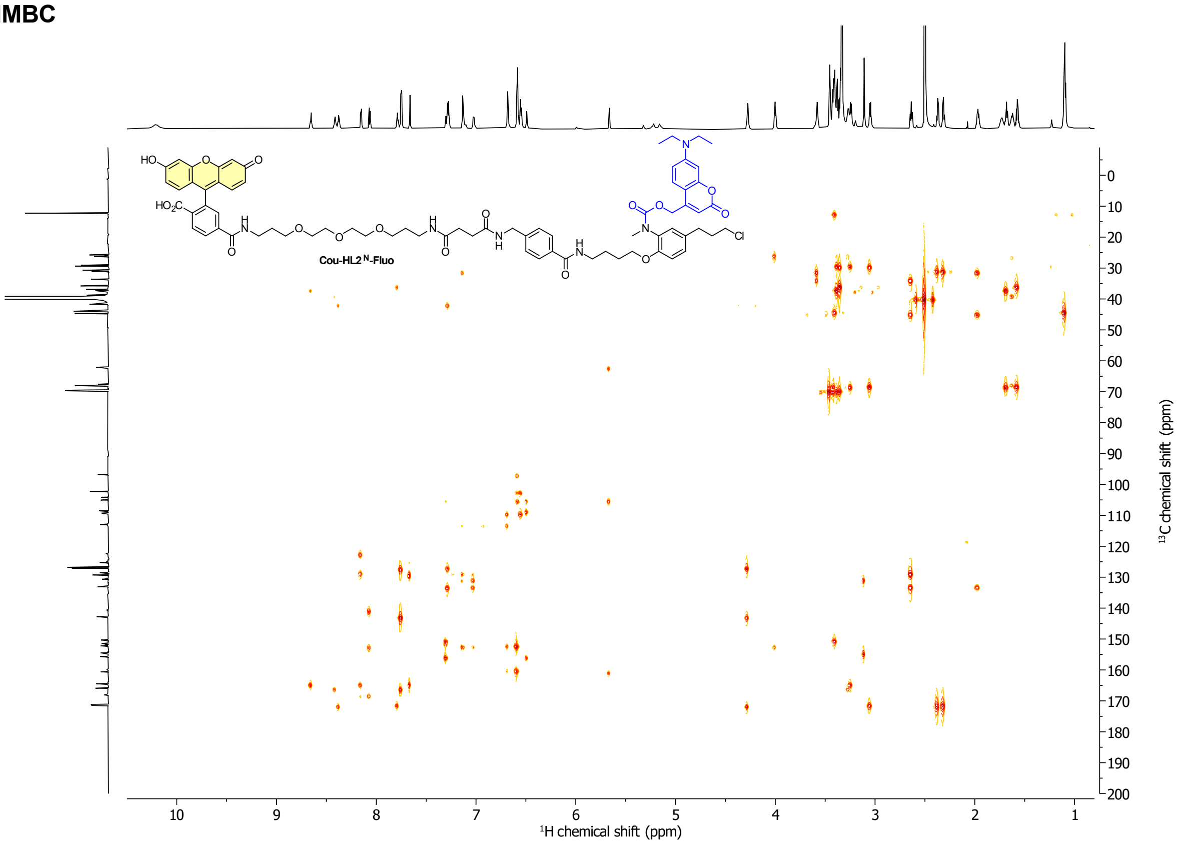

Cou-HL2N-Fluo
¹H chemical shift (ppm)
¹³C chemical shift (ppm)

**Compound 56**

**$^1$H-NMR**

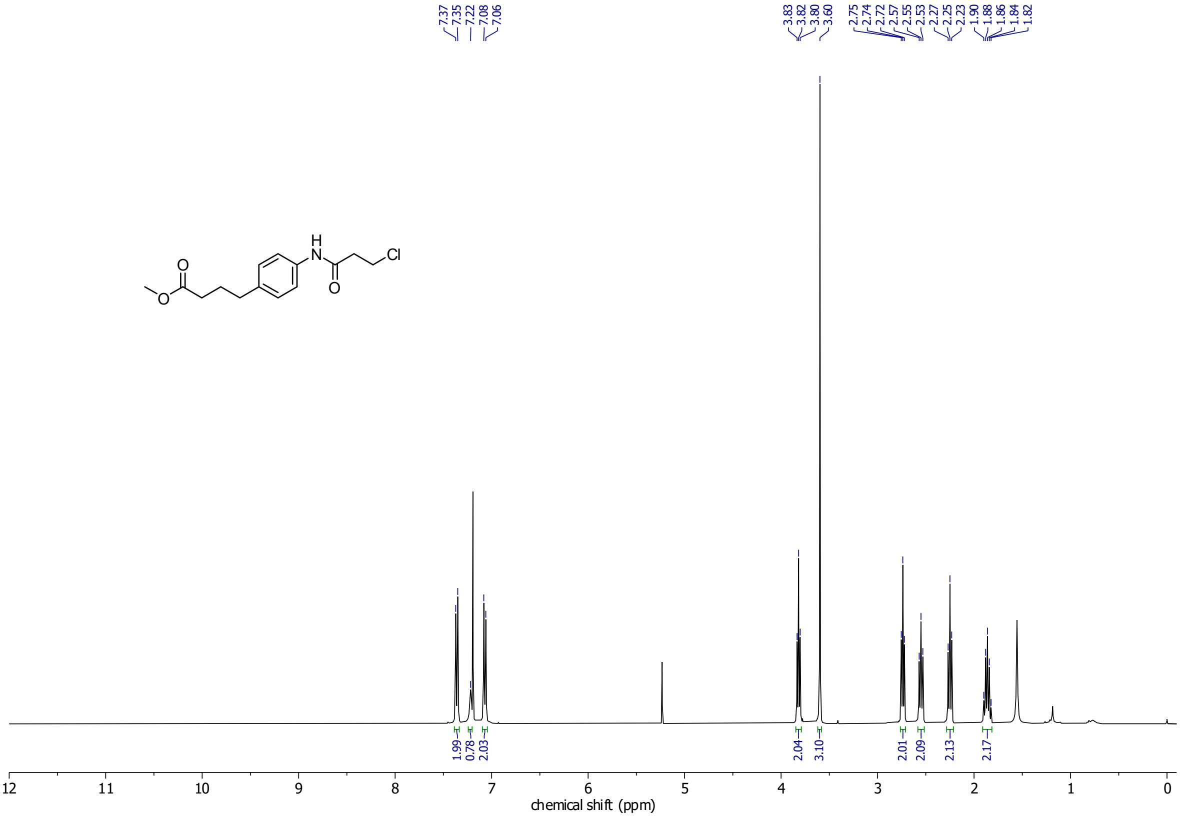

**$^{13}$C-NMR**

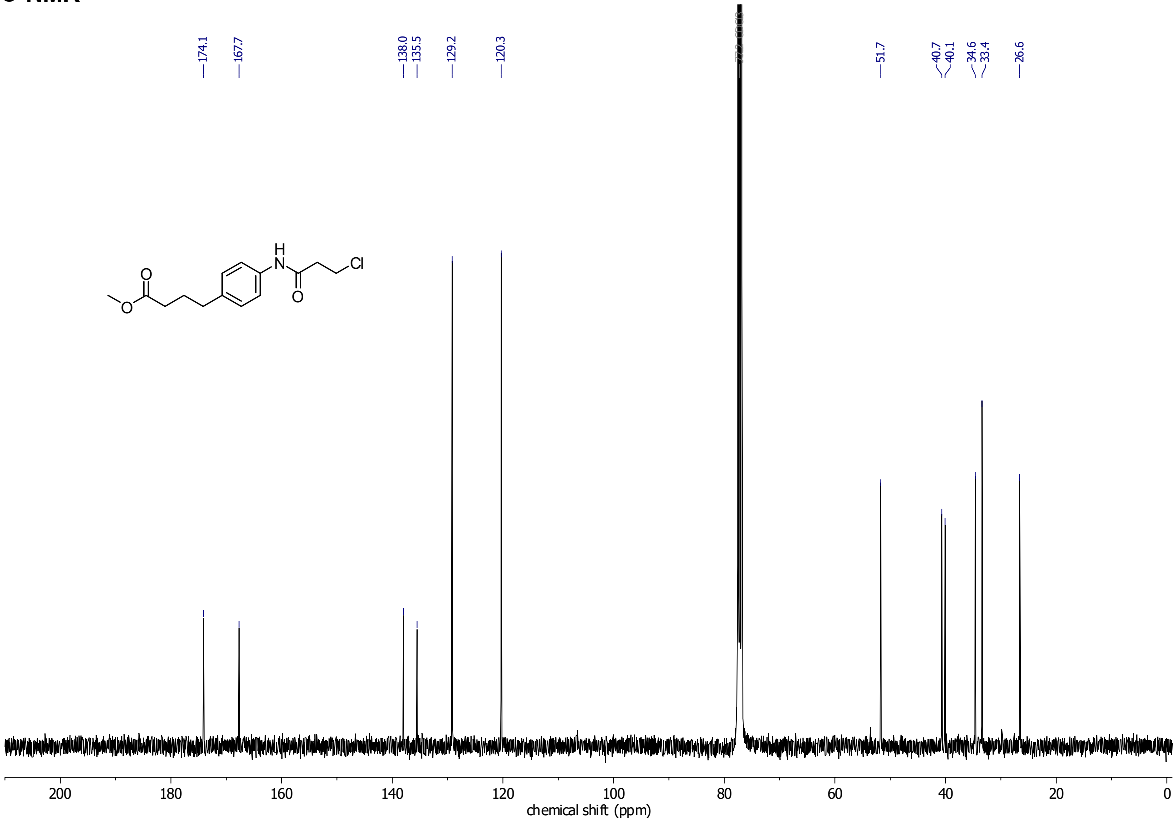

**Compound 60**

**$^{1}$H-NMR**

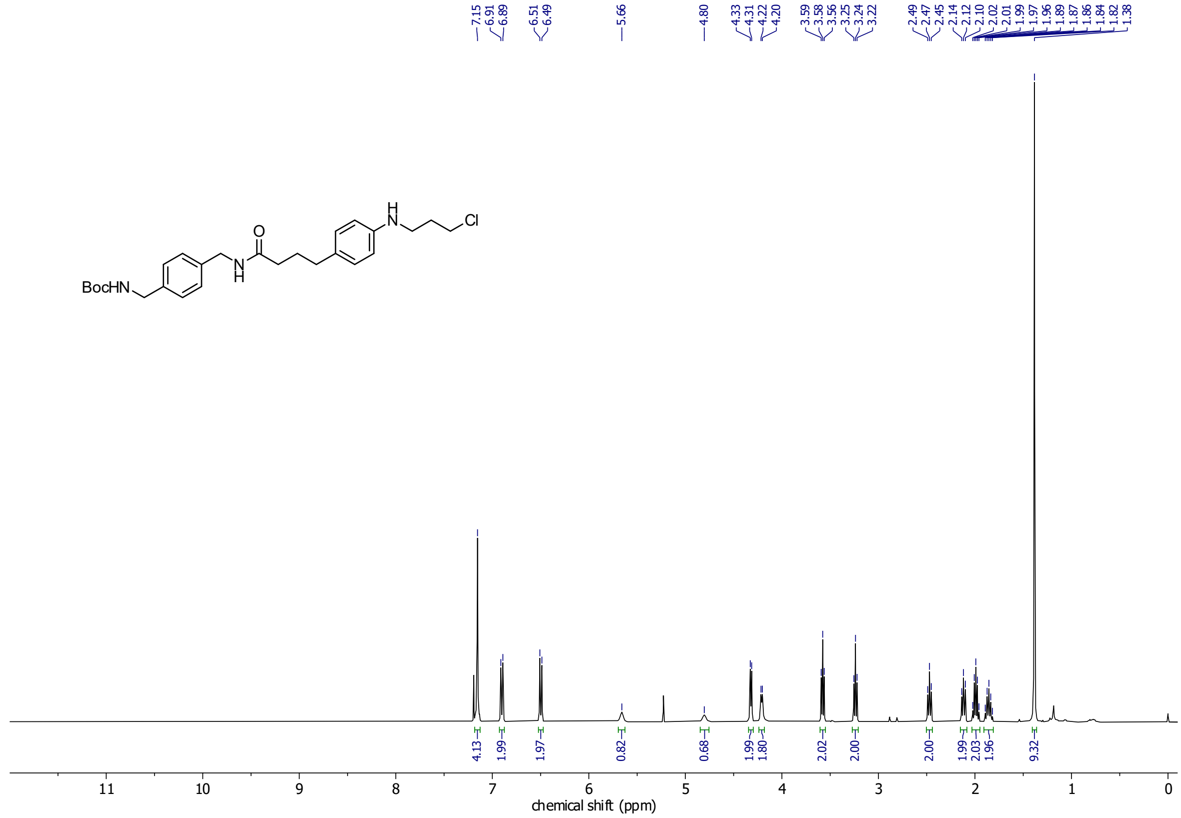

**$^{13}$C-NMR**

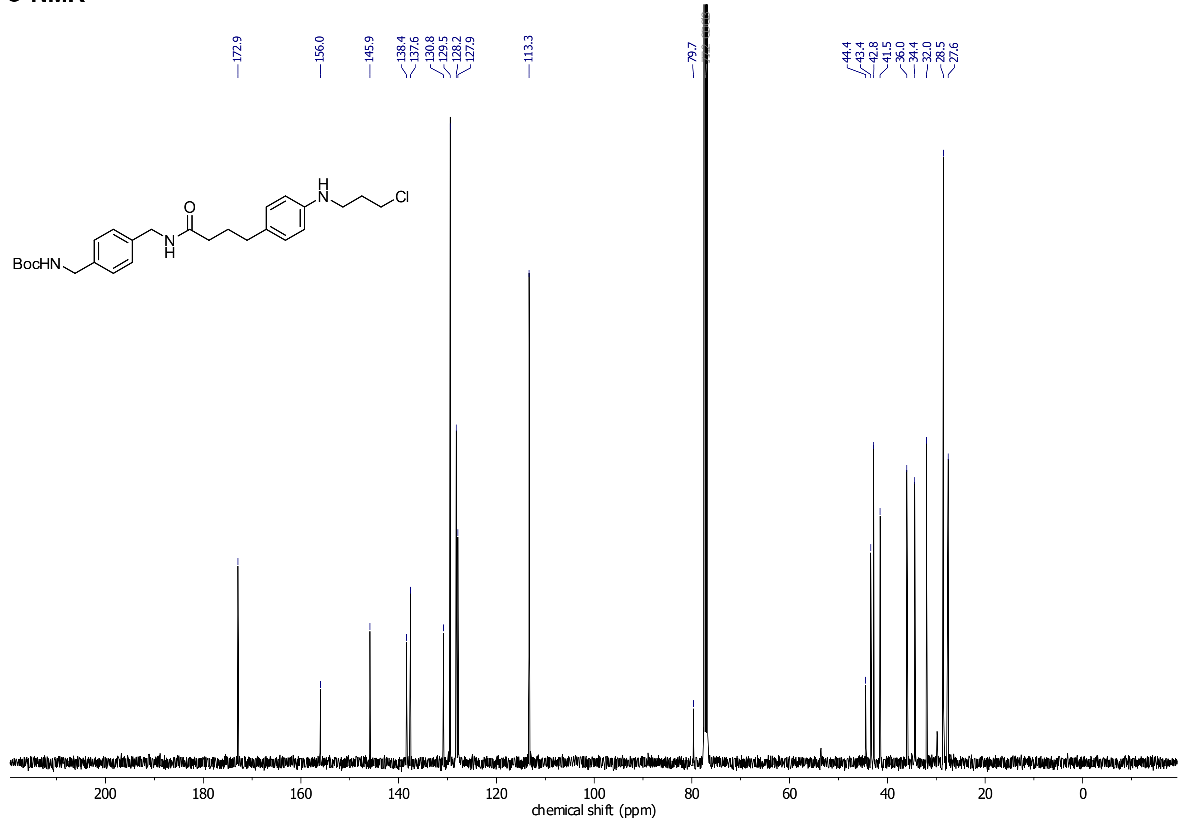

**HL3-Fluo**
**$^1$H-NMR**

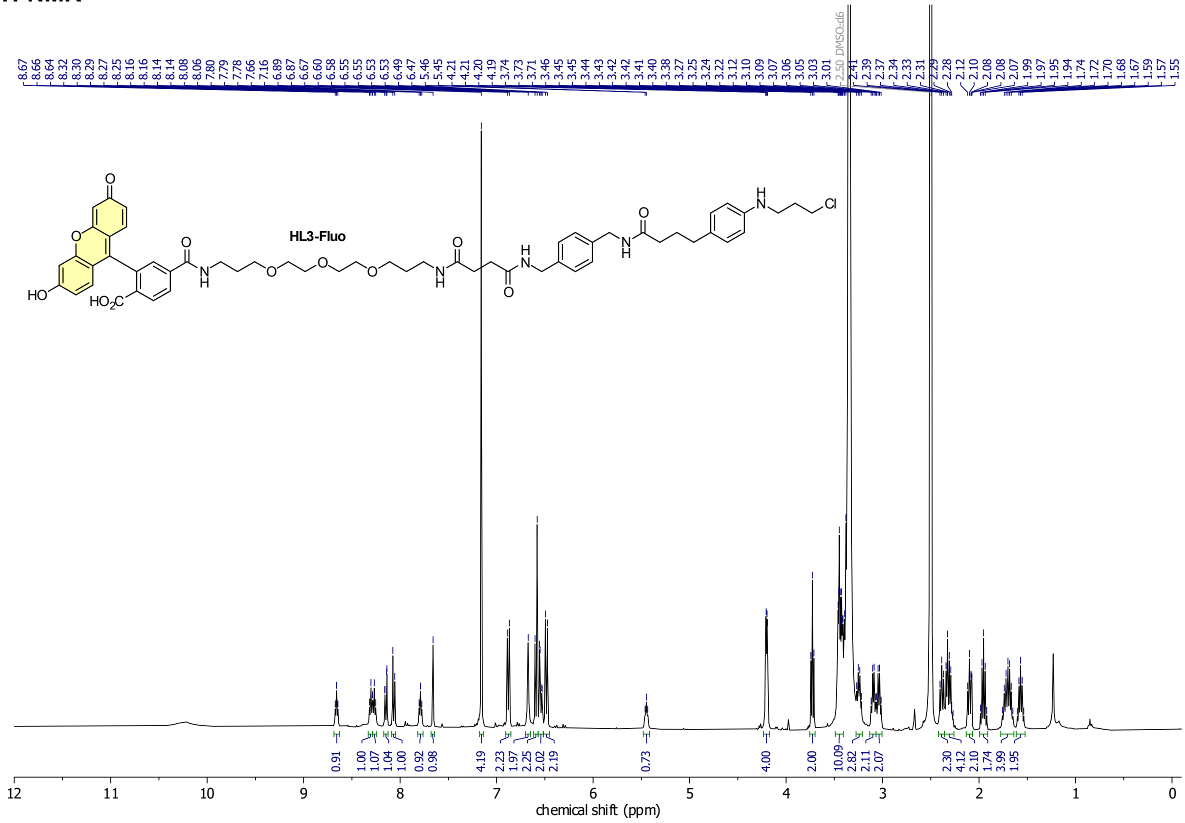

**Cou-HL3-Fluo**
**$^1$H-NMR**

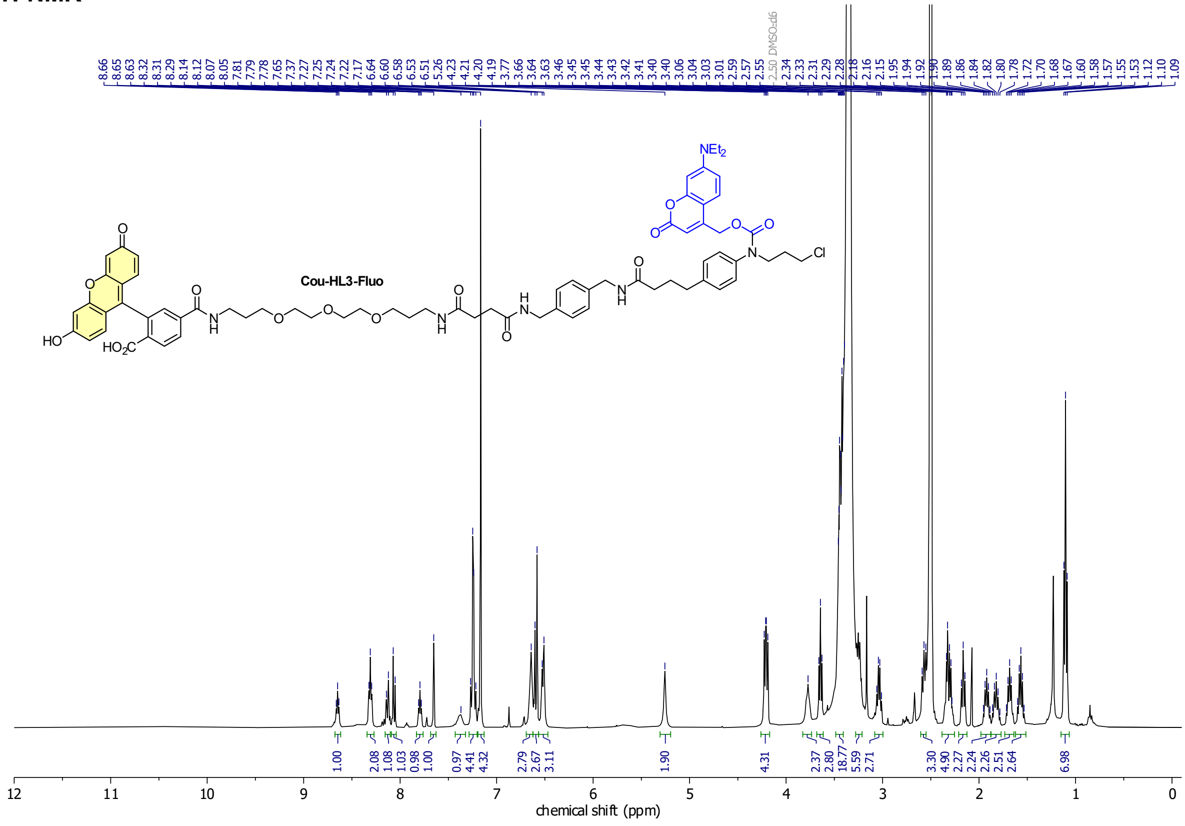

**Compound 64**

**$^{1}$H-NMR**

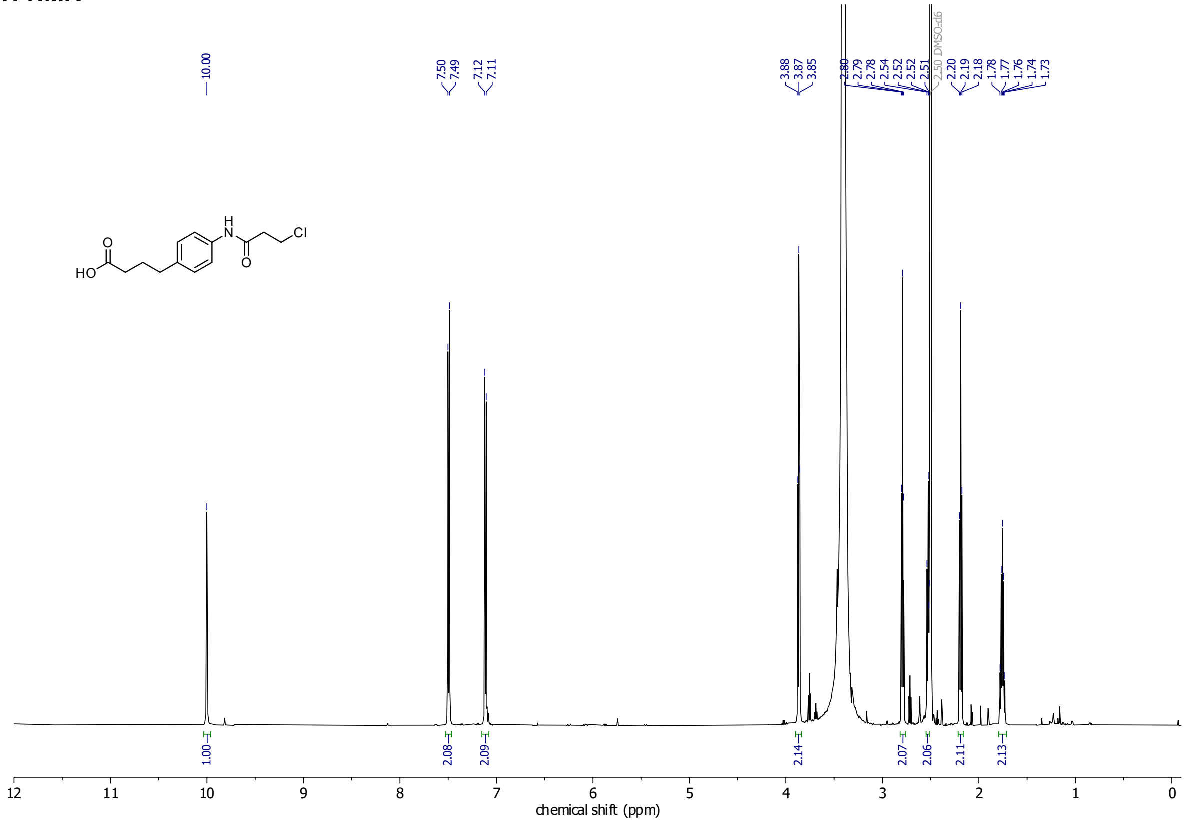

**$^{13}$C-NMR**

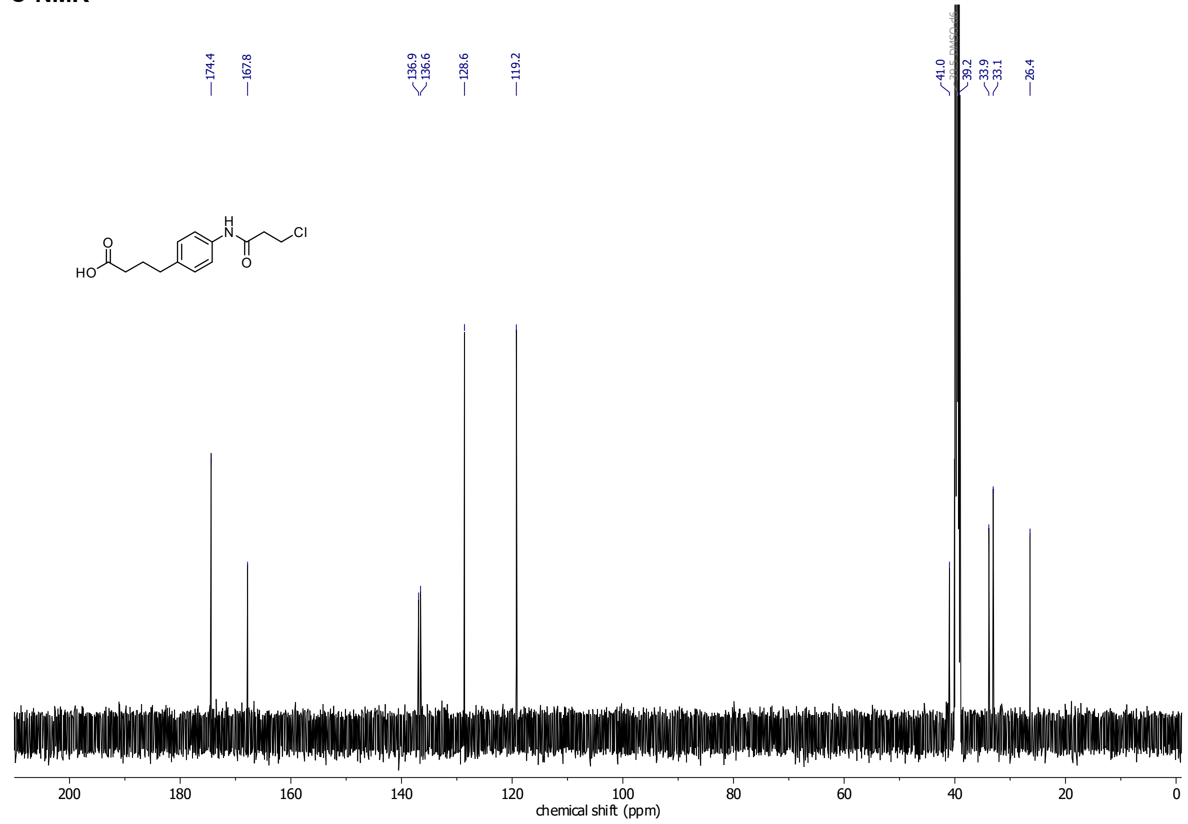

**HSQC**

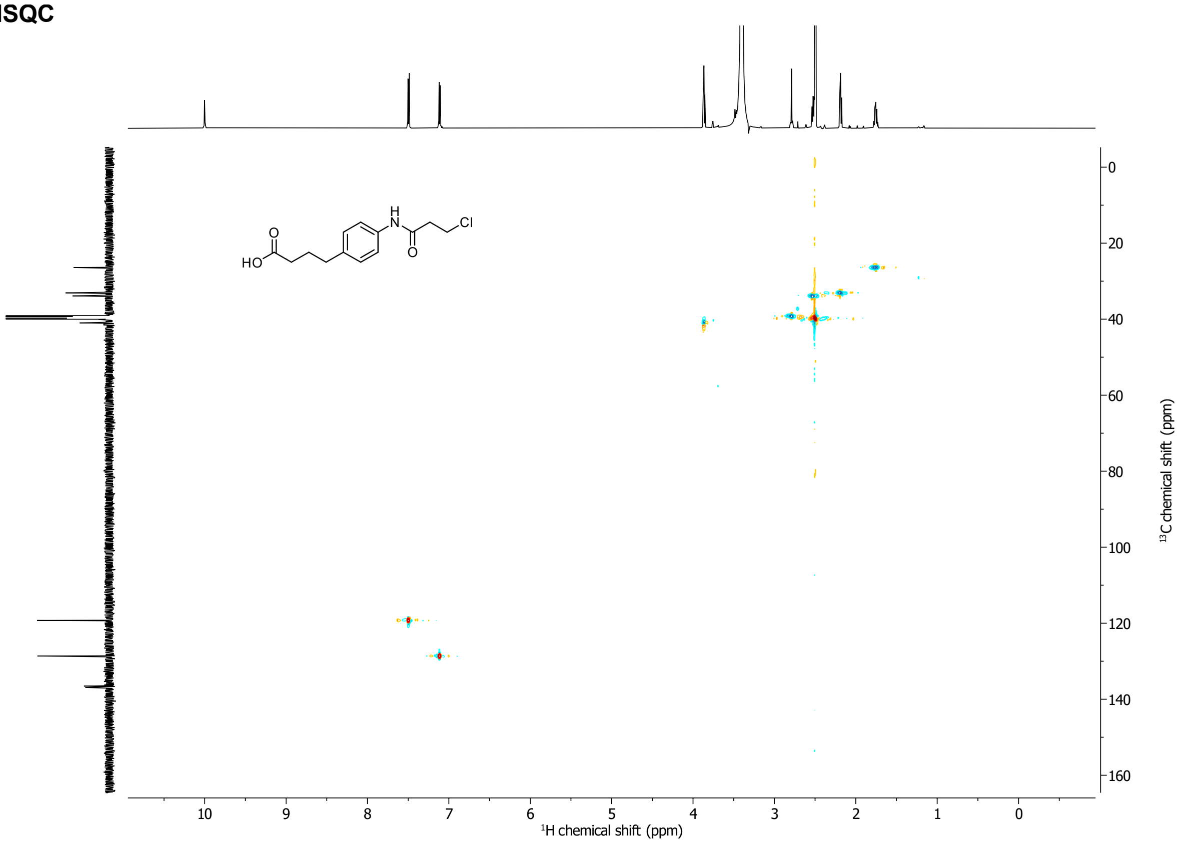

**Compound 65**

**¹H-NMR**

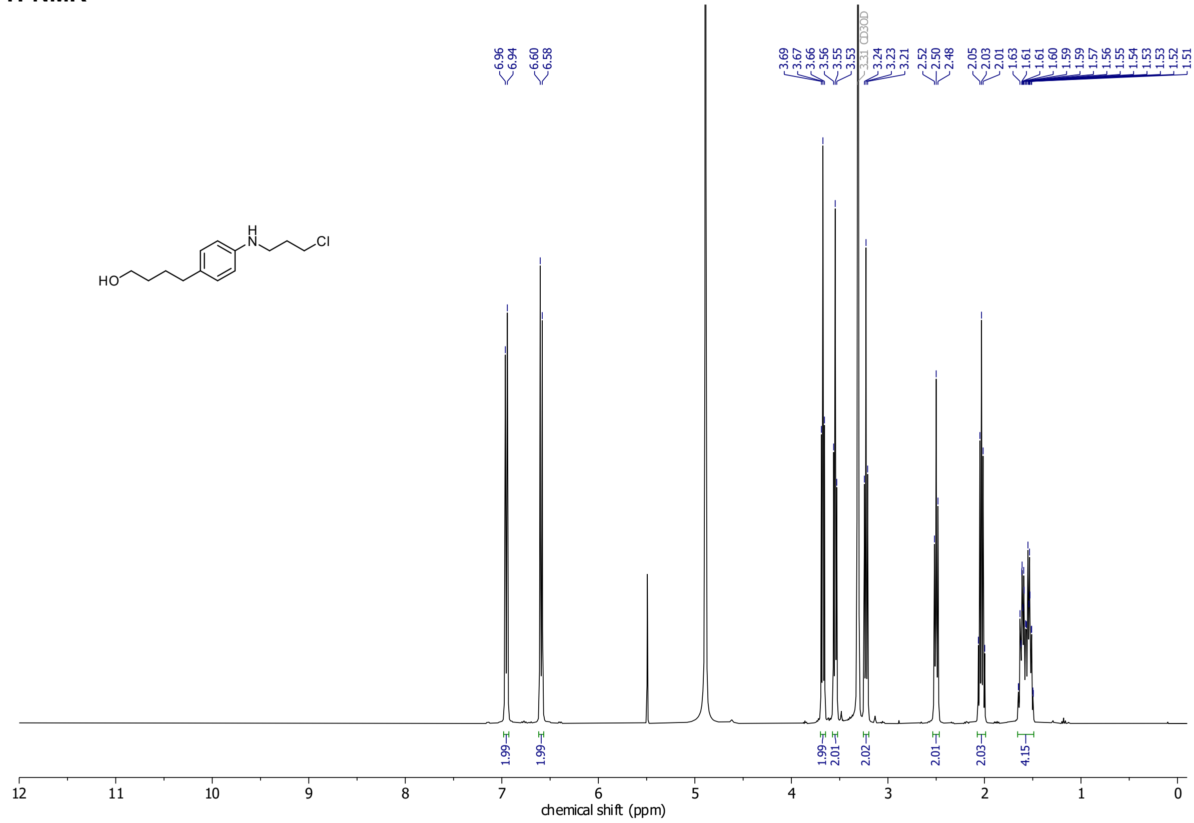

**¹³C-NMR**

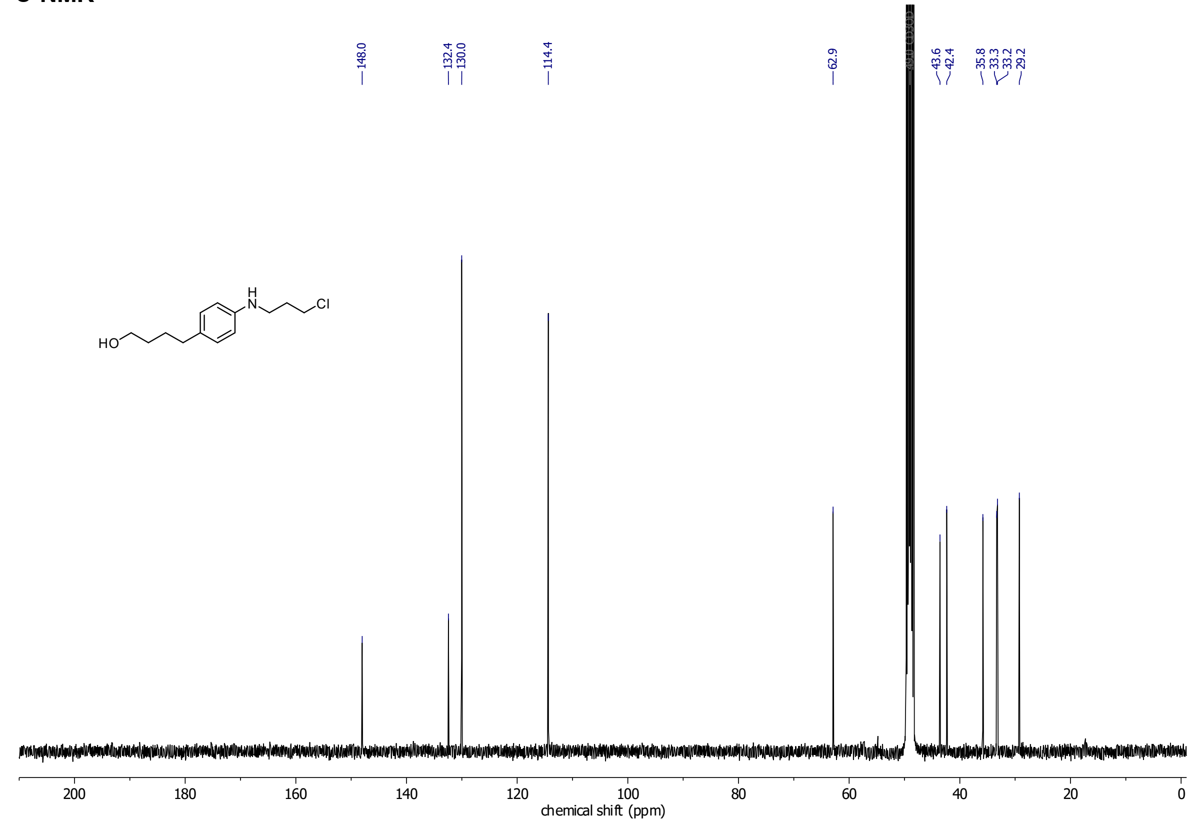

**Compound 66**
**$^{1}$H-NMR**

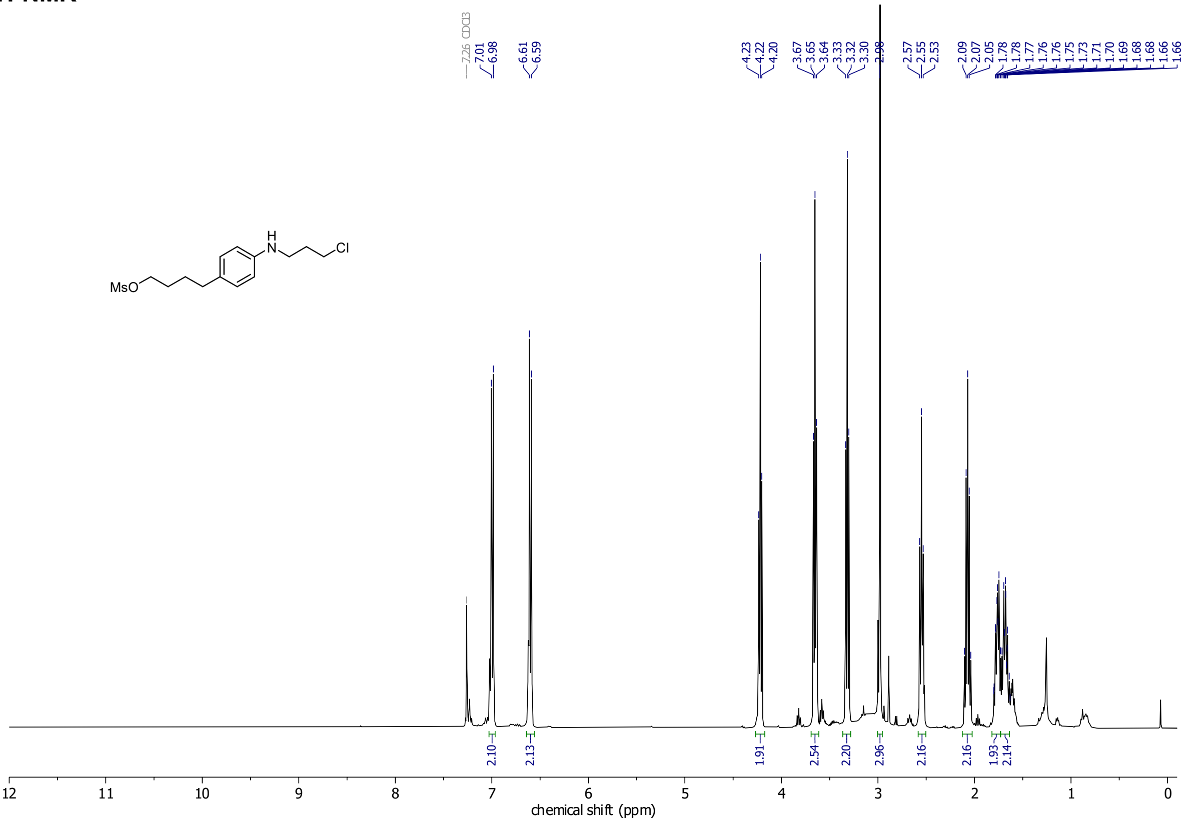

**Compound 67**

**$^1$H-NMR**

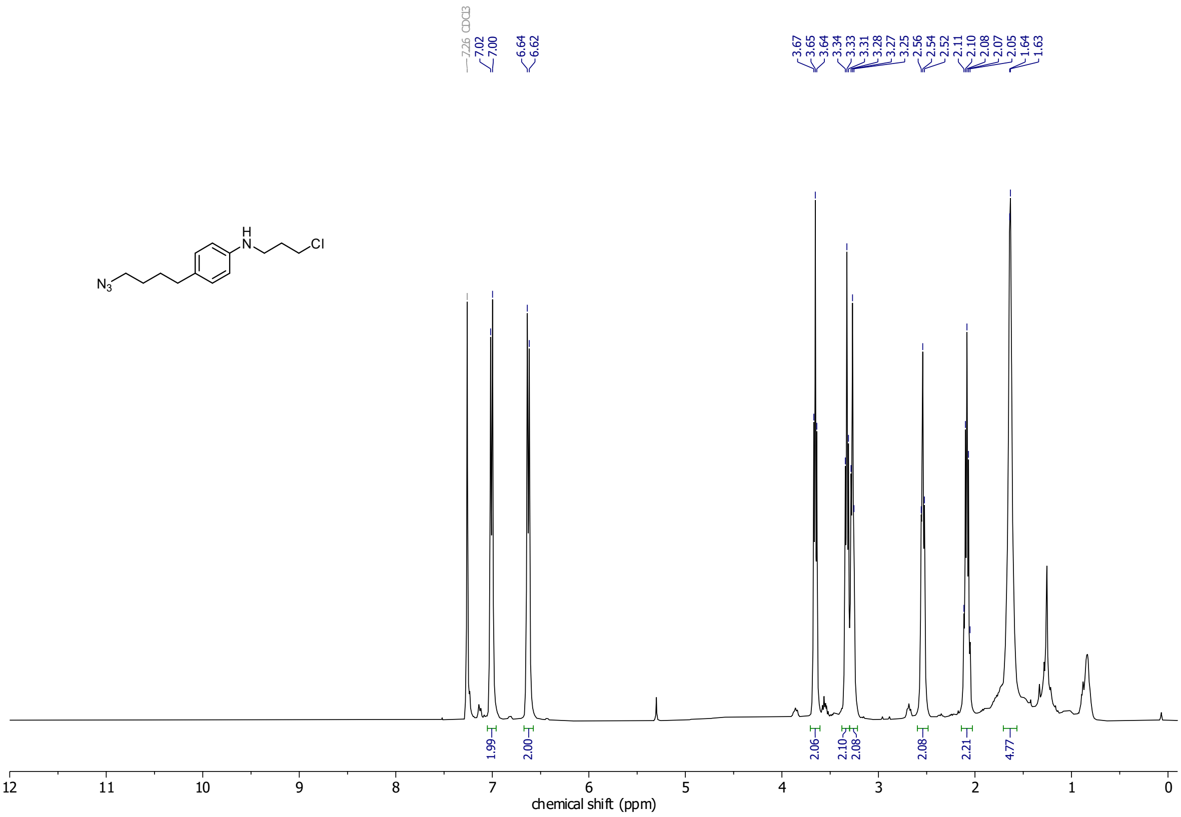

**$^{13}$C-NMR**

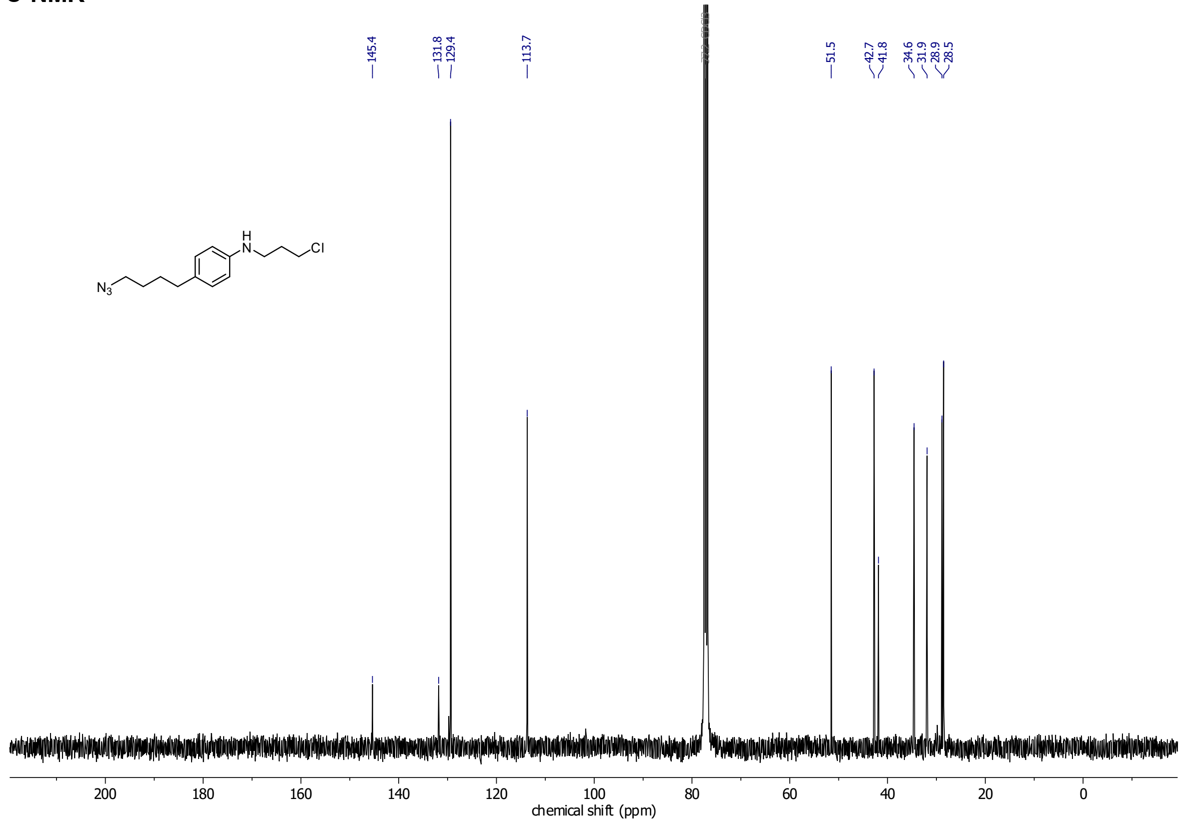

**CHalo-Fluo**

**$^1$H-NMR**

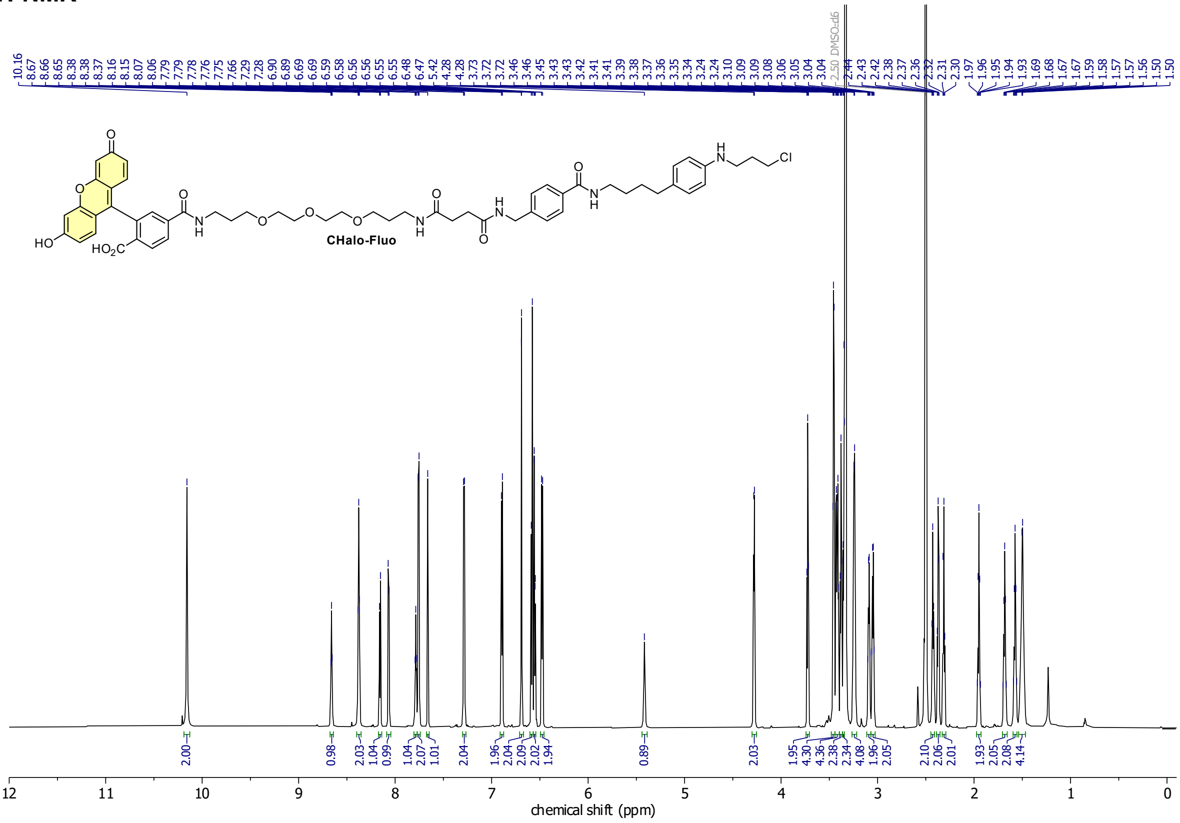

**$^{13}$C-NMR**

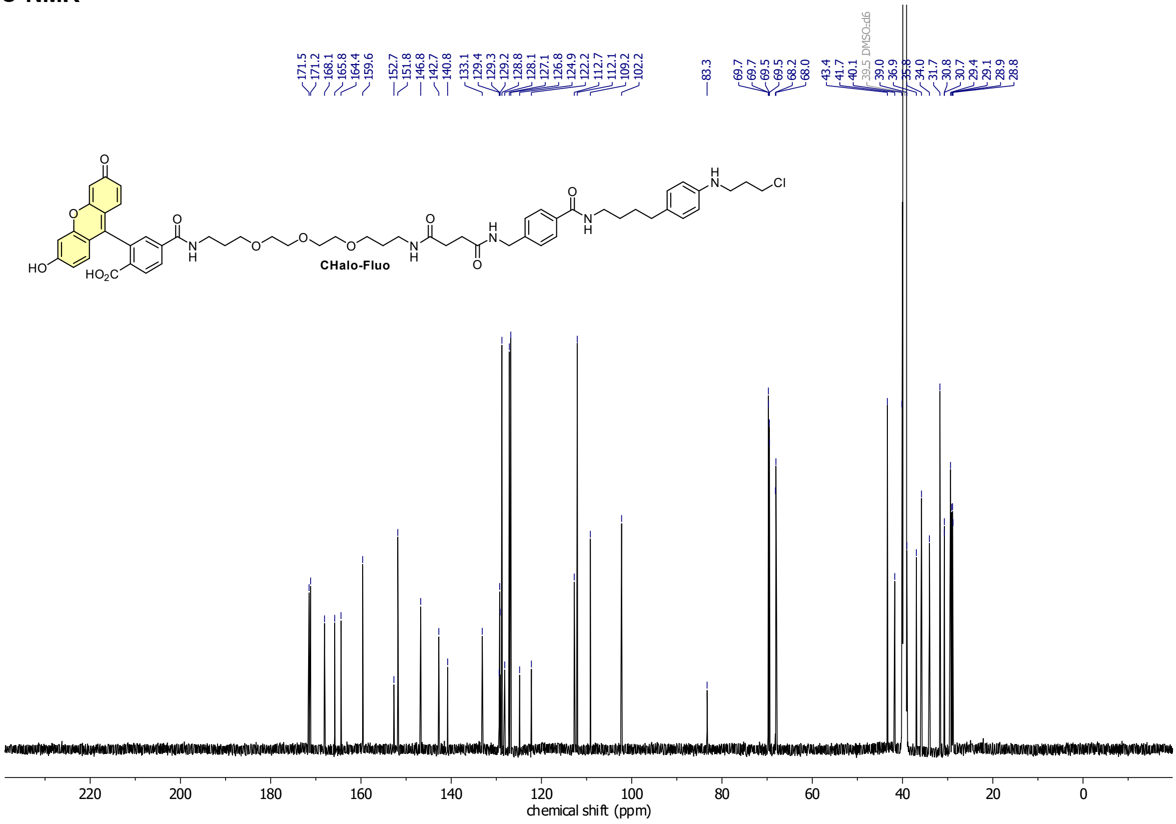

**Cou-CHalo-Fluo**

**$^1$H-NMR**

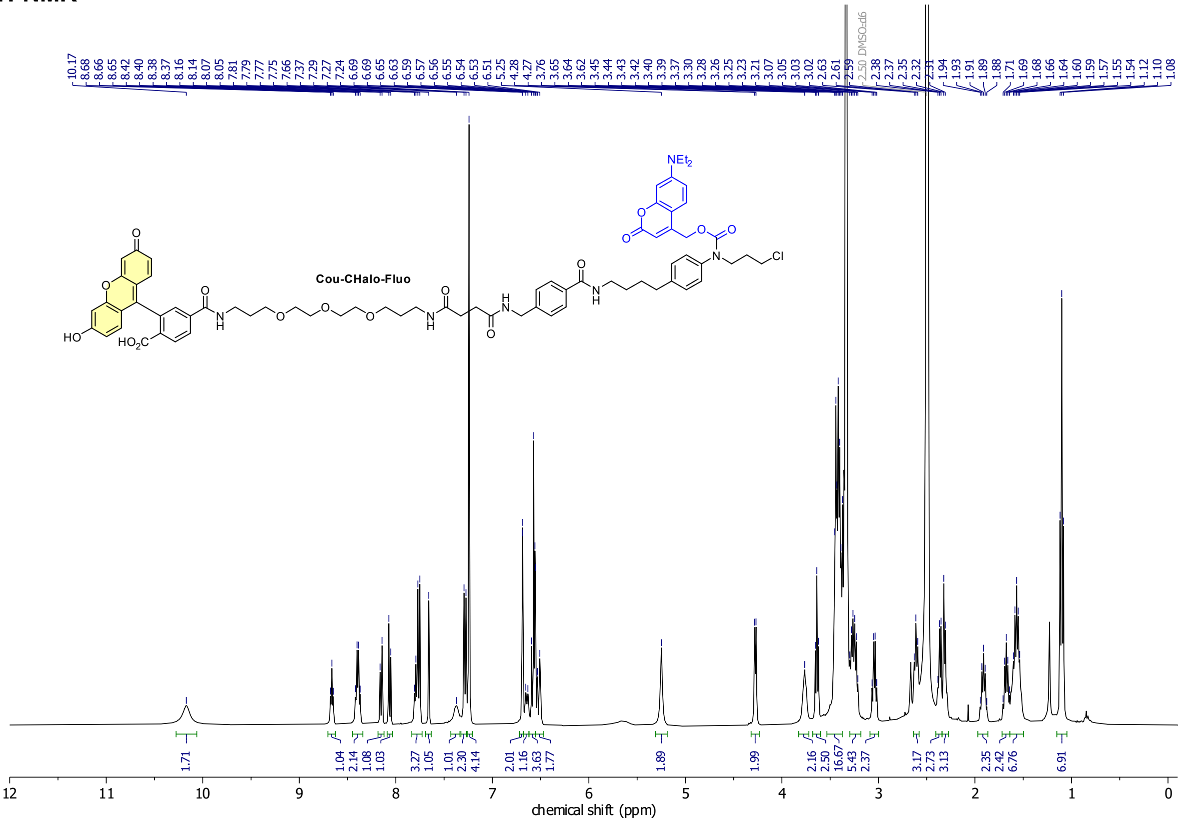
